# Discovery of positive and purifying selection in metagenomic time series of hypermutator microbial populations

**DOI:** 10.1101/2020.05.23.112508

**Authors:** Rohan Maddamsetti, Nkrumah A. Grant

## Abstract

A general method to infer both positive and purifying selection during the real-time evolution of hypermutator pathogens would be broadly useful. To this end, we introduce a simple test to infer mode of selection (STIMS) from metagenomic time series of evolving microbial populations. We test STIMS on metagenomic data generated by simulations of bacterial evolution, and on metagenomic data spanning 62,750 generations of Lenski’s long-term evolution experiment with *Escherichia coli* (LTEE). In both cases, STIMS recovers signals of positive and purifying selection on gold standard sets of genes. Using STIMS, we find strong evidence of ongoing positive selection on key regulators of the *E. coli* gene regulatory network, even in some hypermutator populations. STIMS also detects positive selection on regulatory genes in hypermutator populations of *Pseudomonas aeruginosa* that adapted to subinhibitory concentrations of colistin – an antibiotic of last resort – for just twenty-six days of laboratory evolution. Our results show that the fine-tuning of gene regulatory networks is a general mechanism for rapid and ongoing adaptation. The simplicity of STIMS, together with its intuitive visual interpretation, make it a useful test for positive and purifying selection in metagenomic data sets that track the evolution of hypermutator populations in real-time.

## INTRODUCTION

Organisms often evolve defects in DNA repair and recombination pathways that cause very high mutation rates. Hypermutability is readily observed in cancer [1–3] and in opportunistic pathogens attacking immunocompromised individuals [4]. Hypermutability is also associated with the evolution of antibiotic resistance, including multi-drug-resistant tuberculosis [5]. Hypermutability can increase the rate of beneficial mutations, but it can also obscure the genomic basis of adaptation [6], because vast numbers of nearly-neutral mutations hitchhike to high frequencies with the beneficial mutations that drive the selection dynamics. In this regime, called “emergent neutrality” [7], it is challenging to identify selection on particular genes, unless those genes are under very strong selection [8, 9]. As such, there is a need for methods that can resolve positive and purifying selection in hypermutator populations, due to their importance in basic research, genetic engineering, and medicine. To address this research gap, we present a simple test to infer mode of selection (STIMS) from metagenomic time series of large asexual hypermutator populations, and use this method to study positive and purifying selection in bacterial populations that evolved hypermutability during long-term experimental evolution.

We developed STIMS in the context of Lenski’s long-term evolution experiment with *Escherichia coli* (LTEE) [10, 11]. The LTEE has become an important test bed for many fundamental questions in evolutionary biology, due to its simplicity (daily serial transfer of twelve populations) and comprehensive record of frozen bacterial samples. Previous studies have used the LTEE as a model system to study the tempo and mode of both genomic [12–16] and phenotypic evolution [10, 17–22]. Many of those previous studies focused on the evolutionary dynamics underlying adaptive evolution. The evolution of hypermutability in six of the LTEE populations has obscured the genomic signatures of adaptation [6, 12, 15, 16, 23], again due to emergent neutrality. STIMS gains statistical power to infer selection despite emergent neutrality, by aggregating mutations within a focal set of genes, counting their occurrence, then comparing the counts to a background distribution of the number of mutations per gene set.

This work expands on previous observations suggesting the action of purifying selection on hypermutator populations of the LTEE [21]. In that work, we found that aerobic-specific genes were depleted in mutations compared to randomized sets of genes in three hypermutator populations: Ara−2, Ara+3, and Ara+6. That finding suggests that aerobic-specific genes may have transitioned from positive selection to purifying selection in those LTEE populations [21]. Our work also complements recent findings of purifying selection affecting protein evolution [24] and protein-protein-interaction network evolution in the LTEE [25].

Here, we demonstrate that STIMS recovers signals of positive and purifying selection in evolutionary simulations of haploid populations under a Wright-Fisher model, and show that STIMS recovers signals of positive and purifying selection on gold-standard sets of genes with *a priori* evidence of those selection pressures in the LTEE. We then use STIMS to examine the *tempo* of molecular evolution in modules of co-regulated genes [26–28], and to generate testable hypotheses about the *mode* (i.e. purifying or positive selection) of evolution on those gene modules in the LTEE. Finally, we use STIMS to test for positive selection on regulatory genes in a 26-day evolution experiment involving adaptation of hypermutator *Pseudomonas aeruginosa* to subinhibitory concentrations of colistin, an antibiotic of last resort. These results indicate that STIMS is a useful and general test for selection, even over relatively short timescales of molecular evolution.

## RESULTS

### Simulations demonstrate the conditions in which STIMS is an effective test for positive and purifying selection

The key premise of STIMS is that the tempo of molecular evolution differs across different sets of genes, due to variation in the underlying selection pressures affecting those sets of genes. As input, STIMS requires a list of mutations that have been observed in metagenomic sequencing of a haploid population, sampled over time (Figure 1). STIMS then calculates a simple summary statistic of the evolutionary dynamics: the number of observed mutations in a query gene set, normalized by total gene length. A *p-*value is then calculated using the non-parametric bootstrap, which is simply a count of the times that a random gene set accumulates more mutations than the query gene set (Figure 2). The same logic applies when measuring whether a gene set is depleted of mutations: in this case, STIMS counts the number of times that a random selection of genes accumulates fewer mutations than the query gene set. Importantly, the bootstrapped background distribution does *not*, in general, correspond to a neutral null model of evolution: genome-wide positive and purifying selection may affect the background distribution. Accordingly, STIMS is interpreted in relation to the overall tempo at which mutations are observed across the genome in a given population, which is driven by population-genetic parameters such as mutation rate, population size, and the genomic distribution of mutation fitness effects (DFE).

**Figure 1.**
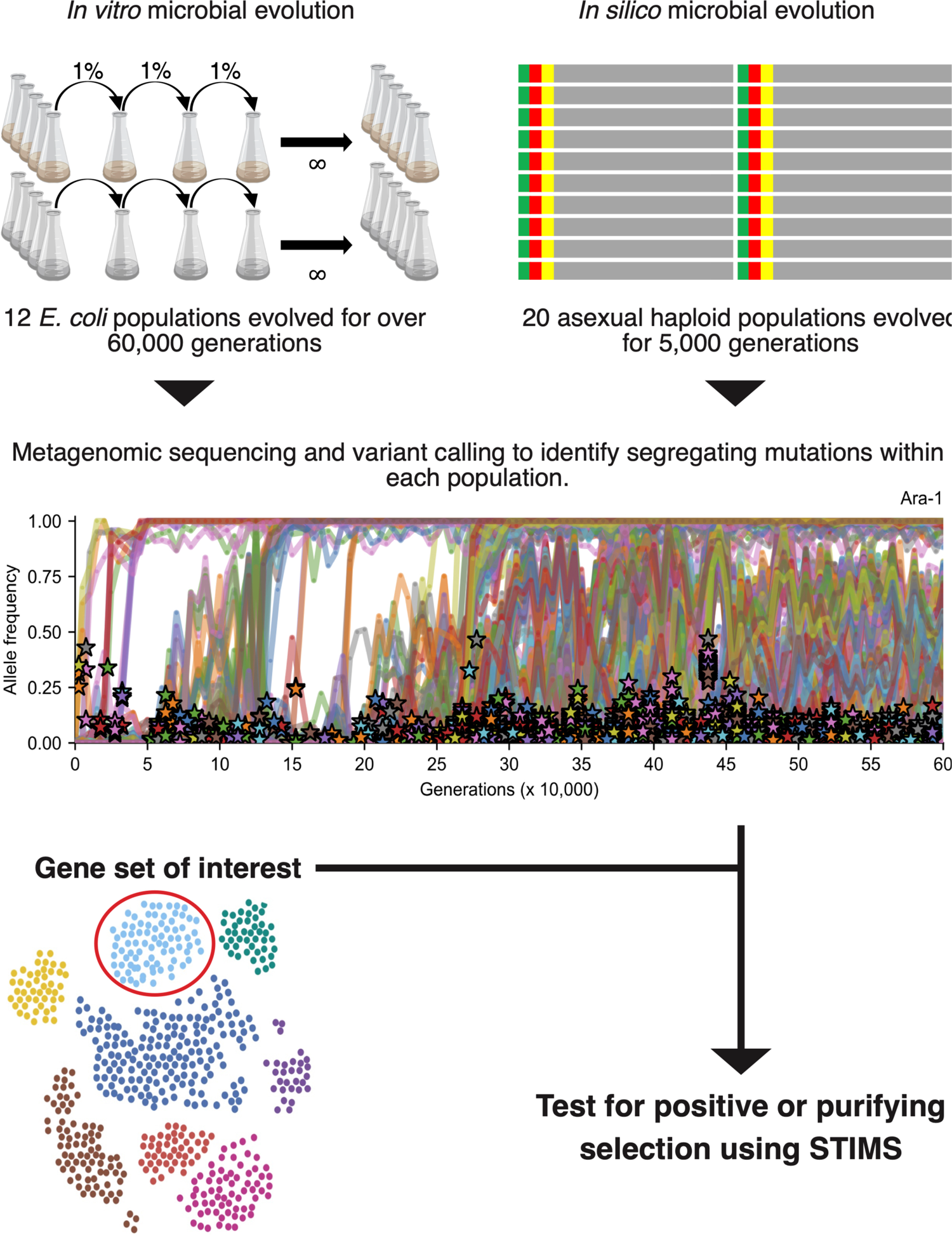
Data analysis workflow. 12 *E. coli* populations have evolved for more than 60,000 generations in Lenski’s long-term evolution experiment (LTEE). We reanalyzed metagenomic time series of the LTEE, described in Good et al. (2017). We also evolved 20 asexual haploid populations *in silico* for 5,000 generations. We identified variants segregating during the simulations to model metagenomic sequencing and variant calling. We designed a simple test to infer mode of selection (STIMS) to test pre-defined query gene sets for positive or purifying selection in the metagenomic time series. See the Materials and Methods for further details.

**Figure 2.**
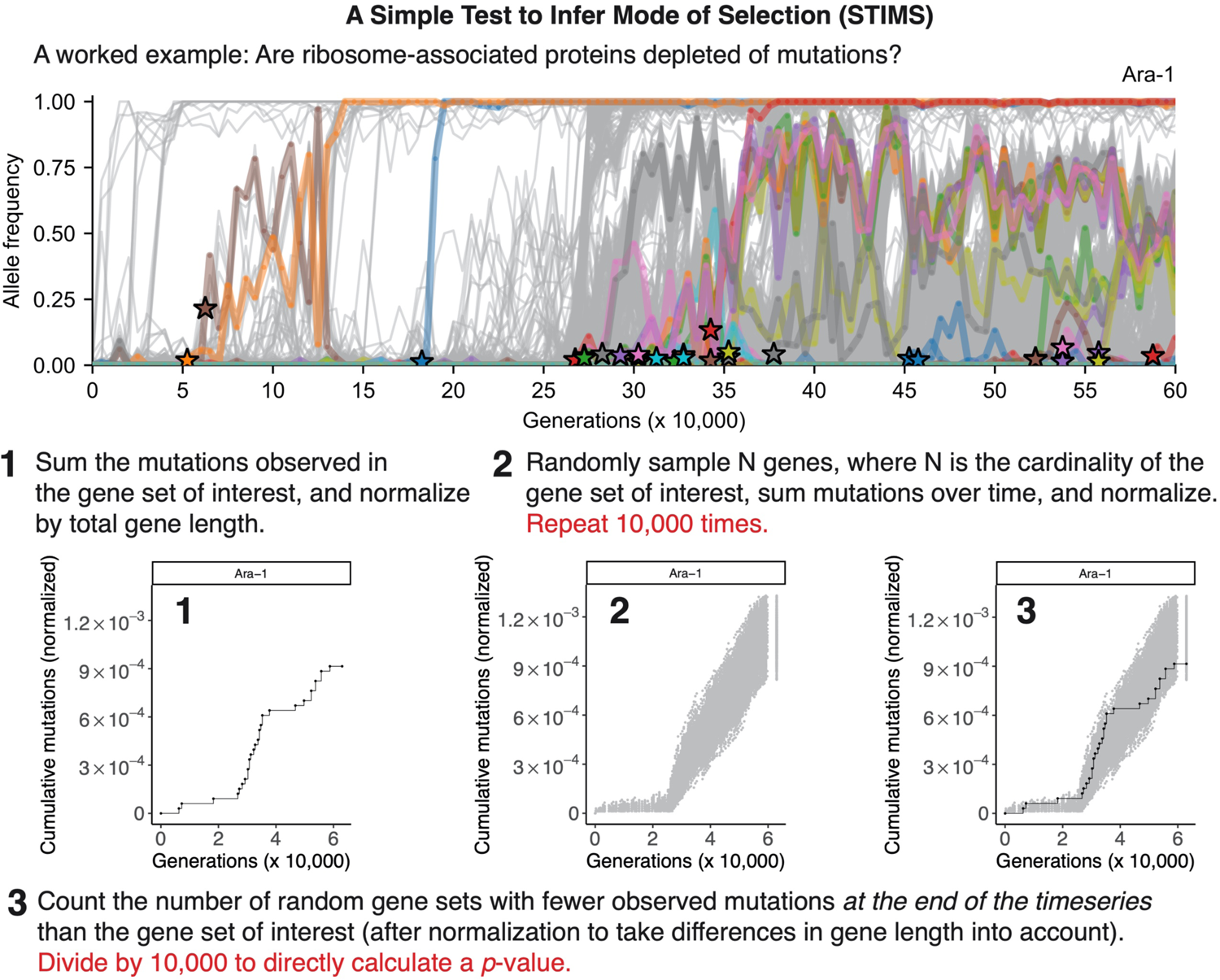
The STIMS algorithm. When examining the evolutionary dynamics of a gene set of interest, one often wants to know whether the set is enriched or depleted of observed mutations. To test for depletion, we use the non-parametric bootstrap to estimate the fraction of randomized gene sets that have fewer mutations than the query gene set. For example, if 11% of randomized gene sets have fewer mutations than the query gene set, then the query gene set is not depleted of mutations (one-tailed *p*-value = 0.11). A similar derivation holds for tests of enrichment. This worked example uses the allele frequency dynamics of ribosome-associated proteins in the Ara−1 population of Lenski’s long term evolution experiment with *Escherichia coli* (LTEE) to illustrate how the STIMS algorithm works. This population evolved a hypermutator phenotype around the 26,250-generation time point.

For this reason, we evolved haploid populations *in silico* using a Wright-Fisher model to examine the conditions under which STIMS can detect positive and purifying selection in idealized populations, in which the strength of selection on various regions of the genome is known *a priori*. We ran 10 replicate simulations of nonmutator haploid populations and 10 replicate simulations of hypermutator haploid populations, using large population sizes, mutation rates parameterized from LTEE measurements, and an idealized genome and DFE based on *E. coli* (Materials and Methods). The results of these simulations, and our analysis of these simulations using STIMS, are shown in Figure 3 for the nonmutator case and in Figure 4 for the hypermutator case.

**Figure 3.**
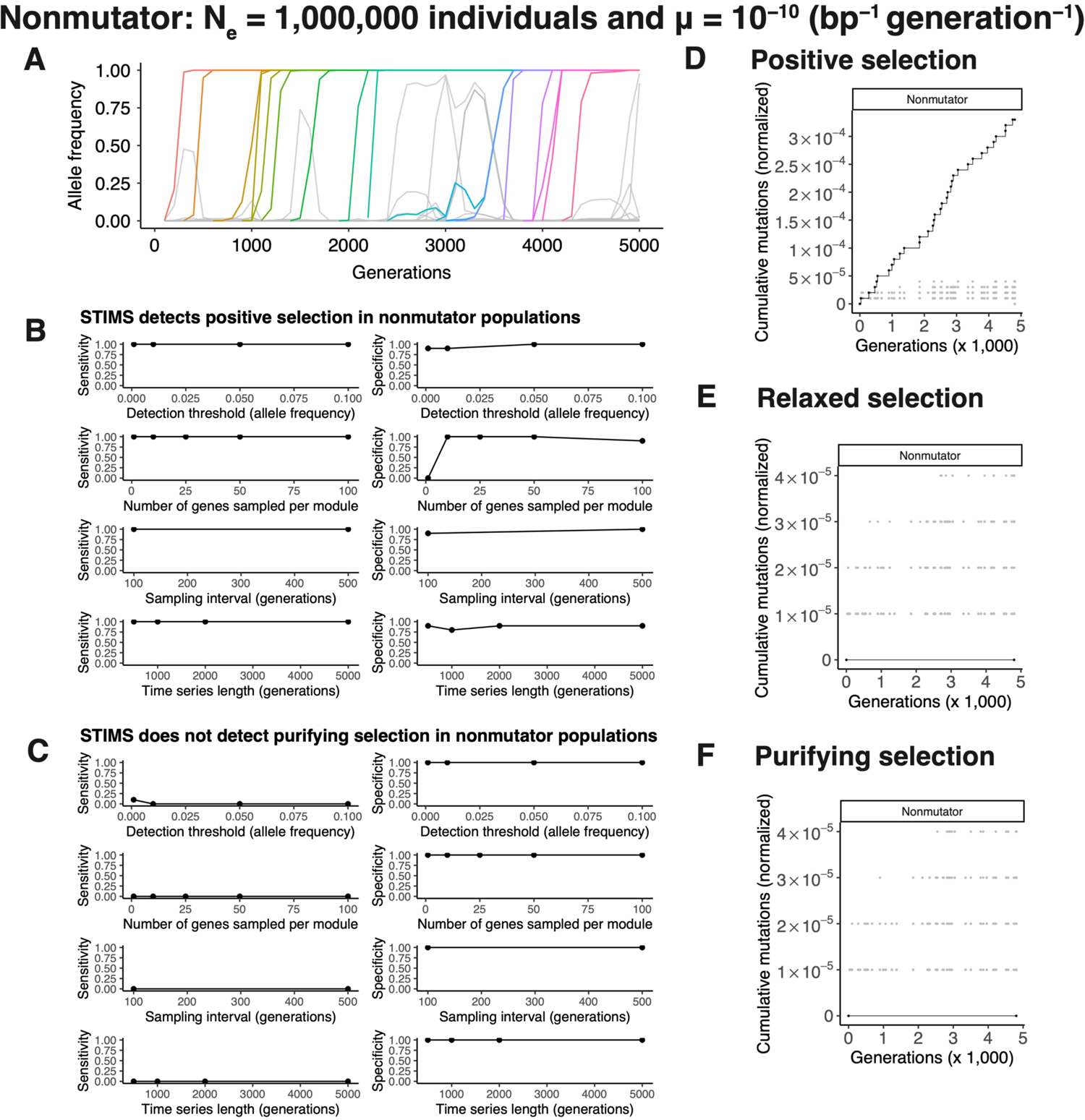
*In silico* evolution experiments demonstrate that STIMS can detect positive but not purifying selection in nonmutator populations. Bacterial evolution was simulated using a Wright-Fisher model. Each simulation lasted 5,000 generations. The effective population size was set to 1 million individuals. Nonmutator bacteria were modeled using a mutation rate of 10^−10^ per base-pair per generation. Each individual contains an idealized genome of 4,000 genes, each 1,000 base-pairs in length, for a genome of 4 million base-pairs. 100 genes are targets of positive selection, 100 genes are evolving completely neutrally (relaxed selection), and 100 genes are evolving under purifying selection. The remaining 3,700 genes evolve nearly neutrally, based on published estimates of the genomic distribution of mutation fitness effects in *Escherichia coli*. See the Materials and Methods for further details. A) Typical allele frequency dynamics of a nonmutator population, showing selective sweeps and clonal interference. B) STIMS shows high sensitivity and specificity for detecting positive selection in nonmutator populations. C) STIMS is unable to detect purifying selection (low sensitivity), but shows no false positives (high specificity). D) Typical result of running STIMS on genes evolving under positive selection in the nonmutator simulations. E) Typical result of running STIMS on genes evolving neutrally in the nonmutator simulations. F) Typical result of running STIMS on genes evolving under purifying selection in the nonmutator simulations.

**Figure 4.**
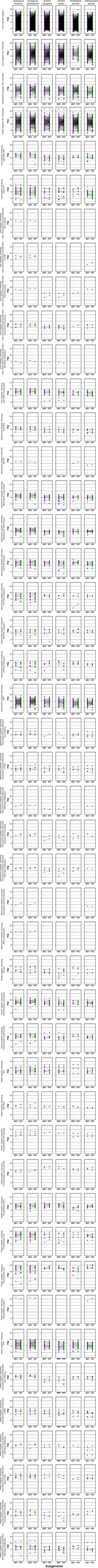
*In silico* evolution experiments demonstrate that STIMS can detect both positive and purifying selection in hypermutator populations. Bacterial evolution was simulated using a Wright-Fisher model. Each simulation lasted 5,000 generations. The effective population size was set to 1 million individuals. Hypermutator bacteria were modeled using a mutation rate of 10^−8^ per base-pair per generation. Each individual contains an idealized genome of 4,000 genes, each 1,000 base-pairs in length, for a genome of 4 million base-pairs. 100 genes are targets of positive selection, 100 genes are evolving completely neutrally (relaxed selection), and 100 genes are evolving under purifying selection. The remaining 3,700 genes evolve nearly neutrally, based on published estimates of the genomic distribution of mutation fitness effects in *Escherichia coli*. See the Materials and Methods for further details. A) Typical allele frequency dynamics of a hypermutator population, showing selective sweeps and clonal interference. B) STIMS shows high sensitivity and specificity for detecting positive selection in hypermutator populations. C) STIMS detects purifying selection with increasing sensitivity at lower allele frequency mutation detection thresholds, more genes sampled from the set of genes evolving under purifying selection, the length of the time series, and shorter intervals between metagenomic sampling, and shows high specificity across all tested parameters. D) Typical result of running STIMS on genes evolving under positive selection in the hypermutator simulations. E) Typical result of running STIMS on genes evolving neutrally in the hypermutator simulations. F) Typical result of running STIMS on genes evolving under purifying selection in the hypermutator simulations.

The dynamics of adaptation in our simulations are strikingly similar to the dynamics observed in the LTEE: both the nonmutator populations (Figure 3A) and the hypermutator populations (Figure 4A) show periodic selective sweeps, cohorts of mutations rising to fixation simultaneously, and clonal interference. These patterns are characteristic of the evolution of large asexual populations in which the supply of beneficial mutations does not limit adaptation [14, 16, 29]. Thus, our *in silico* simulations are representative of the biology we aim to describe.

We designed the genomes of our simulated haploid populations such that we knew *a priori* the mode of selection acting on particular genomic regions. Therefore, we used our simulated data to evaluate how well STIMS could detect selection given that it exists (sensitivity), and avoid false positives (specificity). Our simulations demonstrate that STIMS has both high sensitivity and specificity in detecting positive selection in both nonmutator and hypermutator populations (Figures 3B and 4B). Most mutations observed in the nonmutator populations are beneficial. So, the background distribution for the nonmutator populations reflects positive selection on beneficial driver mutations and the dynamics of passenger mutations hitchhiking with those driver mutations. In the nonmutator populations, not enough mutations are observed across the genome to see which genes are depleted of passenger mutations. Consequently, STIMS cannot detect purifying selection in this context. Nevertheless, STIMS is highly specific. It does not call any false positives in our tests for positive and purifying selection in the nonmutator populations (Figure 3C). Representative visualizations of STIMS when run on nonmutator populations are shown for genes evolving under positive selection in Figure 3D, for genes evolving completely neutrally in Figure 3E, and for genes evolving under purifying selection in Figure 3F. These visualizations show that for nonmutator populations, gene modules with trajectories below the background distribution could be evolving under purifying selection, or could be evolving completely neutrally due to relaxed selection.

This apparent difference from the background distribution is never statistically significant, because most random gene sets have no mutations at all, and these empty sets are not plotted. Gene modules with trajectories above the background distribution are evolving under strong positive selection.

In contrast to the nonmutator populations, our simulations show that STIMS has high sensitivity and specificity in detecting purifying selection in hypermutator populations (Figure 4C). In this case, mutation rates are high enough that passenger mutations arise densely across the genome, allowing for the detection of genes that are depleted of observed passenger mutations due to purifying selection. The sensitivity with which STIMS detects purifying selection increases when 1) mutations can be reliably detected at lower frequencies; 2) the number of genes in the gene set increases; 3) the interval between metagenomic sampling time is smaller; and 4) the length of the time series increases (Figure 4C). All of these experimental variables increase the resolution of the data and increase the ability of STIMS to detect purifying selection in hypermutator populations. Representative visualizations of STIMS when run on hypermutator populations are shown for genes evolving under positive selection in Figure 4D, for genes evolving completely neutrally in Figure 4E, and for genes evolving under purifying selection in Figure 4F.

These results show that STIMS is an effective test for selection, given sufficient data. However, the realized DFE for microbial populations evolving in the laboratory or in nature may deviate substantially from the idealized DFE in the simulations that we have presented. For this reason, we validated STIMS on published metagenomic data tracking the allele frequency dynamics in the LTEE, using three gold standard sets of genes with empirical evidence of relaxed, purifying, and positive selection in the LTEE. These positive controls provide hard evidence that STIMS is effective in practice.

### Control 1: Resolving relaxed selection in the LTEE

As a first positive control, we examined 63 genes that have empirical evidence for neutral fitness effects in REL606 (A. Couce, personal communication). Figure 5 shows the results of applying STIMS on these genes, which we suppose are evolving under relaxed selection in the LTEE. In 5 out of 6 nonmutator populations (top two rows), the trajectory of this gene set falls within the null distribution. The remaining nonmutator population, Ara+5, has no mutations in this gene set, and so falls significantly below the null distribution (two-tailed bootstrap: Bonferroni-corrected *p* < 0.005). In 5 out of 6 hypermutator populations (bottom two rows), the trajectory of this gene set falls within the null distribution. The remaining hypermutator population, Ara+6 lies at the margin of the null distribution (two-tailed bootstrap: Bonferroni-corrected *p* = 0.06). Supplementary Figure S1 shows the result of applying STIMS on these genes, summed across all 12 LTEE populations. Overall, the global tempo of observed mutations in the nonmutator populations is best explained by the occurrence and fixation of beneficial mutations with small numbers of hitchhikers. By contrast, the global tempo of observed mutations in the hypermutator populations is best explained by large numbers of nearly neutral mutations hitchhiking to high frequency with a much smaller number of beneficial mutations [15].

**Figure 5.**
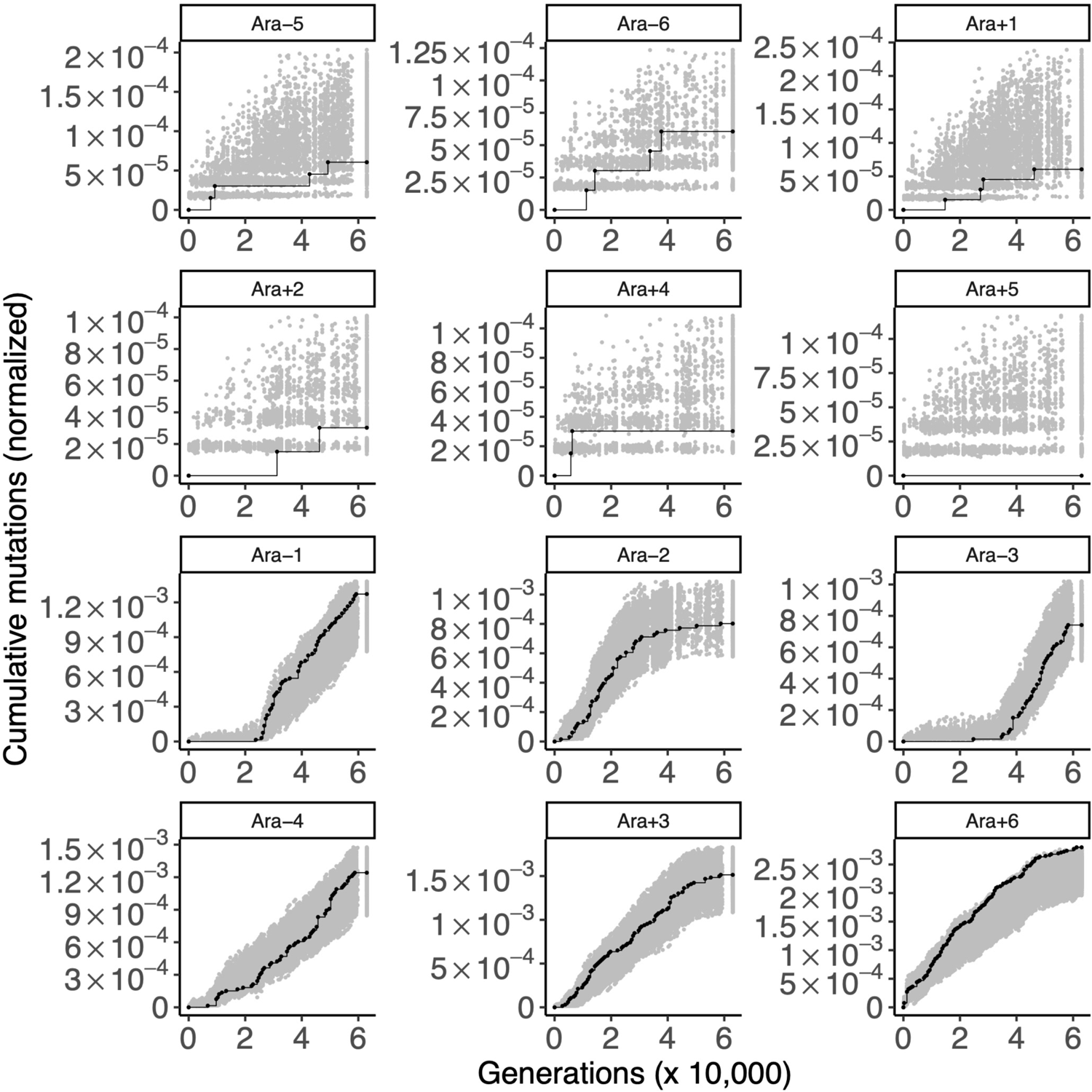
Gold-standard genes under relaxed selection. Each panel shows the cumulative number of mutations in the gold-standard neutral genes (black) in the 12 LTEE populations. For comparison, random sets of 57 genes were sampled 1,000 times, and the cumulative number of mutations in those random gene sets, normalized by gene length, were calculated. The middle 95% of this null distribution is shown in gray, in order to show a two-tailed statistical comparison of the cumulative number of mutations in the gold-standard neutral gene set to the null distribution at *α* = 0.05. The top six panels are populations with the ancestral mutation rate (nonmutator populations) and the bottom six panels are those that evolved elevated mutation rates (hypermutator populations).

### Control 2: Detecting purifying selection in the LTEE

As a second control, we examined genes that were experimentally shown to be essential or nearly essential in REL606 under LTEE conditions [6]. Many of the genes under strongest positive selection in the LTEE are also essential genes [23]. Thus, we excluded 24 essential genes with evidence of parallel evolution in the LTEE (Materials and Methods). We hypothesized that the remaining 491 essential genes would show evidence of purifying selection in the hypermutator populations, such that those loci would be depleted in mutations in comparison to the null distribution (Figure 6 and Supplementary Figure S2). Indeed, these 491 loci show significant depletion of observed mutations in all 6 hypermutator populations (one-tailed bootstrap: *p* < 0.0001 for Ara−1; *p* < 0.01 for Ara−2; *p* < 0.01 for Ara−3; *p* < 0.005 for Ara−4; *p* < 0.02 for Ara+3; *p* < 0.0001 for Ara+6). Additionally, one of the six nonmutator populations, Ara+1 shows a marginally significant depletion of mutations at these loci (one-tailed bootstrap: *p* < 0.05). Ara+1 evolved a insertion-sequence hypermutator phenotype early in its evolutionary history [30–32], so it is plausible that signatures of purifying selection can be seen in Ara+1 [25], even though it did not evolve an elevated point-mutation rate [31].

**Figure 6.**
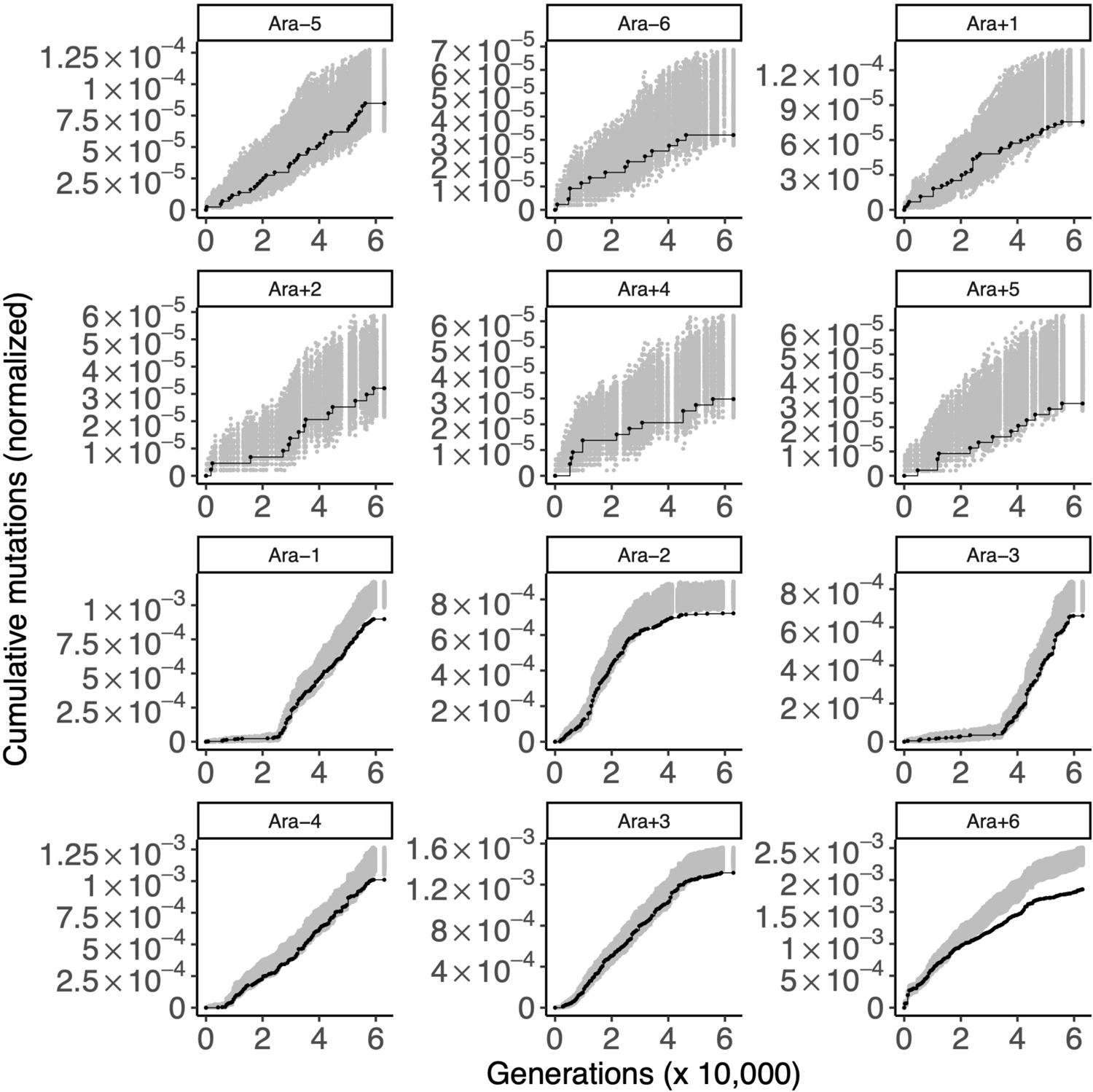
Gold-standard genes under purifying selection. Each panel shows the cumulative number of mutations in the gold standard set of genes under purifying selection (black) in the 12 LTEE populations. For comparison, random sets of 490 genes were sampled 1,000 times, and the cumulative number of mutations in those random gene sets, normalized by gene length, were calculated. The middle 95% of this null distribution is shown in gray. The top six panels are the nonmutator populations and the bottom six panels are the hypermutator populations.

Despite the global signal of purifying selection on these 491 essential genes, it is not clear which specific ones are under purifying selection. The best candidates are the ones that have no mutations whatsoever in the LTEE, while the second-best are those that were only affected by synonymous mutations. We found 65 genes with no observed mutations in the metagenomics data, 18 of which are essential. 105 genes have only synonymous mutations, 30 of which are essential. We examined 491 essential genes, out of 3948 genes in the genome that passed our filters (Materials and Methods). Based on these numbers, there is a significant association between essentiality and having no observed mutations (Fisher’s exact test: *p* = 0.0007) and between essentiality and only having synonymous mutations (Fisher’s exact test: *p* < 10^−5^). The first set of candidates for purifying selection in the LTEE are reported in Supplementary Table S1, and the second set of candidates are reported in Supplementary Table S2. As one would expect, many of these genes encode proteins that catalyze key biological reactions, such as ATP synthase and ribosomal proteins.

### Control 3: Verifying positive selection in the LTEE

As a final control, we examined genes that have previously been shown to be evolving under positive selection in the LTEE. We examined the genes with the strongest signal of parallel evolution in genomes sampled over the first 50,000 generations of the LTEE [15].

We first considered the 50 genes with the most parallelism in the nonmutator populations (Figure 7 and Supplementary Figure S3). STIMS recovered the expected signal of positive selection on these genes in the nonmutator populations (one-tailed bootstrap: *p* < 0.0001 in all cases). STIMS also found an overall signal of positive selection on these genes in the hypermutator populations (one-tailed bootstrap: *p* < 0.05 in all hypermutator populations).

**Figure 7.**
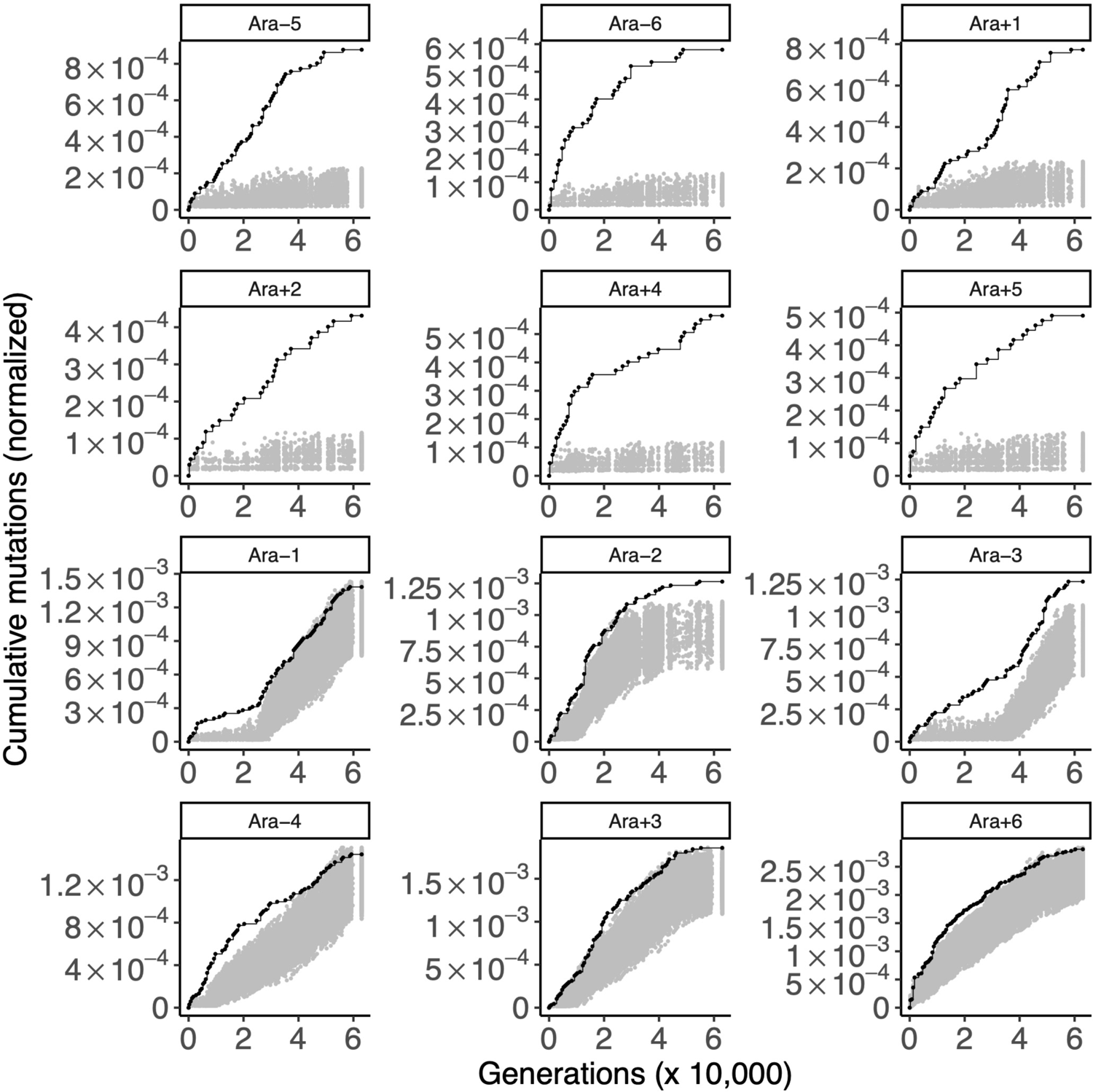
Gold-standard genes under positive selection. Each panel shows the cumulative number of mutations in the gold standard set of genes under positive selection (black) in the 12 LTEE populations. These genes are the 50 genes with the strongest signal of parallel evolution in clones isolated from the six nonmutator populations, as scored by Tenaillon et al. (2016). For comparison, random sets of 50 genes were sampled 1,000 times, and the cumulative number of mutations in those random gene sets, normalized by gene length, were calculated. The middle 95% of this null distribution is shown in gray. The top six panels are the nonmutator populations and the bottom six panels are the hypermutator populations.

In the same fashion, we examined the genes with the strongest signal of parallel evolution in the hypermutator genomes (Supplementary Figure S4 and Supplementary Figure S5). The nonmutator populations accumulate mutations in this gene set at a rate similar to background. As expected, 5 out 6 hypermutator populations show a strong excess of mutations above background (one-tailed bootstrap: *p* < 0.02 in Ara−1, Ara−2, Ara−4, Ara+3, Ara+6). In the remaining population, Ara−3, mutations in this gene set accumulate at the background rate. It is unclear why this population is an outlier, but this finding may be related to the evolution of a citrate metabolic innovation that modified the ecology of this population [33].

### Comparison of STIMS to a test for selection based on the Poisson distribution

We also benchmarked STIMS against a statistical test for selection that uses the Poisson distribution to model the expected distribution of mutations per genomic site per unit time under neutral evolution [16, 34]. In brief, STIMS trades off sensitivity for higher specificity (lower false positive rate) in comparison to the Poisson method, and also has the advantage of showing how the tempo of molecular evolution varies over time in a gene set of interest. These findings, and additional considerations, are described in detail in the Supplementary Text.

#### Application to module decompositions of the E. coli genome

Our control experiments demonstrate that STIMS is capable of recovering signals of purifying and positive selection. In this section, we use STIMS to examine how different modules of genes in the *E. coli* genome are evolving in the LTEE. Our goal is to develop testable hypotheses for which genetic modules and biochemical pathways underlie the ongoing fitness gains observed in the LTEE [35]. We examined three different module decompositions of the *E. coli* genome. The first partitioned the *E. coli* proteome into sectors based on quantitative mass spectrometry [26]; the second reported sets of genes (“eigengenes”) whose expression best predicted *E. coli* growth rates [28]; while the third partitioned the gene regulatory network into independent components based on gene expression [27].

#### Two proteome sectors show evidence of purifying selection

We examined proteome sectors that were identified by quantitative mass spectroscopy during carbon-, nitrogen-, and ribosome-limiting growth conditions [26]. Supplementary Figure S6 shows our findings. One proteome sector was significantly depleted in mutations in Ara−1, Ara+3, and Ara+6, suggesting purifying selection, and also shows evidence of positive selection in Ara+5. This one, called the U-sector (green), contains genes that were not upregulated by any of the growth-limitation treatments in [26]. Rather, the expression of the proteins in the U-sector show a generic positive correlation with growth rate across all of the conditions tested by Hui et al. [26]. When STIMS is applied to these modules, considering mutations summed across *all* LTEE populations, we also see an overall depletion of mutations in proteins in the R-sector (yellow; Supplementary Figure S7). Proteins in the R-sector are upregulated under ribosome-limiting growth conditions, and are associated with translation [26].

#### No evidence of selection on *E. coli* eigengenes

Finally, we examined the sets of genes (called eigengenes), that were most predictive of *E. coli’s* growth rate (and presumably fitness under LTEE conditions), using different *E. coli* strains and growth environments [28]. No eigengenes showed any evidence of selection in the LTEE (Supplementary Figure S8 and S9).

#### Regulators of I-modulons are under positive selection

Sastry et al. (2019) used independent component analysis on the *E. coli* transcriptome, sampled across diverse conditions and strains, to infer 92 independent modules (called I-modulons) in the *E. coli* gene regulatory network (GRN). 61 of the 92 I-modulons correspond to known transcription factors and regulators in the GRN. We asked two questions: first, is there a difference between how the regulators and regulated genes in the I-modulons evolve in the LTEE? Second, are any I-modulons enriched with or depleted of mutations in the LTEE?

We find that transcriptional regulators of the I-modulons are under strong positive selection in the LTEE, while the genes within I-modulons are under much weaker selection (Figure 8). In all 12 populations, regulators of I-modulons are under stronger selection than the genes in the I-modulons themselves. I-modulon regulators are under very strong selection in all 6 nonmutator populations (one-tailed bootstrap: *p* < 0.001 in all nonmutator populations), and show evidence of ongoing positive selection in some hypermutator populations as well (one-tailed bootstrap: *p* = 0.03 in Ara−3, *p ≤* 0.0001 in Ara+3 and Ara+6). By contrast, the evolution of genes within I-modulons fits the null distribution in most populations, although Ara−4 and Ara+3 show evidence of purifying selection. When mutations across all LTEE populations are considered, genes regulated within I-modulons show an overall signal of purifying selection (Supplementary Figure S10). Together, these results indicate that the upper levels of the regulatory hierarchy of the *E. coli* GRN are under strong selection in the LTEE, while the effector genes at the lower levels of the gene regulatory hierarchy are under weaker selection.

**Figure 8.**
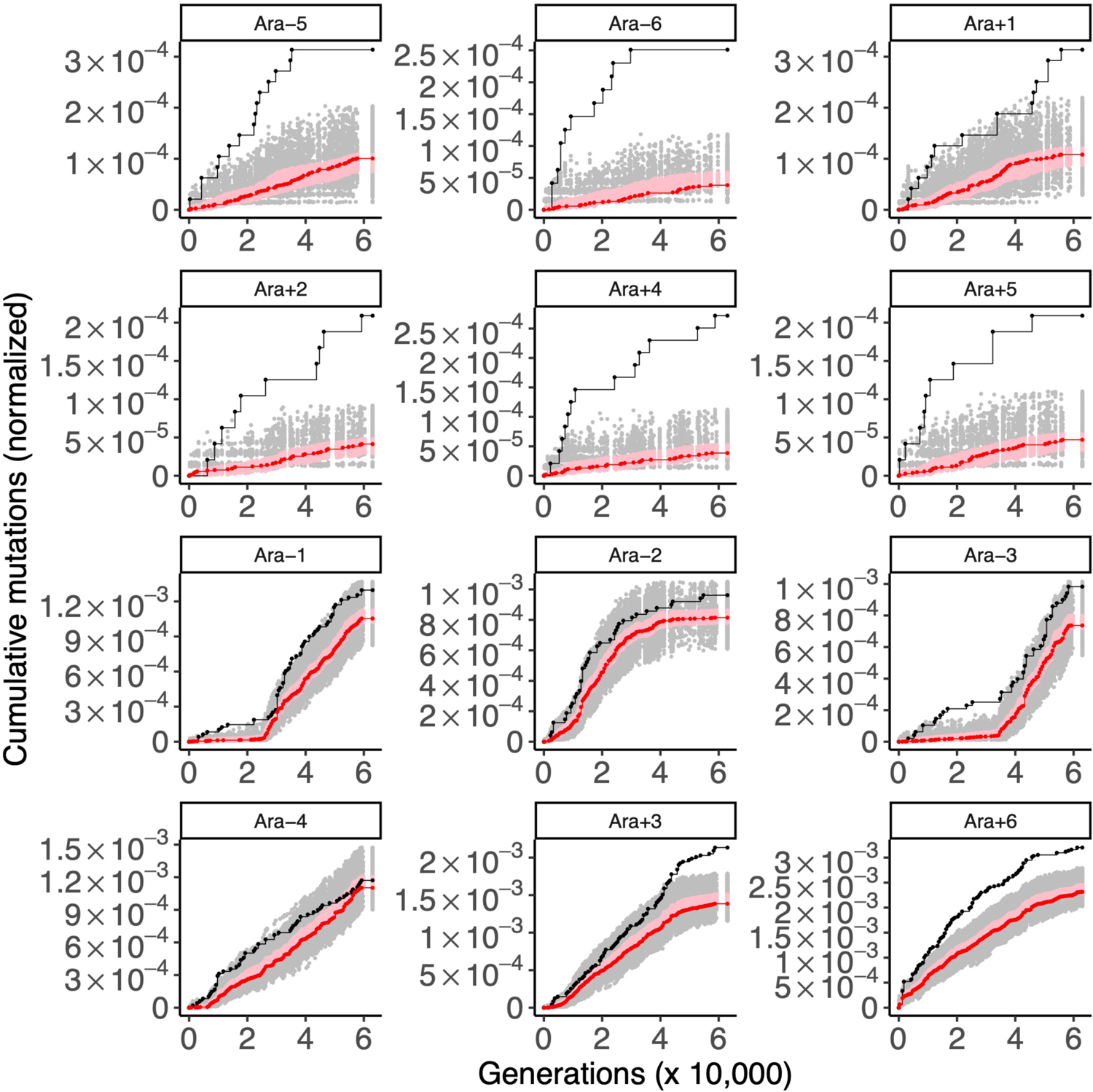
I-modulon regulators evolve under strong positive selection, while genes regulated within I-modulon evolve under weaker selection. Each panel shows the cumulative number of mutations in 70 I-modulon regulators (black) and in 1,394 genes regulated within I-modulons (red) in the 12 LTEE populations. For comparison to the I-modulon regulators, random sets of 70 genes were sampled 1,000 times, and the cumulative number of mutations in those random gene sets, normalized by gene length, were calculated. The middle 95% of the null distribution for I-modulon regulators is shown in gray. A similar procedure was used with random sets of 1,394 genes to make a null distribution in pink to compare to the genes regulated within I-modulons. The top six panels are the nonmutator populations and the bottom six panels are the hypermutator populations.

Figure 8 also shows that the tempo of I-modulon regulator evolution is quite variable across populations. Consider the number of mutations in I-modulon regulators in the nonmutator populations. After 40,000 generations, Ara+1 has 5, Ara+2 has 4, Ara+4 has 2, and Ara+5 has 1, while Ara−5 and Ara−6 have none. Variation in the tempo of evolution of I-modulon regulators is also seen in the hypermutator populations, in particular the strong signal of ongoing positive selection found in Ara+3 and Ara+6, in contrast with the complete lack of signal in Ara−4.

While I-modulons tend to evolve at background rates within individual populations (Figure 8), some specific I-modulons show evidence of idiosyncratic selection in the hypermutator populations. We hypothesize that some of these cases represent historical contingency and epistasis in the LTEE; these hypotheses could be tested in future studies. The results for all I-modulons are provided in Supplementary Files 1 and 2, while particular I-modulons of interest are listed in Supplementary Table 3.

#### Summary of module decomposition results

Altogether, we find that the protein-coding genes with the strongest transcriptional response to changing conditions show little to no evidence of selection in the LTEE. By contrast, top-level transcriptional regulators showed strong signatures of positive selection in all nonmutator populations, as well as in some hypermutator populations. Based on these findings, we hypothesize that the genes that show the strongest transcriptional response to changing conditions tend to be at the lower levels of the gene regulatory network, while natural selection in the LTEE is largely operating on regulators of those genes.

### Cis-regulatory regions of key regulators show significantly more parallel evolution than the cis-regulatory regions of downstream targets

If upper levels of the gene regulatory network are under stronger selection in the LTEE, then *cis*-regulatory regions of I-modulon regulators should also show more evidence of positive selection than the genes within I-modulons. We tested this prediction by examining non-coding mutations associated with I-modulon regulators and their downstream targets. These non-coding mutations occurred within the promoter regions (up to 100 bp upstream) of their annotated gene [16]. There are 48 non-coding mutations associated with 70 I-modulon regulators, and 542 non-coding mutations associated with 1394 genes regulated within I-modulons. These data are consistent with our prediction (Binomial test with 48 successes out of 590 trials and expected probability of success = 70/1464: *p* = 0.0005).

### Application of STIMS to hypermutator populations of Pseudomonas aeruginosa evolving under antibiotic selection

We wanted to know whether STIMS could be applied beyond the LTEE. Unlike the LTEE, however, most evolution experiments do not have genomic, let alone metagenomic, data spanning 30+ years of evolutionary history. We sourced the metagenomic time series data reported by Mehta et al. [8], in which replicate populations of *Pseudomonas aeruginosa* adapted to subinhibitory concentrations of colistin in continuous chemostat culture for one month. We reasoned that the Mehta dataset would be ideal for applying STIMS since both replicate populations in that experiment rapidly evolved hypermutator phenotypes. We asked whether annotated regulatory genes in the *P. aeruginosa* PAO11 genome showed evidence of positive selection in this evolution experiment (Figure 9, Supplementary Figure S11); indeed, STIMS reveals a clear signal of positive selection on the pre-specified set of 424 regulatory genes (one-tailed bootstrap: *p* < 0.001). This finding reveals that STIMS has the power to detect gene sets under strong positive or purifying selection, even when applied to metagenomic time series which span relatively short time periods.

**Figure 9.**
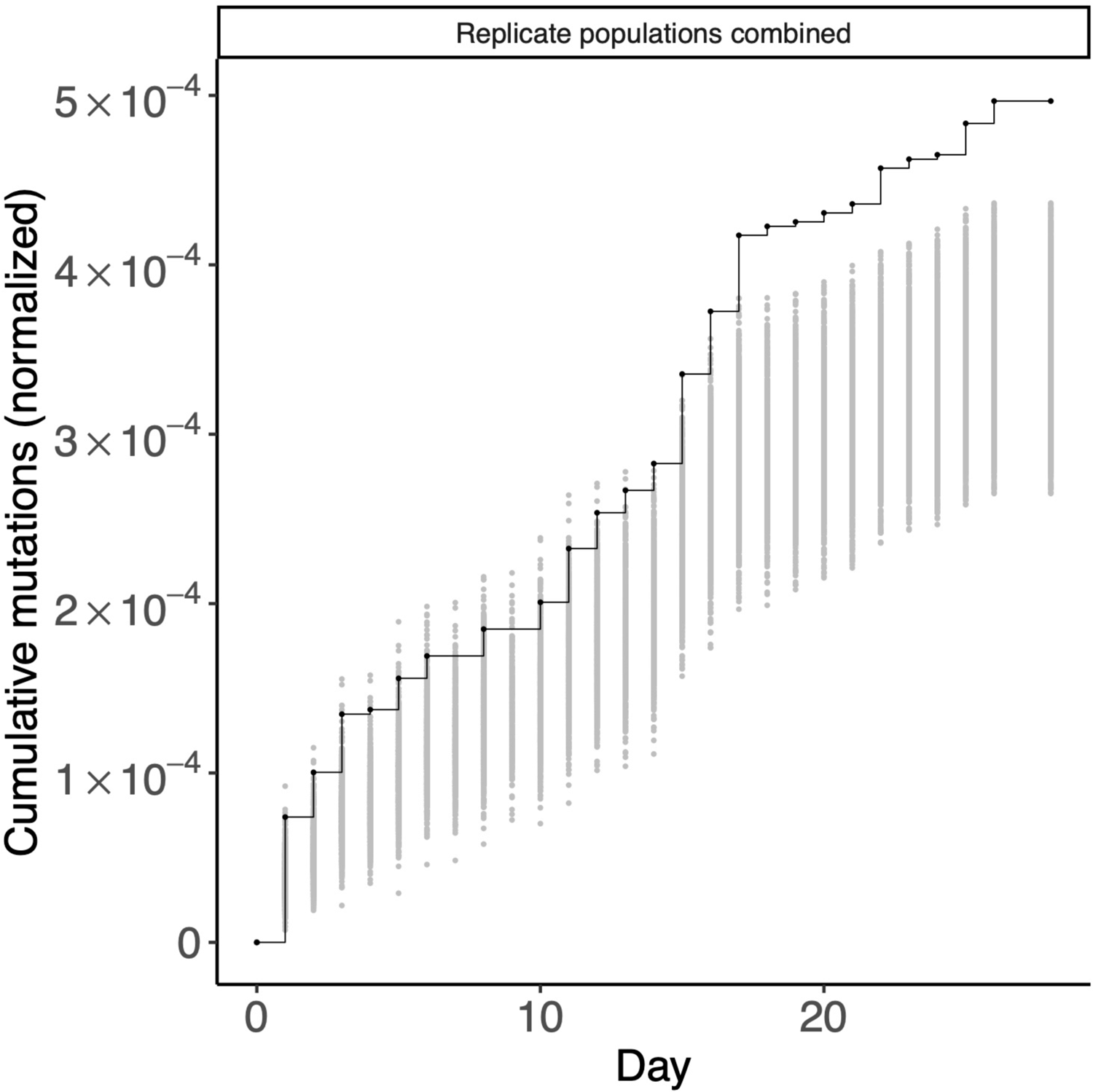
Regulatory genes are under positive selection in hypermutator *Pseudomonas aeruginosa* populations evolving under subinhibitory concentrations of colistin. The cumulative number of mutations in 424 annotated regulatory genes in two replicate populations of hypermutator *P. aeruginosa* is shown in black. For comparison, random sets of 424 genes were sampled 1,000 times, and the cumulative number of mutations in those random gene sets, normalized by gene length, were calculated. The middle 95% of this null distribution is shown in gray.

## DISCUSSION

We show that signals of both positive and purifying selection can be detected in metagenomic time series of large asexual haploid populations, including *Escherichia coli* from Lenski’s long-term evolution experiment (LTEE) and a clinically relevant pathogen, *Pseudomonas aeruginosa*, in a short-term evolution experiment under antibiotic selection.

In contrast to previous studies, which examined global patterns in the tempo and mode of evolution in the LTEE [13, 15, 16], our analysis focuses on functional modules encoded by the *E. coli* genome. We find that the tempo of evolution in particular molecular subsystems gives us insight into the mode of evolution acting on those modules (i.e., relaxed, purifying, or positive selection). We ran simulations to demonstrate that STIMS works in an idealized system in which selection pressures and the genomic DFE can be pre-specified *a priori*, and we ran computational positive control experiments to confirm that STIMS works on genes with prior evidence of relaxed (Figure 5), purifying (Figure 6), and positive selection (Figure 7).

Our findings indicate that the accelerated pace of genomic evolution in the hypermutators, combined with the detailed record of molecular evolution provided by metagenomic time series, may open new opportunities for understanding the genomic basis of adaptive evolution, even though the signal of selection is obscured in hypermutator genomes due to genomic draft [6]. First, the vast number of nearly neutral hitchhikers that are observed in hypermutator populations provides insight into how mutations in modifier alleles affect genome-wide and local mutation rates and biases [31]. Second, regions of the genome that are highly depleted in mutations in the hypermutator populations are strong candidates for purifying selection. Third and most importantly— metagenomic sequencing gives deep sampling of genetic variation off the (eventual) line of descent.

We found compelling evidence of purifying selection in the hypermutator populations of the LTEE. Many of the strongest candidate genes for purifying selection are deeply conserved over evolutionary time, such as those encoding ribosomal subunits. The action of purifying selection on the hypermutator populations is also consistent with the observation that antimutator alleles have fixed in several populations of the LTEE [31, 36], and with the experimental finding that overexpressing RNA chaperones in some hypermutator LTEE strains reduces the mutation load in those strains [37]. Together, these observations suggest that hypermutator lineages are accumulating some deleterious passenger mutations, even as their fitness continues to increase. In any case, our analysis shows that the depletion of mutations in particular genes is not due to the evolution of antimutator alleles: by bootstrapping a null distribution of background rates, STIMS controls for the genome- and population-wide effects of antimutator alleles over time.

The resampling approach used by STIMS can also take the effects of local mutation biases into account [38]. For instance, one can control for local mutation biases by constructing null distributions that directly model chromosomal variation in mutation rates and biases. This can be somewhat complicated, given that mutational biases [6] and regional mutation rates over the chromosome have evolved idiosyncratically across the replicate LTEE populations [31]. Our implementation of STIMS samples gene sets uniformly over the *E. coli* genome: we also implemented a sampling procedure that take the wave pattern of mutation rate variation over the *E. coli* chromosome into account [31, 39, 40], but this more complicated procedure produced the same results as sampling genes uniformly over the chromosome (Materials and Methods).

When we applied STIMS to different module decompositions of the *E. coli* genome, we found compelling evidence of strong positive selection on key global regulators of the *E. coli* gene regulatory network, especially in comparison to the genes that they regulate. Furthermore, we found an excess of mutations in the *cis*-regulatory regions of those regulators in comparison to the genes that they regulate. One explanation could be that mutations that affect the *cis*-regulation and structure of global regulators at higher levels of the GRN cause a cascade of effects on downstream targets, and so are more effective targets for fine-tuning the *E. coli* GRN. Strikingly, we found evidence of continued strong positive selection on key regulatory genes in the two populations with the most mutations in the LTEE: Ara+3 and Ara+6 (Figure 8). By contrast, two nonmutator populations, Ara−5 and Ara−6, showed no mutations at all in I-modulon regulators in the last 20,000 generations of the time course. This suggests that the GRN may initially evolve close to some local fitness maximum (subject to pleiotropic constraints), but then evolve further to compensate for the effects of mutations elsewhere in the genome.

Further work will be needed to test the hypothesis that ongoing evolution of I-modulon regulators is related to compensatory evolution. It is possible that the hypermutator populations are evolving to compensate for an increasing mutation load of deleterious hitchhiker mutations [8]. The appearance of multiple antimutator alleles in the LTEE suggests that that hypermutability has a hidden cost [15, 31, 36], but the magnitude of any mutation load in the LTEE populations remains unknown. Ongoing positive selection on the *E. coli* GRN could also be due to compensatory evolution that is not specifically for hitchhiking deleterious mutations. For instance, it is possible that early beneficial mutations become deleterious due to further mutations [41, 42]. Furthermore, natural selection may greedily favor mutationally accessible but suboptimal trajectories [43, 44] that then open new, idiosyncratic paths for further refinement [45].

Our work has some limitations. By using STIMS to examine the tempo of GRN evolution in the LTEE using the same dataset, we find a number of patterns with an unknown rate of false positives. Therefore, future work is needed to test the specific hypotheses that we have generated, either by using additional time-course data from the LTEE, by analyzing related evolution experiments, or by experimental validation. An important caveat is that deletions that fix in the LTEE can lead to spurious inferences of purifying selection, if they are not taken into account, since deleted genes cannot accumulate mutations. By that same token, gene duplications, amplifications, or other forms of copy number variation could lead to spurious inferences of positive selection, or elevated mutation rates [46, 47]. Those types of mutations appear to be rare in these data, in comparison to the point mutations, indels, and transposon insertions that we count. The complications induced by copy number variation may need to be considered (or safely ignored), depending on the context in which STIMS is used. Finally, STIMS is computationally intensive due to its use of bootstrapping. Often equivalent statistical results can be derived much faster, using exact tests like the binomial test or Fisher’s exact test [8], or by using tests based on the Poisson distribution [34, 48]. The advantage of STIMS is that it provides greater biological insight, by visualizing *how* statistical signatures of positive and purifying selection change in an evolving population over time.

Overall, our work provides greater insight into the mechanistic connection between genotypic and phenotypic evolution in evolution experiments like the LTEE. The genetic architecture underlying fitness improvements in the LTEE likely involves a relatively small number of loci that control global aspects of cellular physiology [15, 16, 23, 49]. These factors may control many downstream pathways, which are being modulated in response to selection in the LTEE. Hence, we hypothesize that the regulation of these pathways is being rewired during the LTEE, often without significantly changing their downstream effectors. This would explain why the modules that show the strongest changes in expression, and best predict growth rate and fitness in response to environmental conditions [26–28] show little evidence of adaptive evolution in the LTEE. Overall, the genotype-phenotype map of *E. coli*, at least in the context of the LTEE, resembles the hierarchical “supervisor-worker” gene architecture proposed by Chen et al. [50] to explain the genetic architecture of quantitative traits in *Saccharomyces cerevisiae*.

As the costs of protein production exerts a strong contrast on cellular energetics, the proper allocation of proteome resources in order to maximize growth, rather the evolution of particular genetic modules, may be the cellular phenotype under strongest selection in the LTEE [51–54]. Future work could test this hypothesis by trying to mimic evolutionary changes in the LTEE by directly perturbing the balance of global physiological and metabolic state variables (chromosome conformation, ppGpp, cAMP levels and redox potential) to maximize growth in the ancestral REL606 strain. Coevolution between the function of those proteome resources, pleiotropic constraints on optimal allocation, as well as feedback with the environment could all play a part in causing the open-ended increases in fitness seen in the LTEE [20, 35].

Finally, we expect our methodology will be broadly useful, especially for workers in the experimental evolution field. To demonstrate the generality of STIMS, we reanalyzed data from a second evolution experiment in which two replicate hypermutator populations of *P. aeruginosa* adapted to subinhibitory concentrations of colistin [8]. Our finding of positive selection on regulatory genes in this experiment validates STIMS as a general method and indicates that the rapid evolution of bacterial gene regulatory networks may be a general mechanism for adaptation during experimental evolution, and perhaps during adaptation to novel environments in general. We expect that our basic idea could be more rigorously justified by deeper theoretical work, and further extended to study evolving populations and communities in both the laboratory and in the wild. Purifying selection is of particular interest in the study of viruses and cancers, for the sake of finding conserved and effective drug targets [55–58]. In particular, we anticipate that STIMS could be applied to clinical time series, such as genomic sampling from cystic fibrosis patients [59, 60], in order to discover gene modules that are under selection in pathogens as they adapt to their host [61, 62].

## MATERIALS AND METHODS

### The STIMS algorithm

In brief, STIMS counts the cumulative number of mutations occurring over time in a module of interest, and compares that number to a null distribution that is constructed by subsampling random sets of genes over the genome, in which the cardinality of these random gene sets is fixed to the cardinality of the module of interest. We normalize the number of observed mutations in a gene set by the total length of that gene set in base-pairs.

In this work, we specifically counted point mutations, small insertions and deletions (indels), and transposon insertions (structural variants) that were detected in metagenomic time series using the *breseq* pipeline [8, 16, 63]. Nonetheless, STIMS does not depend on any specific variant calling software, nor does it specifically require metagenomic time series (genomic time series would suffice). STIMS only requires a list of observed *de novo* mutations over time (i.e. information about the location of each mutation, and the time at which each was first observed in the population).

By bootstrapping a null distribution based on random gene sets, STIMS implicitly controls for genome-wide variation in the mutation rate (since this will affect all genes). In order to control for local variations in mutation rate [31, 39, 40], we subsampled random modules from the same chromosomal neighborhood as the module of interest (dividing the REL606 genome into 46 bins, roughly ∼100,000 bp each), under the assumption that nearby genes have similar mutation rates. This approach gave the same results as sampling genes uniformly across the genome, despite a higher computational cost. For this reason, the results reported here use the simplest method to construct the null distribution, in which genes are sampled without considering their position on the chromosome.

Bootstrapped *p-*values were calculated separately for each population. *p*-values for one-sided tests are exact estimates of the relevant tail of the null distribution, where *p* is the upper-tail (or lower-tail) probability of the event of sampling a normalized cumulative number of mutations under the null distribution that is greater than (or less than) the normalized cumulative number of mutations of the module of interest. Two-tailed tests against the bootstrapped null distribution were calculated with a false-positive (type I error) probability *α* = 0.025 to account for both tails; this is equivalent to multiplying *p-*values by a factor of 2, in a Bonferroni correction for two statistical tests (i.e., on the upper tail and on the lower tail of the null distribution).

In the visualizations shown in the figures, the top 2.5% and bottom 2.5% of points in the null distribution are omitted, such that each panel can be directly interpreted as a randomization test with a false-positive (type I error) probability *α* = 0.05.

### Filtering of genes in REL606 genome

We analyzed the pre-processed LTEE metagenomics data published by [16], and so we applied filters to exclude regions of the REL606 genome that were omitted from that analysis. In particular, pseudogenes and genes in repetitive regions of the genome were excluded. A site was marked as repetitive if (1) it was annotated as a repeat region in the REL606 reference, (2) it was present in the set of masked regions compiled by [15], or (3) it fell within the putative prophage at REL606 genome coordinates 880528–904682.

### Essential genes excluded from purifying selection control experiment

We excluded essential genes that showed parallel evolution in Tenaillon et al. [15]. In that work, a G-score was calculated to measure parallel evolution. G-scores were calculated separately for the nonmutator and hypermutator populations. We ranked all genes in REL606 that passed our filters by their nonmutator and hypermutator G-scores. Essential genes that were in the top 50 of either list were excluded from the purifying selection control experiment. 21 out of the top 50 G-scoring nonmutator genes were essential in REL606, based on the transposon mutagenesis and sequencing experiments carried out by Couce et al. [6]. These genes were: *arcA*, *arcB*, *crp*, *fabF*, *ftsI*, *hflB*, *infB*, *infC*, *mrdA*, *mreB*, *mreC*, *mreD*, *nusA*, *rne*, *rplF*, *rpoB*, *rpsD*, *sapF*, *spoT*, *topA*, and *yabB*. 3 out of the top 50 G-scoring hypermutator genes were essential in REL606. These genes were: *sapA*, *tilS*, and *yhbH*.

### Analysis of regulatory genes in the month-long hypermutator P. aeruginosa evolution experiment

We analyzed the pre-processed metagenomics data from the hypermutator *P. aeruginosa* evolution experiment published by Mehta et al. [8]. We excluded the Pf4 bacteriophage in the *P. aeruginosa* PAO11 reference genome used by Mehta et al. from the STIMS analysis; this region was also omitted from the statistical analysis in the original report [8].

We used the keywords “regulator”, “regulatory”, and “regulation” to search for regulatory genes in the PAO11 genome to identify *de novo* mutations arising in their evolution experiment. This search resulted in a set of 424 regulatory genes, which we then analyzed using STIMS.

### Implementation of a test for selection based on the Poisson distribution

We implemented the Poisson method described by Kinnersley et al. [34]. We summed all mutations across the LTEE associated with genes that passed our filters, and divided by the total length of those genes to calculate the background density of mutations per site, *λ*. For each gene *g* in the genome, we scale *λ* by the length of *g*, to calculate the expected number of mutations at gene *g*: *λ_g_*. The same calculation holds for a set of genes: by multiplying *λ* by the total length of genes in the set, we can calculate the expected number of mutations in the gene set, *λ_set_*. Suppose *x* mutations were counted in the gene set of interest. Then, a one-tailed test for purifying selection in this gene set can be done by calculating a *p-*value from the lower-tail of the Poisson distribution: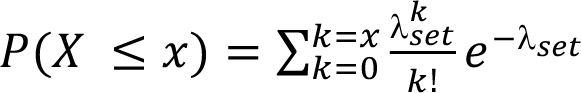. Similarly, a one-tailed test for positive selection in the gene set can be done by directly calculating a *p-*value from the upper-tail of the Poisson distribution: 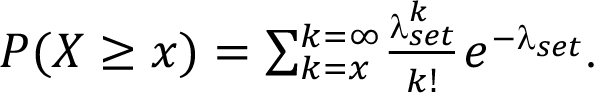

When multiple genes or gene sets were tested for positive or purifying selection, we applied a Benjamini-Hochberg correction for multiple-testing, and report those false-discovery-rate (FDR) corrected *p*-values.

### Wright-Fisher simulations of microbial evolution

We implemented Wright-Fisher evolutionary simulations of asexual haploid populations in SLiM [64]. We simulated populations of 1 million cells, each containing a genome of 1,000 contiguous genes. We tested STIMS in two scenarios: the evolution of a nonmutator populations with a point-mutation rate of 10^−10^ per base-pair per generation, and the evolution of a hypermutator populations with a 100-fold increase in mutation rate: 10^−8^ per base-pair per generation. These population genetic parameters are comparable to the effective population size (3.3 *×* 10^7^) and the ancestral mutation rate (8.9 *×*10^−11^) of Lenski’s LTEE [29], and the 100-fold evolved increases in mutation rates observed in the LTEE [31]. Our simulations assume a background genomic distribution of mutation fitness effects (DFE) in which the vast majority of mutations are nearly neutral, with a long tail of deleterious mutations, based on high-throughput measurements of the DFE in *E. coli* in microfluidic mutation accumulation experiments [65]. We assume that 100 genes are evolving under positive selection, such that 10% of mutations occurring in those genes are beneficial. This assumption implies that 0.0025 of mutations are beneficial in our model, which is comparable to the experimental estimate that 1 out of 150 newly arising mutations in *E. coli* are beneficial in laboratory conditions [66]. Following recent simulations of adaptive evolution in the LTEE, we draw beneficial mutations from an exponential DFE with mean 0.01578 [29]. We further assume that 100 genes are evolving completely neutrally such that mutations in these genes have no fitness effect, and assume that another set of 100 genes are evolving under purifying selection, such that 40% of mutations in those genes cause a 30% decrease in fitness. This last assumption matches the finding that 1% of new mutations in *E. coli* are lethal under laboratory conditions [65].

Ten replicate simulations were run for both the nonmutator and hypermutator populations, and all mutations in each simulation were saved at 100-generation intervals. Mutations were filtered based on a 1% allele frequency threshold, to simulate metagenomic sequencing: this is equivalent to uniform 100*×* short-read sequencing coverage across the genome per sequencing sample, with no sequencing errors. We then ran STIMS on the 100 genes under positive selection, the 100 genes under relaxed selection (those evolving completely neutrally), and the 100 genes under purifying selection. We examined the sensitivity and specificity of STIMS by varying the allele frequency detection threshold, the number of genes sampled per module, the sampling interval in the time series, and the length of the time series, holding all other setting to their default values. Four values of the allele frequency detection limit were examined: 0.01%, 1%, 5%, 10%. The number of genes sampled per module was varied between 1, 25, 50, and 100 genes. The sampling interval was either 100- or 500-generation intervals. The length of the time series was 500, 1000, 2000, or 5000 generations. True positives and false negatives were counted by the number of STIMS tests on the genes evolving under positive or purifying selection that passed a one-sided *p*-value significance threshold of 0.05.

False positives and true negatives were counted by the number of STIMS tests on the genes evolving neutrally that nevertheless passed the same *p*-value significance threshold. Sensitivity (also known as the true positive rate, or as statistical power) was calculated as the number of true positives divided by the sum of true positives and false negatives, out of the 10 tests on the 10 replicate datasets. Specificity (also known as the true negative rate) was calculated as the number of true negatives divided by the sum of true negatives and false positives, out of the 10 tests on the 10 replicate datasets.

### Data and analysis code availability

All analysis codes for this project are available at: www.github.com/rohanmaddamsetti/LTEE-purifying-selection. A stand-alone implementation of STIMS in the Julia programming language is available at: www.github.com/rohanmaddamsetti/STIMS. The sequence data used in our study have already been deposited into a public database. The other data and analysis scripts in this manuscript have been archived on the Zenodo Digital Repository (DOI: http://doi.org/10.5281/zenodo.4783368).

## ACKNOWLEDGEMENTS

We thank Alejandro Couce for sharing the list of gold-standard neutral genes in *E. coli* B strain REL606. We thank Richard Lenski for valuable discussions and advice. We thank Jeffrey Barrick for valuable discussions, advice, and comments on an earlier version of our manuscript. We also thank Benjamin Good for making pre-processed LTEE metagenomic data and analysis scripts accessible for the research community. The LTEE that generated the bacteria we used in this study is supported by a grant from the National Science Foundation (currently DEB-1951307) to Richard Lenski and Jeffrey Barrick. N.A.G. was supported in part by the BEACON Center for the Study of Evolution in Action (NSF cooperative agreement DBI-0939454) and by Michigan State University.

## SUPPLEMENTARY INFORMATION

Supplementary File 1. Results of running STIMS, over all LTEE populations, on each I-modulon in Sastry et al. (2019).

Supplementary File 2. Results of running STIMS, individually on each each LTEE population, on each I-modulon in Sastry et al. (2019).

Supplementary File 3. Results of running Poisson method as a genome-wide screen for positive and purifying selection, over all LTEE populations, and individually on each LTEE population.

## SUPPLEMENTARY TEXT

### Comparison of STIMS to a test for selection based on the Poisson distribution

One class of statistical tests for selection in evolution experiments uses the Poisson distribution to model the expected distribution of mutations per genomic site per unit time under neutral evolution [16, 34]. Using this framework, significant deviations from the Poisson null expectation can indicate positive or purifying selection. In principle, Poisson methods should be simpler to implement and computationally cheaper than STIMS. However, methods based on the Poisson distribution may be sensitive to model misspecification, due to local mutation biases that can cause deviation from the Poisson expectation in the absence of selection [31]. Variations of the bootstrapping approach used by STIMS, on the other hand, can account for such mutational biases through randomization (Materials and Methods).

To better understand the relative strengths and weaknesses of STIMS in comparison to Poisson methods, we implemented the Poisson method used by Kinnersley *et al.* [34], and re-ran all of our positive control experiments on gold-standard sets of genes (Materials and Methods). Overall, both STIMS and the Poisson method give similar results. The Poisson method successfully recovers signals of purifying selection (*p* < 10^−14^) and positive selection (*p* < 10^−50^) on the respective sets of gold-standard genes. However, the Poisson method finds a weak signal of positive selection on the gold-standard set of genes under relaxed selection (*p* = 0.0397). The Poisson method finds a signal of purifying selection on most proteome sectors (FDR-corrected *p* < 10^−11^ for U-sector; FDR-corrected *p* < 0.00001 for A-sector; FDR-corrected *p* = 0.000162 for R-sector; FDR-corrected *p =* 0.00182 for S-sector; FDR-corrected *p* = 0.00217 for O-sector; FDR-corrected *p* = 0.391 for C-sector). Eigengenes show little to no evidence of selection under the Poisson method (FDR-corrected *p* = 0.0327 for Eigengene 2; the rest are nonsignificant). I-modulon regulators show evidence of very strong positive selection (*p* < 10^−13^), while 16 I-modulons show evidence of either positive or purifying selection at a Benjamini-Hochberg corrected *p*-value threshold of 0.01 (Supplementary Table 4). 14 out of these 16 I-modulons are significant by STIMS, while the other two, “flu-yeeRS” and “Pyruvate”, trend toward significance (Supplementary File S1). Finally, we used the Poisson method as a genome-wide screen for positive and purifying selection, both over all LTEE populations, and on each population individually. We report these results at a Benjamini-Hochberg corrected *p*-value threshold of 0.01 in Supplementary File S3. Most of the genes showing evidence of positive selection have been reported before [15, 16, 23]. However, this screen finds 7 genes with a significant signal of purifying selection across the LTEE, none of which have been reported before: *acnB* (aconitate hydratase), *ybjL* (hypothetical protein), *yhdP* (conserved membrane protein predicted transporter), *recG* (ATP-dependent DNA helicase), *cyoB* (cytochrome o ubiquinol oxidase subunit I), *wcaJ* (predicted UDP-glucose lipid carrier transferase), and *atpD* (F0F1 ATP synthase subunit beta). All of these genes are longer than 1,300 bp, and three of them (*ybjL*, *wcaJ*, and *atpD*) have no observed mutations at all.

Altogether, this detailed comparison indicates the Poisson method is faster and is more practical for genome-wide scans for positive and purifying selection on single genes, but that its sensitivity comes at the price of more false positives. STIMS is more conservative than the Poisson method, and gives results that are more consistent with ground truth, as defined by the sets of gold-standard genes that we used for empirical validation. STIMS, unlike the Poisson method, also provides a picture of the tempo of evolution in a gene set over time, as well as changes in statistical significance over time. These visualizations are especially useful for observing *how* the tempo of evolutionary change in a gene set of interest changes over time, and for generating further hypotheses and predictions for empirical validation (Supplementary Files 1 and 2).

## SUPPLEMENTARY TABLES AND FIGURES

**Supplementary Table S1.** Essential genes in REL606 with no observed mutations in the LTEE metagenomics data. encoded by prophage CP-933O

| Gene | Locus tag | Gene length | Product |
| --- | --- | --- | --- |
| *rlpB* | ECB_00610 | 582 | minor lipoprotein |
| *rplJ* | ECB_03861 | 498 | 50S ribosomal protein L10 |
| *ECB_01539* | ECB_01539 | 408 | protein similar to DicA regulator of DicB<br>encoded by prophage CP-933O |
| *sdhD* | ECB_00682 | 348 | succinate dehydrogenase cytochrome b556<br>small membrane subunit |
| *rplV* | ECB_03166 | 333 | 50S ribosomal protein L22 |
| *rpsJ* | ECB_03172 | 312 | 30S ribosomal protein S10 |
| *rpsN* | ECB_03158 | 306 | 30S ribosomal protein S14 |
| *yrbB* | ECB_03056 | 294 | hypothetical protein |
| *groES* | ECB_04012 | 294 | co-chaperonin GroES |
| *rpsS* | ECB_03167 | 279 | 30S ribosomal protein S19 |
| *rpsT* | ECB_00027 | 264 | 30S ribosomal protein S20 |
| *rpsQ* | ECB_03162 | 255 | 30S ribosomal protein S17 |
| *atpE* | ECB_03621 | 240 | F0F1 ATP synthase subunit C |
| *rpmC* | ECB_03163 | 192 | 50S ribosomal protein L29 |
| *ydaE* | ECB_01329 | 171 | conserved protein |
| *rpmJ* | ECB_03150 | 117 | 50S ribosomal protein L36 |
| *ybgT* | ECB_00694 | 114 | hypothetical protein |
| *rhoL* | ECB_03660 | 102 | rho operon leader peptide |

**Supplementary Table S2.** Essential genes in REL606 with only synonymous mutations in the LTEE metagenomics data.

| Gene | Locus tag | Gene length | Product |
| --- | --- | --- | --- |
| *atpD* | ECB_03616 | 1383 | F0F1 ATP synthase subunit beta |
| *icdA* | ECB_01134 | 1251 | isocitrate dehydrogenase |
| *thyA* | ECB_02675 | 795 | thymidylate synthase |
| *plsC* | ECB_02890 | 738 | 1-acyl-sn-glycerol-3-phosphate acyltransferase |
| *pdxH* | ECB_01608 | 657 | pyridoxamine 5'-phosphate oxidase |
| *rplD* | ECB_03170 | 606 | 50S ribosomal protein L4 |
| *lpcA* | ECB_00217 | 579 | phosphoheptose isomerase |
| *rimM* | ECB_02497 | 549 | 16S rRNA-processing protein |
| *rplE* | ECB_03159 | 540 | 50S ribosomal protein L5 |
| *sixA* | ECB_02264 | 486 | phosphohistidine phosphatase |
| *folK* | ECB_00141 | 480 | 2-amino-4-hydroxy-6-hydroxymethyldihydropteridine<br>pyrophosphokinase |
| *accB* | ECB_03113 | 471 | acetyl-CoA carboxylase |
| *holD* | ECB_04247 | 414 | DNA polymerase III subunit psi |
| *rpsI* | ECB_03090 | 393 | 30S ribosomal protein S9 |
| *iscU* | ECB_02421 | 387 | scaffold protein |
| *rplQ* | ECB_03145 | 384 | 50S ribosomal protein L17 |
| *panD* | ECB_00130 | 381 | aspartate 1-decarboxylase precursor |
| *rpsL* | ECB_03193 | 375 | 30S ribosomal protein S12 |
| *rplN* | ECB_03161 | 372 | 50S ribosomal protein L14 |
| *rnpA* | ECB_03587 | 360 | ribonuclease P |
| *rplT* | ECB_01685 | 357 | 50S ribosomal protein L20 |
| *yadR* | ECB_00155 | 345 | hypothetical protein |
| *fdx* | ECB_02417 | 336 | [2Fe-2S] ferredoxin |
| *secG* | ECB_03040 | 333 | protein-export membrane protein |
| *ybaB* | ECB_00422 | 330 | hypothetical protein |
| *acpP* | ECB_01090 | 237 | acyl carrier protein |
| *rpmI* | ECB_01686 | 198 | 50S ribosomal protein L35 |
| *rpmH* | ECB_03586 | 141 | 50S ribosomal protein L34 |
| *pheL* | ECB_02487 | 48 | pheA gene leader peptide |
| *trpL* | ECB_01239 | 45 | trp operon leader peptide |

**Supplementary Table S3.**
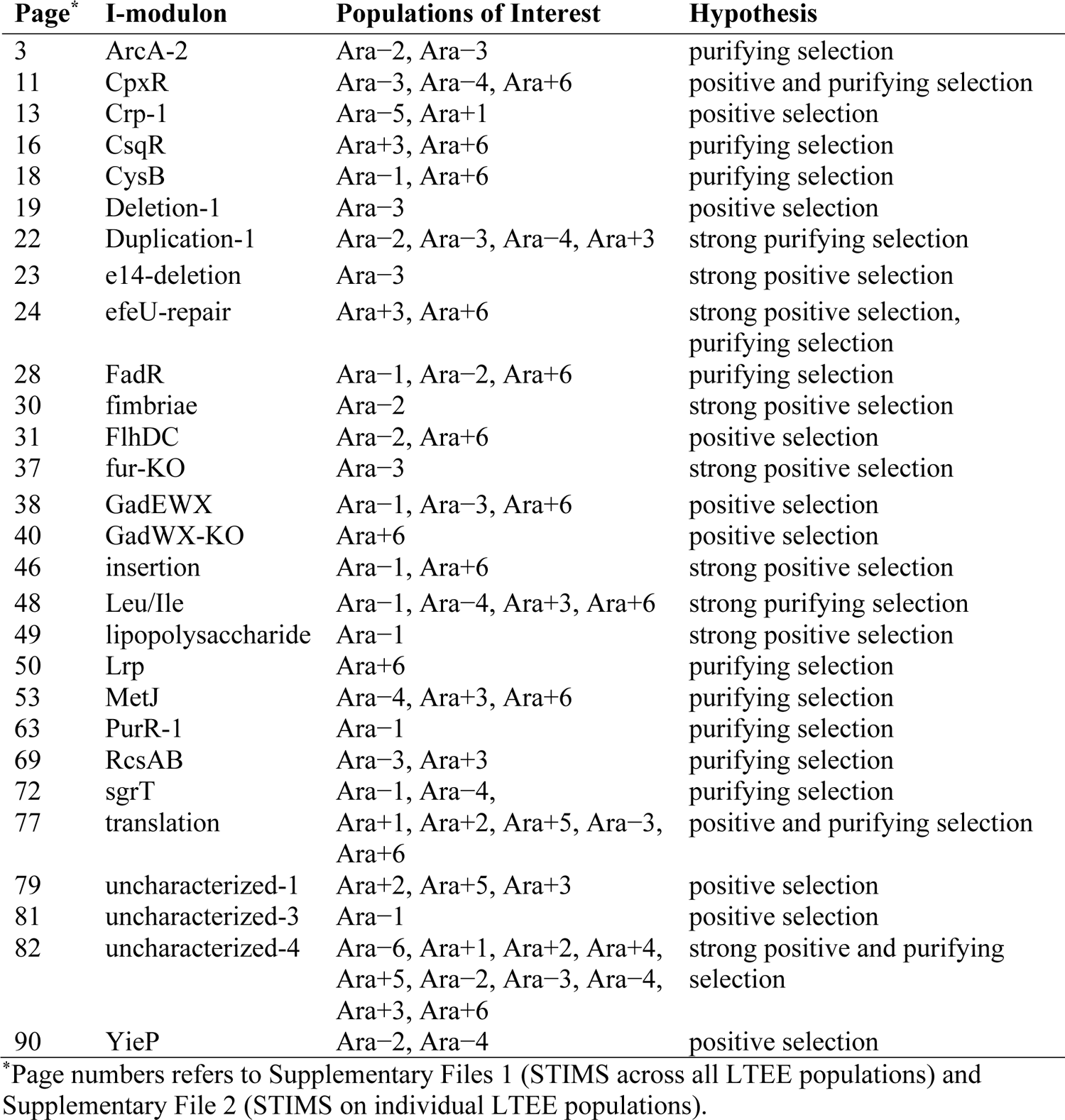
I-modulons showing evidence of selection, historical contingency and epistasis in the LTEE, using STIMS.

| Page* | I-modulon | Populations of Interest | Hypothesis |
| --- | --- | --- | --- |
| 3 | ArcA-2 | Ara-2, Ara-3 | purifying selection |
| 11 | CpxR | Ara-3, Ara-4, Ara+6 | positive and purifying selection |
| 13 | Crp-1 | Ara-5, Ara+1 | positive selection |
| 16 | CsqR | Ara+3, Ara+6 | purifying selection |
| 18 | CysB | Ara-1, Ara+6 | purifying selection |
| 19 | Deletion-1 | Ara-3 | positive selection |
| 22 | Duplication-1 | Ara-2, Ara-3, Ara-4, Ara+3 | strong purifying selection |
| 23 | e14-deletion | Ara-3 | strong positive selection |
| 24 | efeU-repair | Ara+3, Ara+6 | strong positive selection, purifying selection |
| 28 | FadR | Ara-1, Ara-2, Ara+6 | purifying selection |
| 30 | fimbriae | Ara-2 | strong positive selection |
| 31 | FlhDC | Ara-2, Ara+6 | positive selection |
| 37 | fur-KO | Ara-3 | strong positive selection |
| 38 | GadEWX | Ara-1, Ara-3, Ara+6 | positive selection |
| 40 | GadWX-KO | Ara+6 | positive selection |
| 46 | insertion | Ara-1, Ara+6 | strong positive selection |
| 48 | Leu/Ile | Ara-1, Ara-4, Ara+3, Ara+6 | strong purifying selection |
| 49 | lipopolysaccharide | Ara-1 | strong positive selection |
| 50 | Lrp | Ara+6 | purifying selection |
| 53 | MetJ | Ara-4, Ara+3, Ara+6 | purifying selection |
| 63 | PurR-1 | Ara-1 | purifying selection |
| 69 | RcsAB | Ara-3, Ara+3 | purifying selection |
| 72 | sgrT | Ara-1, Ara-4, | purifying selection |
| 77 | translation | Ara+1, Ara+2, Ara+5, Ara-3, Ara+6 | positive and purifying selection |
| 79 | uncharacterized-1 | Ara+2, Ara+5, Ara+3 | positive selection |
| 81 | uncharacterized-3 | Ara-1 | positive selection |
| 82 | uncharacterized-4 | Ara-6, Ara+1, Ara+2, Ara+4, Ara+5, Ara-2, Ara-3, Ara-4, Ara+3, Ara+6 | strong positive and purifying selection |
| 90 | YieP | Ara-2, Ara-4 | positive selection |

\*Page numbers refers to Supplementary Files 1 (STIMS across all LTEE populations) and Supplementary File 2 (STIMS on individual LTEE populations).

**Supplementary Table S4.**
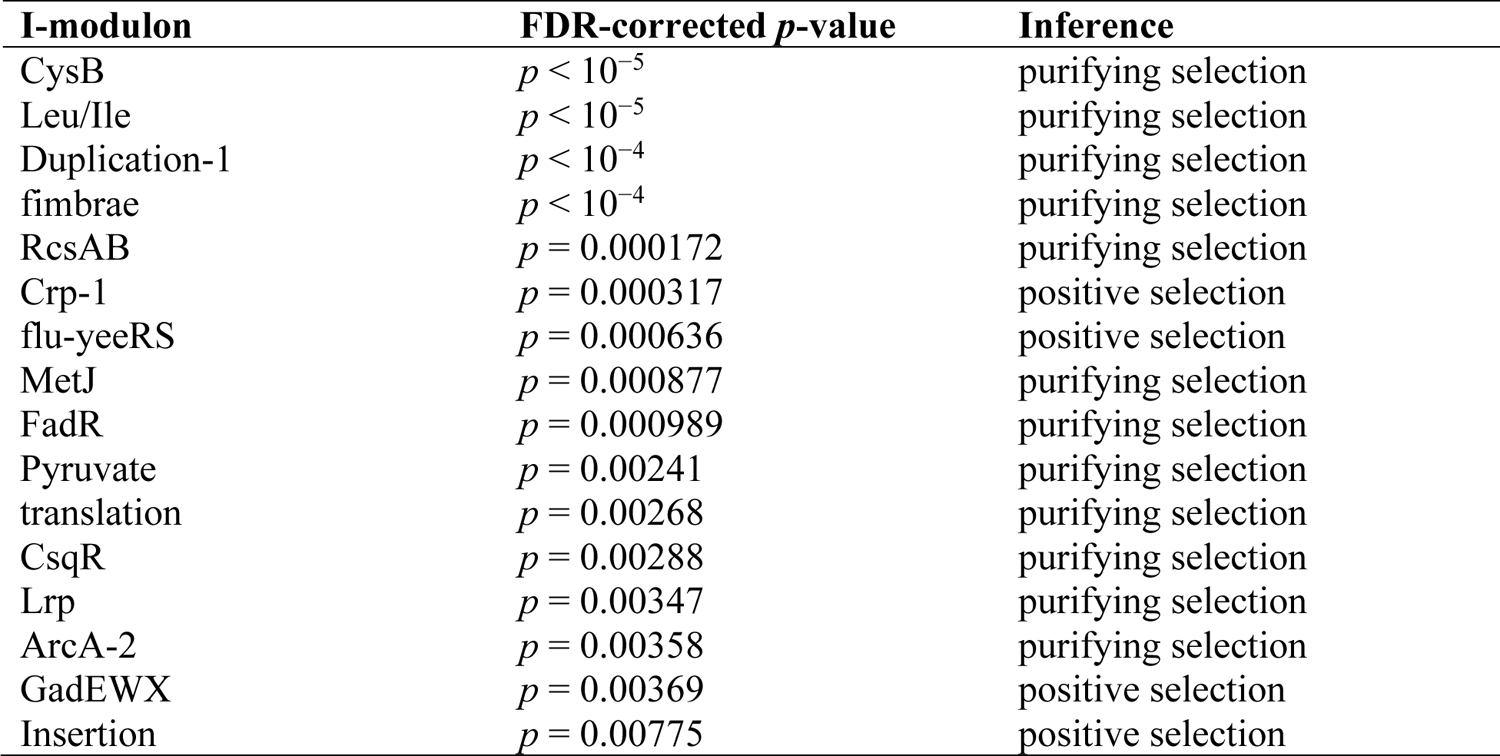
I-modulons showing evidence of selection across all LTEE populations based on the Poisson method, at a FDR-corrected *p*-value threshold of 0.01.

**Supplementary Figure S1.**
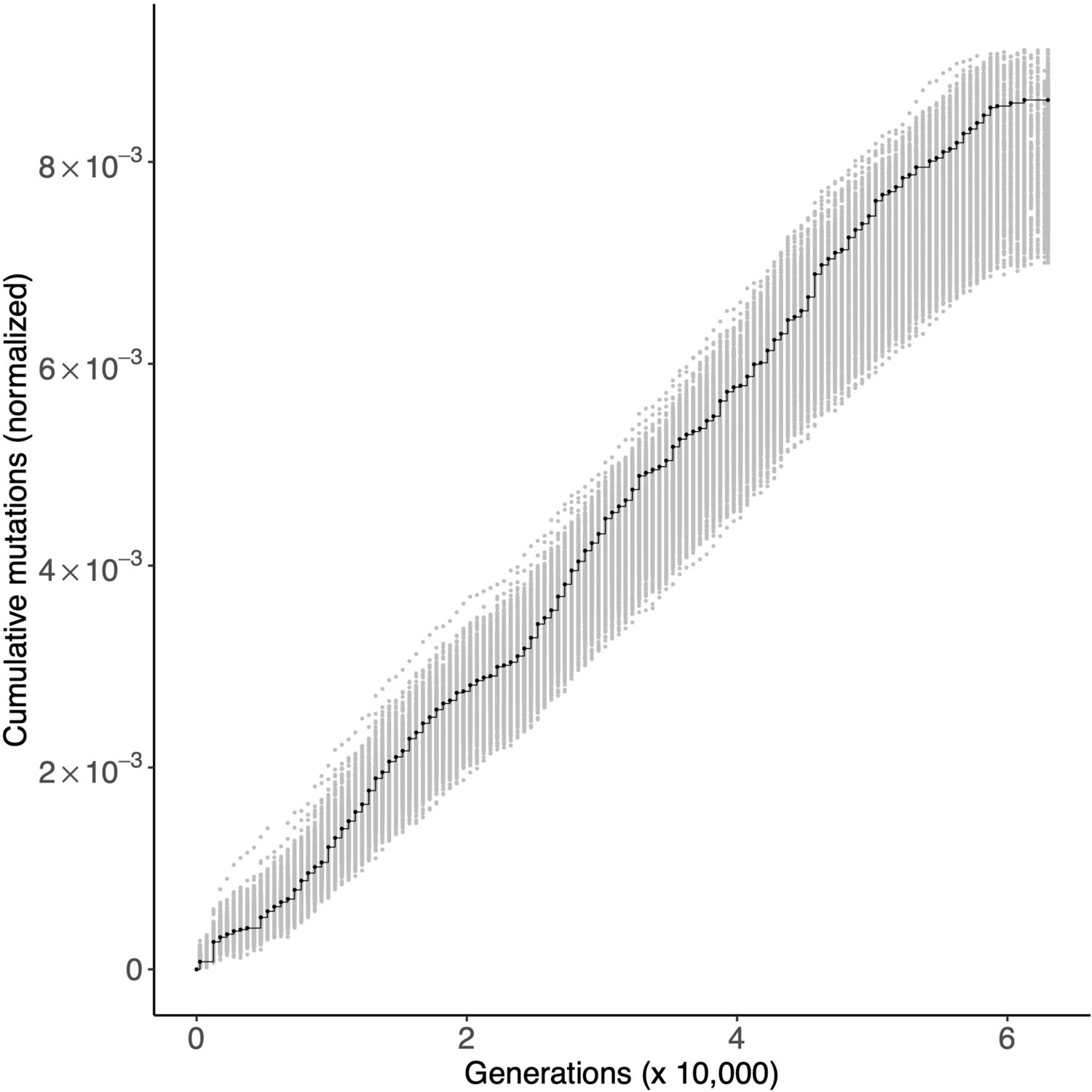
Gold-standard genes under relaxed selection. Each panel shows the cumulative number of mutations in the gold-standard neutral genes (black) in all 12 LTEE populations. For comparison, random sets of 57 genes were sampled 1,000 times, and the cumulative number of mutations in those random gene sets, normalized by gene length, were calculated. The middle 95% of this null distribution is shown in gray, in order to show a two-tailed statistical comparison of the cumulative number of mutations in the gold-standard neutral gene set to the null distribution at *α*= 0.05.

**Supplementary Figure S2.**
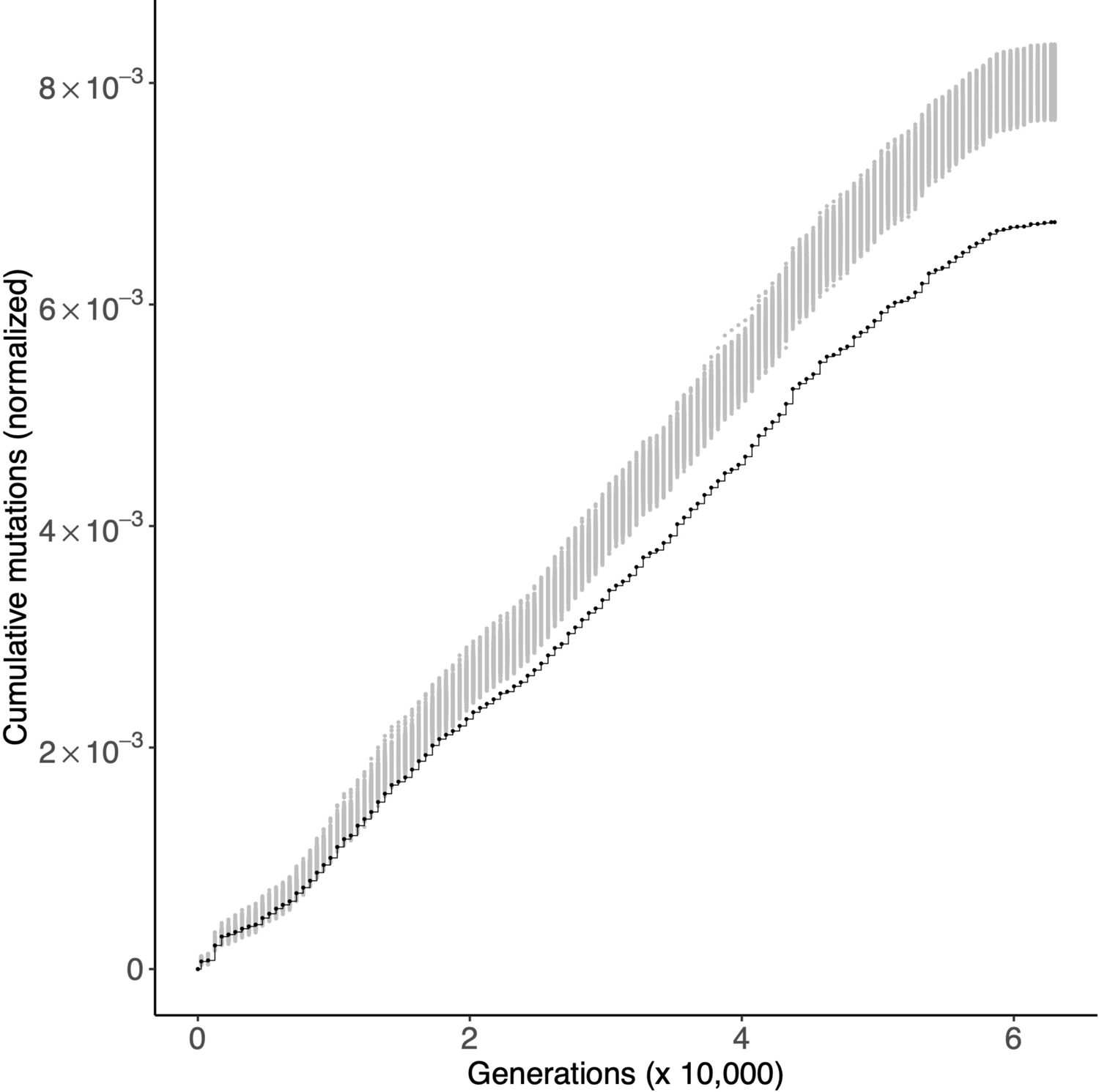
Gold-standard genes under purifying selection. Each panel shows the cumulative number of mutations in the gold standard set of genes under purifying selection (black) in all 12 LTEE populations. For comparison, random sets of 490 genes were sampled 1,000 times, and the cumulative number of mutations in those random gene sets, normalized by gene length, were calculated. The middle 95% of this null distribution is shown in gray.

**Supplementary Figure S3.**
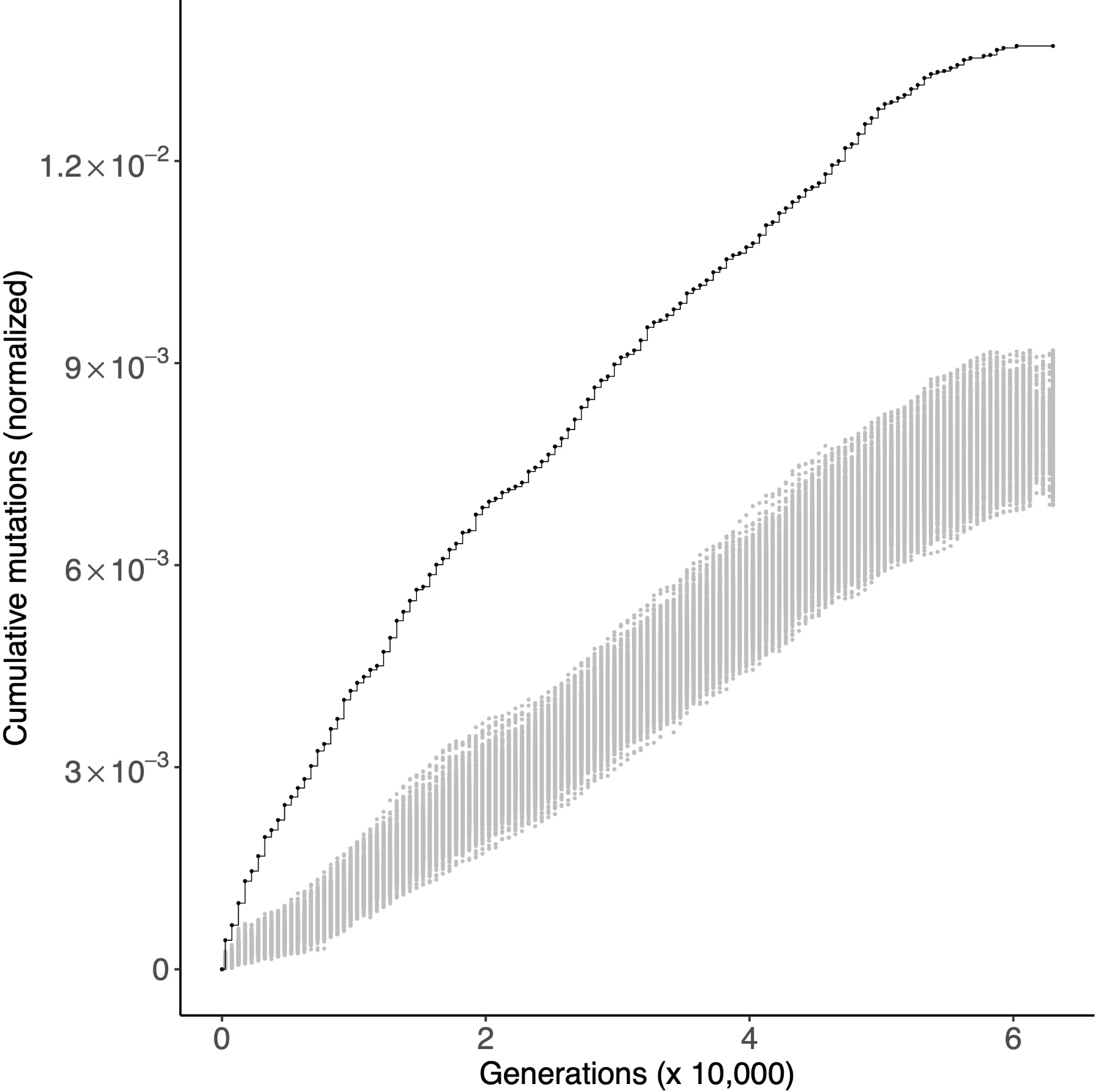
Gold-standard genes under positive selection. Each panel shows the cumulative number of mutations in the gold standard set of genes under positive selection (black) in all 12 LTEE populations. These genes are the 50 genes with the strongest signal of parallel evolution in clones isolated from the six nonmutator populations, as scored by Tenaillon et al. (2016). For comparison, random sets of 50 genes were sampled 1,000 times, and the cumulative number of mutations in those random gene sets, normalized by gene length, were calculated. The middle 95% of this null distribution is shown in gray.

**Supplementary Figure S4.**
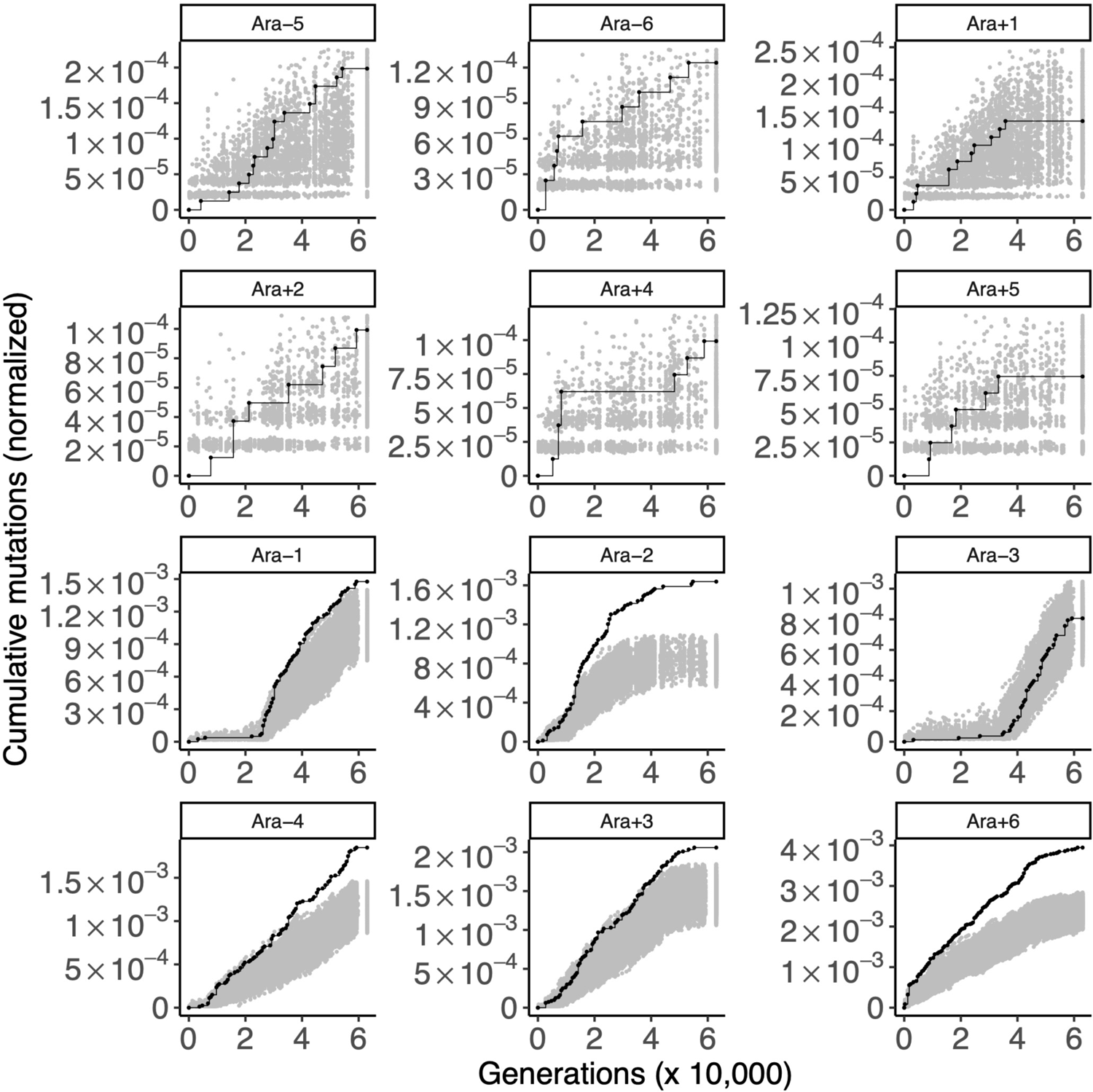
Genes evolving in parallel in hypermutator populations. Each panel shows the cumulative number of mutations in the 50 genes with the strongest signal of parallel evolution in clones isolated from the six hypermutator populations, as scored by Tenaillon et al. (2016), in the 12 LTEE populations. These curves are drawn in black. For comparison, random sets of 50 genes were sampled 1,000 times, and the cumulative number of mutations in those random gene sets, normalized by gene length, were calculated. The middle 95% of this null distribution is shown in gray. The top six panels are the nonmutator populations and the bottom six panels are the hypermutator populations.

**Supplementary Figure S5.**
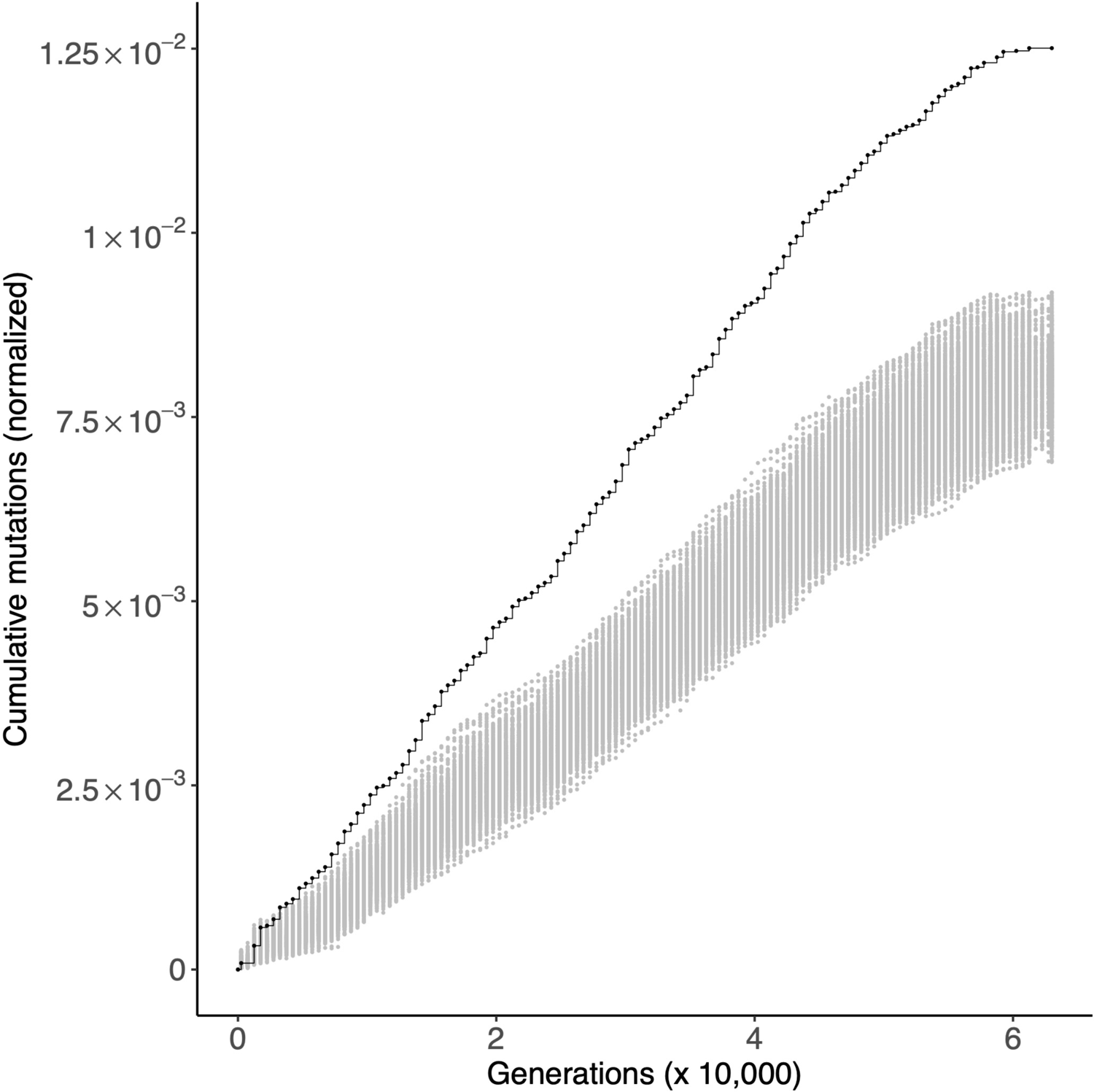
Genes evolving in parallel in hypermutator populations. Each panel shows the cumulative number of mutations in the 50 genes with the strongest signal of parallel evolution in clones isolated from the six hypermutator populations, as scored by Tenaillon et al. (2016), in all 12 LTEE populations. These curves are drawn in black. For comparison, random sets of 50 genes were sampled 1,000 times, and the cumulative number of mutations in those random gene sets, normalized by gene length, were calculated. The middle 95% of this null distribution is shown in gray.

**Supplementary Figure S6.**
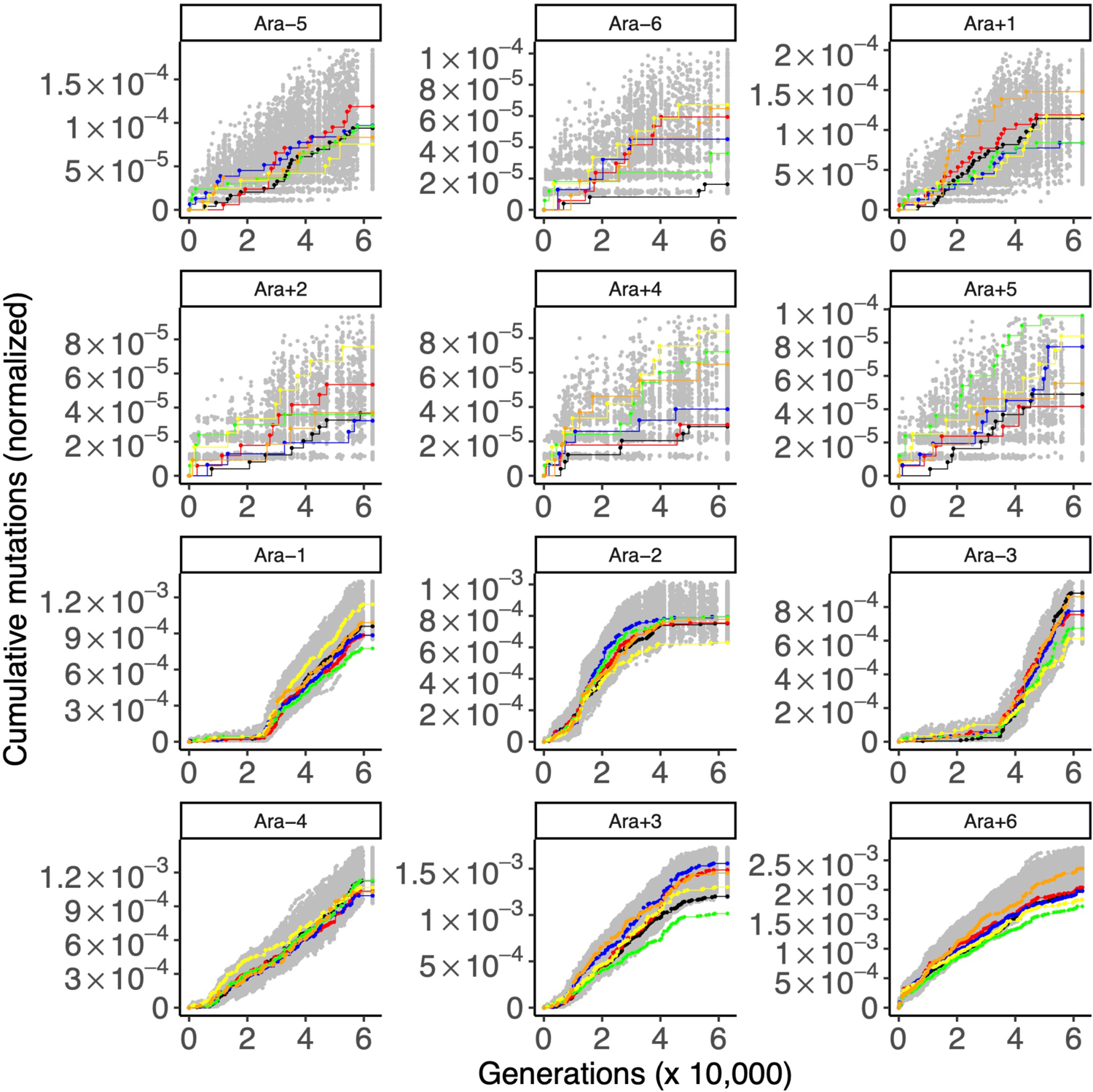
Purifying selection on the proteome U-sector. Each panel shows the cumulative number of mutations in the A-sector (black), S-sector (red), O-sector (blue), U-sector (green), R-sector (yellow), and C-sector (orange) in the 12 LTEE populations. For comparison, random sets of 92 genes (i.e. the smallest proteome sector cardinality) were sampled 1,000 times, and the cumulative number of mutations in those random gene sets, normalized by gene length, were calculated. The middle 95% of this null distribution is shown in gray. The top six panels are the nonmutator populations and the bottom six panels are the hypermutator populations.

**Supplementary Figure S7.**
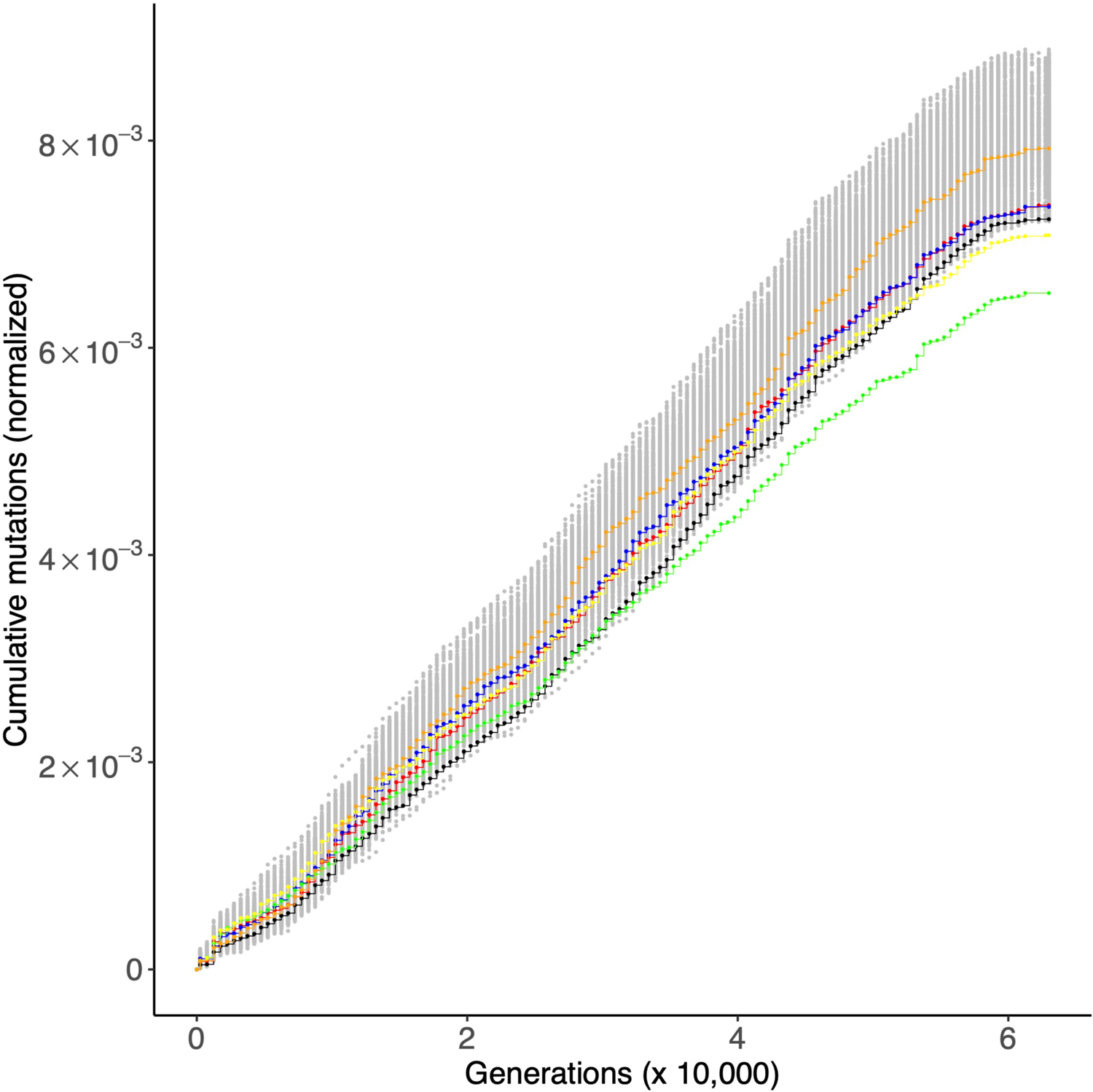
Purifying selection on the proteome U-sector and R-sector. Each panel shows the cumulative number of mutations in the A-sector (black), S-sector (red), O-sector (blue), U-sector (green), R-sector (yellow), and C-sector (orange) in all 12 LTEE populations. For comparison, random sets of 92 genes (i.e. the smallest proteome sector cardinality) were sampled 1,000 times, and the cumulative number of mutations in those random gene sets, normalized by gene length, were calculated. The middle 95% of this null distribution is shown in gray.

**Supplementary Figure S8.**
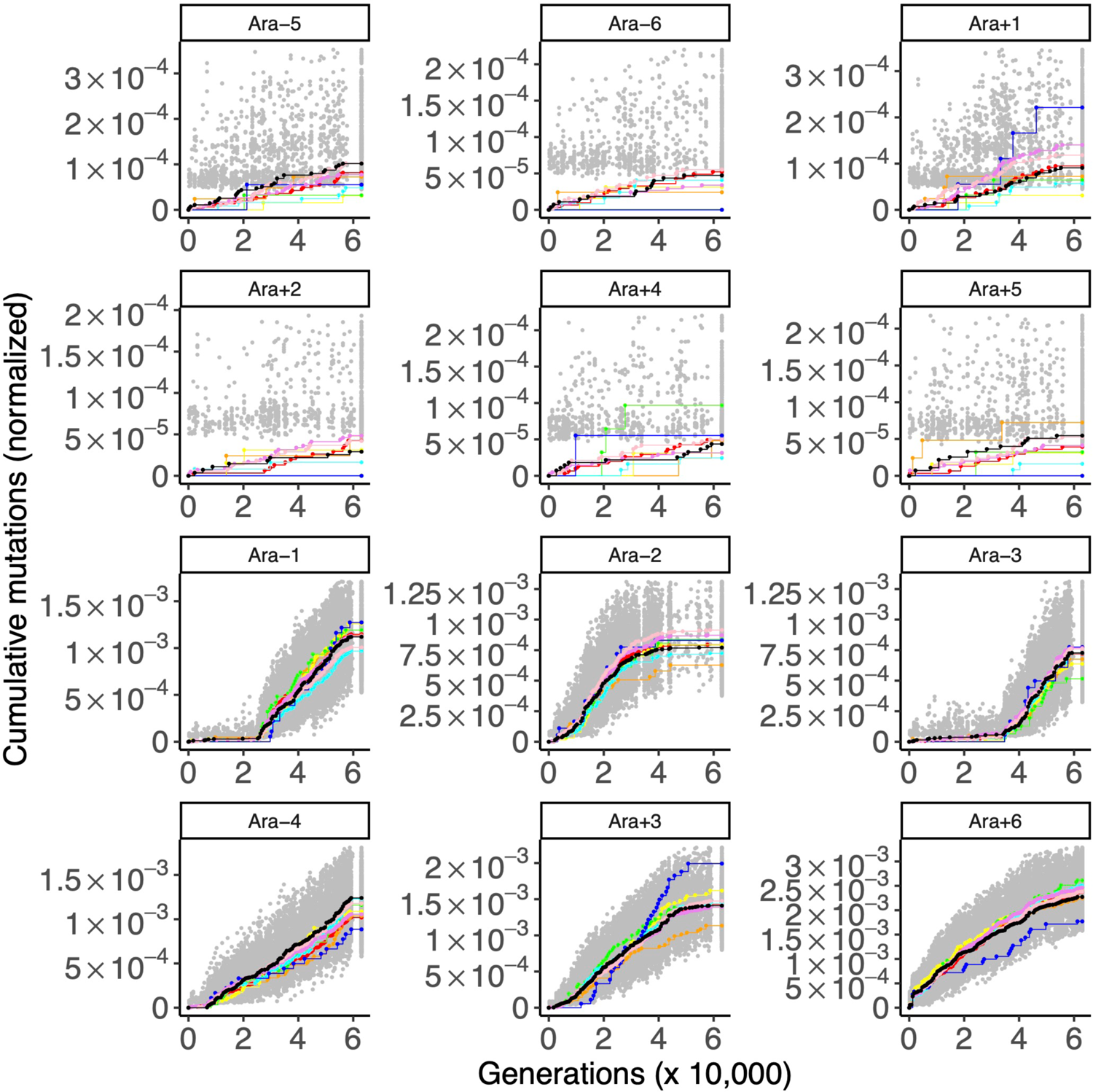
No evidence of selection on *E. coli* eigengenes. Each panel shows the cumulative number of mutations in eigengene 1 (red), eigengene 2 (orange), eigengene 3 (yellow), eigengene 4 (green), eigengene 5 (cyan), eigengene 6 (blue), eigengene 7 (violet), eigengene 8 (pink), and eigengene 9 (black) in the 12 LTEE populations. For comparison, random sets of 15 genes (i.e. the smallest eigengene cardinality) were sampled 1,000 times, and the cumulative number of mutations in those random gene sets, normalized by gene length, were calculated. The middle 95% of this null distribution is shown in gray. The top six panels are the nonmutator populations and the bottom six panels are the hypermutator populations.

**Supplementary Figure S9.**
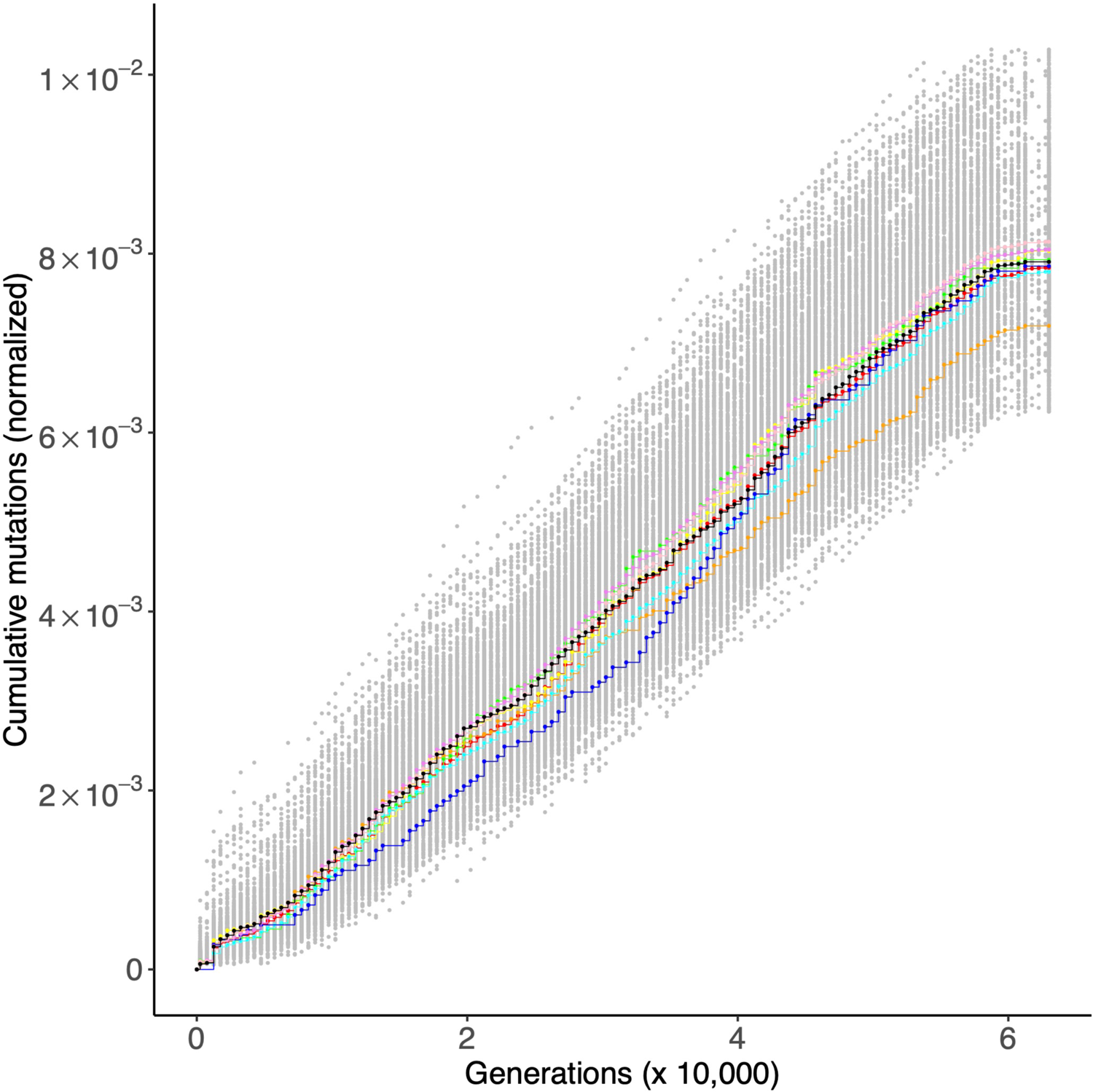
No evidence of selection on *E. coli* eigengenes. Each panel shows the cumulative number of mutations in eigengene 1 (red), eigengene 2 (orange), eigengene 3 (yellow), eigengene 4 (green), eigengene 5 (cyan), eigengene 6 (blue), eigengene 7 (violet), eigengene 8 (pink), and eigengene 9 (black) in the 12 LTEE populations. For comparison, random sets of 15 genes (i.e. the smallest eigengene cardinality) were sampled 1,000 times, and the cumulative number of mutations in those random gene sets, normalized by gene length, were calculated.

**Supplementary Figure S10.**
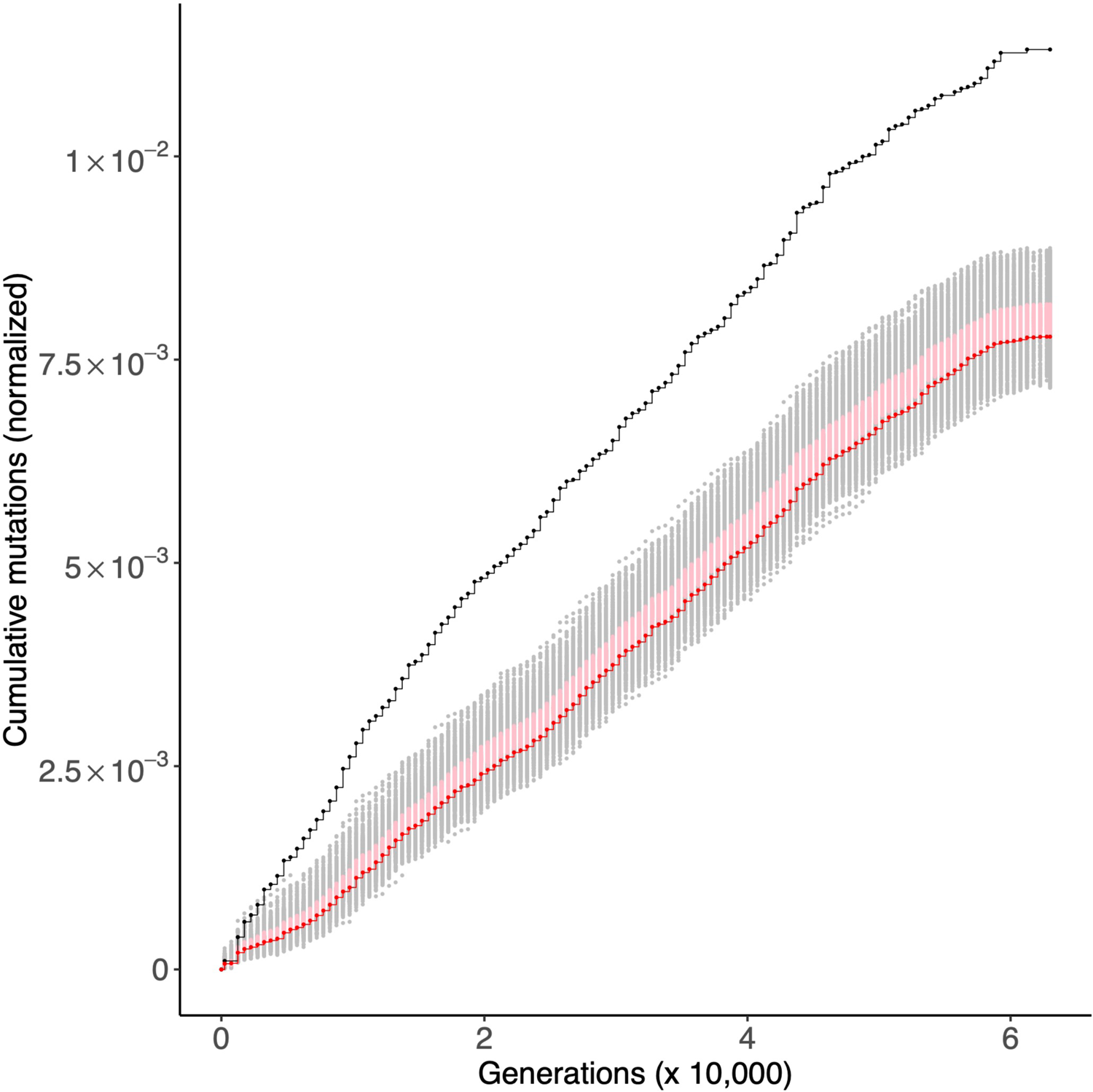
I-modulon regulators evolve under strong positive selection, while genes regulated within I-modulon evolve under purifying selection. Each panel shows the cumulative number of mutations in 70 I-modulon regulators (black) and in 1,394 genes regulated by I-modulon regulators (red) in the 12 LTEE populations. For comparison to the I-modulon regulators, random sets of 70 genes were sampled 1,000 times, and the cumulative number of mutations in those random gene sets, normalized by gene length, were calculated. The middle 95% of this null distribution is shown in gray. A similar procedure was used with random sets of 1,394 genes to make a comparable distribution in pink to compare to the genes regulated by I-modulon regulators.

**Supplementary Figure S11.**
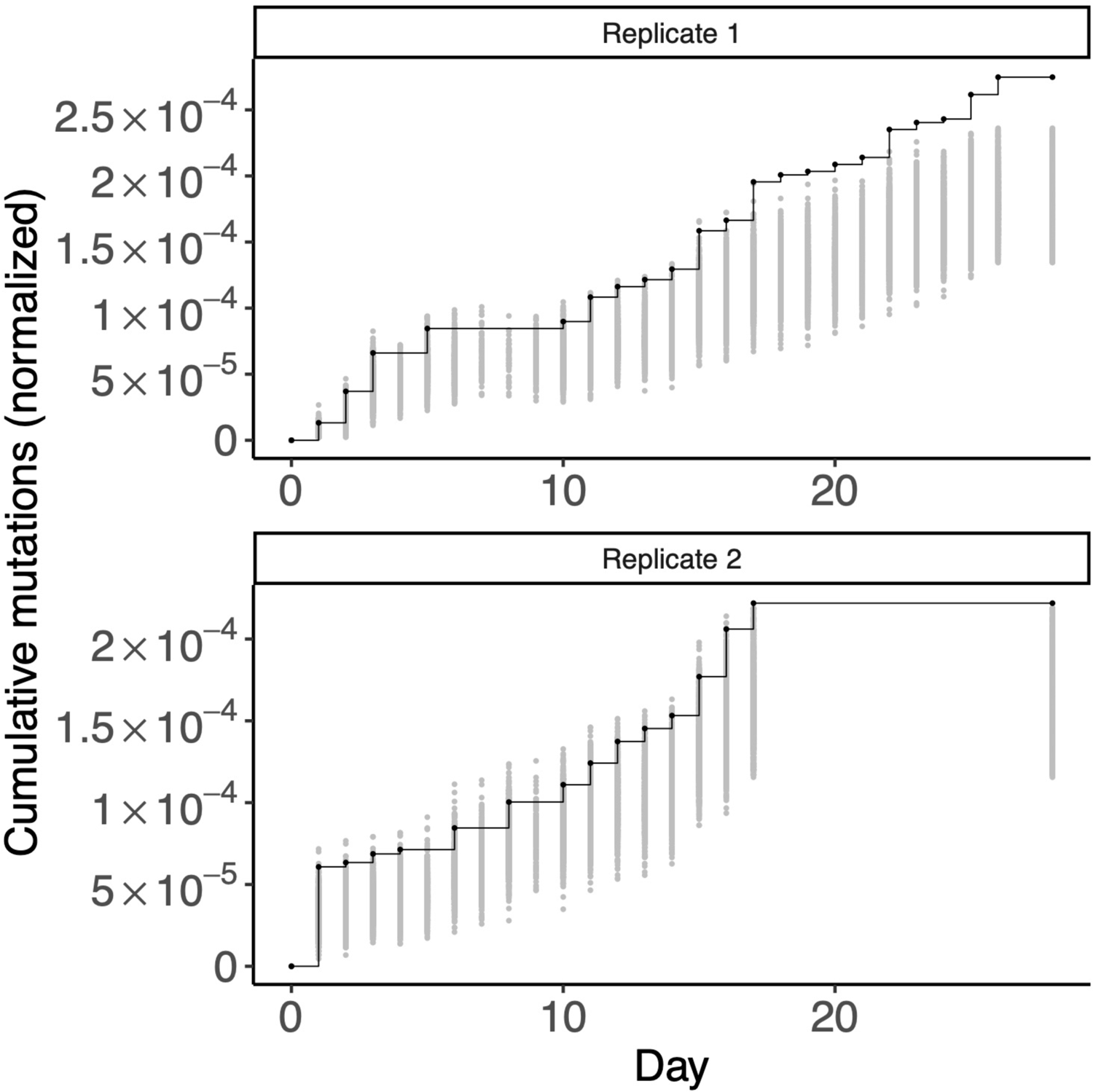
Regulatory genes are under positive selection in hypermutator *Pseudomonas aeruginosa* populations evolving under subinhibitory concentrations of colistin. Each panel shows the cumulative number of mutations in 424 annotated regulatory genes (black), in two replicate populations of hypermutator *P. aeruginosa*. For comparison, random sets of 424 genes were sampled 1,000 times, and the cumulative number of mutations in those random gene sets, normalized by gene length, were calculated. The middle 95% of this null distribution is shown in gray.

## REFERENCES

1. Jeggo PA, Pearl LH, Carr AM. DNA repair, genome stability and cancer: a historical perspective. Nature Reviews Cancer. 2016;16(1):35.

2. Loeb LA. A mutator phenotype in cancer. Cancer research. 2001;61(8):3230–9.

3. Modrich P. Mismatch repair, genetic stability, and cancer. Science. 1994;266(5193):1959-61.

4. Feliziani S, Marvig RL, Lujan AM, Moyano AJ, Di Rienzo JA, Johansen HK, et al. Coexistence and within-host evolution of diversified lineages of hypermutable Pseudomonas aeruginosa in long-term cystic fibrosis infections. PLoS Genet. 2014;10(10):e1004651.

5. Ford CB, Shah RR, Maeda MK, Gagneux S, Murray MB, Cohen T, et al. Mycobacterium tuberculosis mutation rate estimates from different lineages predict substantial differences in the emergence of drug-resistant tuberculosis. Nature genetics. 2013;45(7):784–90.

6. Couce A, Caudwell LV, Feinauer C, Hindré T, Feugeas J-P, Weigt M, et al. Mutator genomes decay, despite sustained fitness gains, in a long-term experiment with bacteria. Proceedings of the National Academy of Sciences. 2017;114(43):E9026–E35.

7. Schiffels S, Szöllősi GJ, Mustonen V, Lässig M. Emergent neutrality in adaptive asexual evolution. Genetics. 2011;189(4):1361–75.

8. Mehta HH, Prater AG, Beabout K, Elworth RA, Karavis M, Gibbons HS, et al. The essential role of hypermutation in rapid adaptation to antibiotic stress. Antimicrobial agents and chemotherapy. 2019;63(7):e00744–19.

9. Hammerstrom TG, Beabout K, Clements TP, Saxer G, Shamoo Y. Acinetobacter baumannii repeatedly evolves a hypermutator phenotype in response to tigecycline that effectively surveys evolutionary trajectories to resistance. PloS one. 2015;10(10):e0140489.

10. Lenski RE, Rose MR, Simpson SC, Tadler SC. Long-term experimental evolution in Escherichia coli. I. Adaptation and divergence during 2,000 generations. The American Naturalist. 1991;138(6):1315-41.

11. Lenski RE. Experimental evolution and the dynamics of adaptation and genome evolution in microbial populations. The ISME journal. 2017;11(10):2181–94.

12. Barrick JE, Lenski RE. Genome-wide mutational diversity in an evolving population of Escherichia coli. Cold Spring Harbor symposia on quantitative biology. 2009;74:119–29.

13. Barrick JE, Yu DS, Yoon SH, Jeong H, Oh TK, Schneider D, et al. Genome evolution and adaptation in a long-term experiment with Escherichia coli. Nature. 2009;461(7268):1243-7.

14. Maddamsetti R, Lenski RE, Barrick JE. Adaptation, clonal interference, and frequency-dependent interactions in a long-term evolution experiment with Escherichia coli. Genetics. 2015;200(2):619–31.

15. Tenaillon O, Barrick JE, Ribeck N, Deatherage DE, Blanchard JL, Dasgupta A, et al. Tempo and mode of genome evolution in a 50,000-generation experiment. Nature. 2016;536(7615):165-70.

16. Good BH, McDonald MJ, Barrick JE, Lenski RE, Desai MM. The dynamics of molecular evolution over 60,000 generations. Nature. 2017;551(7678):45-50.

17. Vasi F, Travisano M, Lenski RE. Long-term experimental evolution in Escherichia coli. II. Changes in life-history traits during adaptation to a seasonal environment. The american naturalist. 1994;144(3):432–56.

18. Lenski RE, Mongold JA, Sniegowski PD, Travisano M, Vasi F, Gerrish PJ, et al. Evolution of competitive fitness in experimental populations of E. coli: what makes one genotype a better competitor than another? Antonie van Leeuwenhoek. 1998;73(1):35–47.

19. Elena SF, Cooper VS, Lenski RE. Punctuated evolution caused by selection of rare beneficial mutations. Science. 1996;272(5269):1802-4.

20. Wiser MJ, Ribeck N, Lenski RE. Long-term dynamics of adaptation in asexual populations. Science. 2013;342(6164):1364-7.

21. Grant NA, Maddamsetti R, Lenski RE. Maintenance of metabolic plasticity despite relaxed selection in a long-term evolution experiment with Escherichia coli. The American Naturalist. 2021;0(ja):null. doi: 10.1086/714530.

22. Grant NA, Magid AA, Franklin J, Dufour Y, Lenski RE. Changes in Cell Size and Shape During 50,000 Generations of Experimental Evolution with Escherichia coli. Journal of Bacteriology. 2021;203(10).

23. Maddamsetti R, Hatcher PJ, Green AG, Williams BL, Marks DS, Lenski RE. Core genes evolve rapidly in the long-term evolution experiment with Escherichia coli. Genome biology and evolution. 2017;9(4):1072–83.

24. Maddamsetti R. Universal constraints on protein evolution in the long-term evolution experiment with Escherichia coli. Genome Biology and Evolution. 2021. doi: 10.1093/gbe/evab070.

25. Maddamsetti R. Selection maintains protein interactome resilience in the long-term evolution experiment with Escherichia coli. Genome Biology and Evolution. 2021. doi: 10.1093/gbe/evab074.

26. Hui S, Silverman JM, Chen SS, Erickson DW, Basan M, Wang J, et al. Quantitative proteomic analysis reveals a simple strategy of global resource allocation in bacteria. Molecular systems biology. 2015;11(2).

27. Sastry AV, Gao Y, Szubin R, Hefner Y, Xu S, Kim D, et al. The Escherichia coli transcriptome mostly consists of independently regulated modules. Nature communications. 2019;10(1):1–14.

28. Wytock TP, Motter AE. Predicting growth rate from gene expression. Proceedings of the National Academy of Sciences. 2019;116(2):367–72.

29. Izutsu M, Lake DM, Matson ZW, Dodson JP, Lenski RE. Effects of periodic bottlenecks on the dynamics of adaptive evolution in microbial populations. bioRxiv. 2021.

30. Papadopoulos D, Schneider D, Meier-Eiss J, Arber W, Lenski RE, Blot M. Genomic evolution during a 10,000-generation experiment with bacteria. Proceedings of the National Academy of Sciences. 1999;96(7):3807–12.

31. Maddamsetti R, Grant NA. Divergent evolution of mutation rates and biases in the long-term evolution experiment with Escherichia coli. Genome Biology and Evolution. 2020. doi: 10.1093/gbe/evaa178.

32. Consuegra J, Gaffé J, Lenski RE, Hindré T, Barrick JE, Tenaillon O, et al. Insertion-sequence-mediated mutations both promote and constrain evolvability during a long-term experiment with bacteria. Nature communications. 2021;12(1):1–12.

33. Turner CB, Blount ZD, Mitchell DH, Lenski RE. Evolution and coexistence in response to a key innovation in a long-term evolution experiment with Escherichia coli. bioRxiv. 2015:020958.

34. Kinnersley M, Schwartz K, Yang D-D, Sherlock G, Rosenzweig F. Evolutionary dynamics and structural consequences of de novo beneficial mutations and mutant lineages arising in a constant environment. BMC biology. 2021;19(1):1–21.

35. Lenski RE, Wiser MJ, Ribeck N, Blount ZD, Nahum JR, Morris JJ, et al. Sustained fitness gains and variability in fitness trajectories in the long-term evolution experiment with Escherichia coli. Proceedings of the Royal Society B: Biological Sciences. 2015;282(1821):20152292.

36. Wielgoss S, Barrick JE, Tenaillon O, Wiser MJ, Dittmar WJ, Cruveiller S, et al. Mutation rate dynamics in a bacterial population reflect tension between adaptation and genetic load. Proceedings of the National Academy of Sciences. 2013;110(1):222–7.

37. Rudan M, Schneider D, Warnecke T, Krisko A. RNA chaperones buffer deleterious mutations in E. coli. Elife. 2015;4:e04745.

38. Bailey SF, Guo Q, Bataillon T. Identifying drivers of parallel evolution: A regression model approach. Genome biology and evolution. 2018;10(10):2801–12.

39. Foster PL, Hanson AJ, Lee H, Popodi EM, Tang H. On the mutational topology of the bacterial genome. G3: Genes, Genomes, Genetics. 2013;3(3):399-407.

40. Niccum BA, Lee H, MohammedIsmail W, Tang H, Foster PL. The Symmetrical Wave Pattern of Base-Pair Substitution Rates across the Escherichia coli Chromosome Has Multiple Causes. mBio. 2019;10(4).

41. Quandt EM, Gollihar J, Blount ZD, Ellington AD, Georgiou G, Barrick JE. Fine-tuning citrate synthase flux potentiates and refines metabolic innovation in the Lenski evolution experiment. Elife. 2015;4:e09696.

42. Zee PC, Mendes-Soares H, Yu YTN, Kraemer SA, Keller H, Ossowski S, et al. A shift from magnitude to sign epistasis during adaptive evolution of a bacterial social trait. Evolution. 2014;68(9):2701–8.

43. Venkataram S, Monasky R, Sikaroodi SH, Kryazhimskiy S, Kacar B. Evolutionary stalling and a limit on the power of natural selection to improve a cellular module. Proceedings of the National Academy of Sciences. 2020;117(31):18582–90.

44. Rodrigues JV, Shakhnovich EI. Adaptation to mutational inactivation of an essential gene converges to an accessible suboptimal fitness peak. Elife. 2019;8.

45. Kaznatcheev A. Computational complexity as an ultimate constraint on evolution. Genetics. 2019;212(1):245–65.

46. Andersson DI, Slechta ES, Roth JR. Evidence that gene amplification underlies adaptive mutability of the bacterial lac operon. Science. 1998;282(5391):1133-5.

47. Hendrickson H, Slechta ES, Bergthorsson U, Andersson DI, Roth JR. Amplification– mutagenesis: evidence that “directed” adaptive mutation and general hypermutability result from growth with a selected gene amplification. Proceedings of the National Academy of Sciences. 2002;99(4):2164–9.

48. Good BH, Desai MM. Evolution of mutation rates in rapidly adapting asexual populations. Genetics. 2016;204(3):1249–66.

49. Philippe N, Crozat E, Lenski RE, Schneider D. Evolution of global regulatory networks during a long-term experiment with Escherichia coli. Bioessays. 2007;29(9):846–60.

50. Chen H, Wu C-I, He X. The genotype–phenotype relationships in the light of natural selection. Molecular biology and evolution. 2018;35(3):525–42.

51. Scott M, Gunderson CW, Mateescu EM, Zhang Z, Hwa T. Interdependence of cell growth and gene expression: origins and consequences. Science. 2010;330(6007):1099-102.

52. Erickson DW, Schink SJ, Patsalo V, Williamson JR, Gerland U, Hwa T. A global resource allocation strategy governs growth transition kinetics of Escherichia coli. Nature. 2017;551(7678):119-23.

53. Klumpp S, Hwa T. Bacterial growth: global effects on gene expression, growth feedback and proteome partition. Current opinion in biotechnology. 2014;28:96–102.

54. Basan M, Hui S, Okano H, Zhang Z, Shen Y, Williamson JR, et al. Overflow metabolism in Escherichia coli results from efficient proteome allocation. Nature. 2015;528(7580):99-104.

55. Mathis RA, Sokol ES, Gupta PB. Identification of Genes under Purifying Selection in Human Cancers. BioRxiv. 2017:129205.

56. Weghorn D, Sunyaev S. Bayesian inference of negative and positive selection in human cancers. Nature genetics. 2017;49(12):1785–8.

57. Patel SJ, Sanjana NE, Kishton RJ, Eidizadeh A, Vodnala SK, Cam M, et al. Identification of essential genes for cancer immunotherapy. Nature. 2017;548(7669):537-42.

58. Li X, Giorgi EE, Marichannegowda MH, Foley B, Xiao C, Kong X-P, et al. Emergence of SARS-CoV-2 through recombination and strong purifying selection. Science Advances. 2020;6(27):eabb9153.

59. Lieberman TD, Michel J-B, Aingaran M, Potter-Bynoe G, Roux D, Davis MR, et al. Parallel bacterial evolution within multiple patients identifies candidate pathogenicity genes. Nature genetics. 2011;43(12):1275–80.

60. Caballero JD, Clark ST, Coburn B, Zhang Y, Wang PW, Donaldson SL, et al. Selective sweeps and parallel pathoadaptation drive Pseudomonas aeruginosa evolution in the cystic fibrosis lung. MBio. 2015;6(5).

61. Chung H, Lieberman TD, Vargas SO, Flett KB, McAdam AJ, Priebe GP, et al. Global and local selection acting on the pathogen Stenotrophomonas maltophilia in the human lung. Nature communications. 2017;8:14078.

62. Cooper VS, Honsa E, Rowe H, Deitrick C, Iverson AR, Whittall JJ, et al. Experimental Evolution in vivo to Identify Selective Pressures During Pneumococcal Colonization. Msystems. 2020;5(3).

63. Deatherage DE, Traverse CC, Wolf LN, Barrick JE. Detecting rare structural variation in evolving microbial populations from new sequence junctions using breseq. Frontiers in genetics. 2015;5:468.

64. Haller BC, Messer PW. SLiM 3: forward genetic simulations beyond the Wright–Fisher model. Molecular biology and evolution. 2019;36(3):632–7.

65. Robert L, Ollion J, Robert J, Song X, Matic I, Elez M. Mutation dynamics and fitness effects followed in single cells. Science. 2018;359(6381):1283-6.

66. Perfeito L, Fernandes L, Mota C, Gordo I. Adaptive mutations in bacteria: high rate and small effects. science. 2007;317(5839):813-5.

