## Supplementary file S1 for "Discovery of positive and purifying selection in metagenomic time series of hypermutator microbial populations"

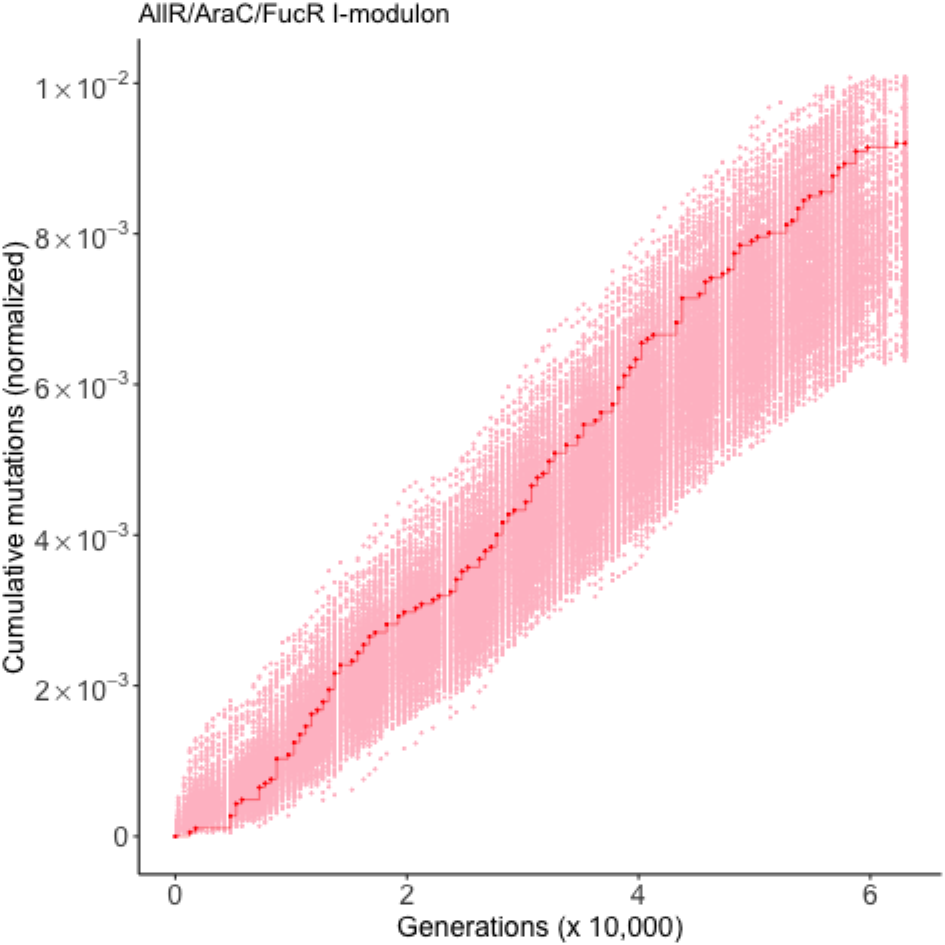
AllR/AraC/FucR I-modulon
Cumulative mutations (normalized)
$1\times10^{-2}$
$8\times10^{-3}$
$6\times10^{-3}$
$4\times10^{-3}$
$2\times10^{-3}$
0
0
2
4
6
Generations (x 10,000)

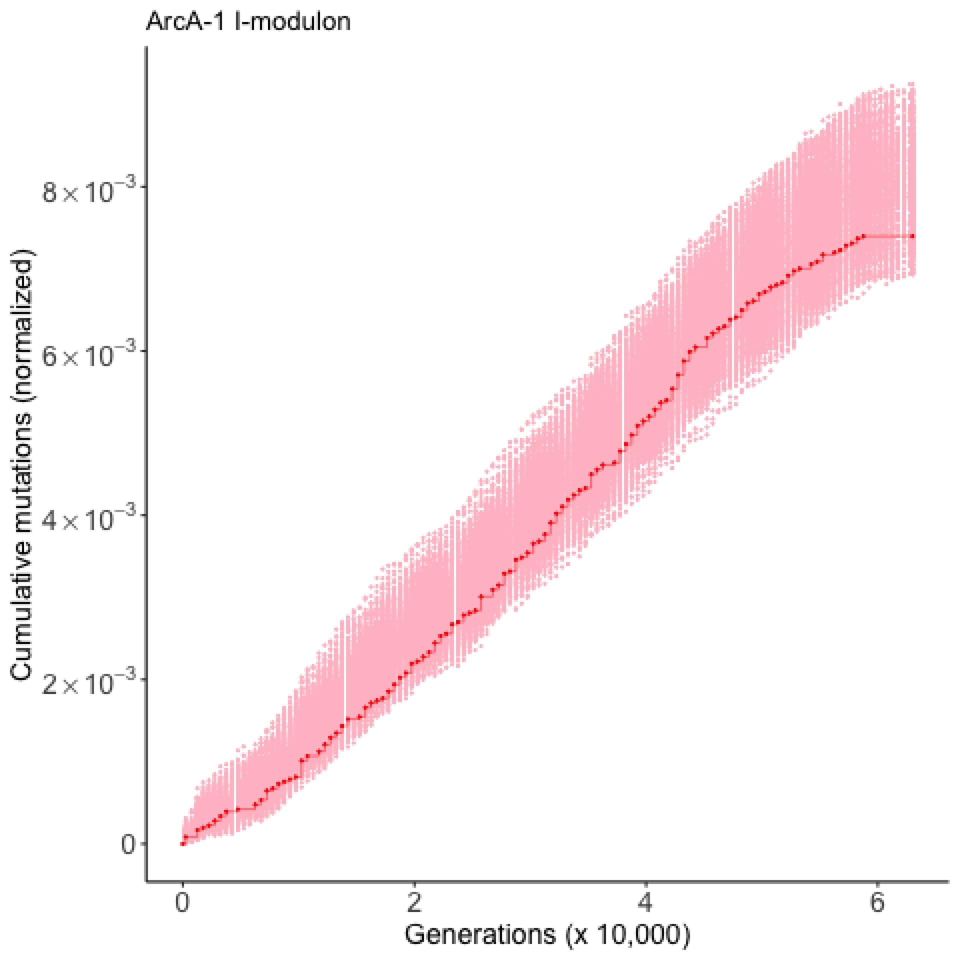

ArcA-1 I-modulon
Cumulative mutations (normalized)
8×10^-3
6×10^-3
4×10^-3
2×10^-3
0
0
2
4
6
Generations (x 10,000)

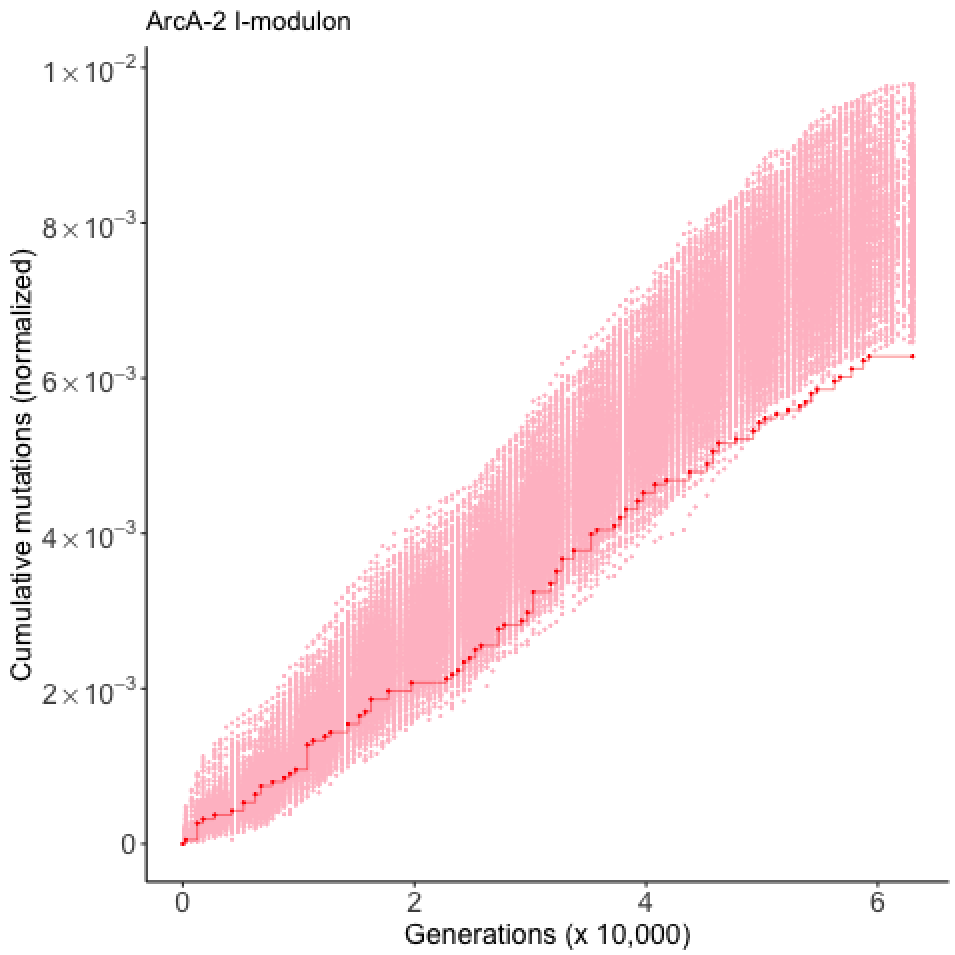
ArcA-2 I-modulon
Cumulative mutations (normalized)
1×10^−2
8×10^−3
6×10^−3
4×10^−3
2×10^−3
0
0
2
4
6
Generations (x 10,000)

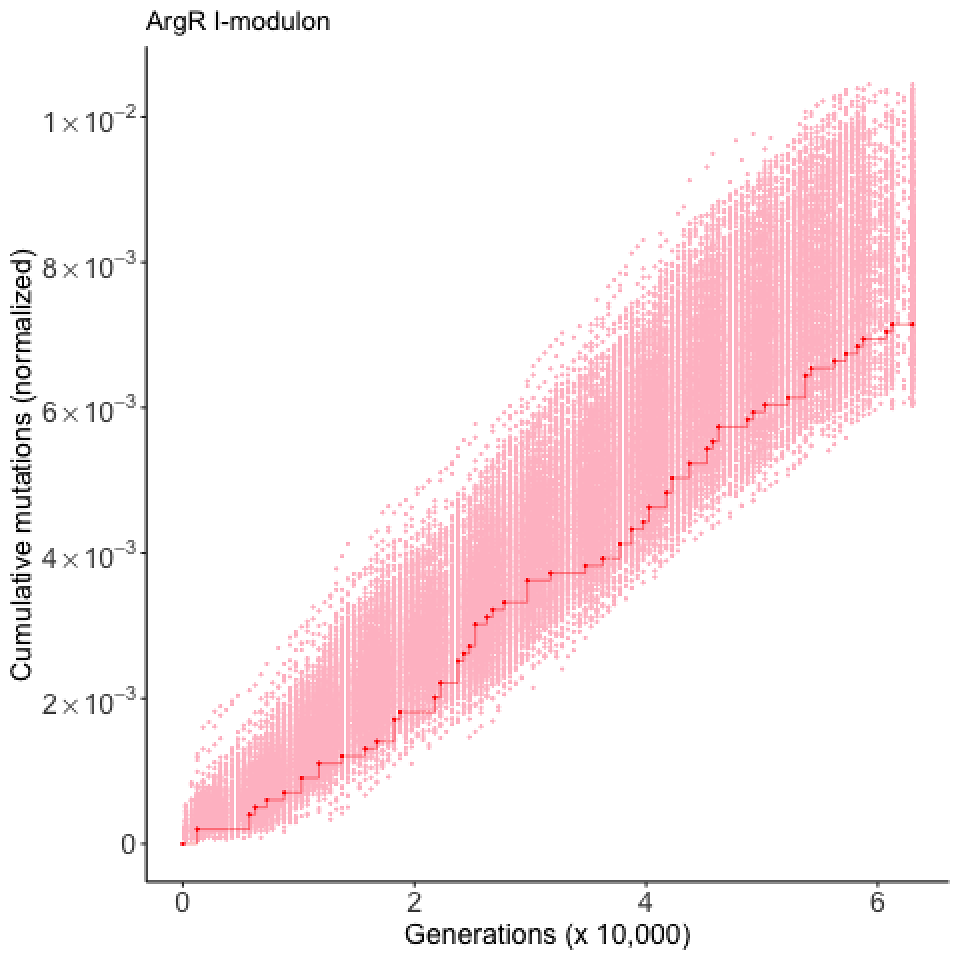
ArgR I-modulon
Cumulative mutations (normalized)
$1\times10^{-2}$
$8\times10^{-3}$
$6\times10^{-3}$
$4\times10^{-3}$
$2\times10^{-3}$
0
0
2
4
6
Generations (x 10,000)

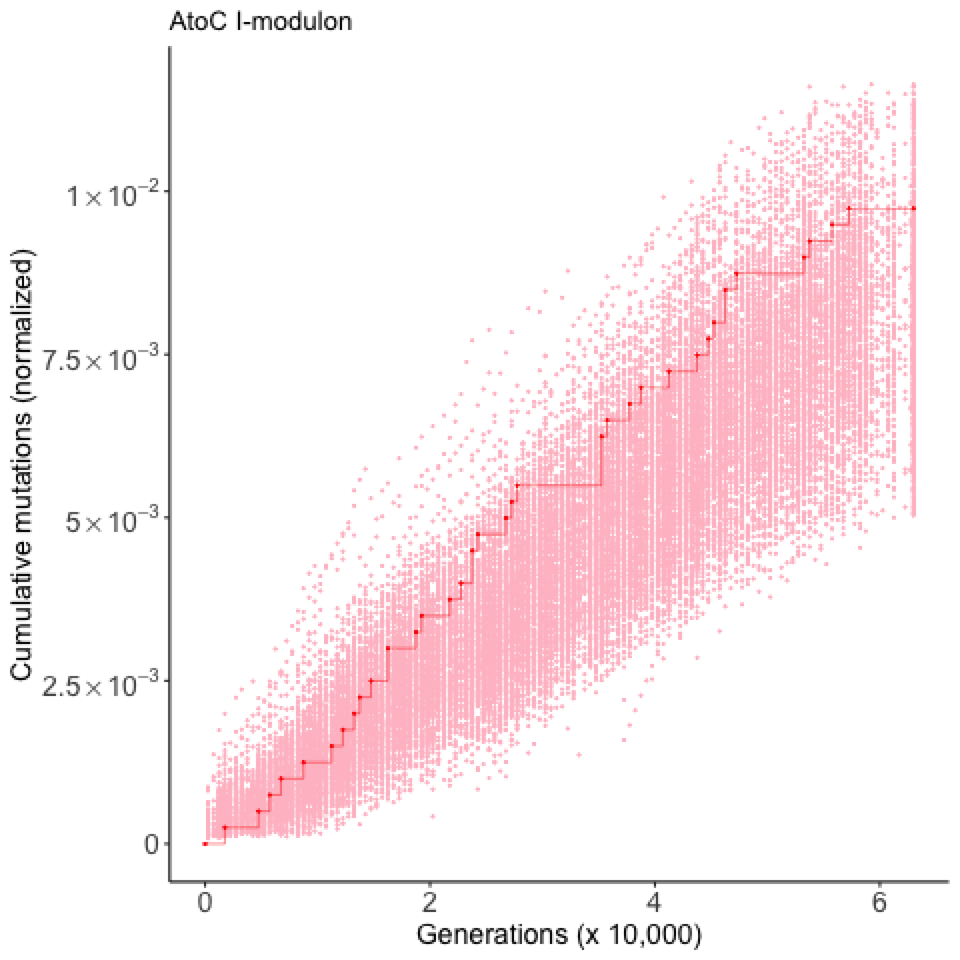

AtoC I-modulon
Cumulative mutations (normalized)
$1\times10^{-2}$
$7.5\times10^{-3}$
$5\times10^{-3}$
$2.5\times10^{-3}$
0
0
2
4
6
Generations (x 10,000)

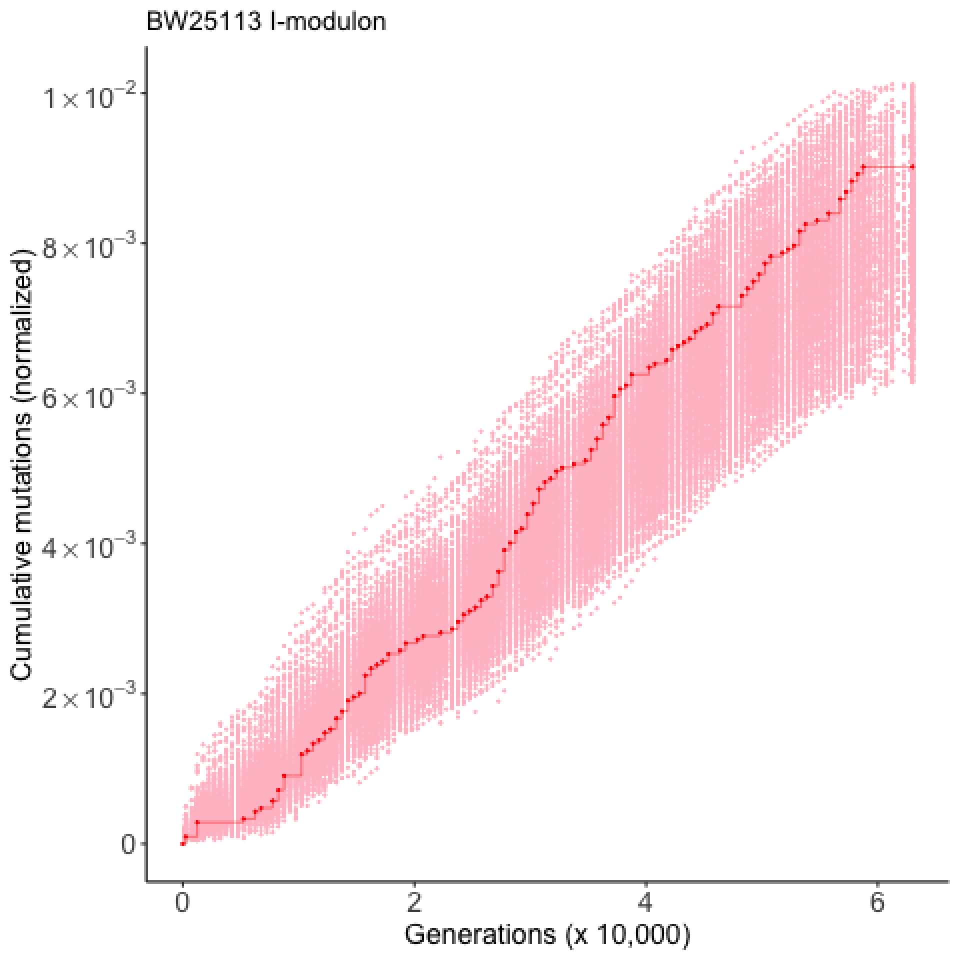

BW25113 I-modulon
Cumulative mutations (normalized)
$1\times10^{-2}$
$8\times10^{-3}$
$6\times10^{-3}$
$4\times10^{-3}$
$2\times10^{-3}$
0
0
2
4
6
Generations (x 10,000)

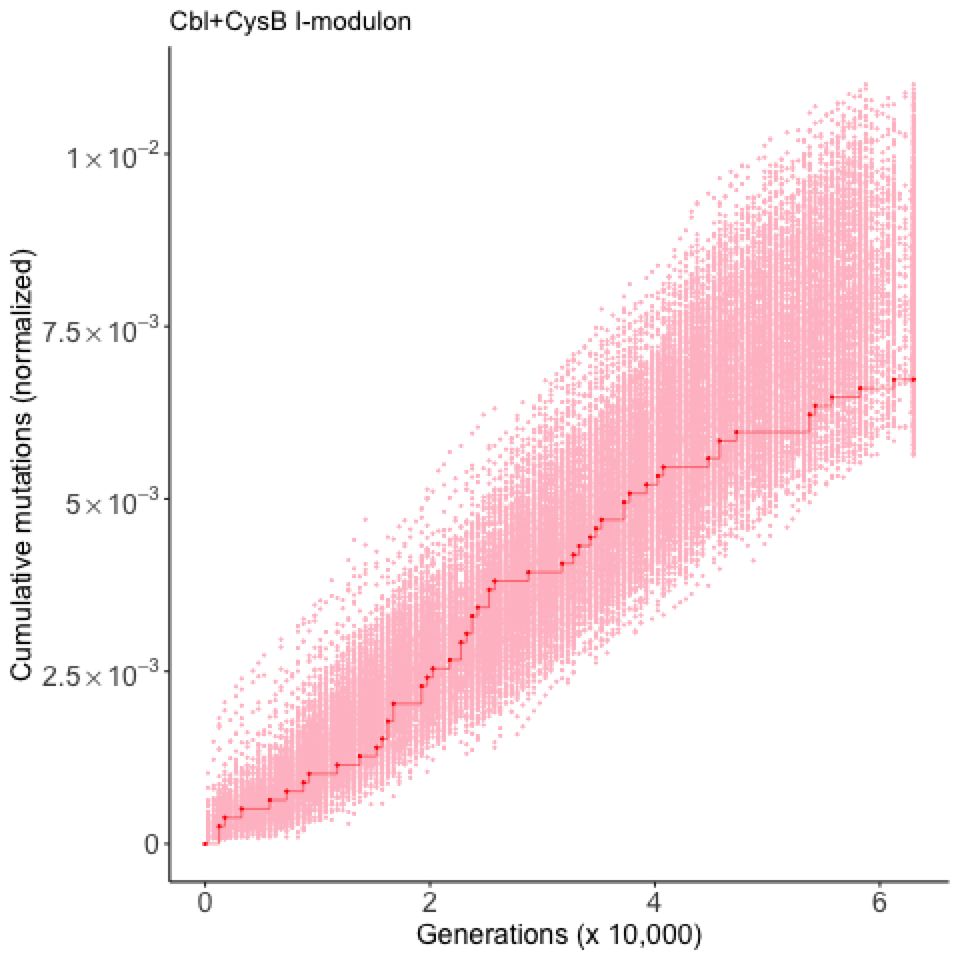

Cbl+CysB I-modulon
Cumulative mutations (normalized)
$1\times10^{-2}$
$7.5\times10^{-3}$
$5\times10^{-3}$
$2.5\times10^{-3}$
0
0
2
4
6
Generations (x 10,000)

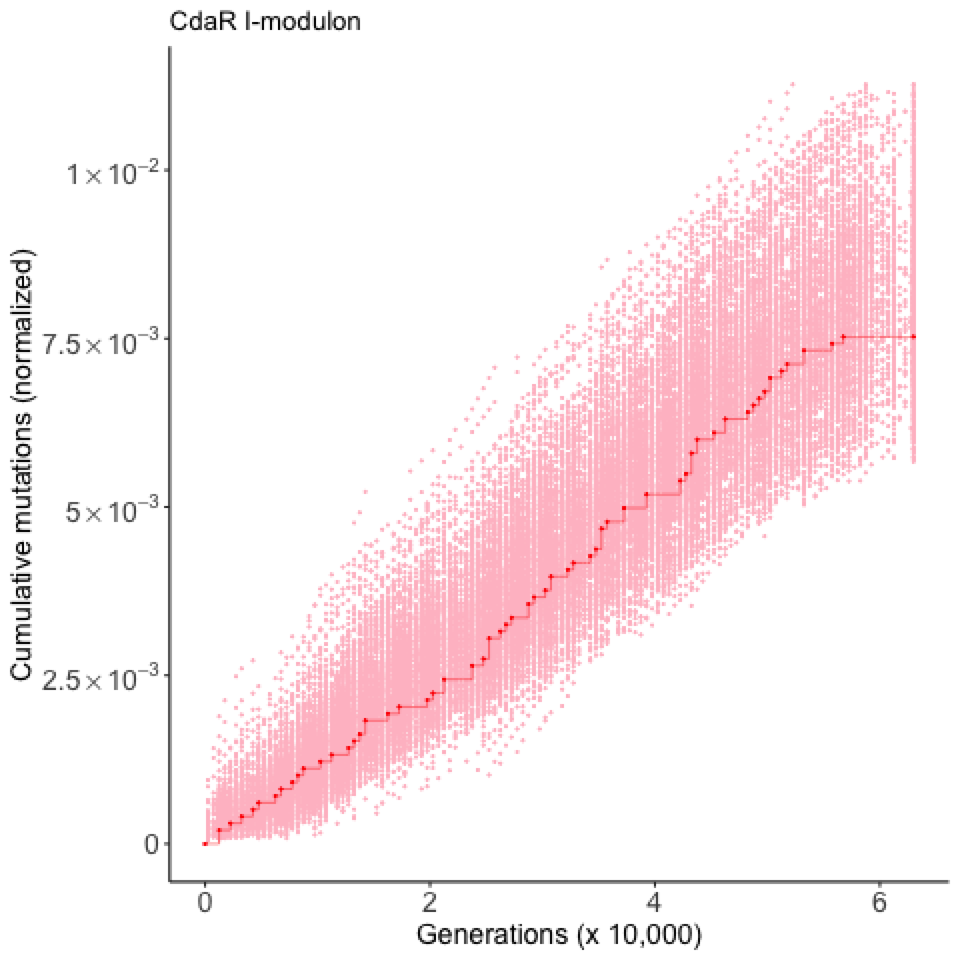
CdaR I-modulon
Cumulative mutations (normalized)
$1\times10^{-2}$
$7.5\times10^{-3}$
$5\times10^{-3}$
$2.5\times10^{-3}$
0
0
2
4
6
Generations (x 10,000)

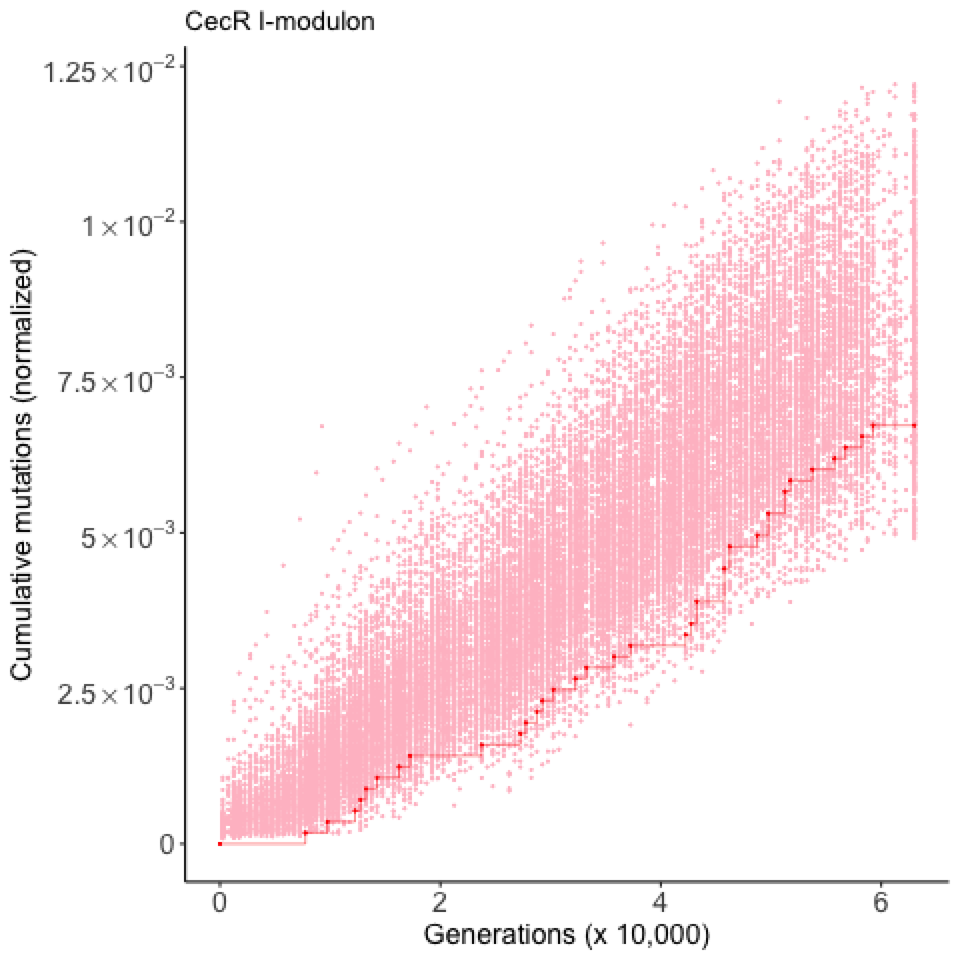

CecR I-modulon
Cumulative mutations (normalized)
1.25×10^-2
1×10^-2
7.5×10^-3
5×10^-3
2.5×10^-3
0
0
2
4
6
Generations (x 10,000)

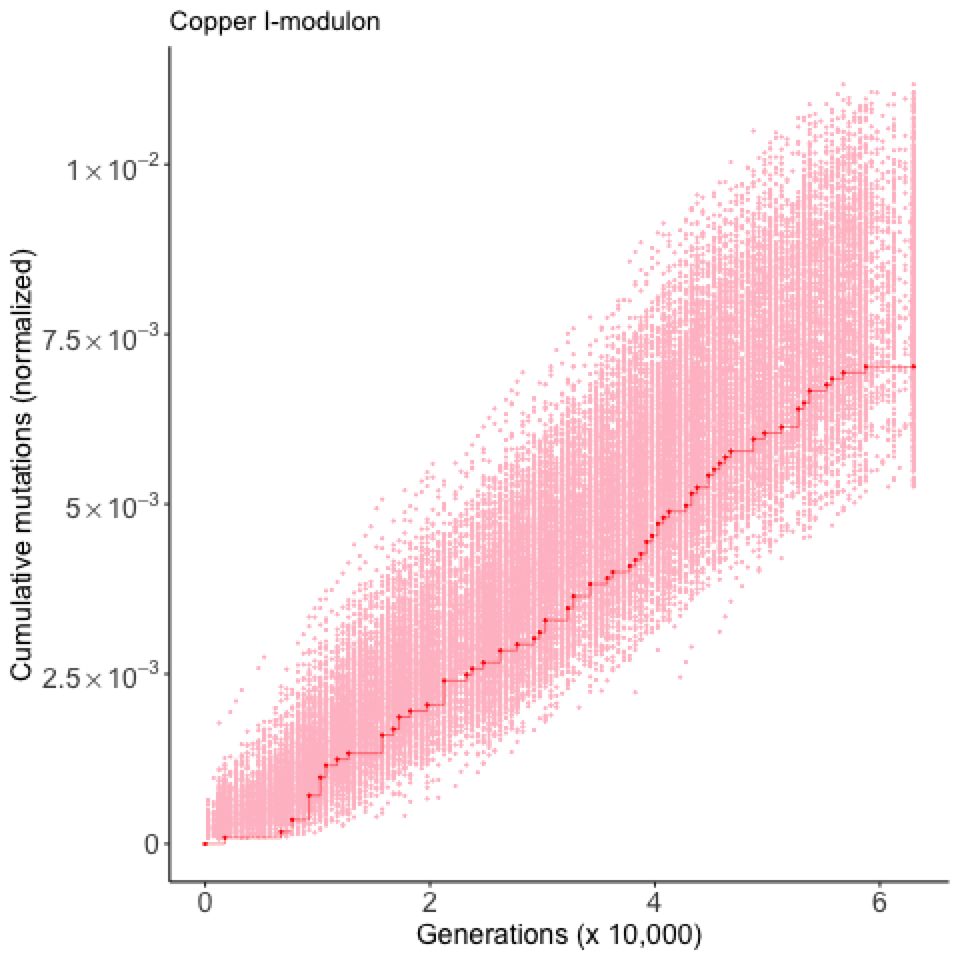

Copper I-modulon
Cumulative mutations (normalized)
1×10^−2
7.5×10^−3
5×10^−3
2.5×10^−3
0
0
2
4
6
Generations (x 10,000)

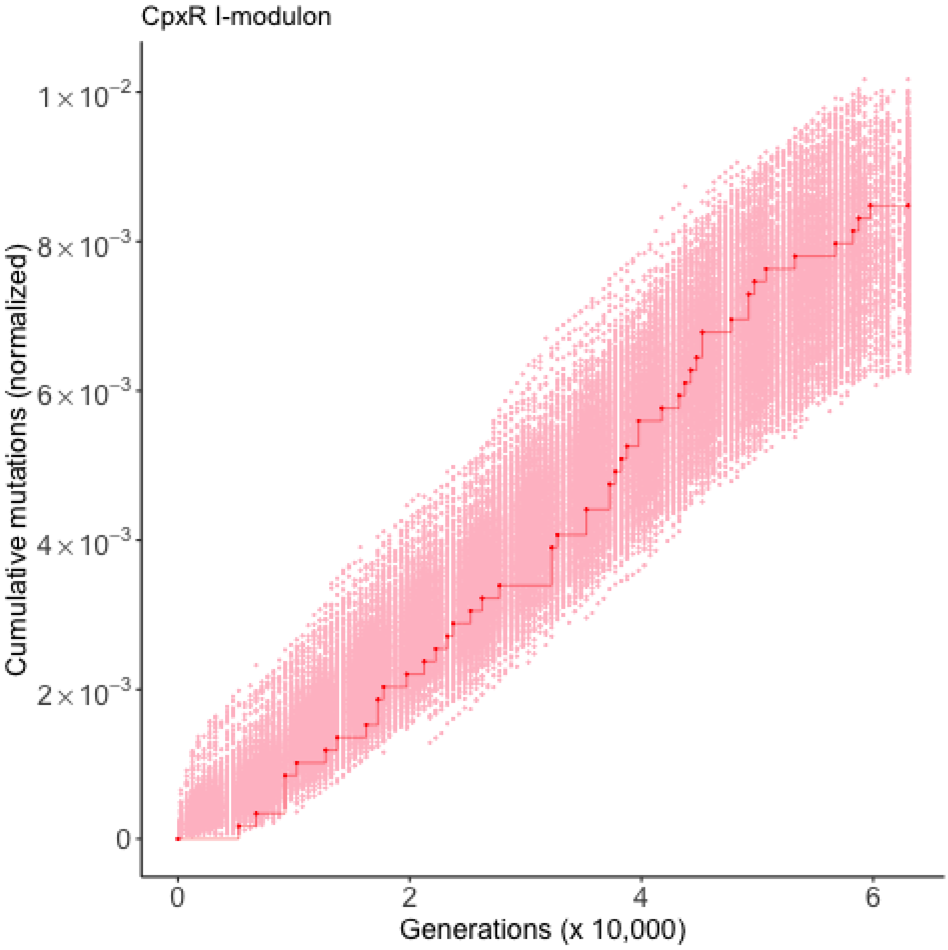

CpxR I-modulon
Cumulative mutations (normalized)
$1\times10^{-2}$
$8\times10^{-3}$
$6\times10^{-3}$
$4\times10^{-3}$
$2\times10^{-3}$
0
0
2
4
6
Generations (x 10,000)

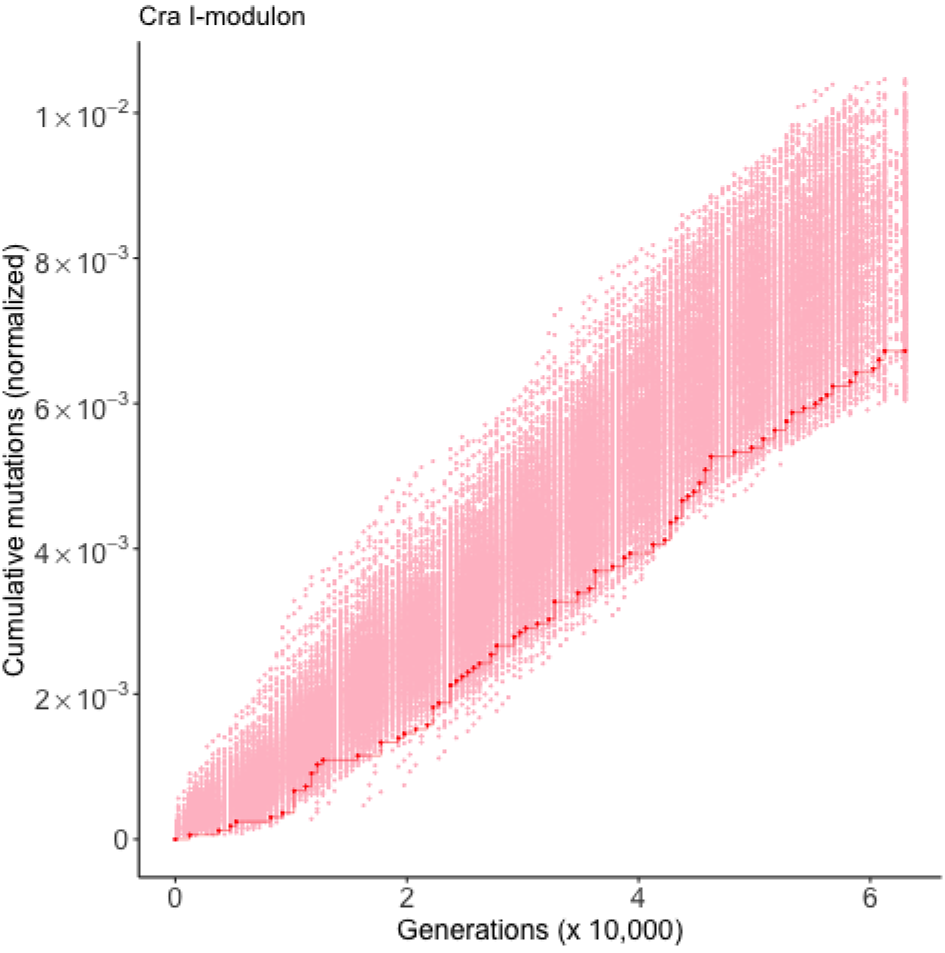
Cra I-modulon
Cumulative mutations (normalized)
$1\times10^{-2}$
$8\times10^{-3}$
$6\times10^{-3}$
$4\times10^{-3}$
$2\times10^{-3}$
0
0
2
4
6
Generations (x 10,000)

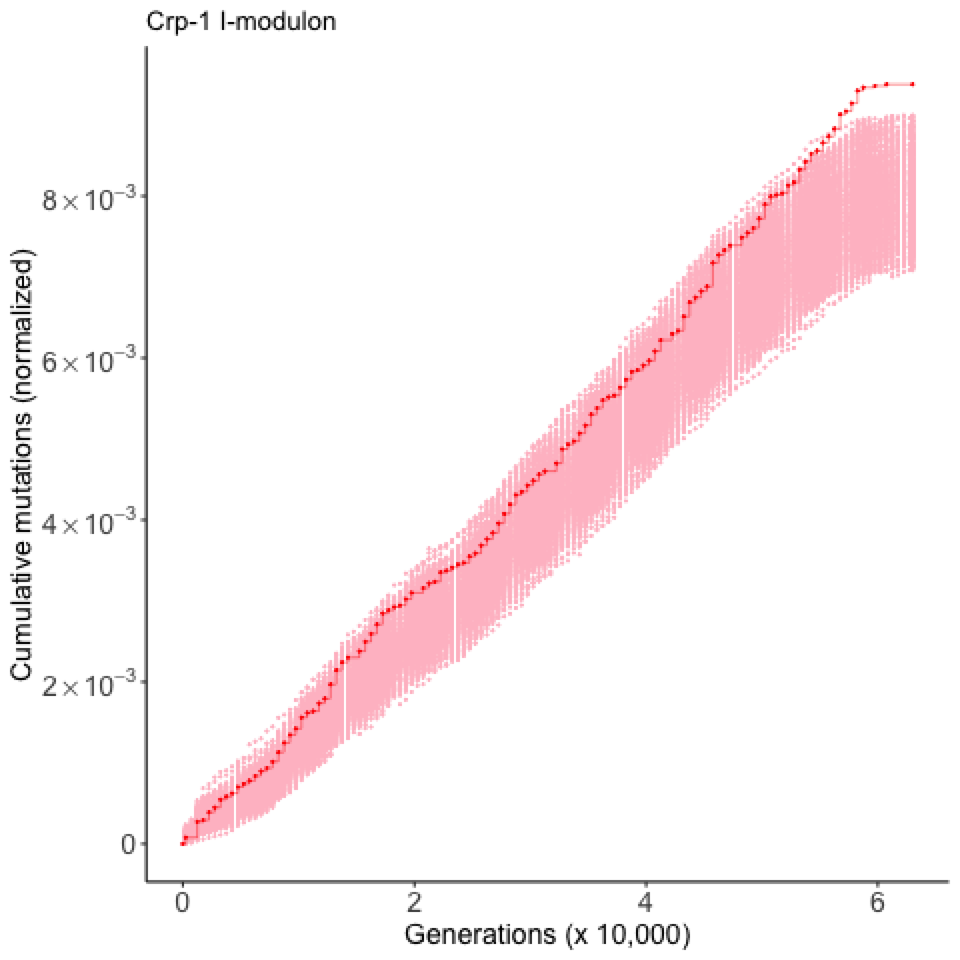
Crp-1 I-modulon
Cumulative mutations (normalized)
$8\times10^{-3}$
$6\times10^{-3}$
$4\times10^{-3}$
$2\times10^{-3}$
0
0
2
4
6
Generations (x 10,000)

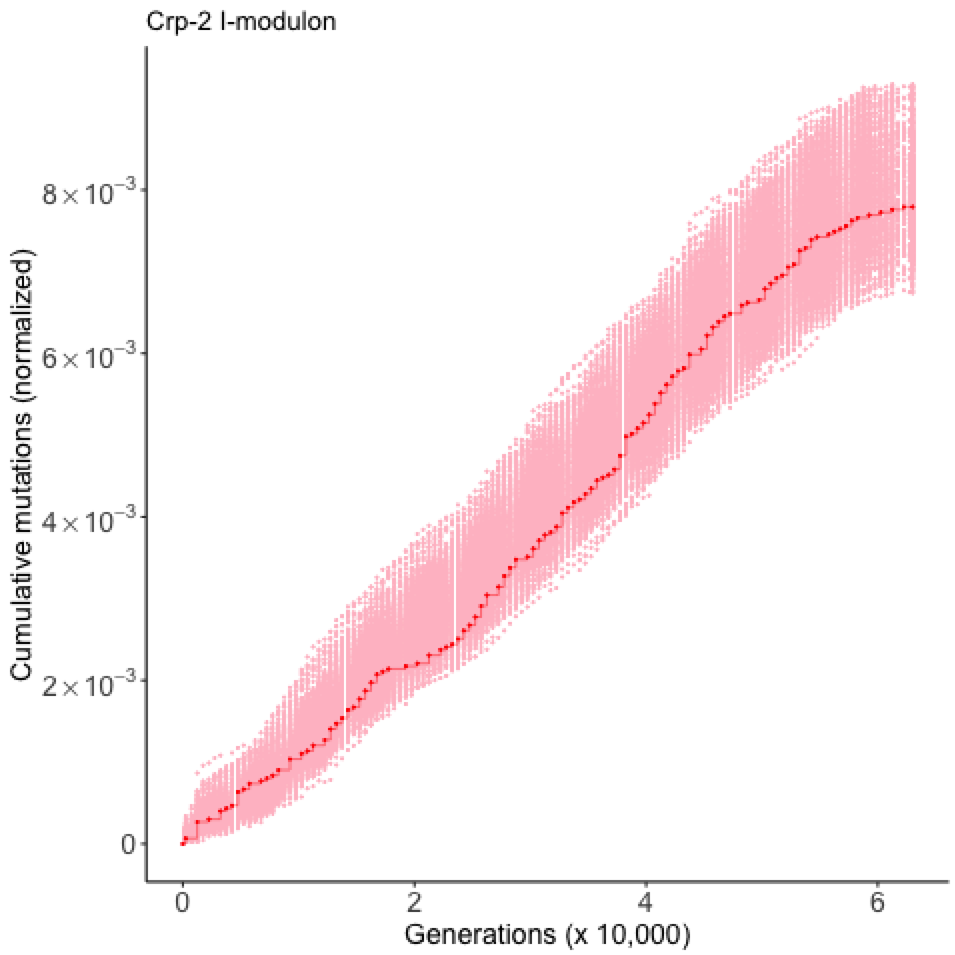

Crp-2 I-modulon
Cumulative mutations (normalized)
8×10⁻³
6×10⁻³
4×10⁻³
2×10⁻³
0
0
2
4
6
Generations (x 10,000)

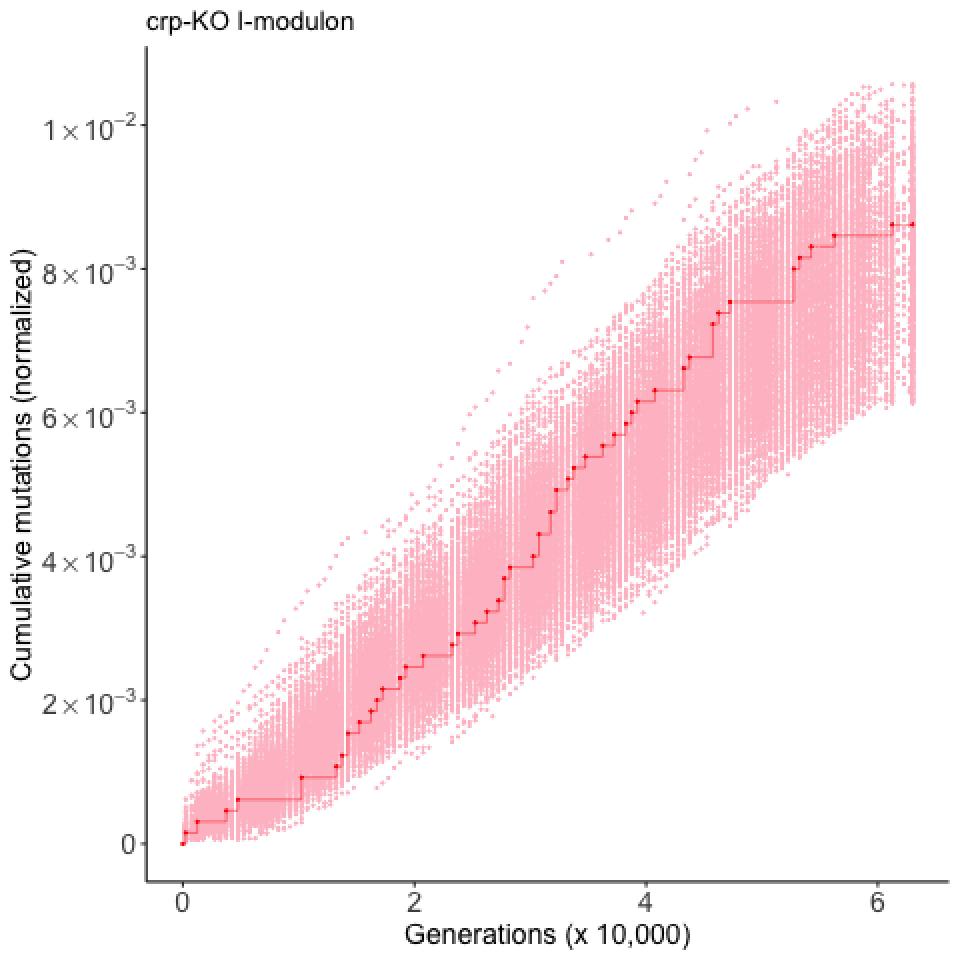

crp-KO I-modulon
Cumulative mutations (normalized)
1×10⁻²
8×10⁻³
6×10⁻³
4×10⁻³
2×10⁻³
0
0
2
4
6
Generations (x 10,000)

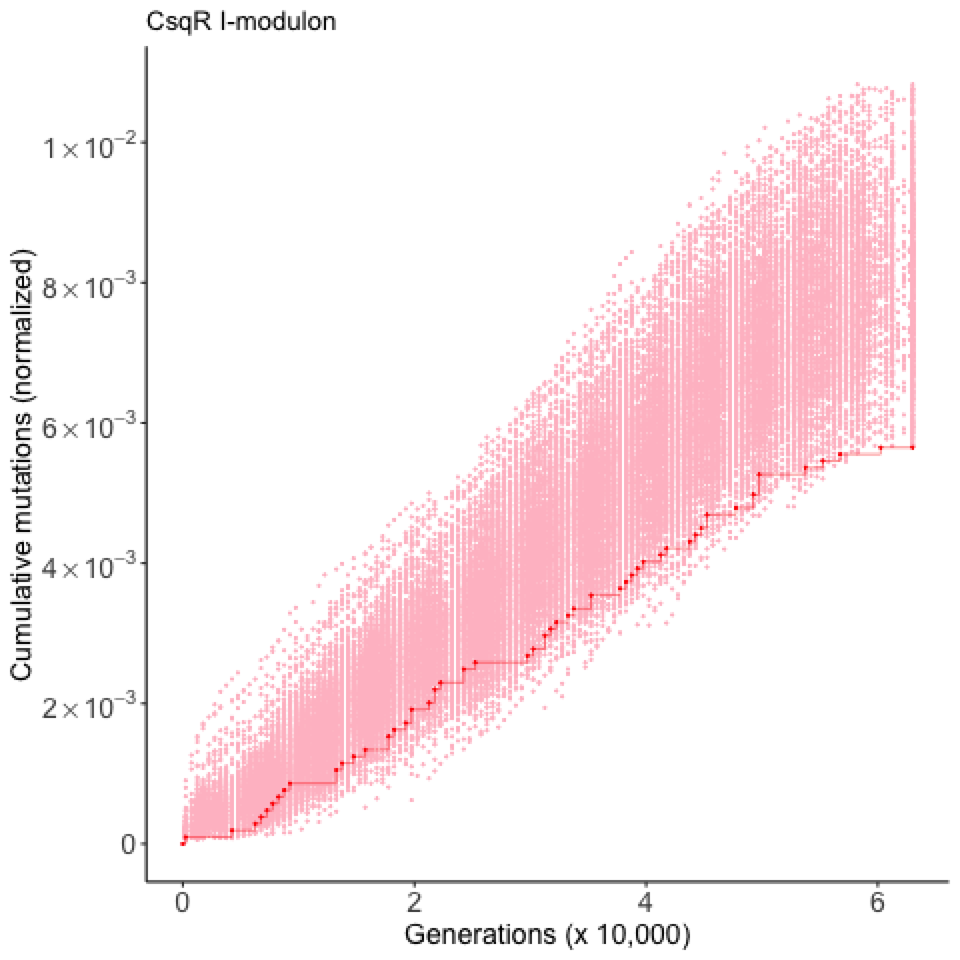
CsqR I-modulon
Cumulative mutations (normalized)
Generations (x 10,000)
0
2
4
6
$1\times10^{-2}$
$8\times10^{-3}$
$6\times10^{-3}$
$4\times10^{-3}$
$2\times10^{-3}$
0

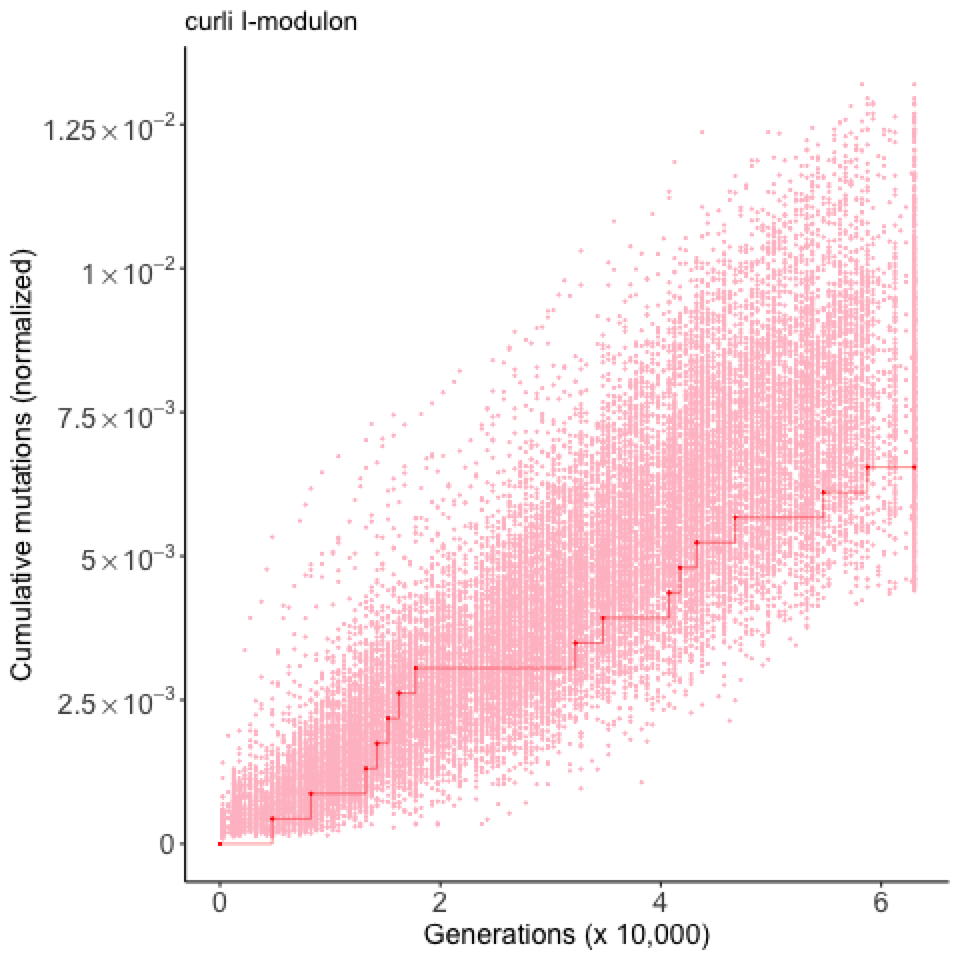

curli I-modulon
Cumulative mutations (normalized)
1.25×10⁻²
1×10⁻²
7.5×10⁻³
5×10⁻³
2.5×10⁻³
0
0
2
4
6
Generations (x 10,000)

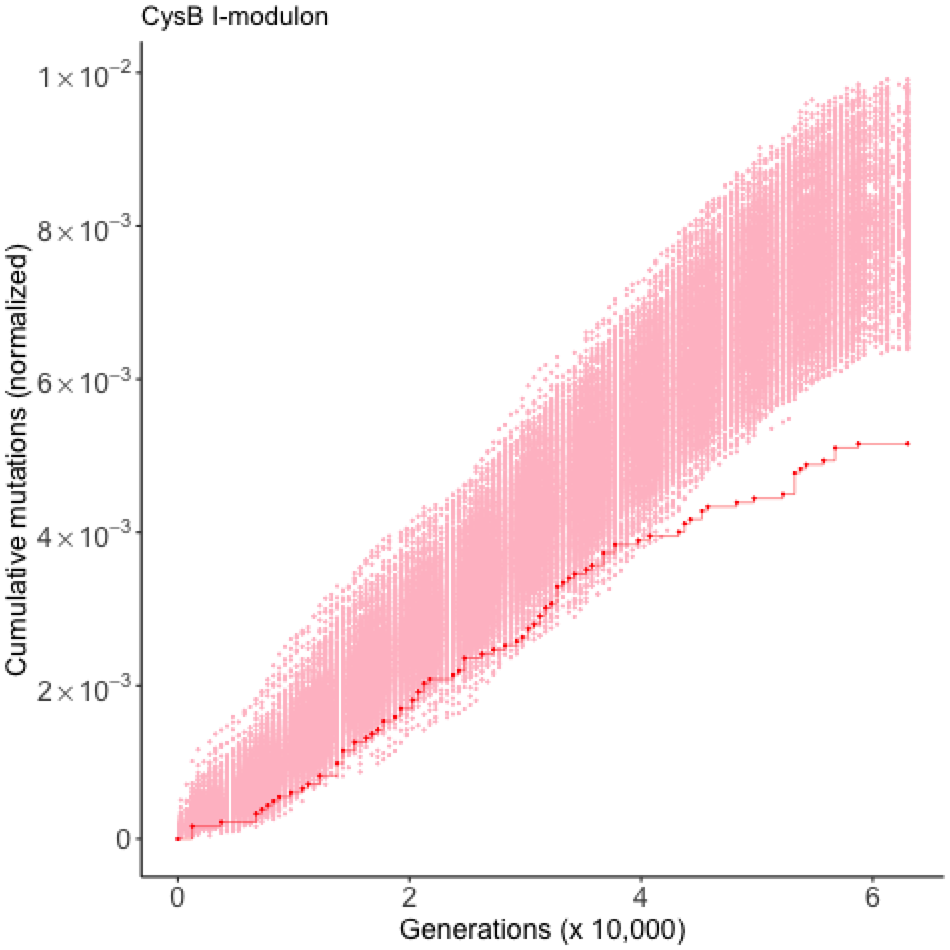

CysB I-modulon
Cumulative mutations (normalized)
$1\times10^{-2}$
$8\times10^{-3}$
$6\times10^{-3}$
$4\times10^{-3}$
$2\times10^{-3}$
0
0
2
4
6
Generations (x 10,000)

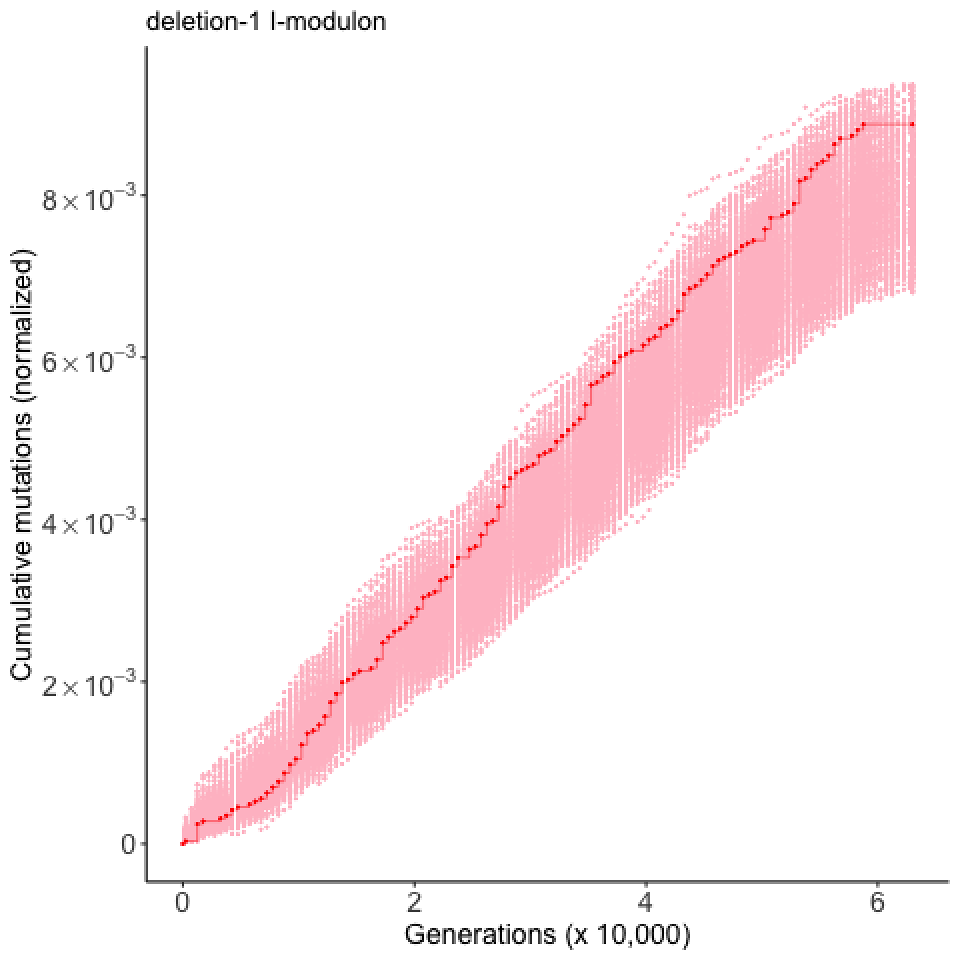

deletion-1 I-modulon
Cumulative mutations (normalized)
8×10^−3
6×10^−3
4×10^−3
2×10^−3
0
0
2
4
6
Generations (x 10,000)

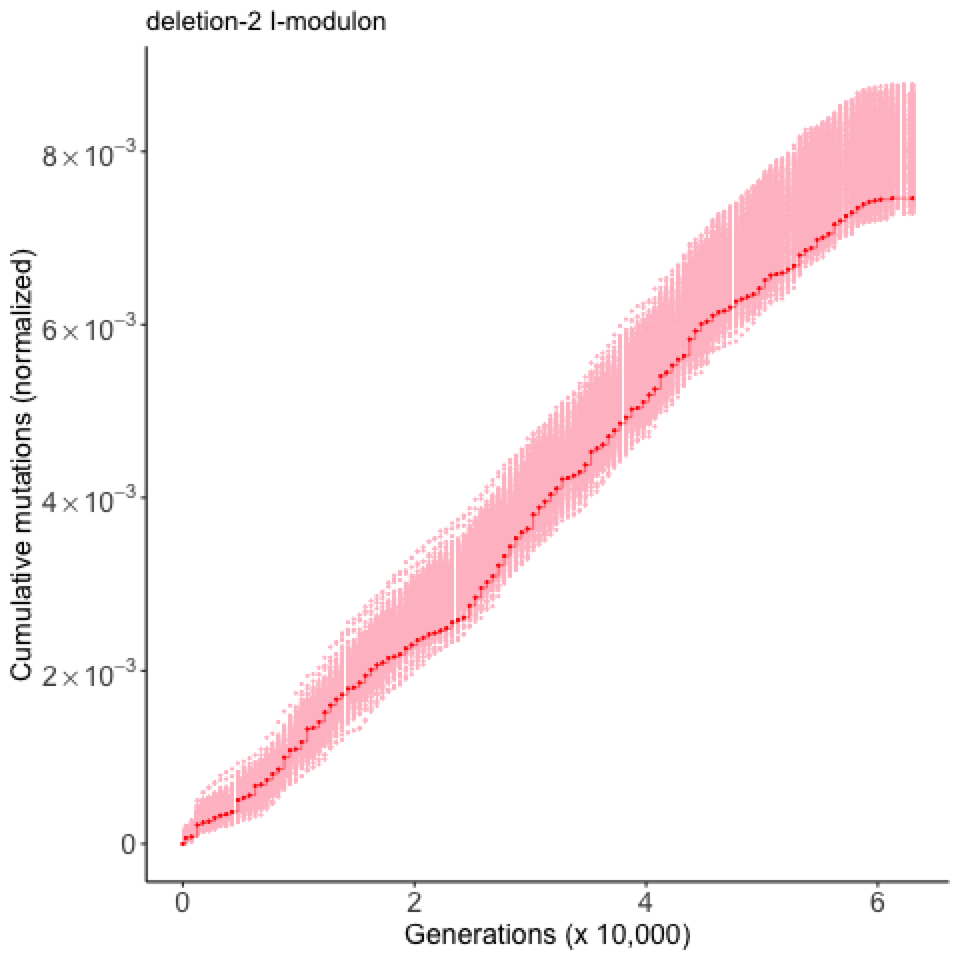

deletion-2 I-modulon
Cumulative mutations (normalized)
$8\times10^{-3}$
$6\times10^{-3}$
$4\times10^{-3}$
$2\times10^{-3}$
0
0
2
4
6
Generations (x 10,000)

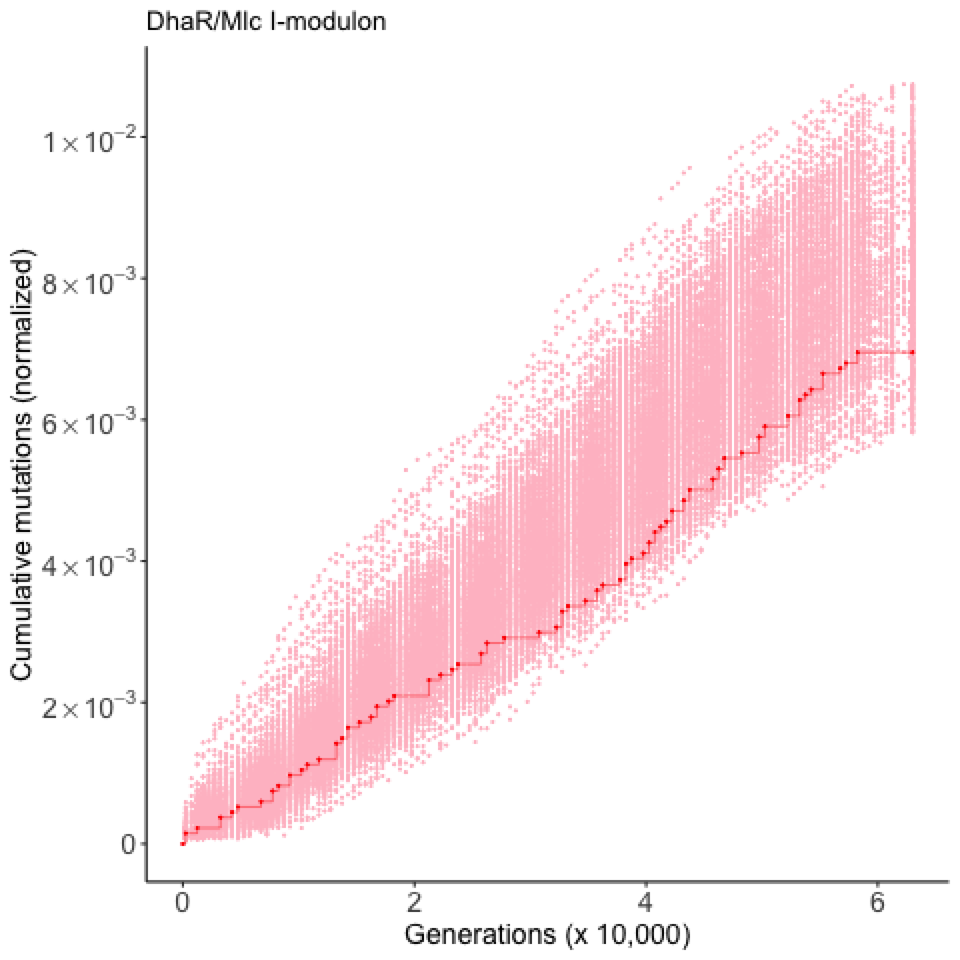

DhaR/Mlc I-modulon
Cumulative mutations (normalized)
$1\times10^{-2}$
$8\times10^{-3}$
$6\times10^{-3}$
$4\times10^{-3}$
$2\times10^{-3}$
0
0
2
4
6
Generations (x 10,000)

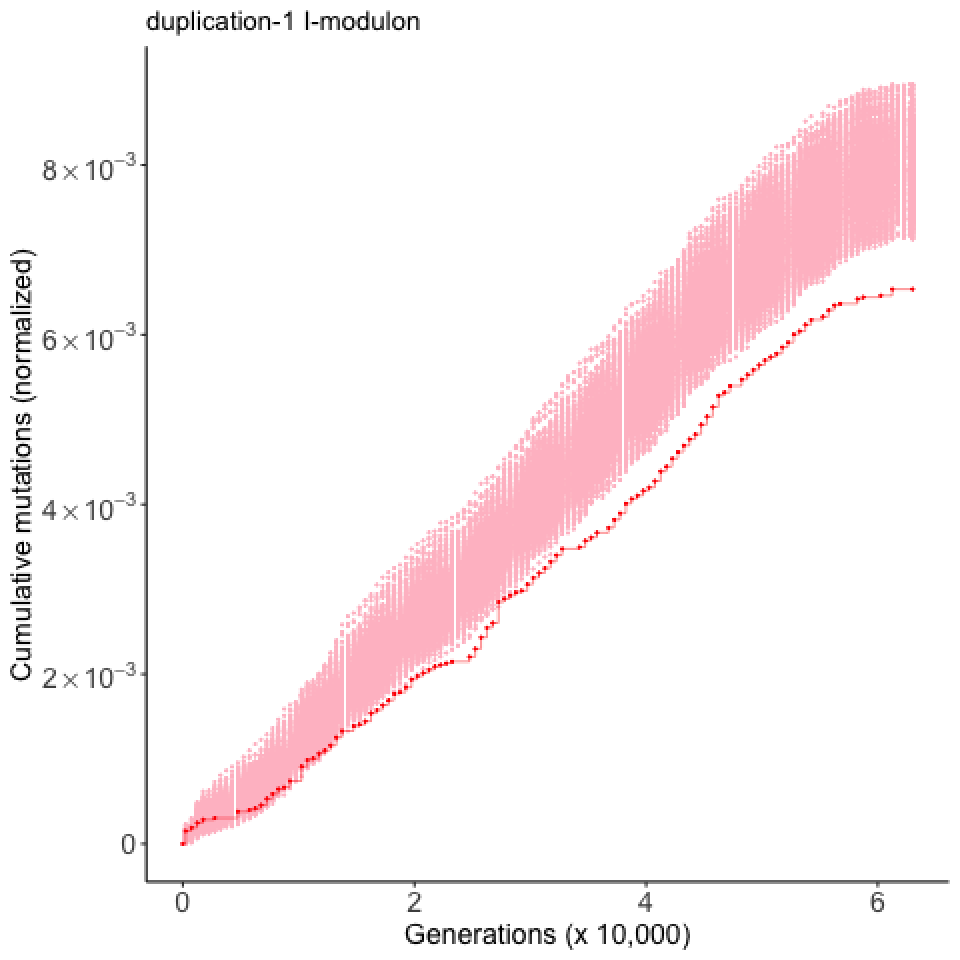
duplication-1 I-modulon
Cumulative mutations (normalized)
$8\times10^{-3}$
$6\times10^{-3}$
$4\times10^{-3}$
$2\times10^{-3}$
0
0
2
4
6
Generations (x 10,000)

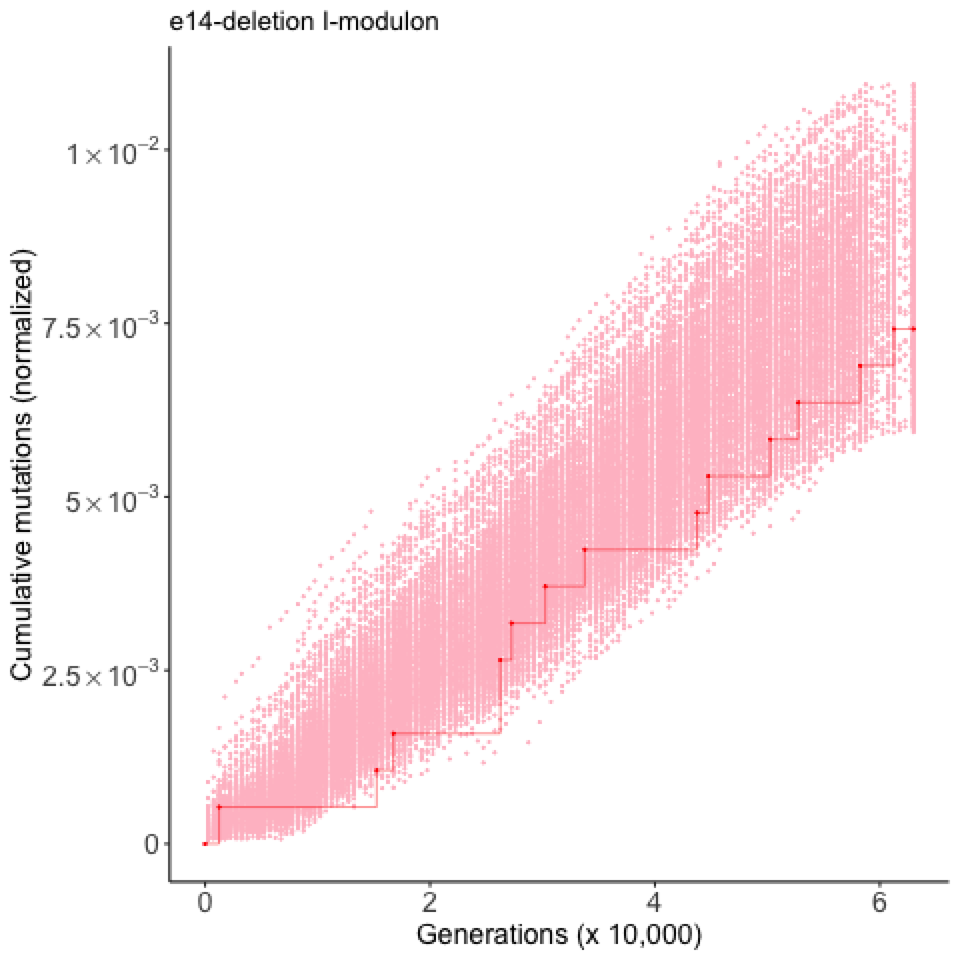

e14-deletion I-modulon
Cumulative mutations (normalized)
1×10^-2
7.5×10^-3
5×10^-3
2.5×10^-3
0
0
2
4
6
Generations (x 10,000)

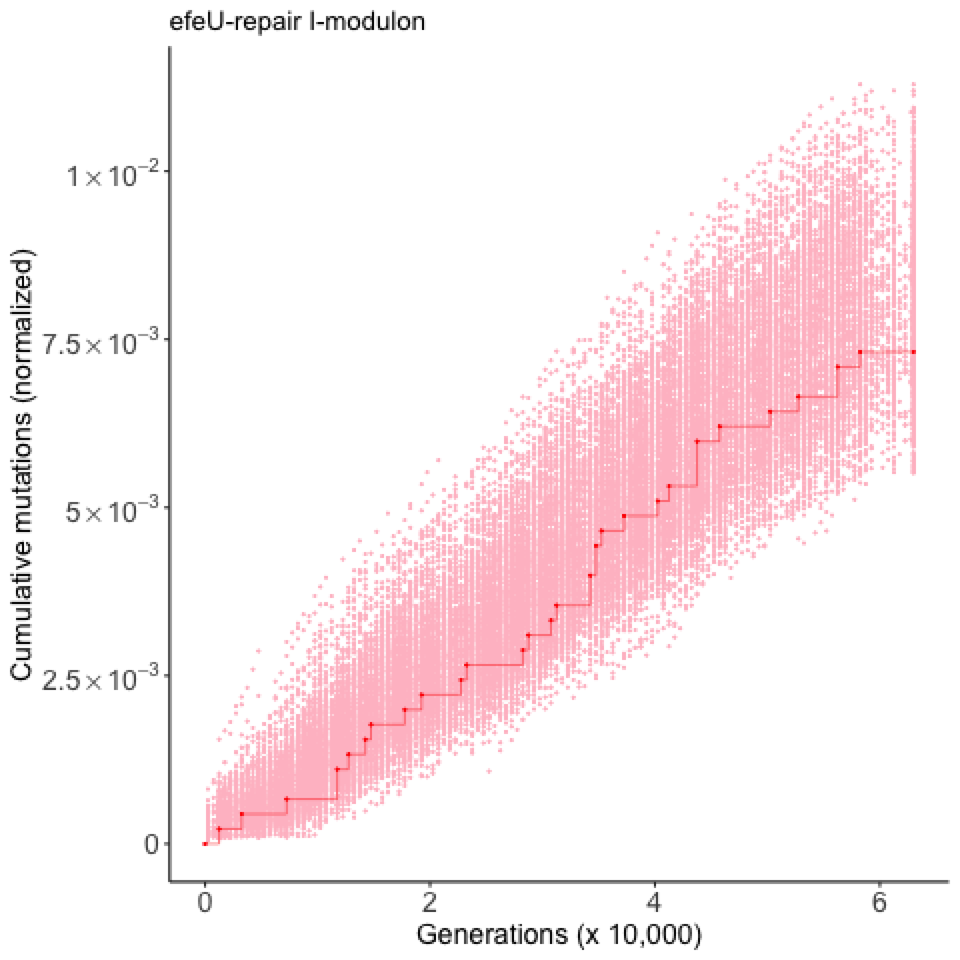

efeU-repair I-modulon
Cumulative mutations (normalized)
$1\times10^{-2}$
$7.5\times10^{-3}$
$5\times10^{-3}$
$2.5\times10^{-3}$
0
0
2
4
6
Generations (x 10,000)

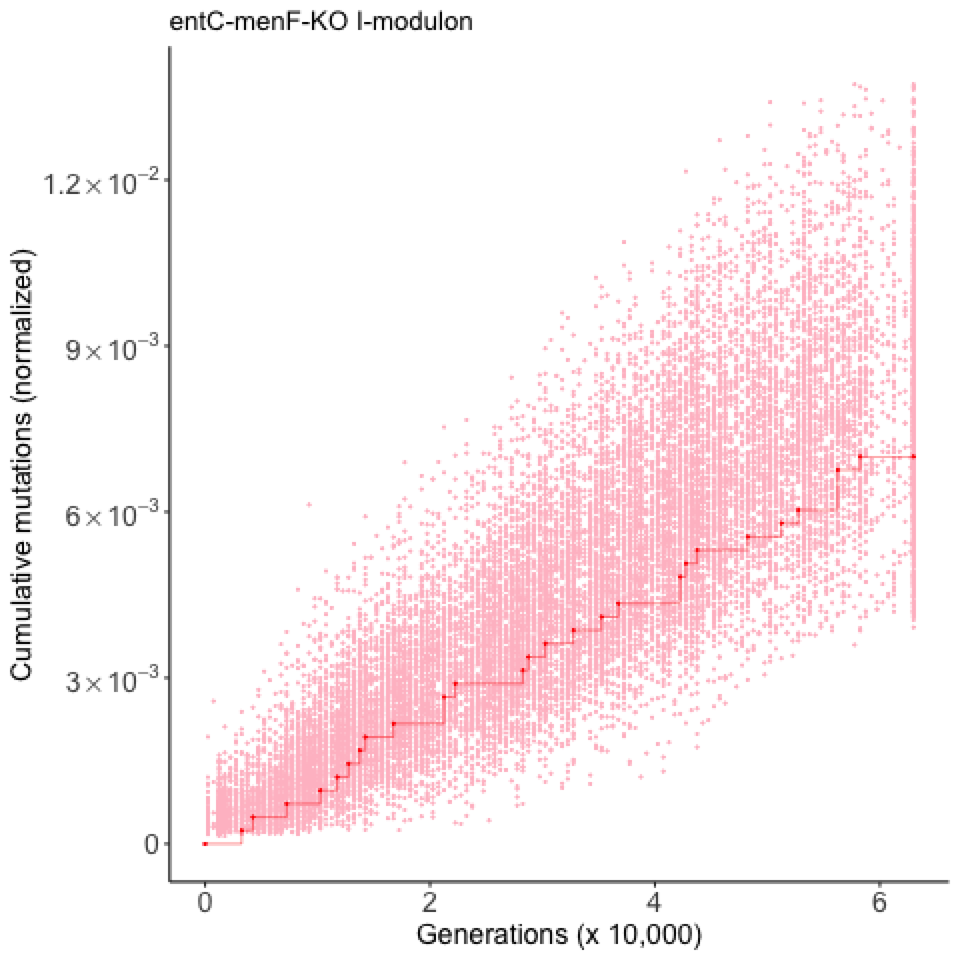
entC-menF-KO I-modulon
Cumulative mutations (normalized)
1.2×10^-2
9×10^-3
6×10^-3
3×10^-3
0
0
2
4
6
Generations (x 10,000)

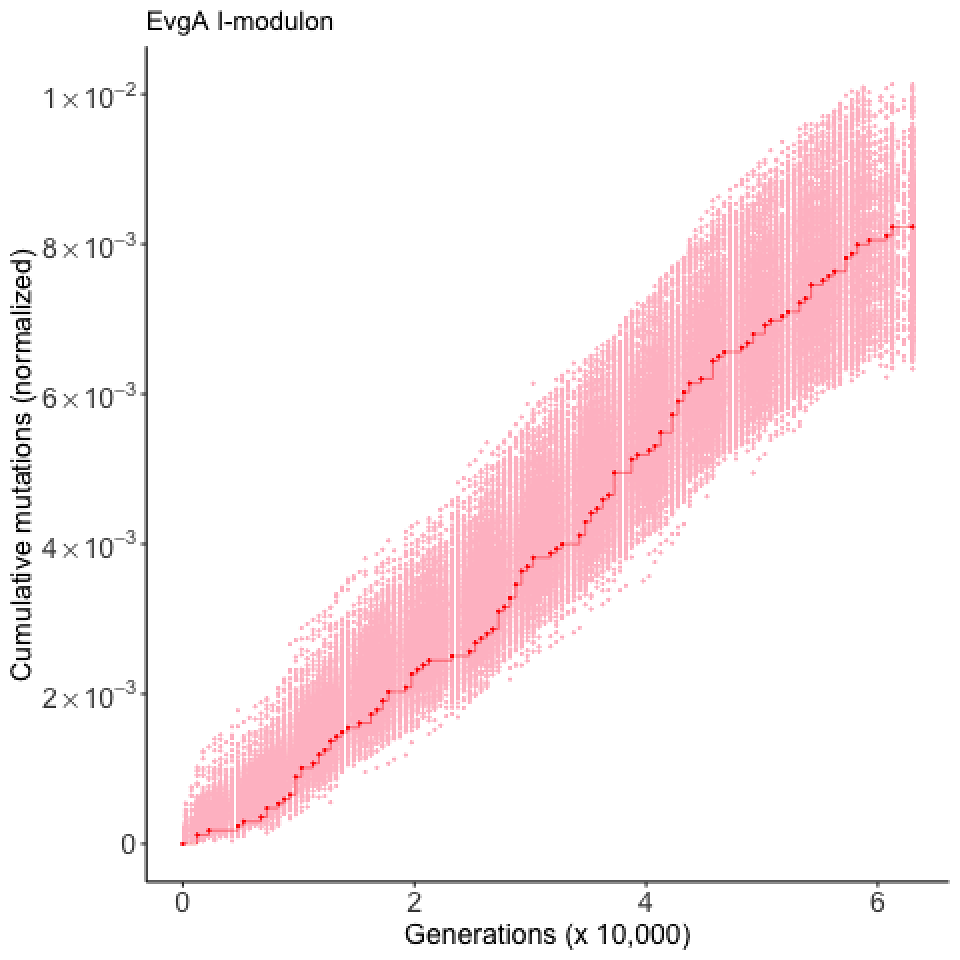

EvgA I-modulon
Cumulative mutations (normalized)
$1\times10^{-2}$
$8\times10^{-3}$
$6\times10^{-3}$
$4\times10^{-3}$
$2\times10^{-3}$
0
0
2
4
6
Generations (x 10,000)

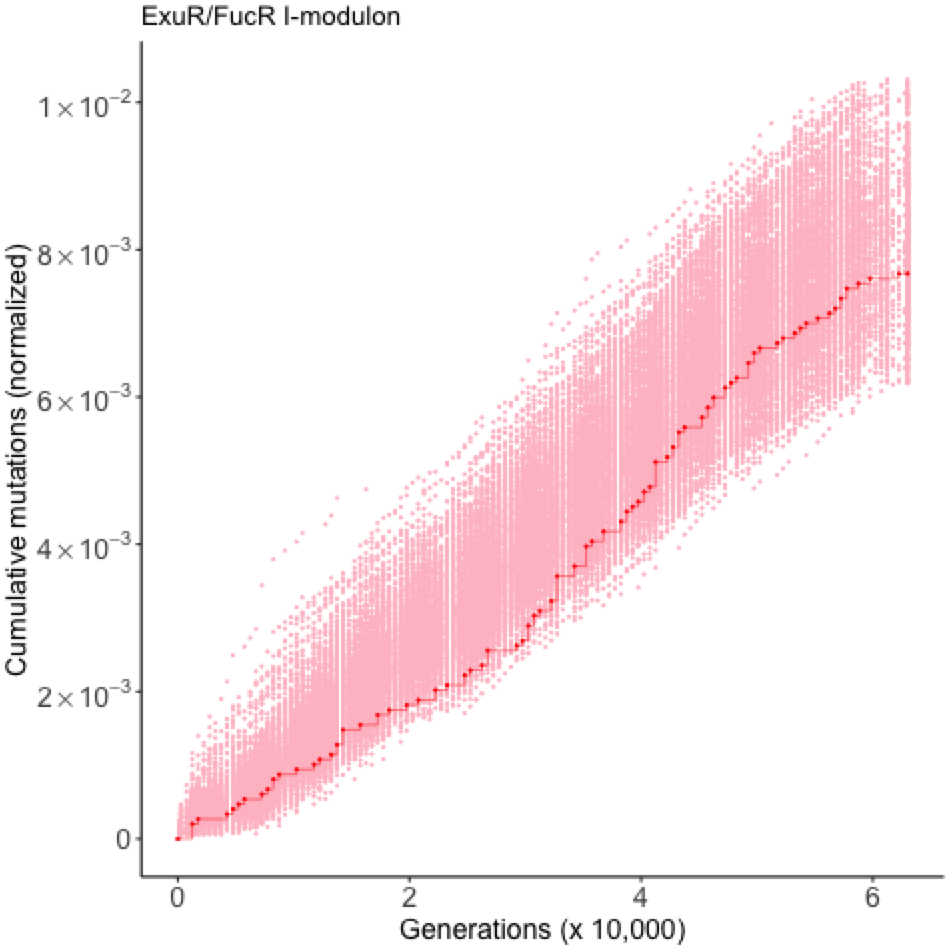

ExuR/FucR I-modulon
Cumulative mutations (normalized)
$1\times10^{-2}$
$8\times10^{-3}$
$6\times10^{-3}$
$4\times10^{-3}$
$2\times10^{-3}$
0
0
2
4
6
Generations (x 10,000)

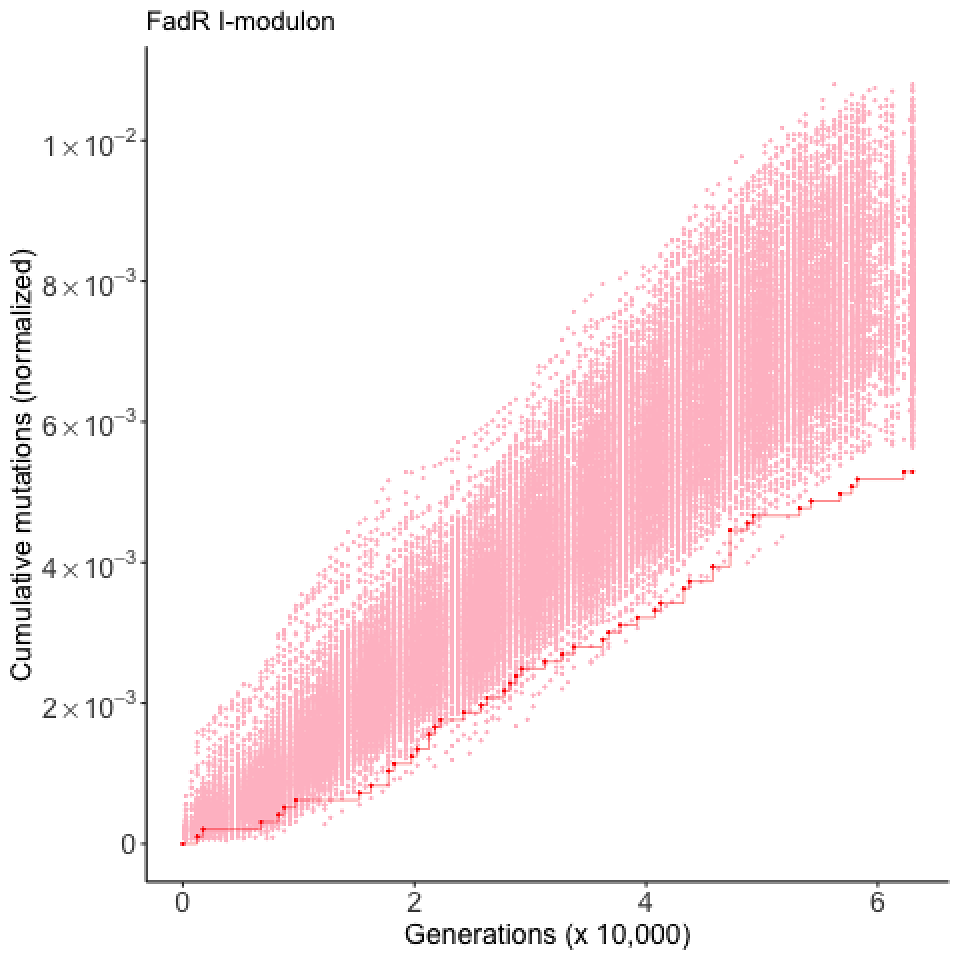

FadR I-modulon
Cumulative mutations (normalized)
1×10⁻²
8×10⁻³
6×10⁻³
4×10⁻³
2×10⁻³
0
0
2
4
6
Generations (x 10,000)

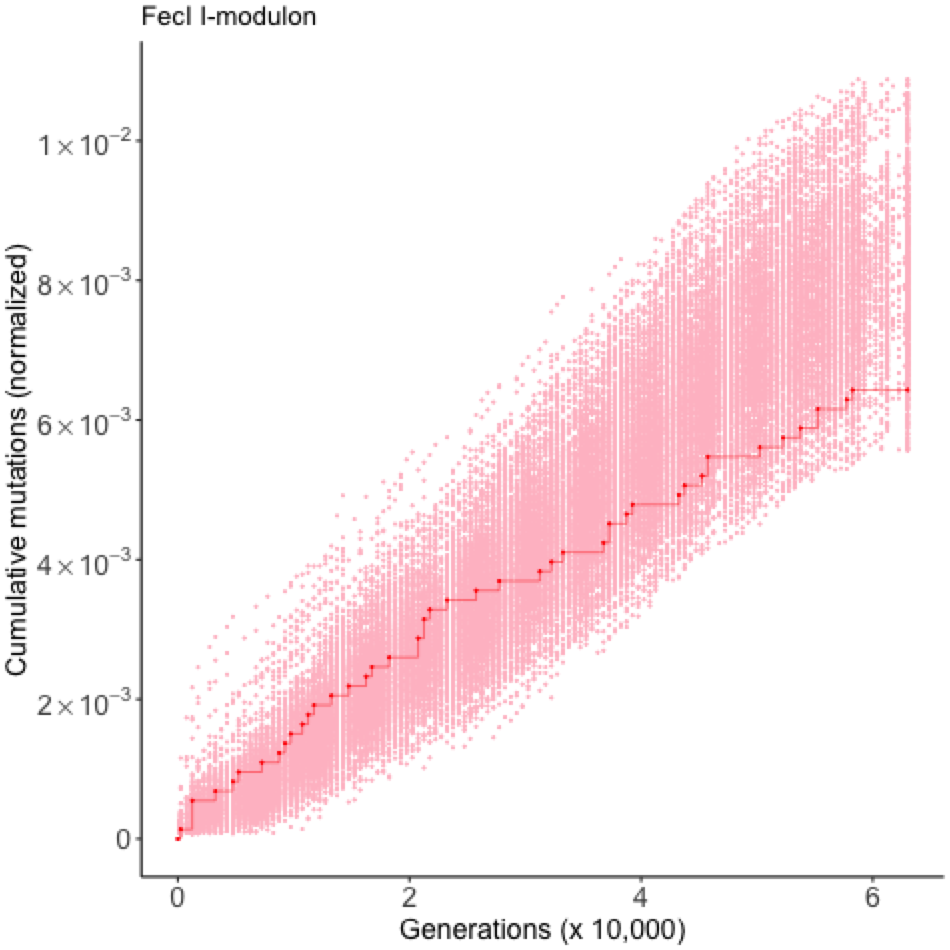

FecI I-modulon
Cumulative mutations (normalized)
$1\times10^{-2}$
$8\times10^{-3}$
$6\times10^{-3}$
$4\times10^{-3}$
$2\times10^{-3}$
0
0
2
4
6
Generations (x 10,000)

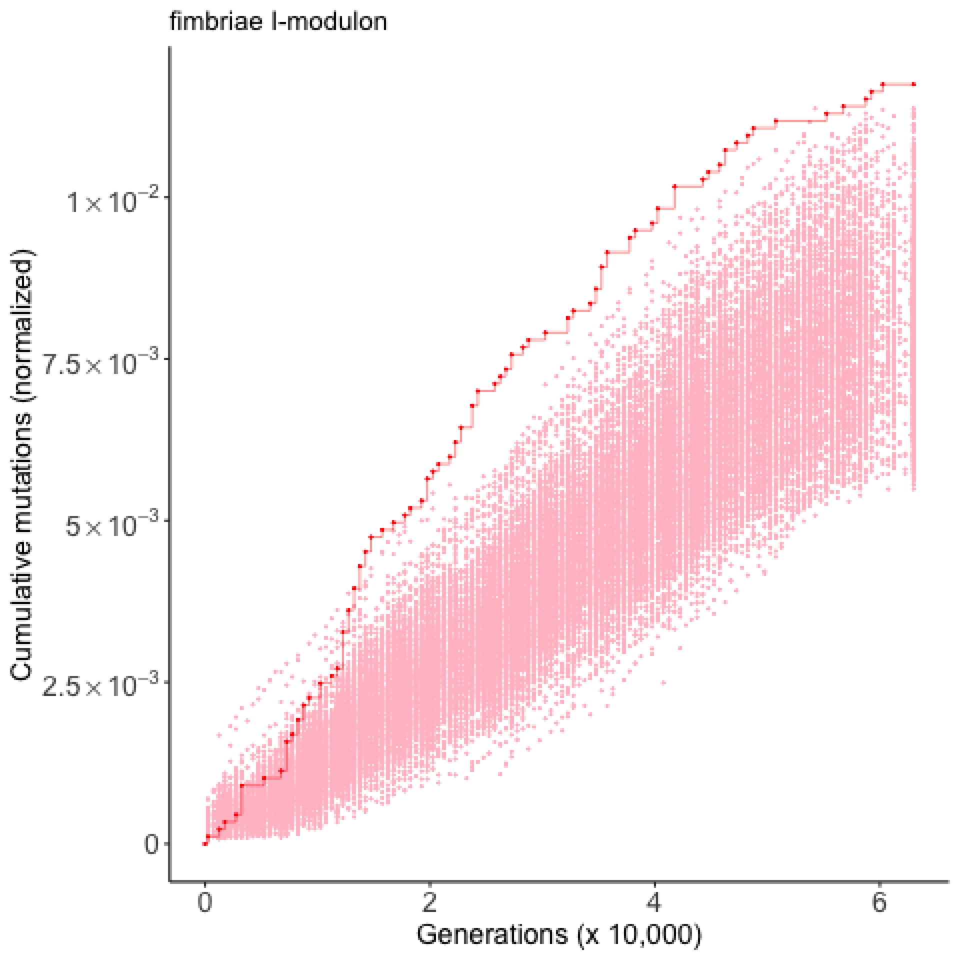

fimbriae I-modulon
Cumulative mutations (normalized)
1×10^−2
7.5×10^−3
5×10^−3
2.5×10^−3
0
0
2
4
6
Generations (x 10,000)

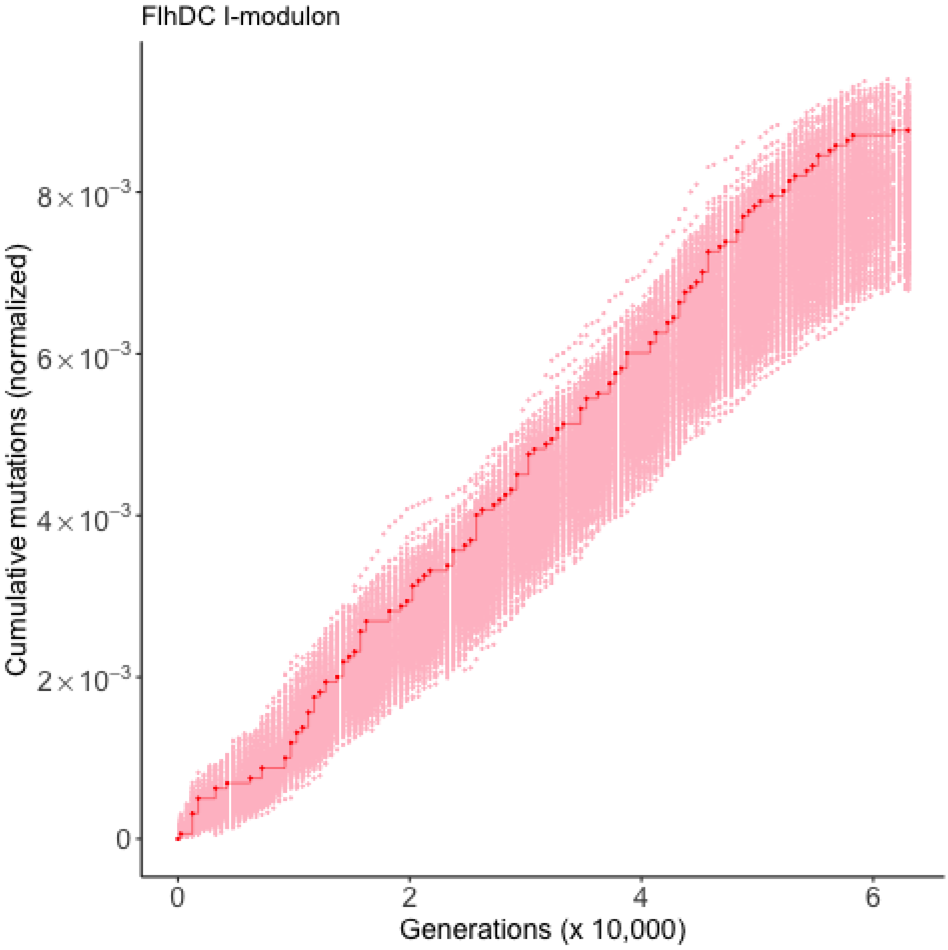

FlhDC I-modulon
Cumulative mutations (normalized)
8×10⁻³
6×10⁻³
4×10⁻³
2×10⁻³
0
0
2
4
6
Generations (x 10,000)

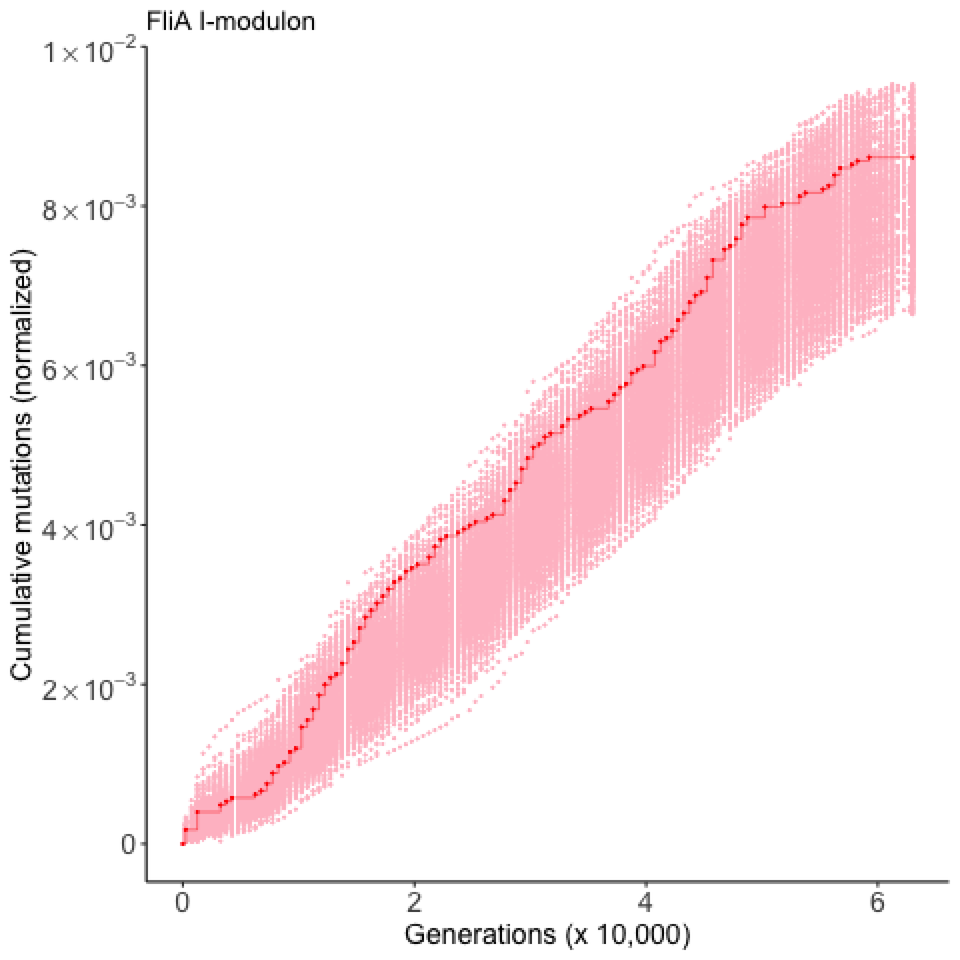

FliA I-modulon
Cumulative mutations (normalized)
$1\times10^{-2}$
$8\times10^{-3}$
$6\times10^{-3}$
$4\times10^{-3}$
$2\times10^{-3}$
0
0
2
4
6
Generations (x 10,000)

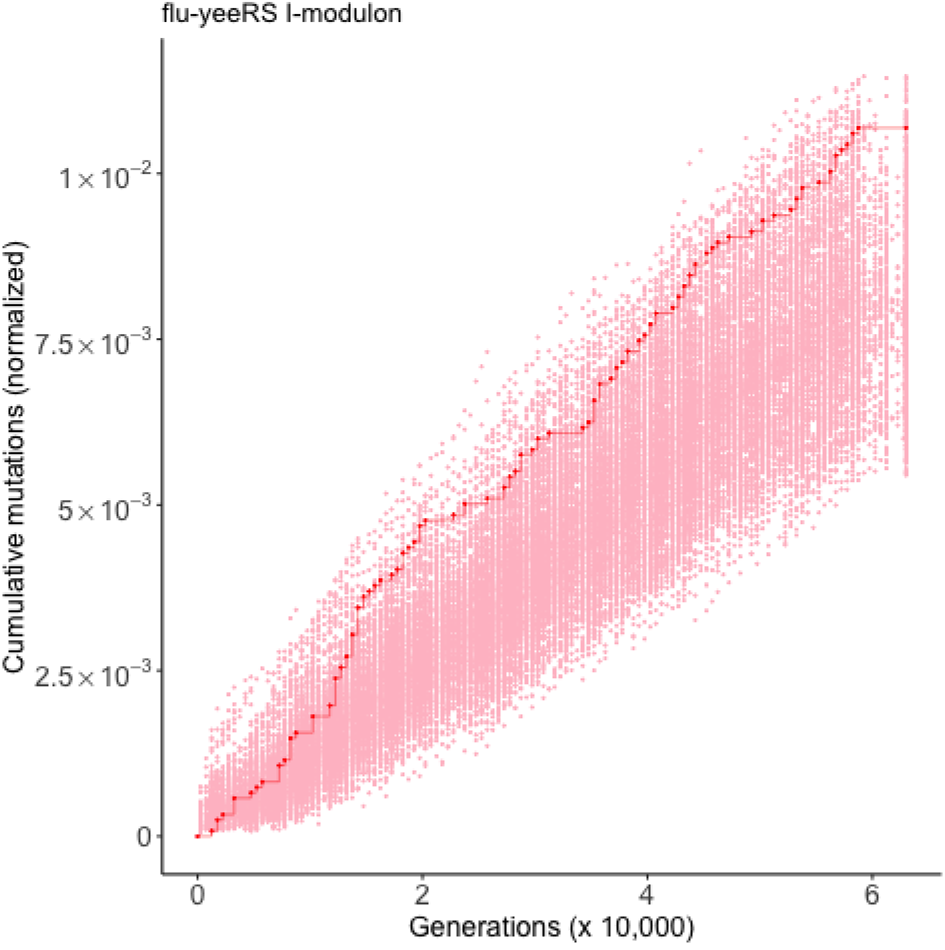
flu-yeeRS I-modulon
Cumulative mutations (normalized)
$1\times10^{-2}$
$7.5\times10^{-3}$
$5\times10^{-3}$
$2.5\times10^{-3}$
0
0
2
4
6
Generations (x 10,000)

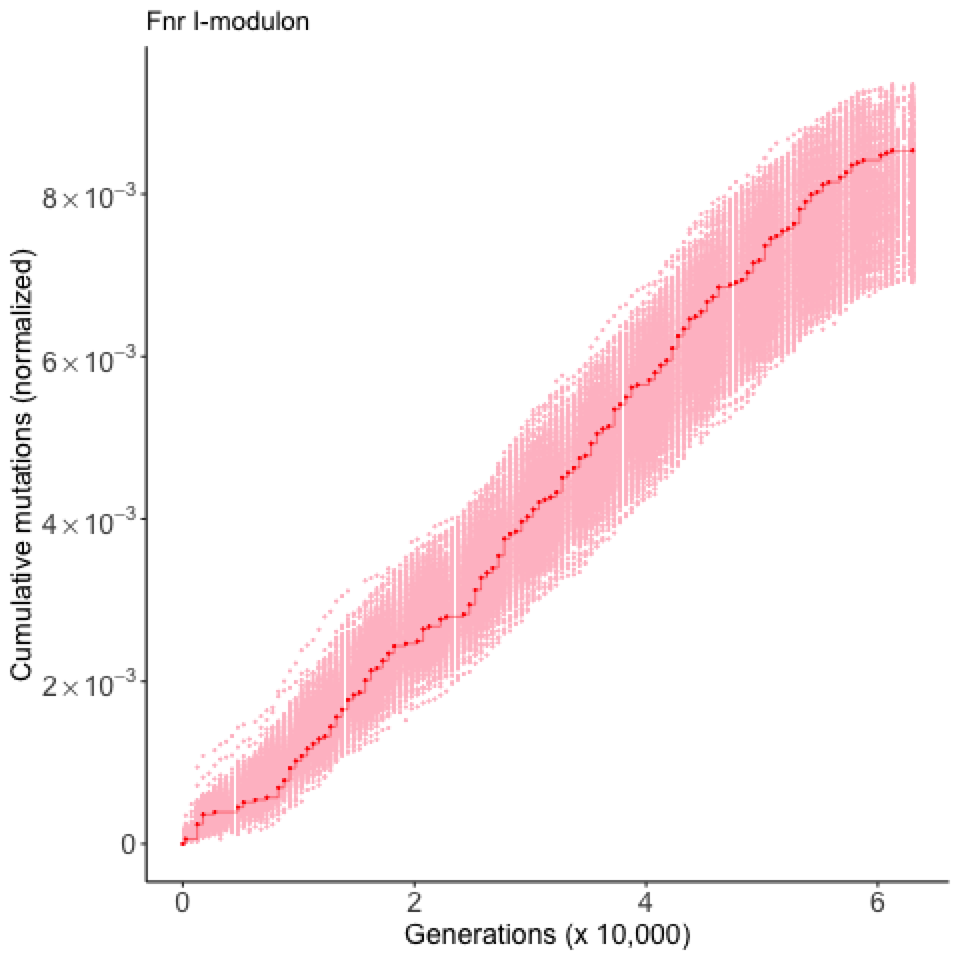

Fnr I-modulon
Cumulative mutations (normalized)
8×10⁻³
6×10⁻³
4×10⁻³
2×10⁻³
0
0
2
4
6
Generations (x 10,000)

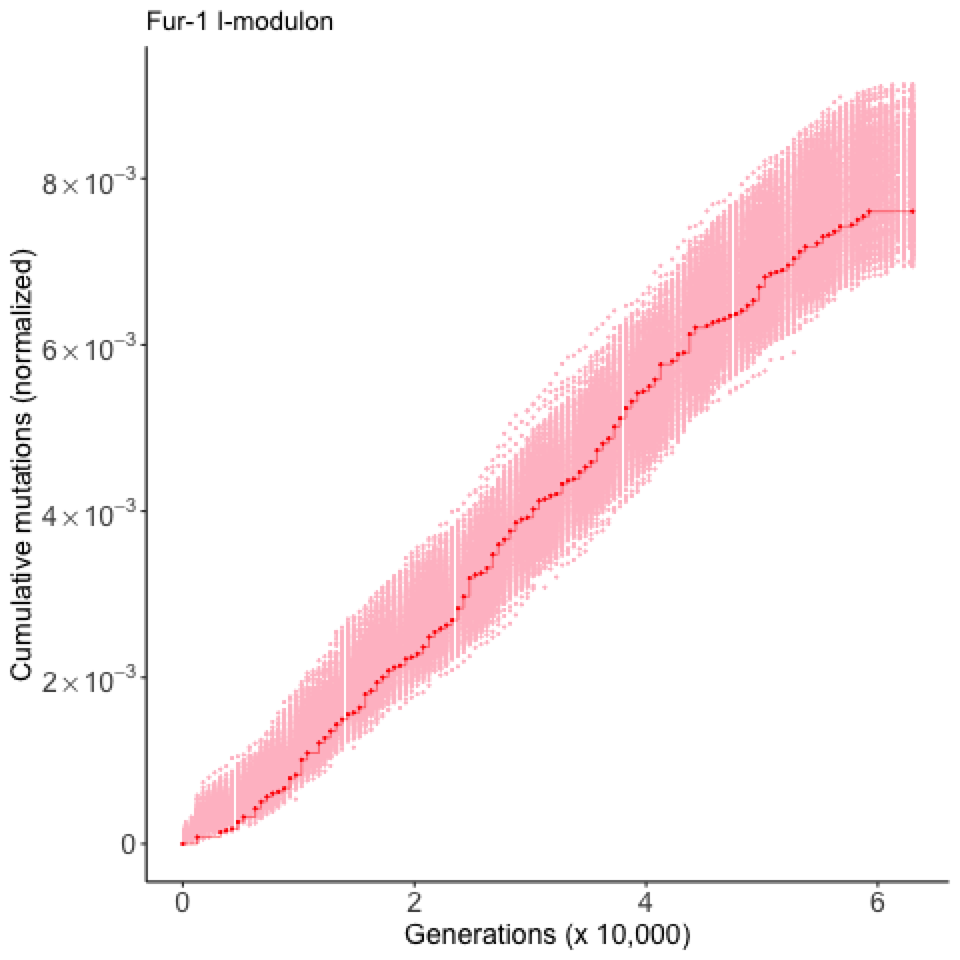

Fur-1 I-modulon
Cumulative mutations (normalized)
$8\times10^{-3}$
$6\times10^{-3}$
$4\times10^{-3}$
$2\times10^{-3}$
0
0
2
4
6
Generations (x 10,000)

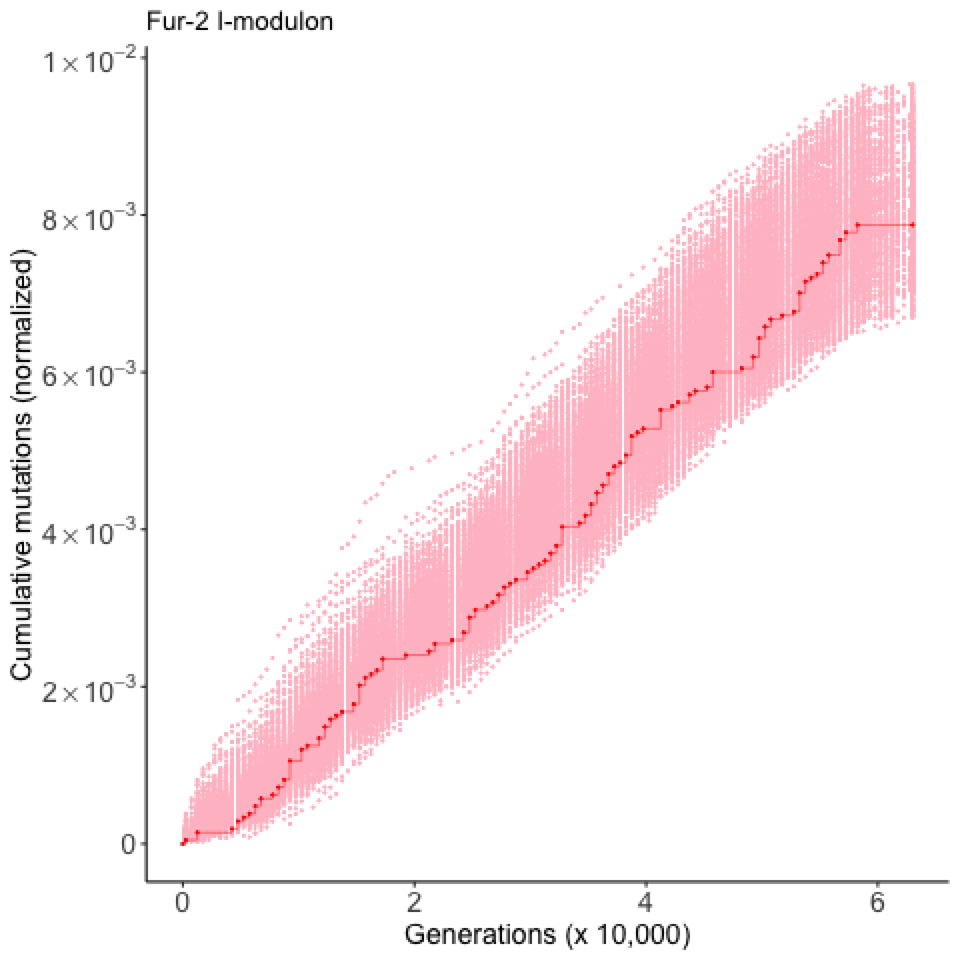

Fur-2 I-modulon
$1\times10^{-2}$
$8\times10^{-3}$
$6\times10^{-3}$
$4\times10^{-3}$
$2\times10^{-3}$
0
Cumulative mutations (normalized)
0
2
4
6
Generations (x 10,000)

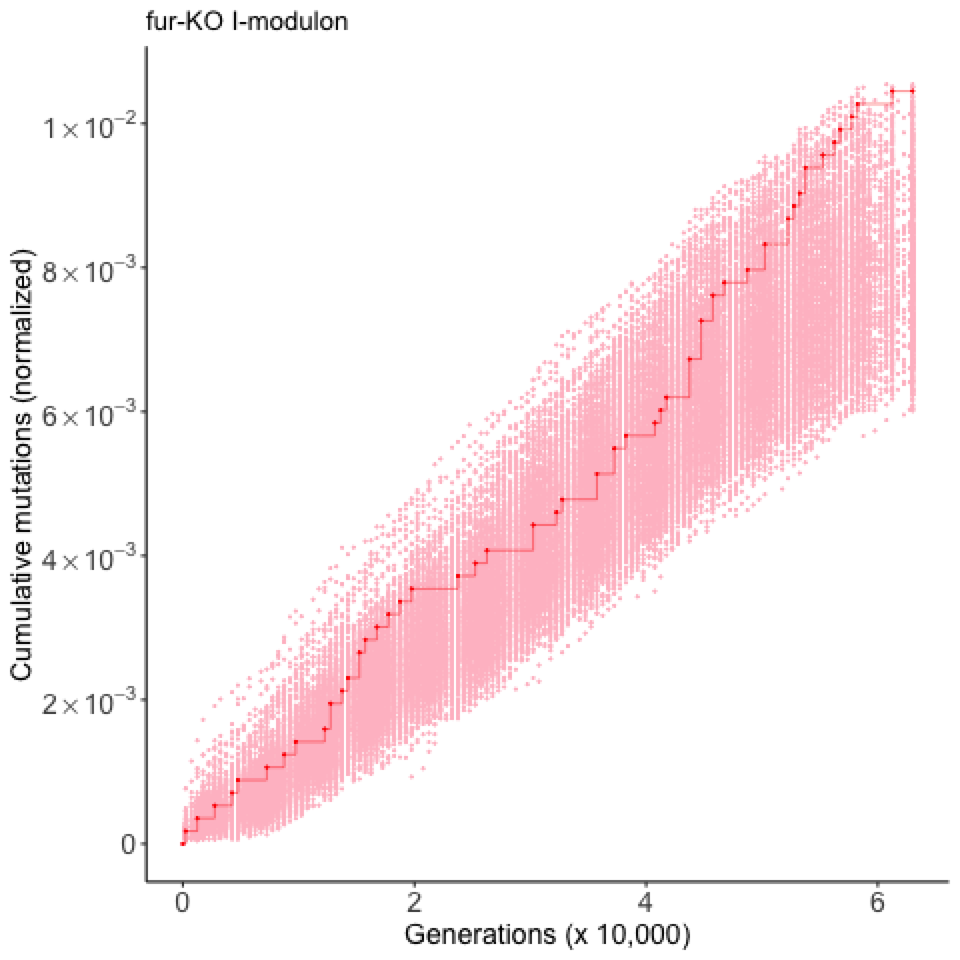
fur-KO I-modulon
Cumulative mutations (normalized)
$1\times10^{-2}$
$8\times10^{-3}$
$6\times10^{-3}$
$4\times10^{-3}$
$2\times10^{-3}$
0
0
2
4
6
Generations (x 10,000)

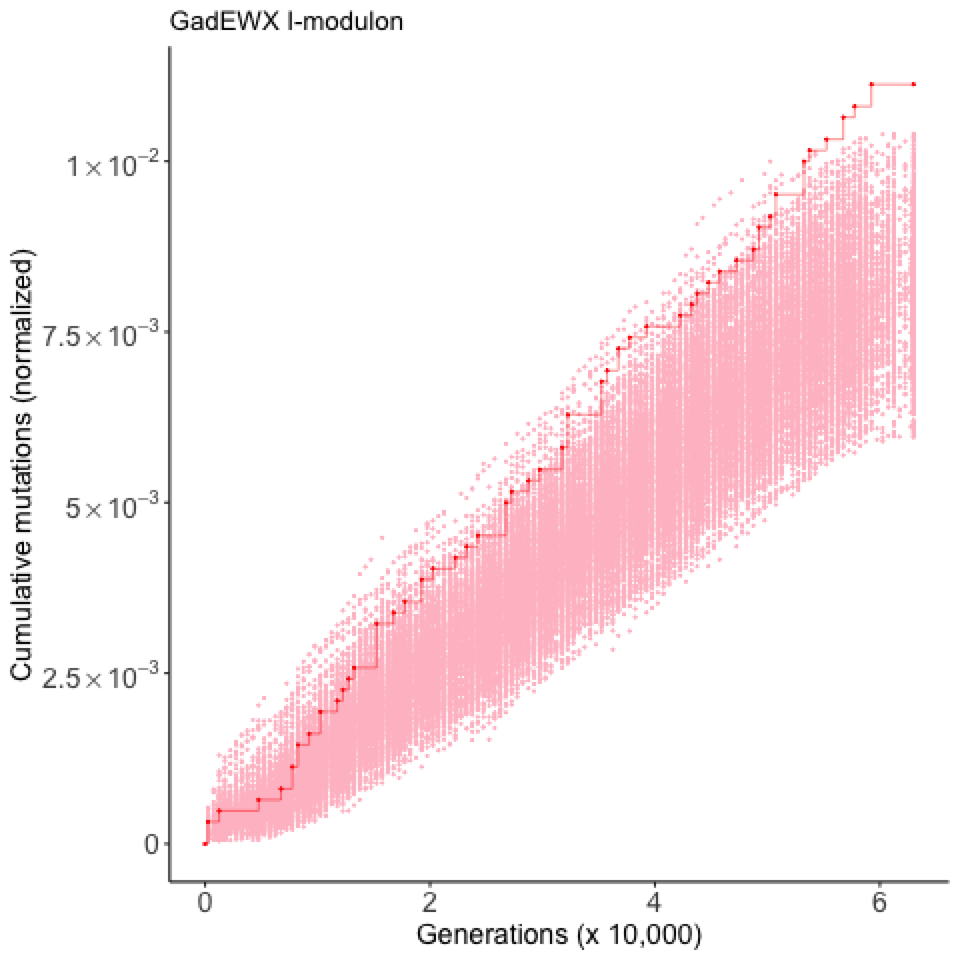
GadEWX I-modulon
Cumulative mutations (normalized)
$1\times10^{-2}$
$7.5\times10^{-3}$
$5\times10^{-3}$
$2.5\times10^{-3}$
0
0
2
4
6
Generations (x 10,000)

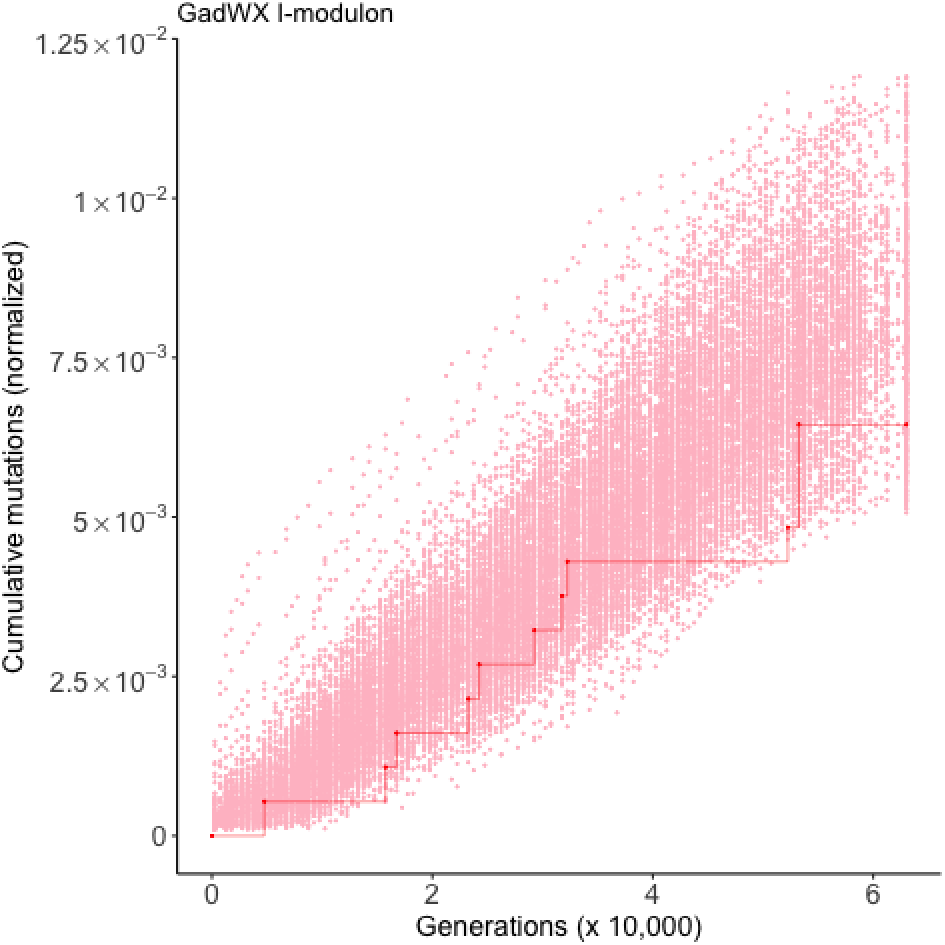
GadWX I-modulon
1.25×10^−2
1×10^−2
7.5×10^−3
5×10^−3
2.5×10^−3
0
Cumulative mutations (normalized)
0
2
4
6
Generations (x 10,000)

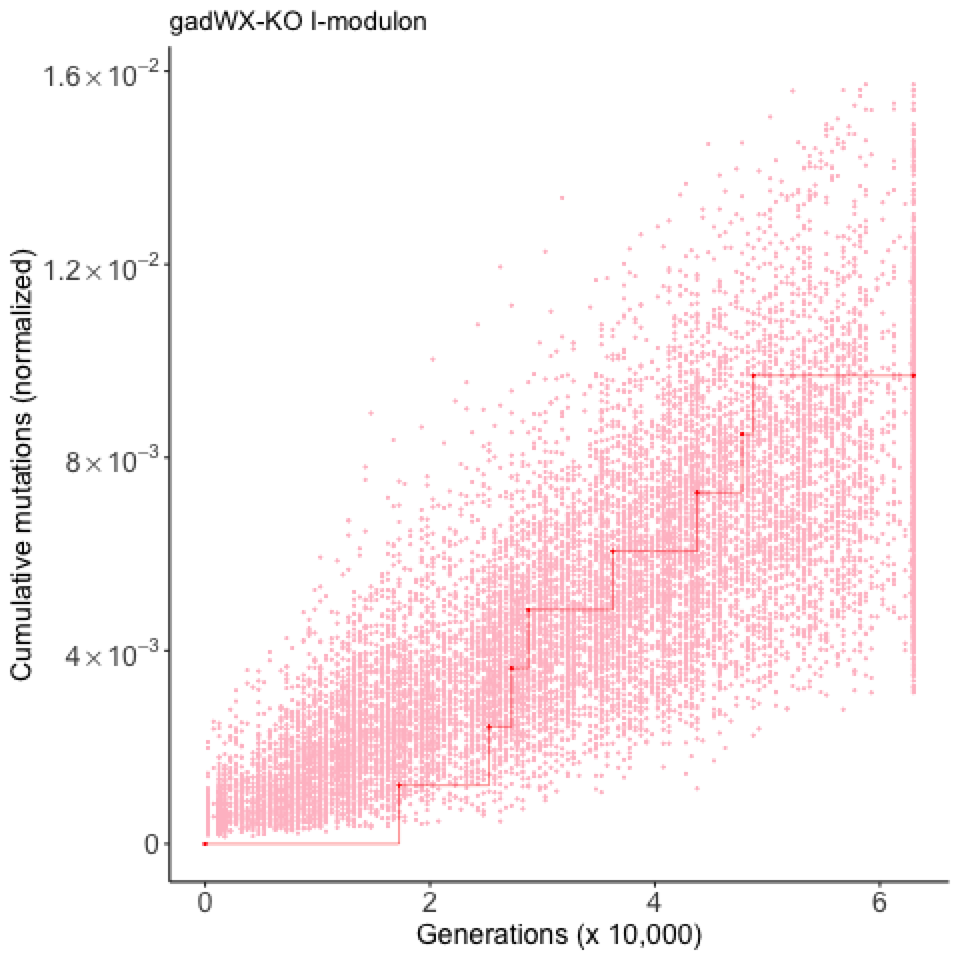

gadWX-KO I-modulon
Cumulative mutations (normalized)
1.6×10⁻²
1.2×10⁻²
8×10⁻³
4×10⁻³
0
0
2
4
6
Generations (x 10,000)

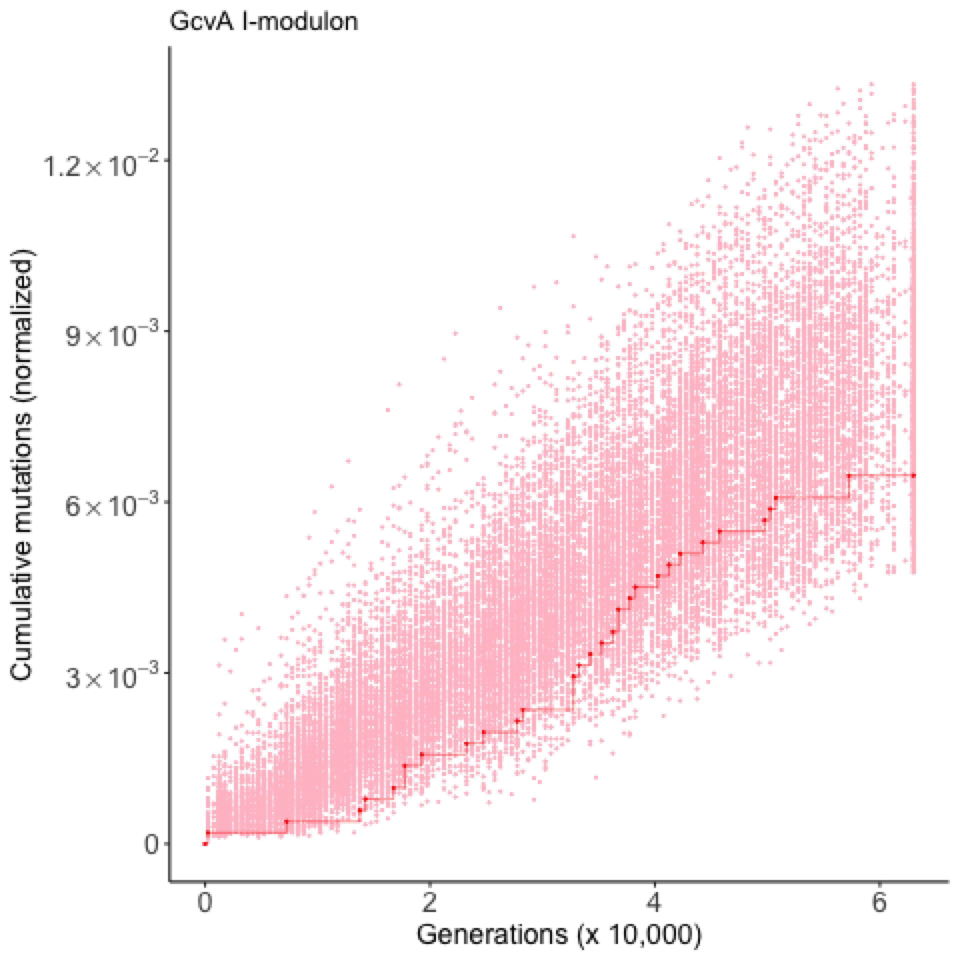
GcvA I-modulon
Cumulative mutations (normalized)
$1.2\times10^{-2}$
$9\times10^{-3}$
$6\times10^{-3}$
$3\times10^{-3}$
0
0
2
4
6
Generations (x 10,000)

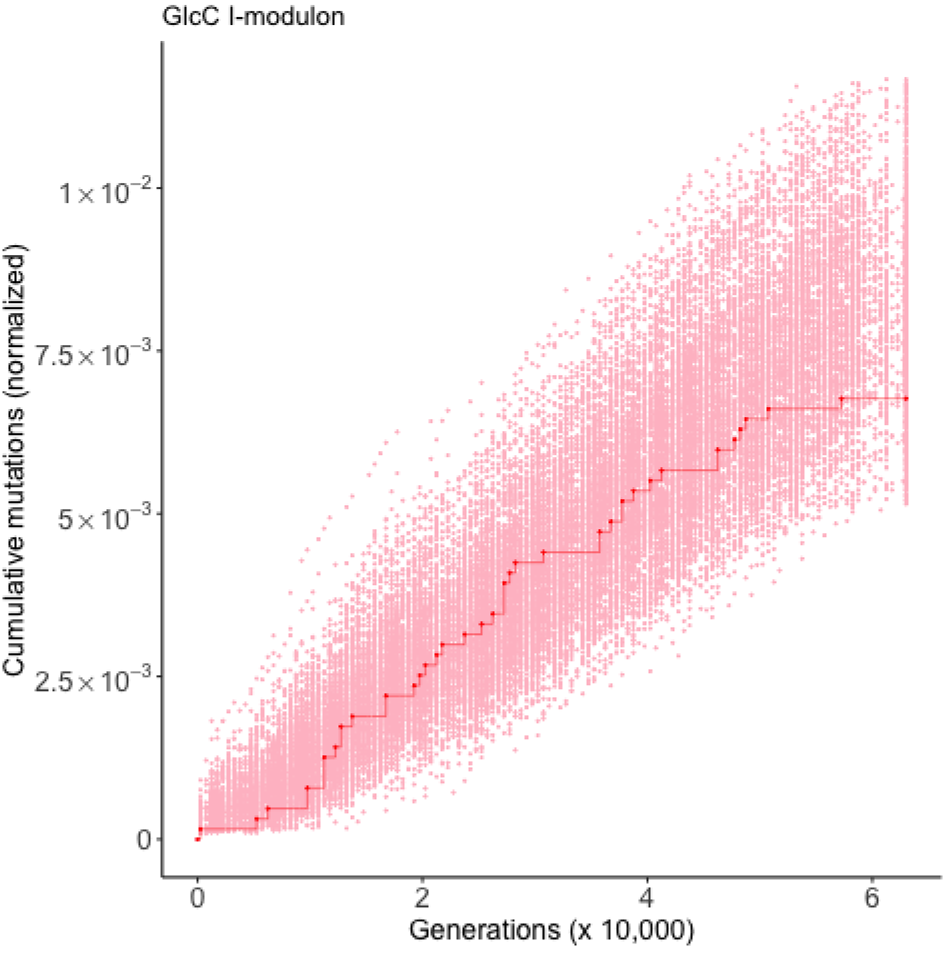
GlcC I-modulon
Cumulative mutations (normalized)
1×10^-2
7.5×10^-3
5×10^-3
2.5×10^-3
0
0
2
4
6
Generations (x 10,000)

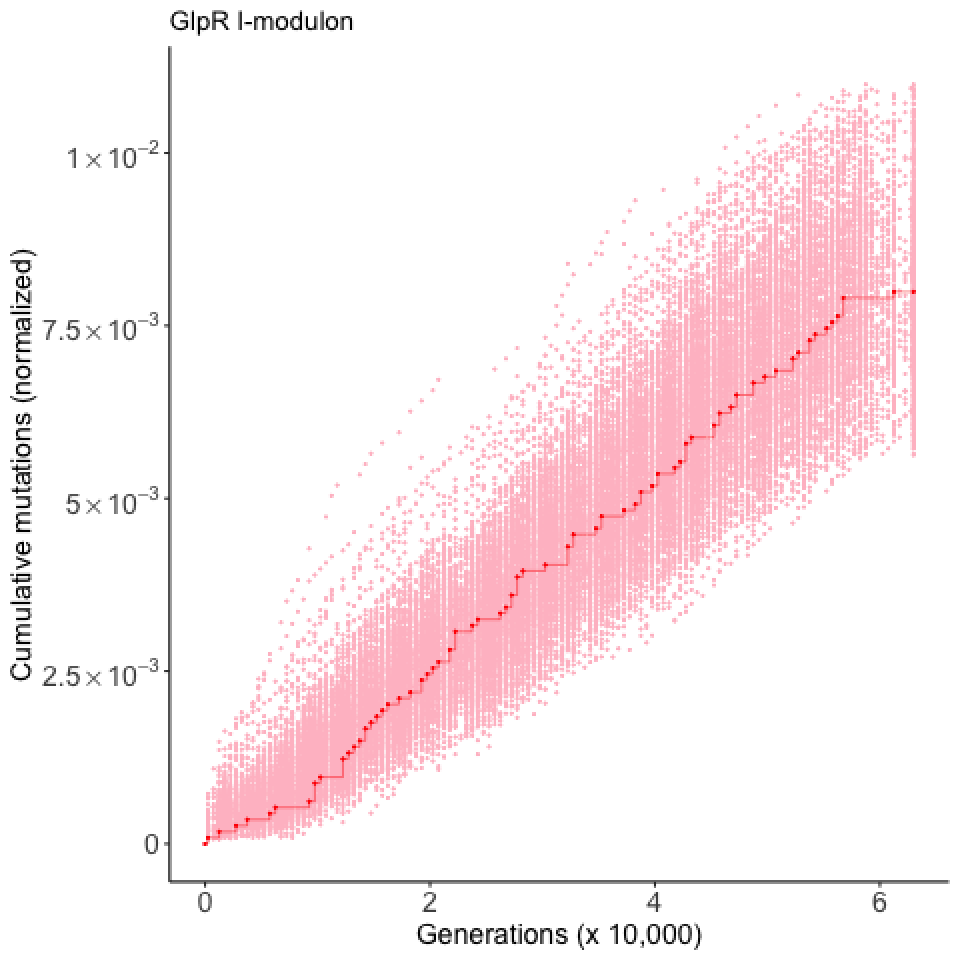

GlpR I-modulon
Cumulative mutations (normalized)
$1\times10^{-2}$
$7.5\times10^{-3}$
$5\times10^{-3}$
$2.5\times10^{-3}$
0
0
2
4
6
Generations (x 10,000)

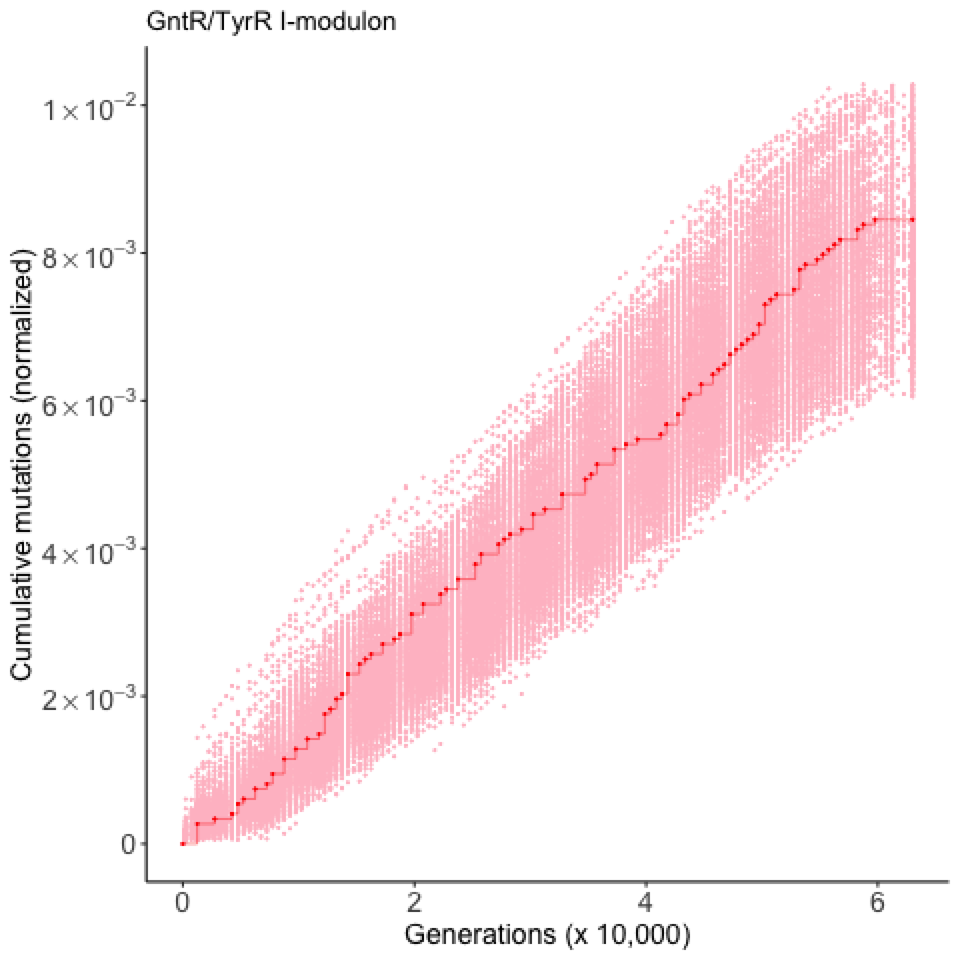
GntR/TyrR I-modulon
Cumulative mutations (normalized)
$1\times10^{-2}$
$8\times10^{-3}$
$6\times10^{-3}$
$4\times10^{-3}$
$2\times10^{-3}$
0
0
2
4
6
Generations (x 10,000)

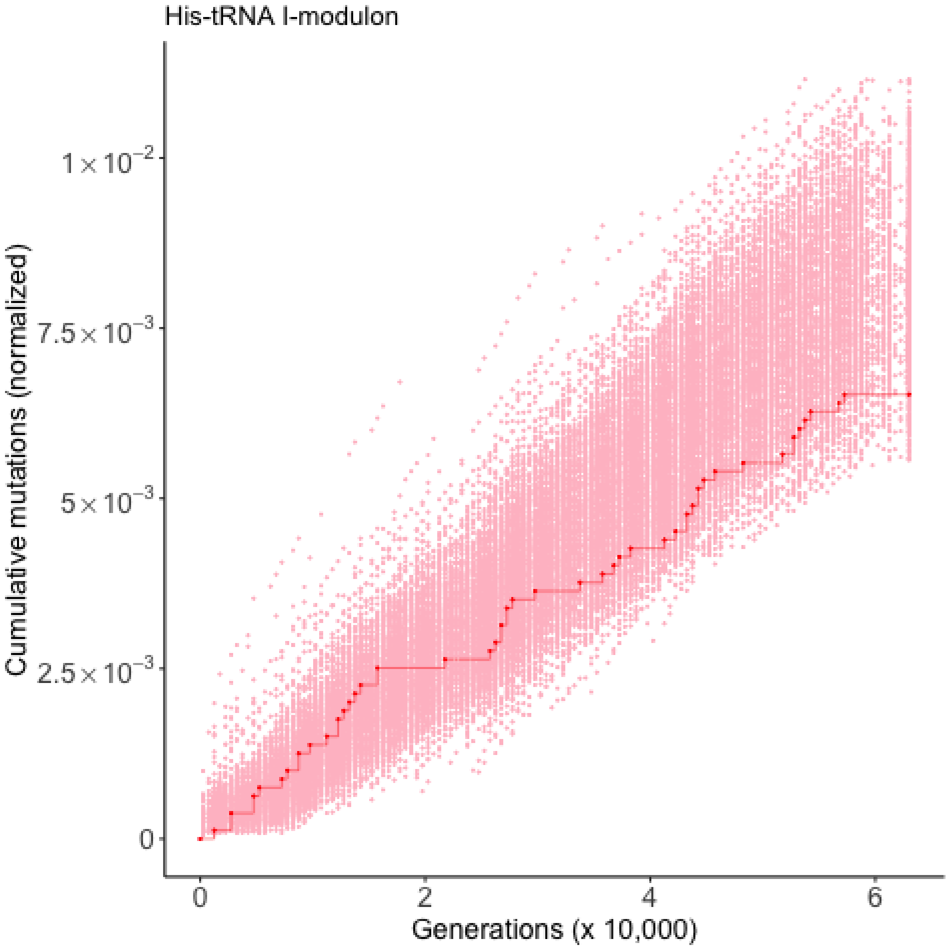

His-tRNA I-modulon
Cumulative mutations (normalized)
$1\times10^{-2}$
$7.5\times10^{-3}$
$5\times10^{-3}$
$2.5\times10^{-3}$
0
0
2
4
6
Generations (x 10,000)

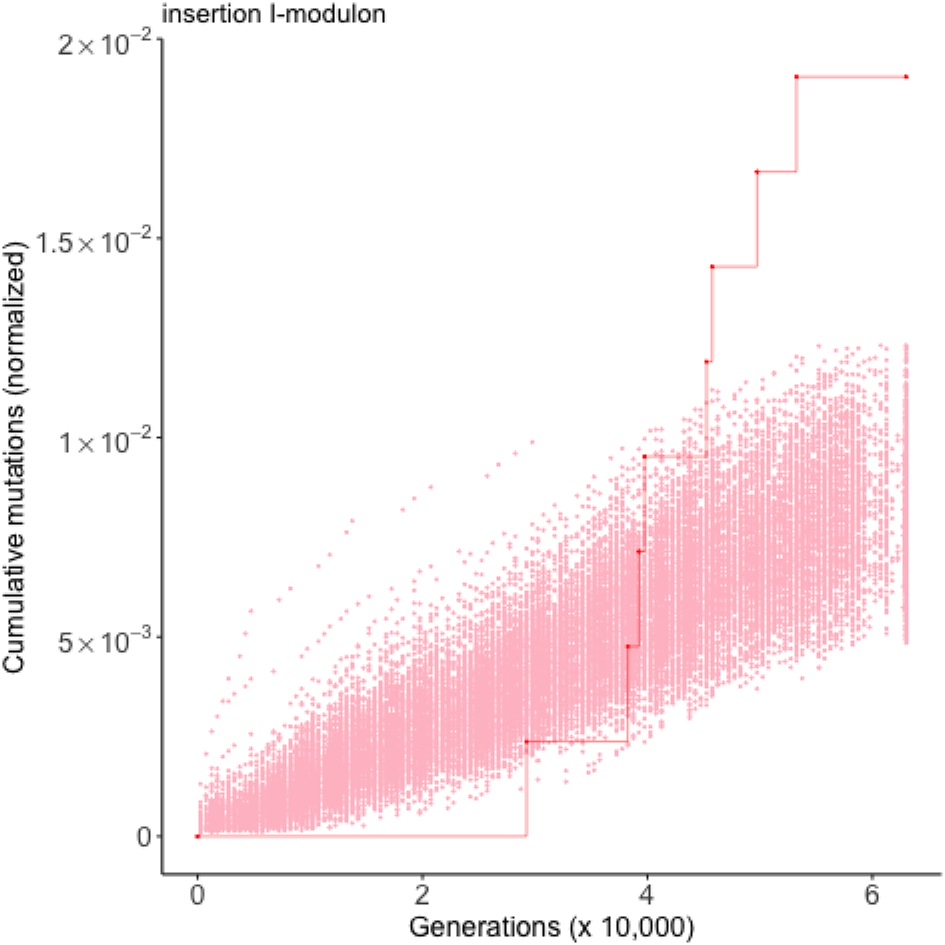

insertion I-modulon
$2\times10^{-2}$
$1.5\times10^{-2}$
$1\times10^{-2}$
$5\times10^{-3}$
0
0
2
4
6
Generations (x 10,000)
Cumulative mutations (normalized)

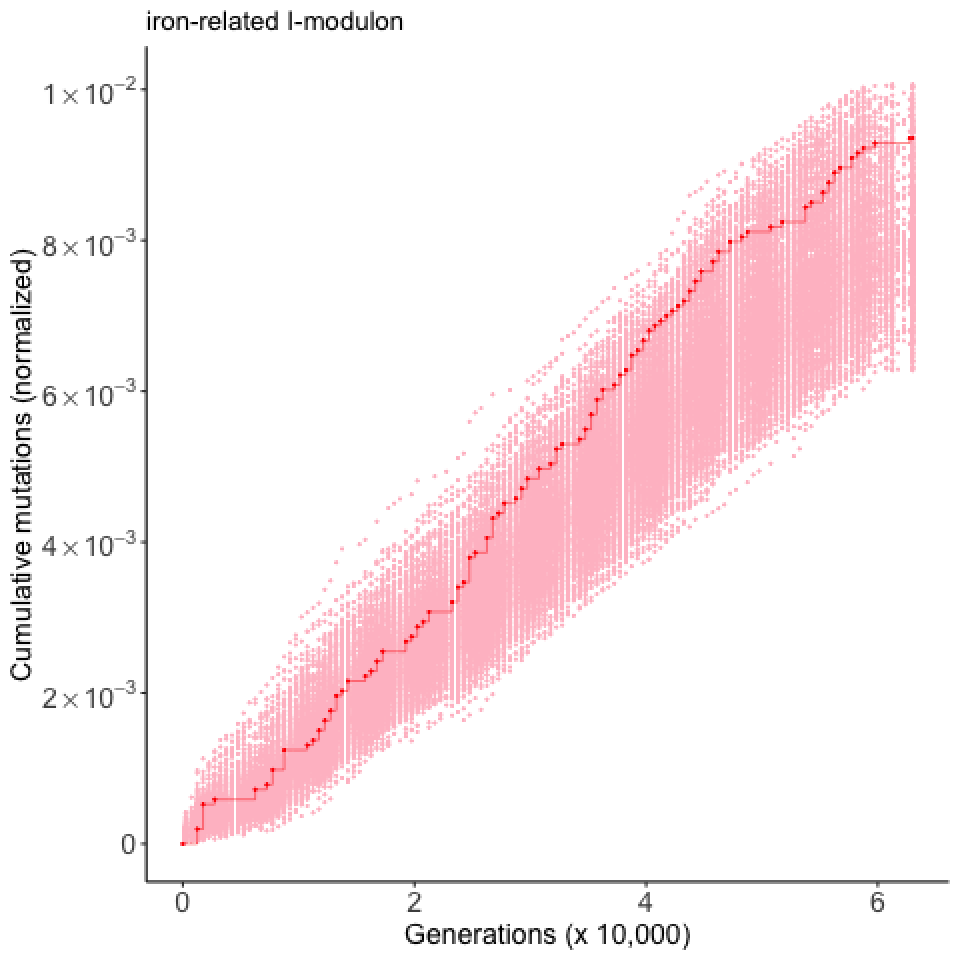

iron-related I-modulon
Cumulative mutations (normalized)
$1\times10^{-2}$
$8\times10^{-3}$
$6\times10^{-3}$
$4\times10^{-3}$
$2\times10^{-3}$
0
0
2
4
6
Generations (x 10,000)

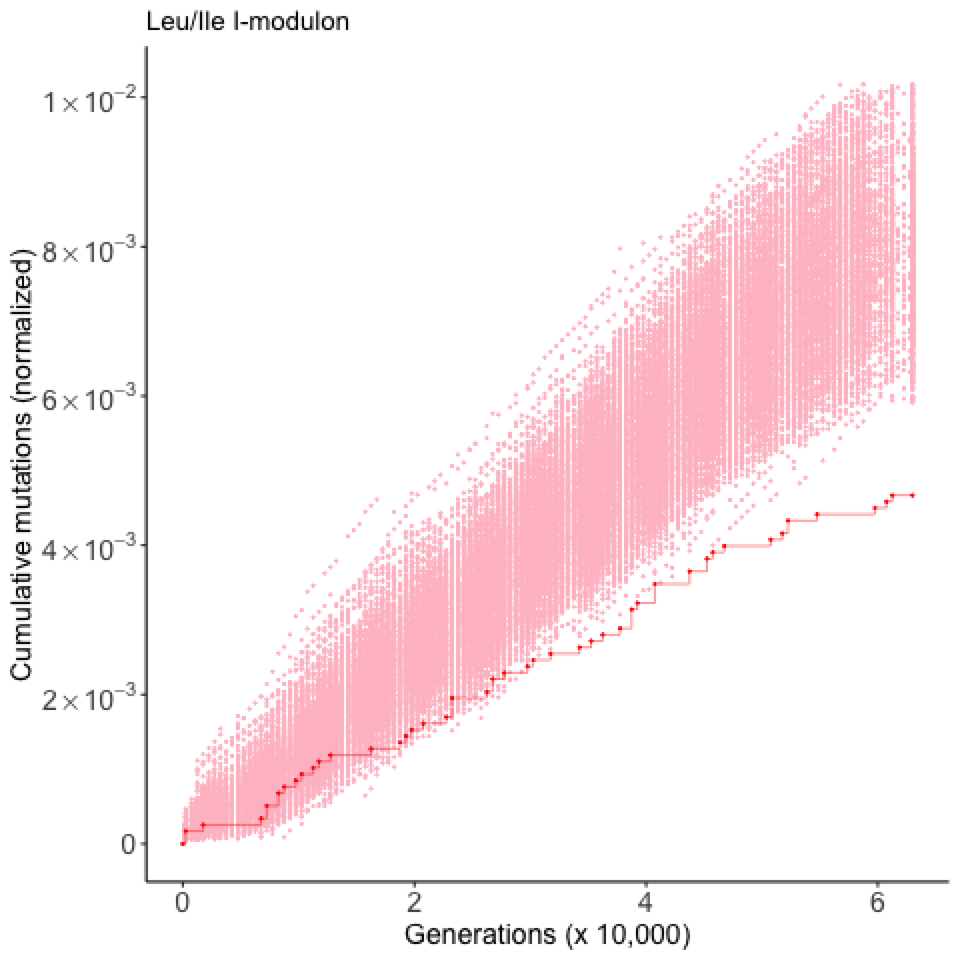

Leu/Ile I-modulon
Cumulative mutations (normalized)
1×10⁻²
8×10⁻³
6×10⁻³
4×10⁻³
2×10⁻³
0
0
2
4
6
Generations (x 10,000)

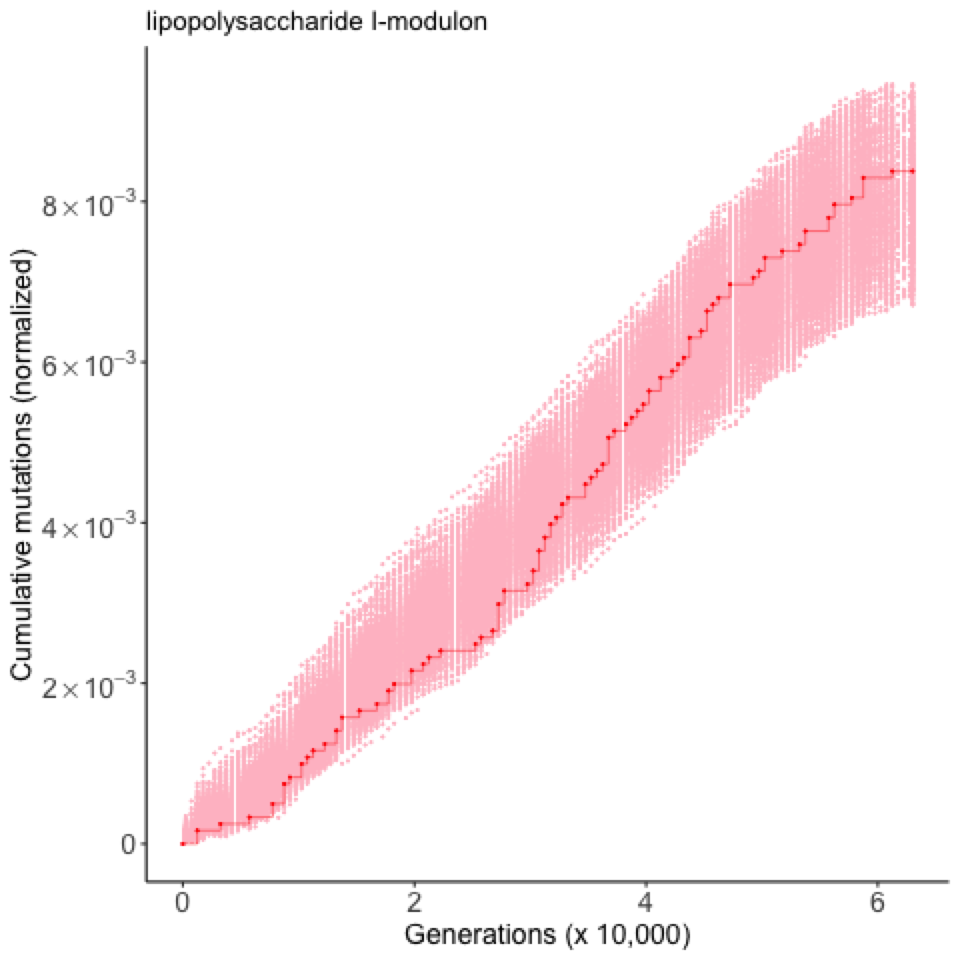

lipopolysaccharide I-modulon
Cumulative mutations (normalized)
$8\times10^{-3}$
$6\times10^{-3}$
$4\times10^{-3}$
$2\times10^{-3}$
0
0
2
4
6
Generations (x 10,000)

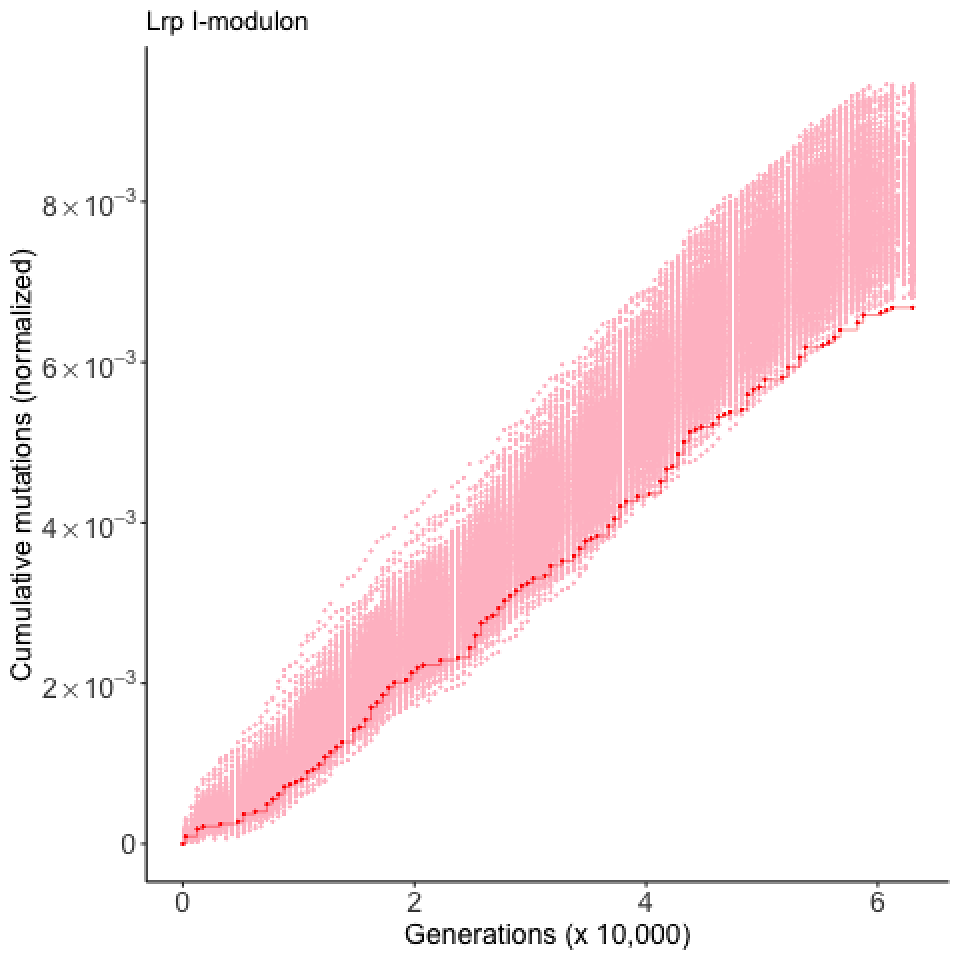

Lrp I-modulon
Cumulative mutations (normalized)
$8 \times 10^{-3}$
$6 \times 10^{-3}$
$4 \times 10^{-3}$
$2 \times 10^{-3}$
0
0
2
4
6
Generations (x 10,000)

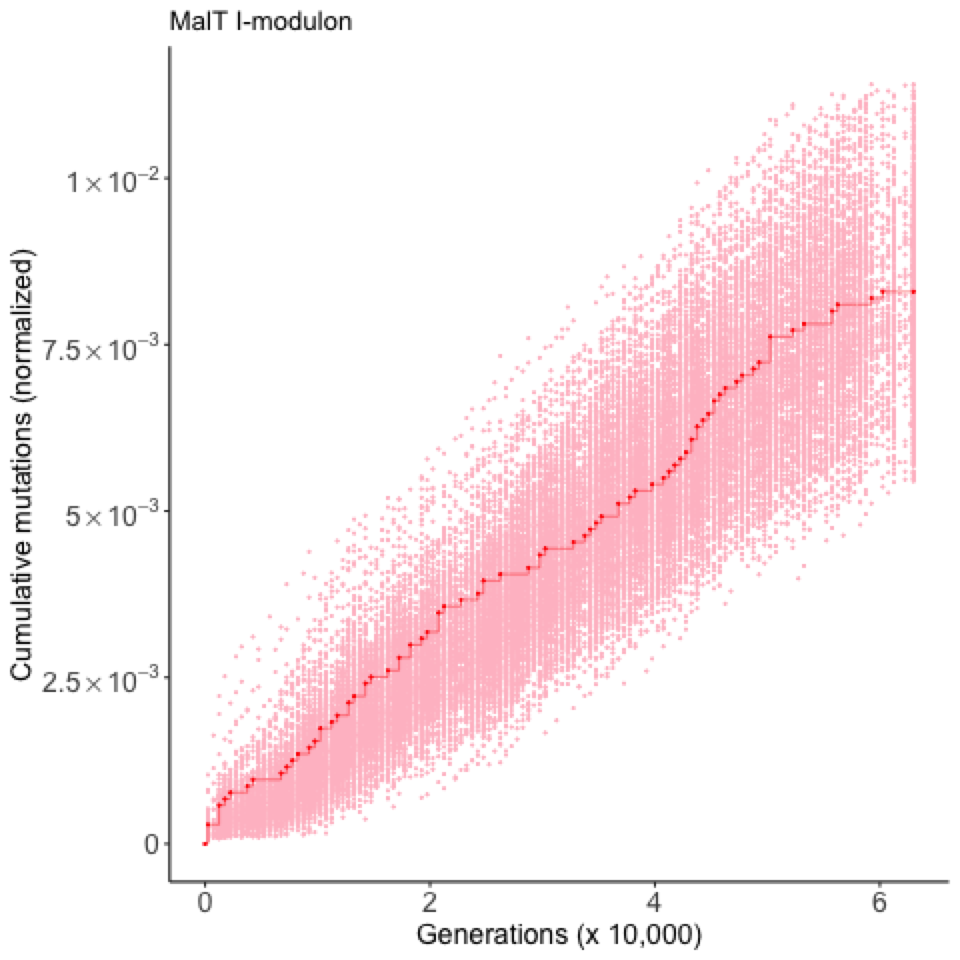

MalT I-modulon
Cumulative mutations (normalized)
1×10^-2
7.5×10^-3
5×10^-3
2.5×10^-3
0
0
2
4
6
Generations (x 10,000)

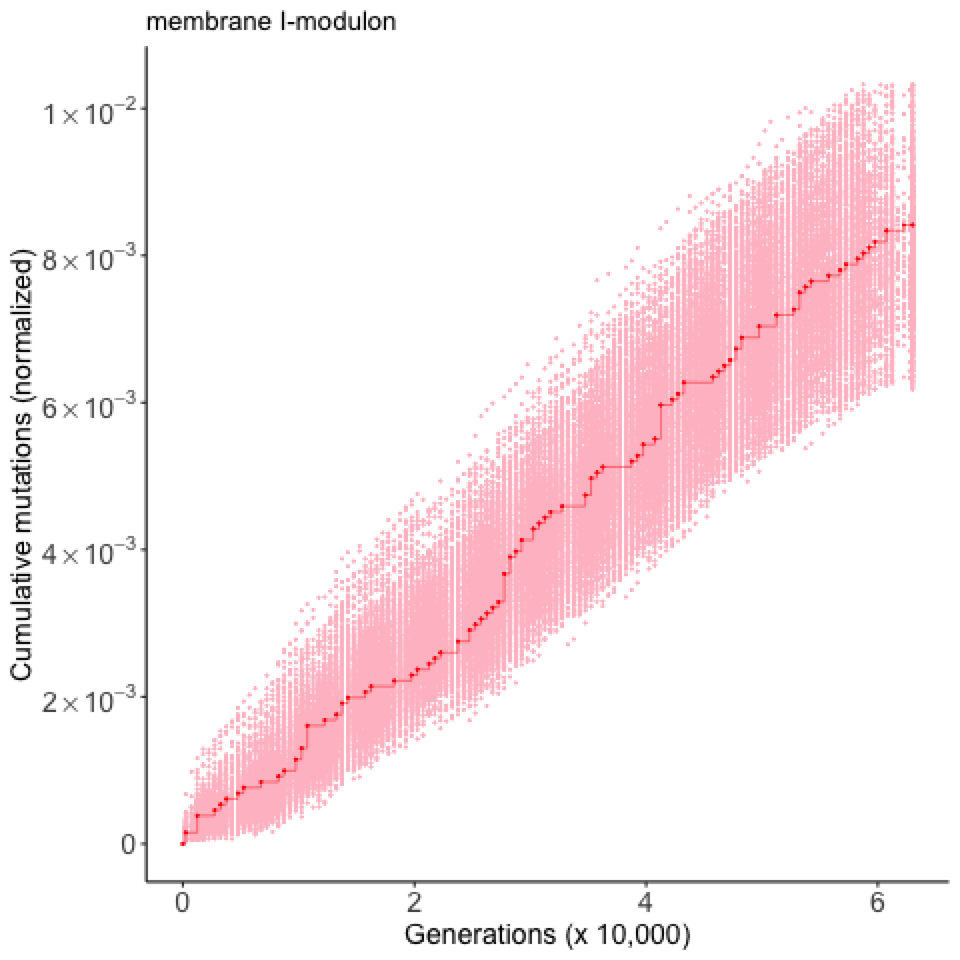
membrane I-modulon
Cumulative mutations (normalized)
$1\times10^{-2}$
$8\times10^{-3}$
$6\times10^{-3}$
$4\times10^{-3}$
$2\times10^{-3}$
0
0
2
4
6
Generations (x 10,000)

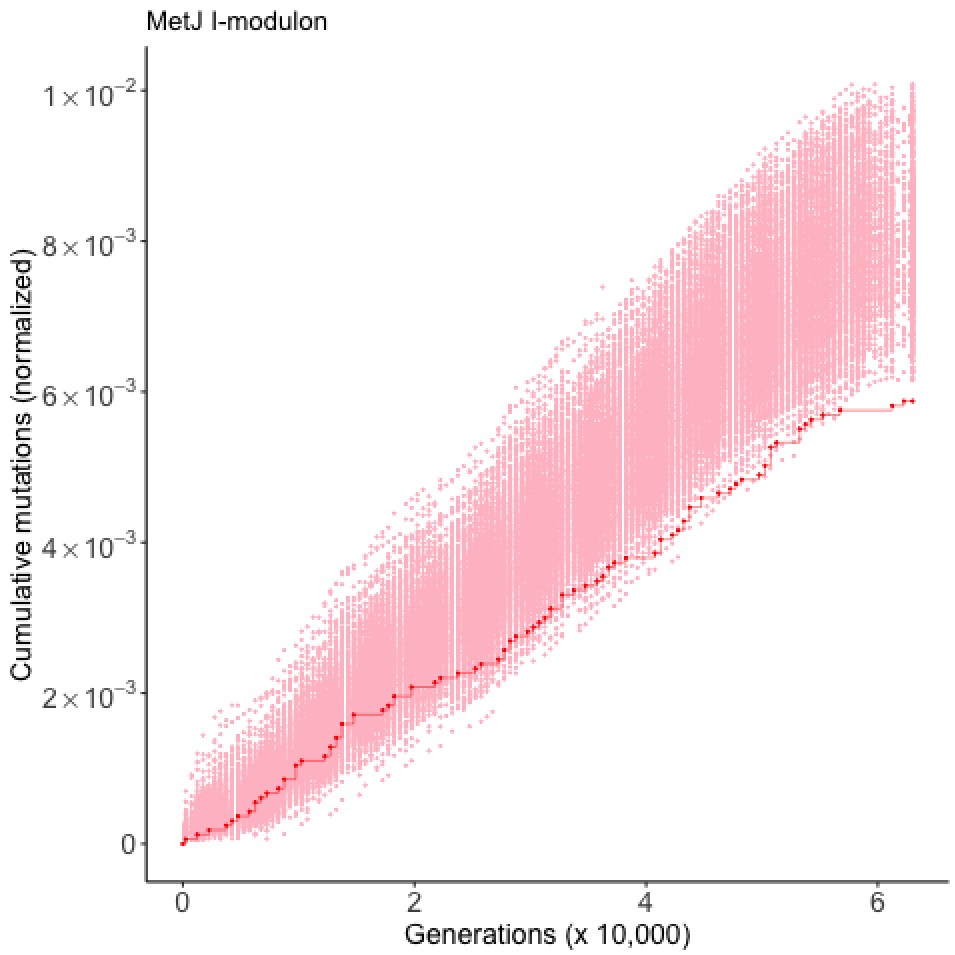

MetJ I-modulon
Cumulative mutations (normalized)
1×10^-2
8×10^-3
6×10^-3
4×10^-3
2×10^-3
0
0
2
4
6
Generations (x 10,000)

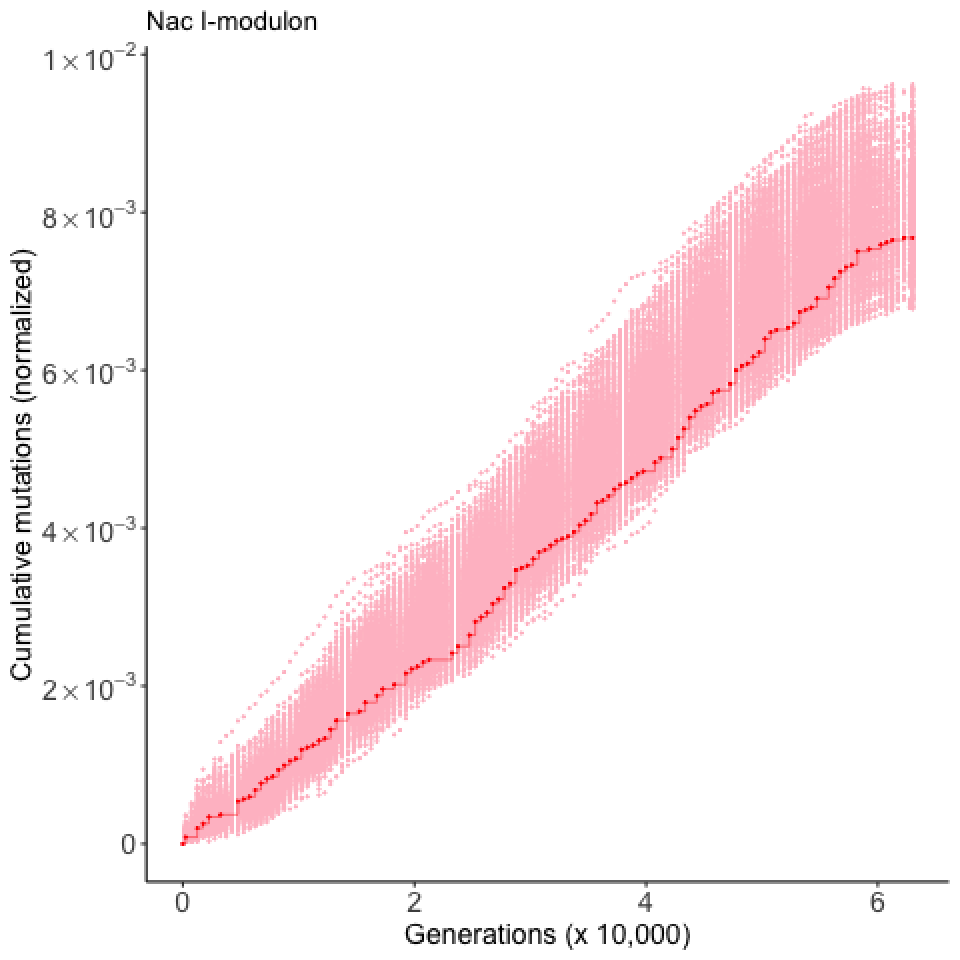

Nac I-modulon
Cumulative mutations (normalized)
$1\times10^{-2}$
$8\times10^{-3}$
$6\times10^{-3}$
$4\times10^{-3}$
$2\times10^{-3}$
0
0
2
4
6
Generations (x 10,000)

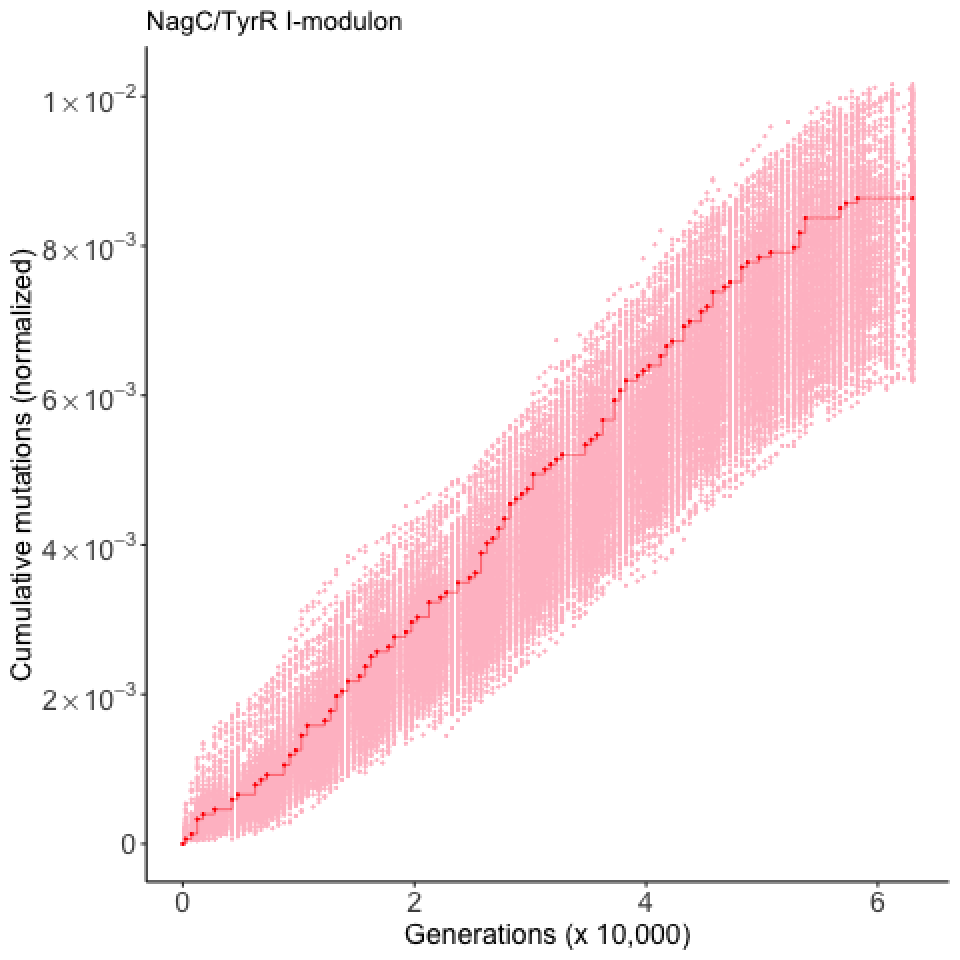
NagC/TyrR I-modulon
Cumulative mutations (normalized)
$1\times10^{-2}$
$8\times10^{-3}$
$6\times10^{-3}$
$4\times10^{-3}$
$2\times10^{-3}$
0
0
2
4
6
Generations (x 10,000)

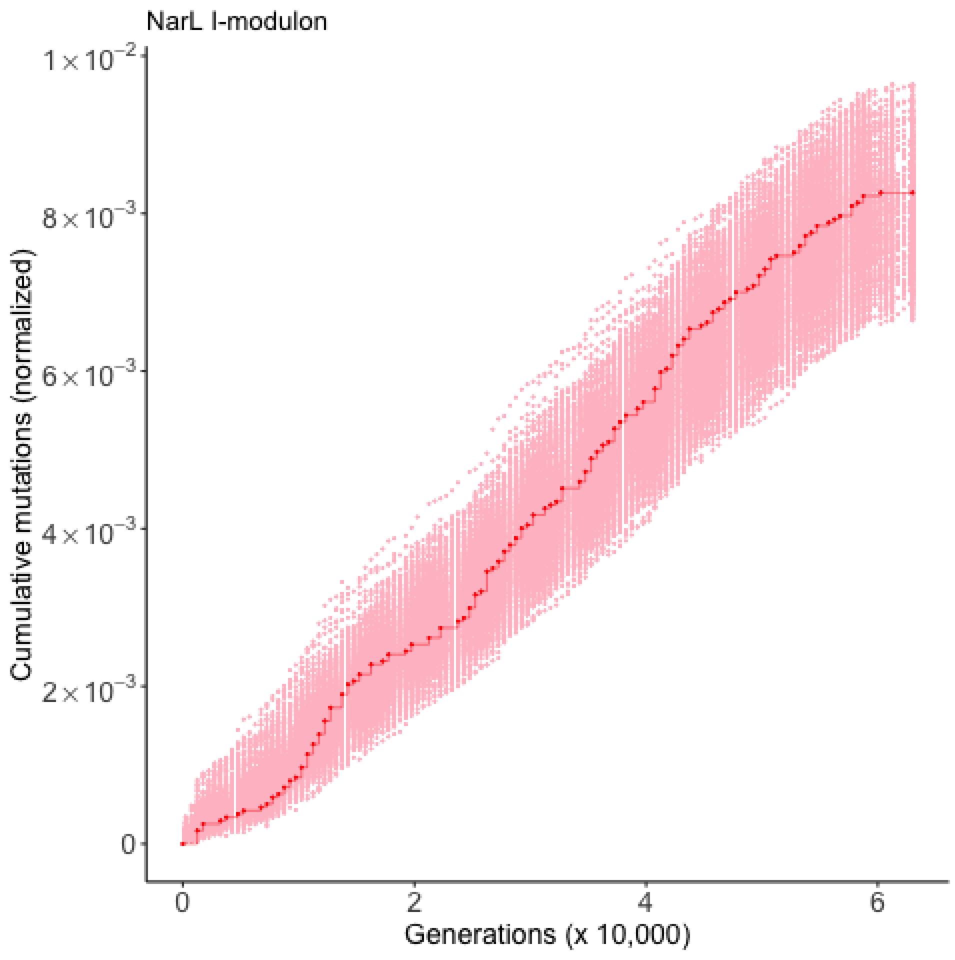

NarL I-modulon
Cumulative mutations (normalized)
$1\times10^{-2}$
$8\times10^{-3}$
$6\times10^{-3}$
$4\times10^{-3}$
$2\times10^{-3}$
0
0
2
4
6
Generations (x 10,000)

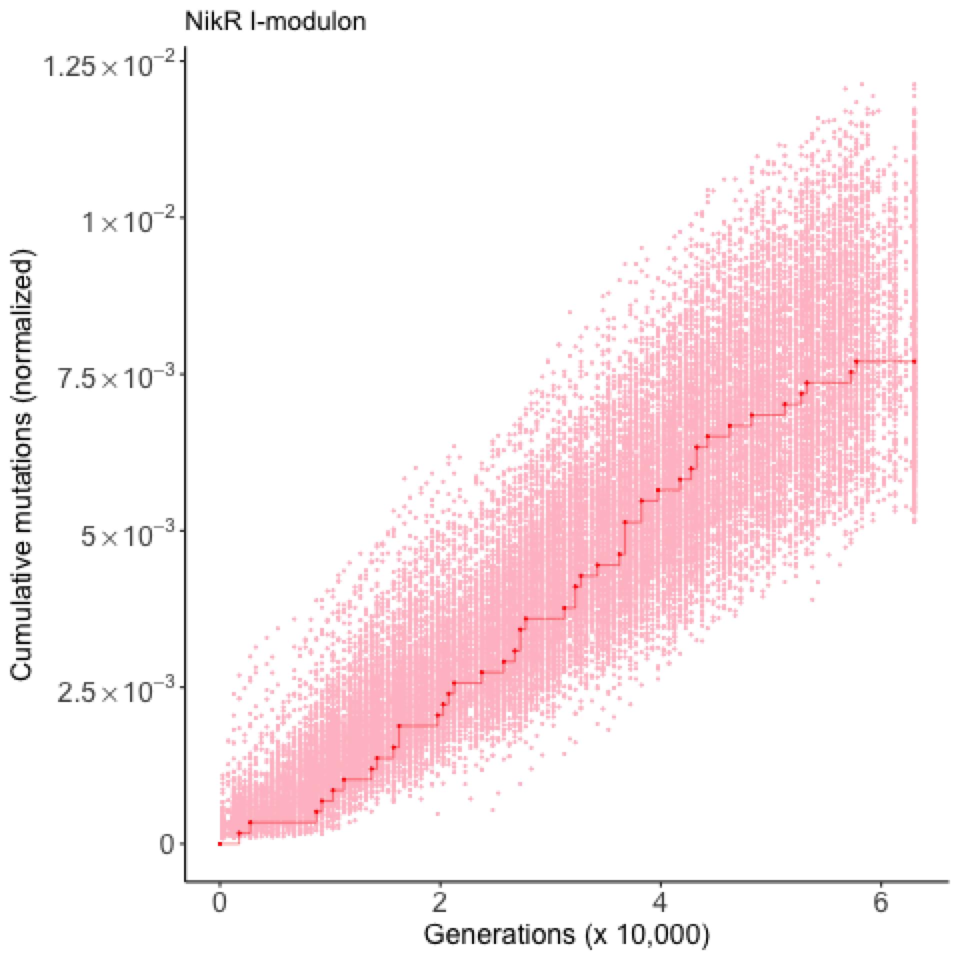

NikR I-modulon
Cumulative mutations (normalized)
$1.25 \times 10^{-2}$
$1 \times 10^{-2}$
$7.5 \times 10^{-3}$
$5 \times 10^{-3}$
$2.5 \times 10^{-3}$
0
0
2
4
6
Generations (x 10,000)

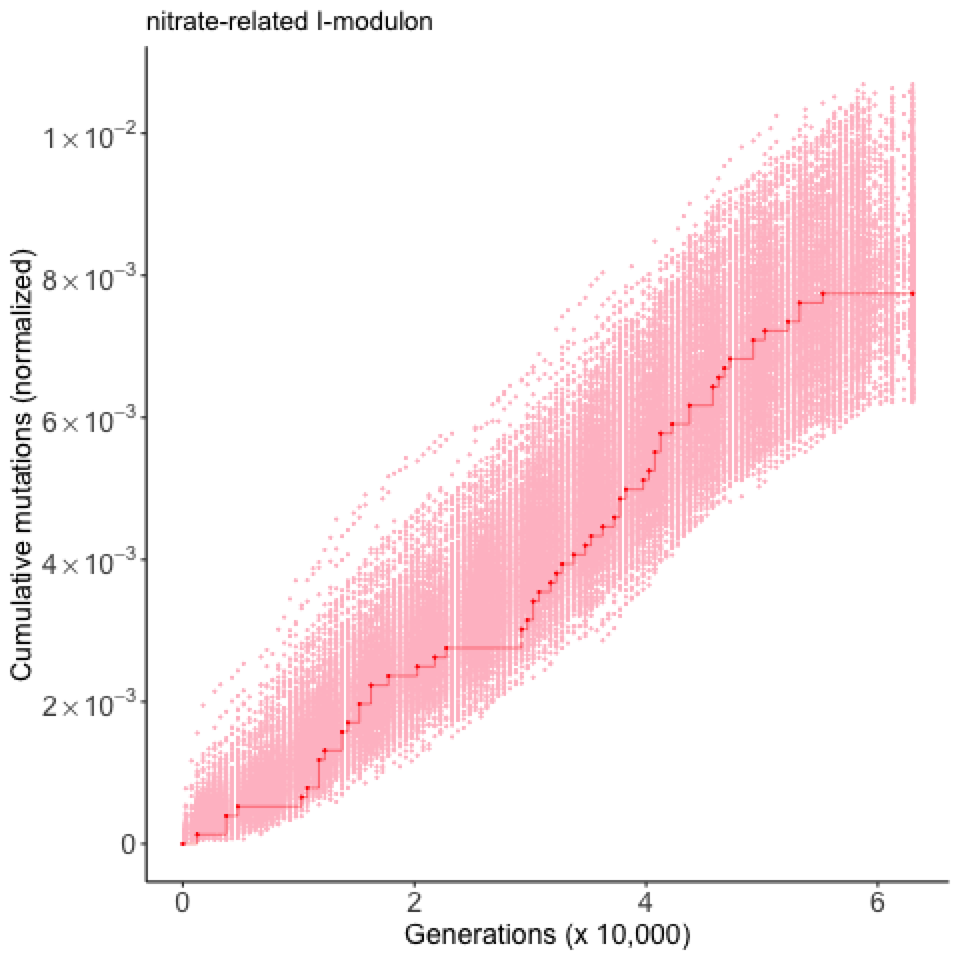

nitrate-related I-modulon
Cumulative mutations (normalized)
$1\times10^{-2}$
$8\times10^{-3}$
$6\times10^{-3}$
$4\times10^{-3}$
$2\times10^{-3}$
0
0
2
4
6
Generations (x 10,000)

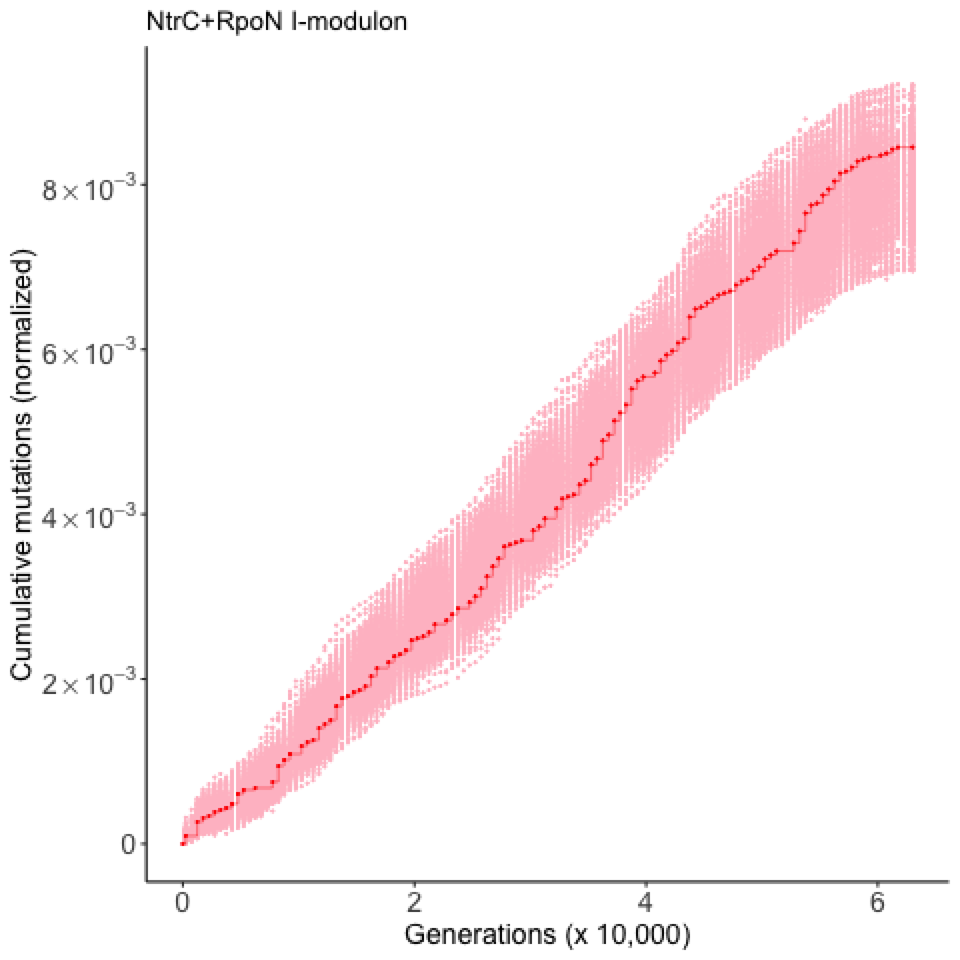

NtrC+RpoN I-modulon
Cumulative mutations (normalized)
8×10⁻³
6×10⁻³
4×10⁻³
2×10⁻³
0
0
2
4
6
Generations (x 10,000)

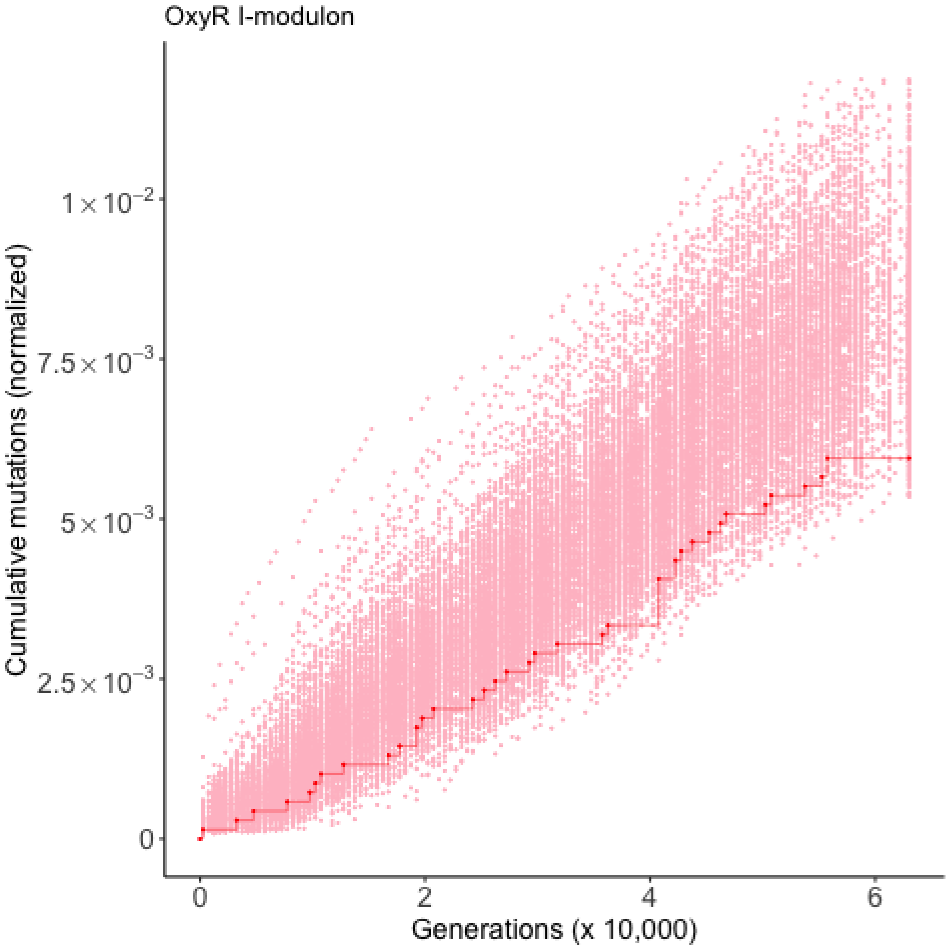
OxyR I-modulon
Cumulative mutations (normalized)
$1\times10^{-2}$
$7.5\times10^{-3}$
$5\times10^{-3}$
$2.5\times10^{-3}$
0
0
2
4
6
Generations (x 10,000)

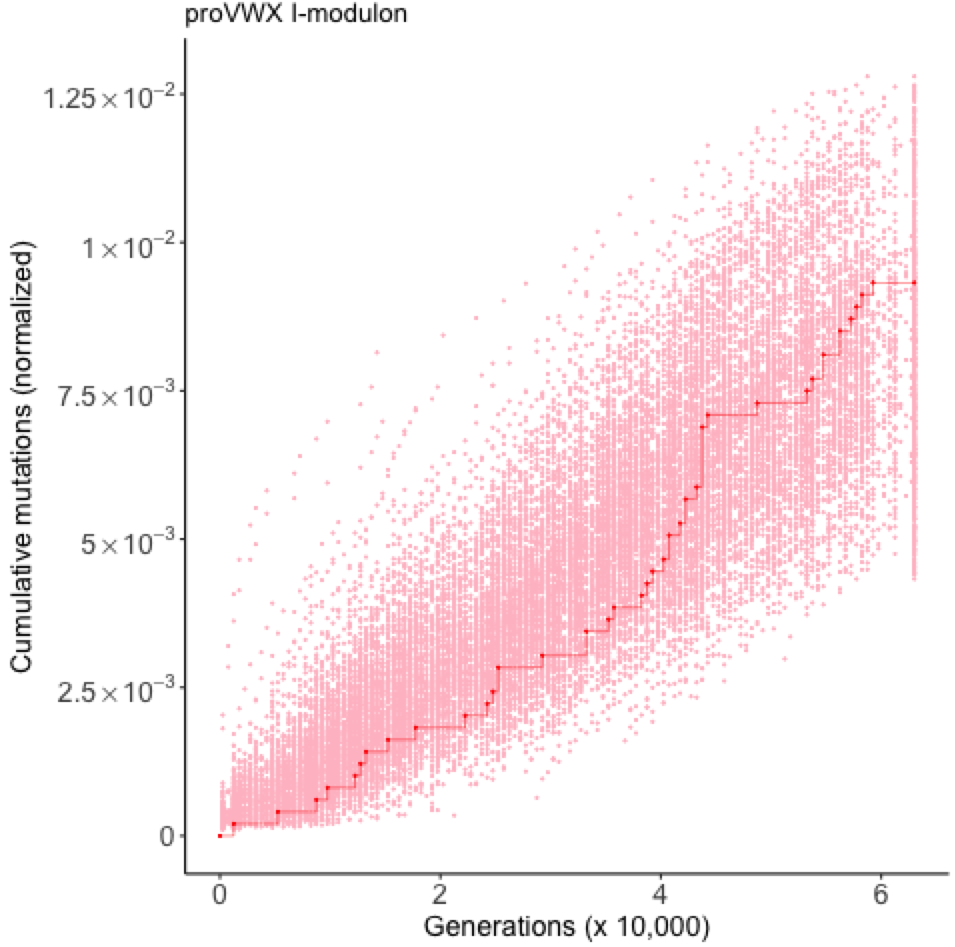
proVWX I-modulon
Cumulative mutations (normalized)
1.25×10^-2
1×10^-2
7.5×10^-3
5×10^-3
2.5×10^-3
0
0
2
4
6
Generations (x 10,000)

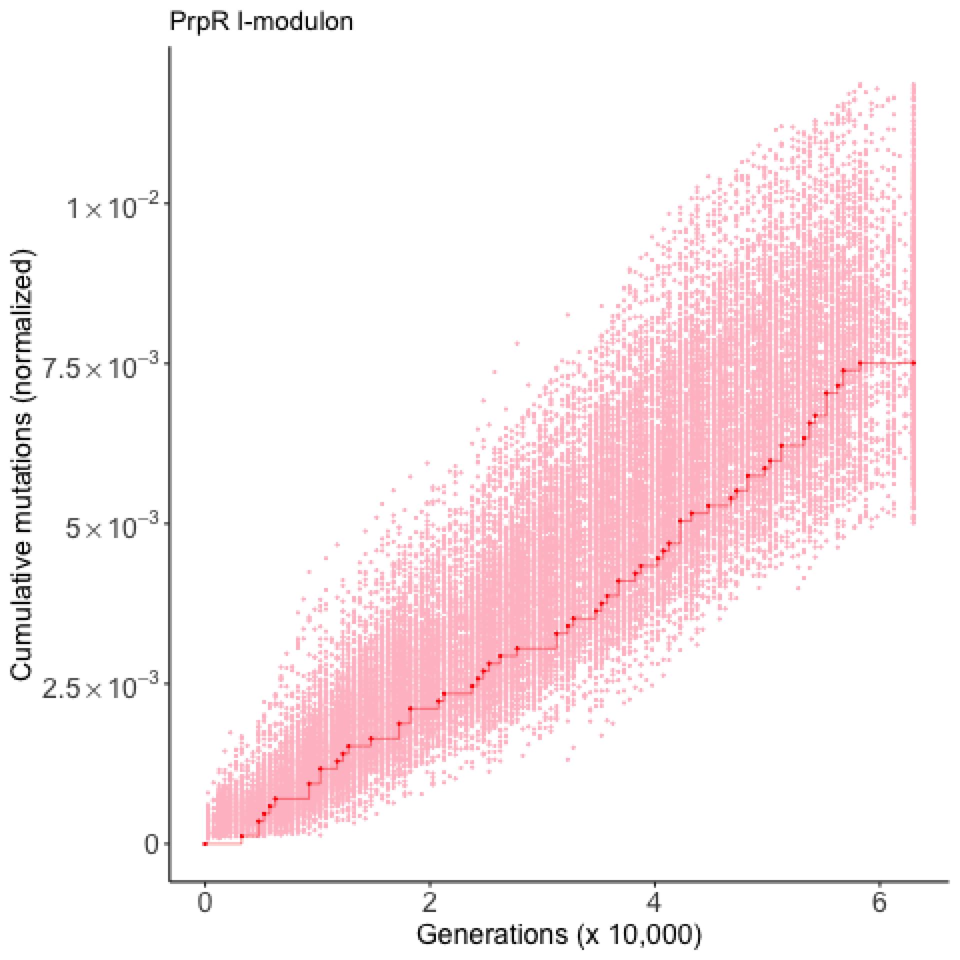
PrpR I-modulon
Cumulative mutations (normalized)
$1\times10^{-2}$
$7.5\times10^{-3}$
$5\times10^{-3}$
$2.5\times10^{-3}$
0
0
2
4
6
Generations (x 10,000)

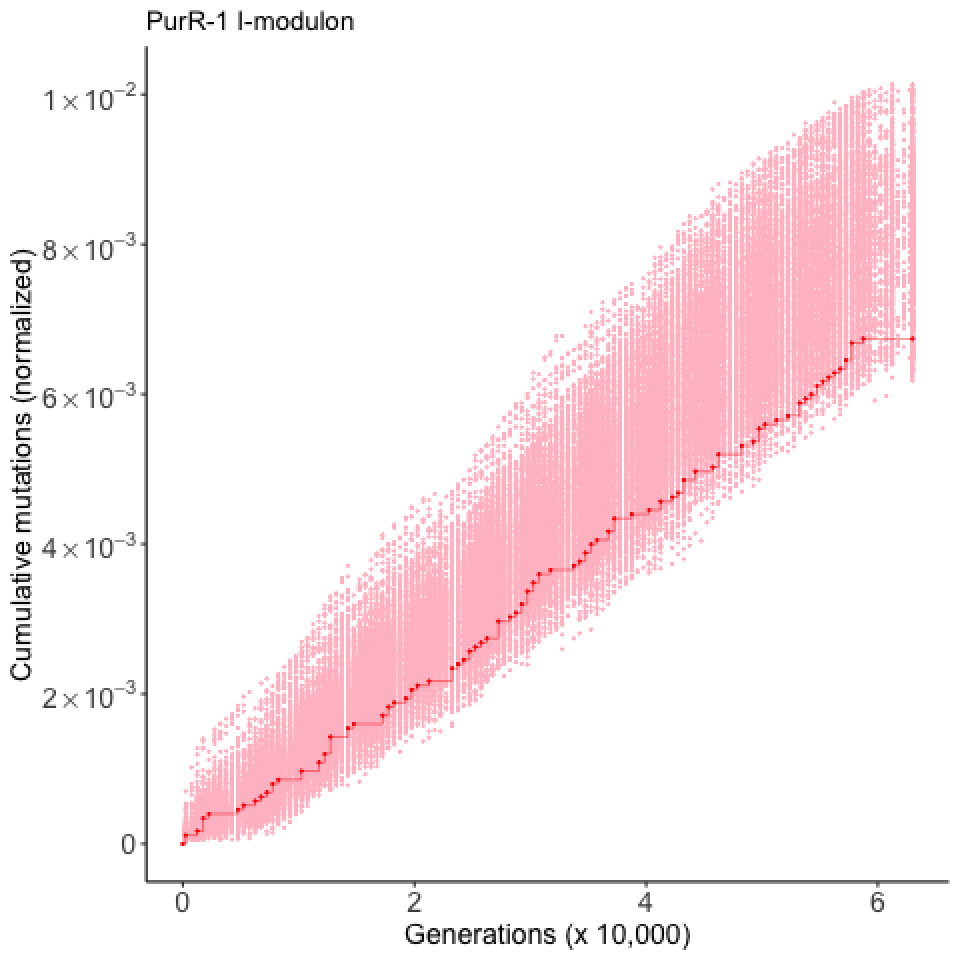
PurR-1 I-modulon
Cumulative mutations (normalized)
$1\times10^{-2}$
$8\times10^{-3}$
$6\times10^{-3}$
$4\times10^{-3}$
$2\times10^{-3}$
0
0
2
4
6
Generations (x 10,000)

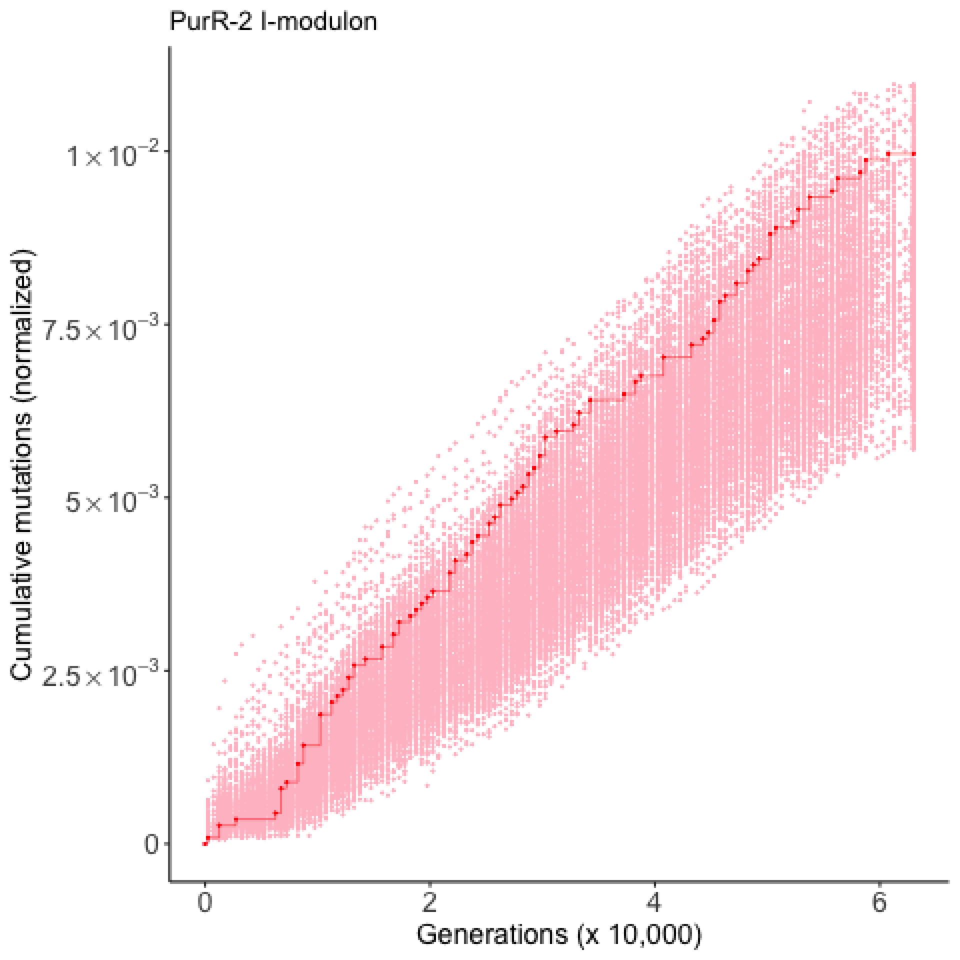

PurR-2 I-modulon
Cumulative mutations (normalized)
$1\times10^{-2}$
$7.5\times10^{-3}$
$5\times10^{-3}$
$2.5\times10^{-3}$
0
0
2
4
6
Generations (x 10,000)

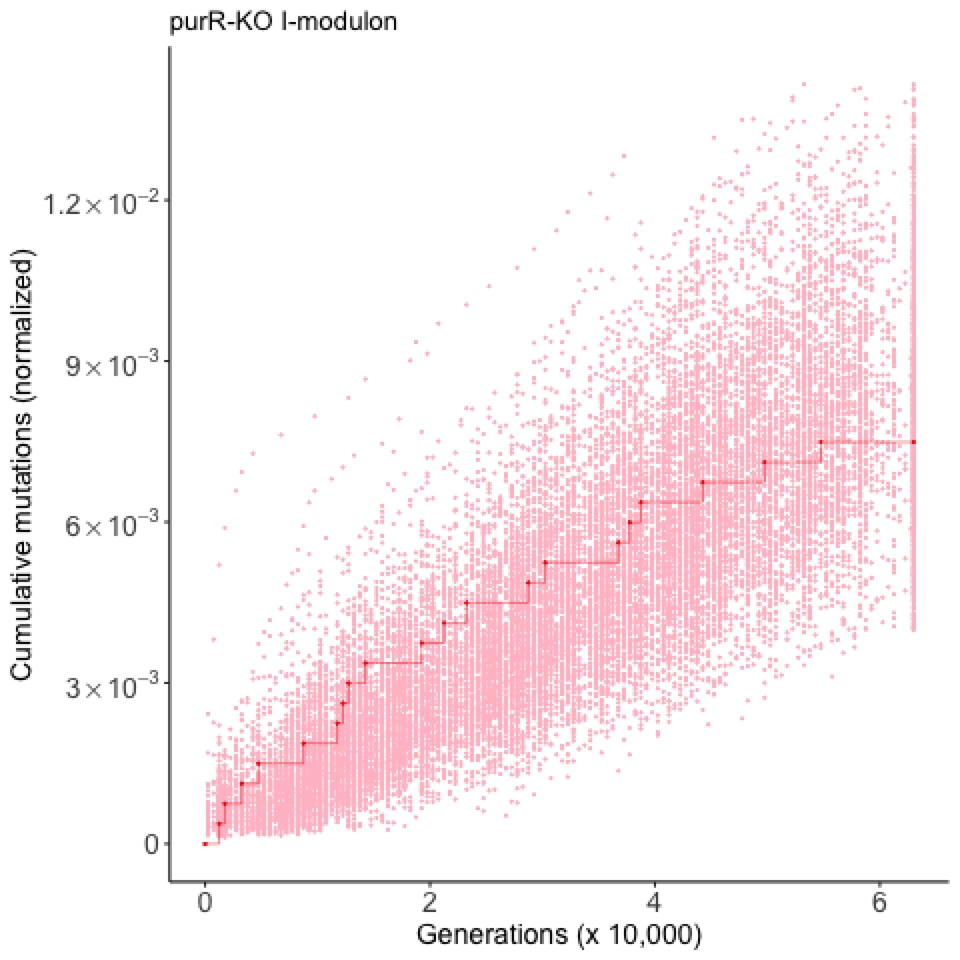

purR-KO I-modulon
Cumulative mutations (normalized)
$1.2\times10^{-2}$
$9\times10^{-3}$
$6\times10^{-3}$
$3\times10^{-3}$
0
0
2
4
6
Generations (x 10,000)

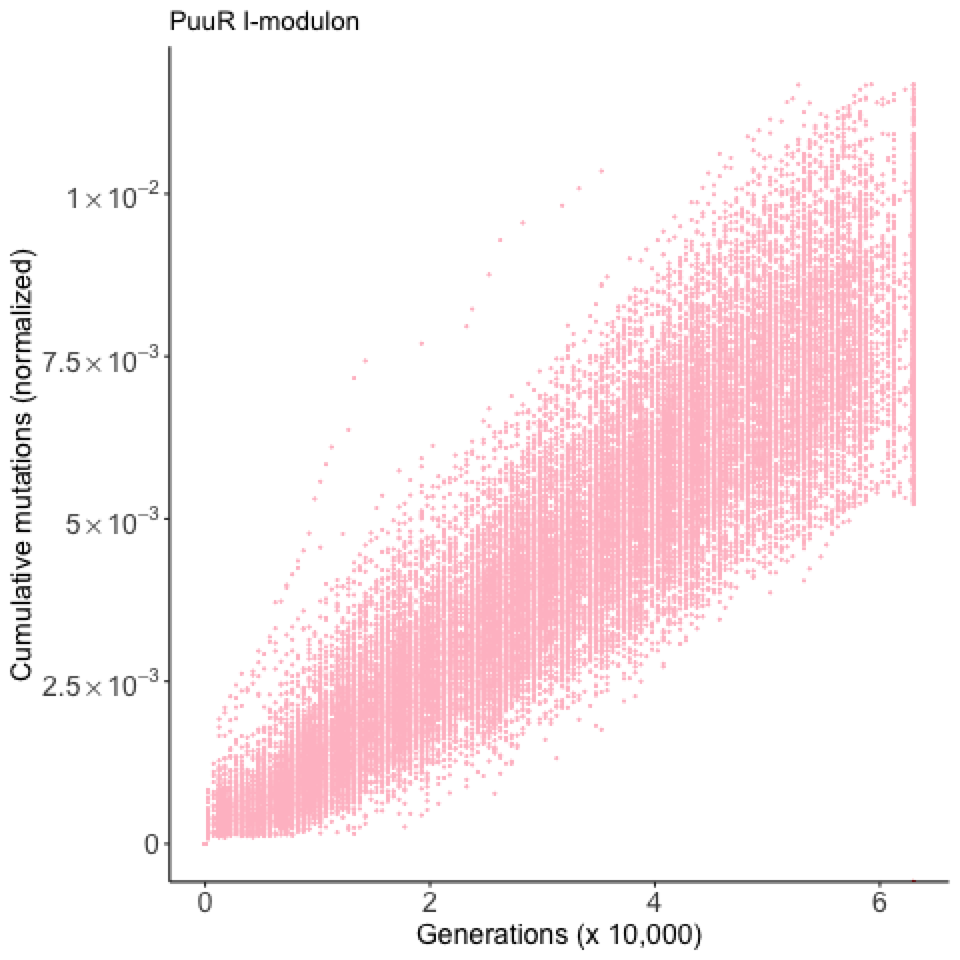
PuuR I-modulon
Cumulative mutations (normalized)
$1\times10^{-2}$
$7.5\times10^{-3}$
$5\times10^{-3}$
$2.5\times10^{-3}$
0
0
2
4
6
Generations (x 10,000)

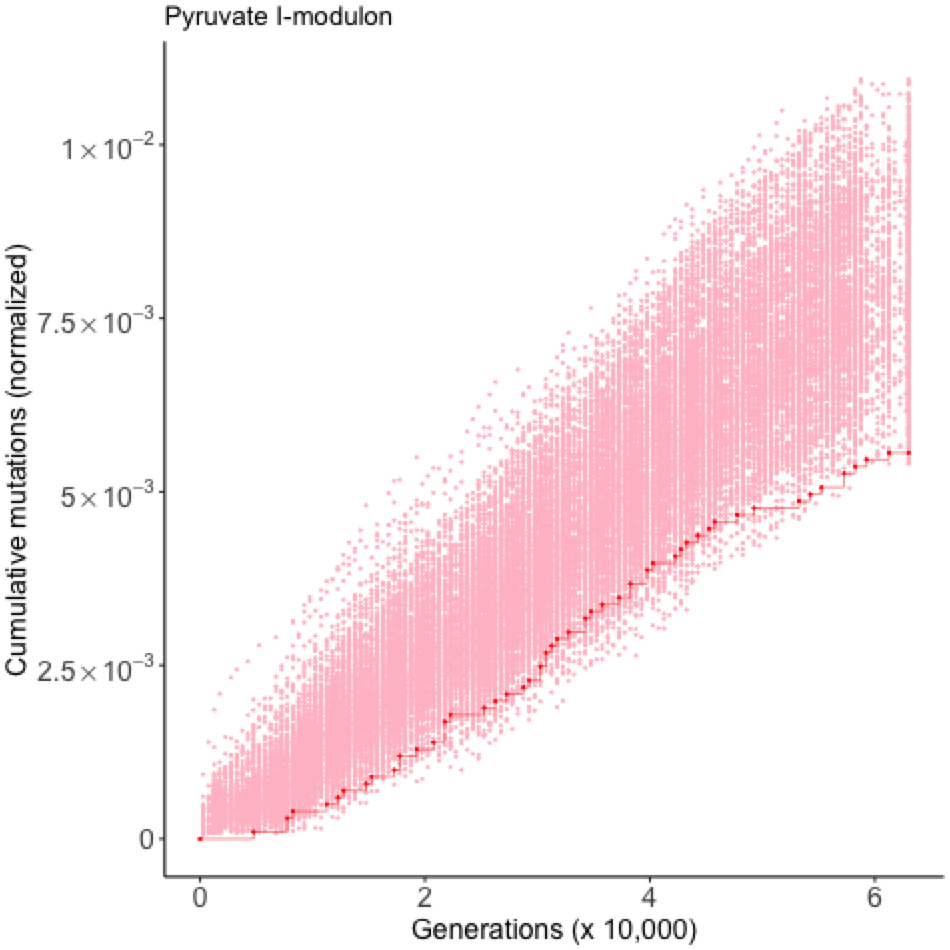
Pyruvate I-modulon
Cumulative mutations (normalized)
$1\times10^{-2}$
$7.5\times10^{-3}$
$5\times10^{-3}$
$2.5\times10^{-3}$
0
0
2
4
6
Generations (x 10,000)

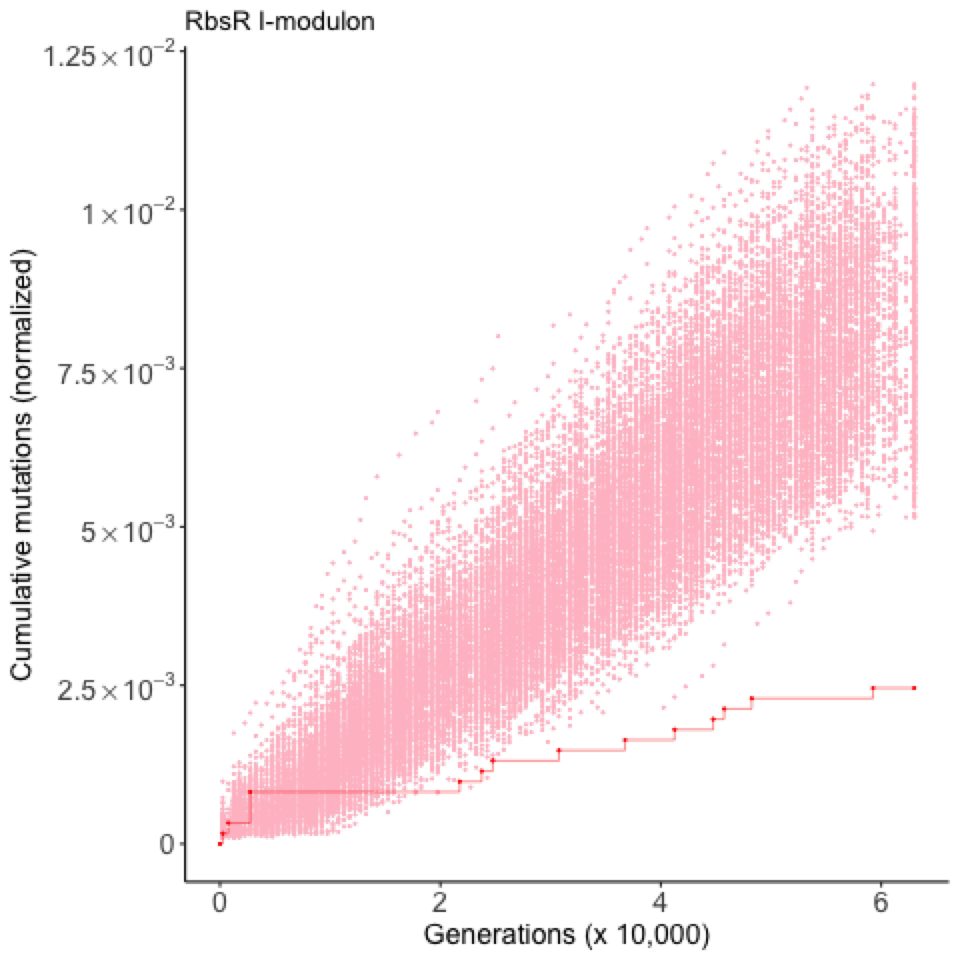

RbsR I-modulon
Cumulative mutations (normalized)
1.25×10^-2
1×10^-2
7.5×10^-3
5×10^-3
2.5×10^-3
0
0
2
4
6
Generations (x 10,000)

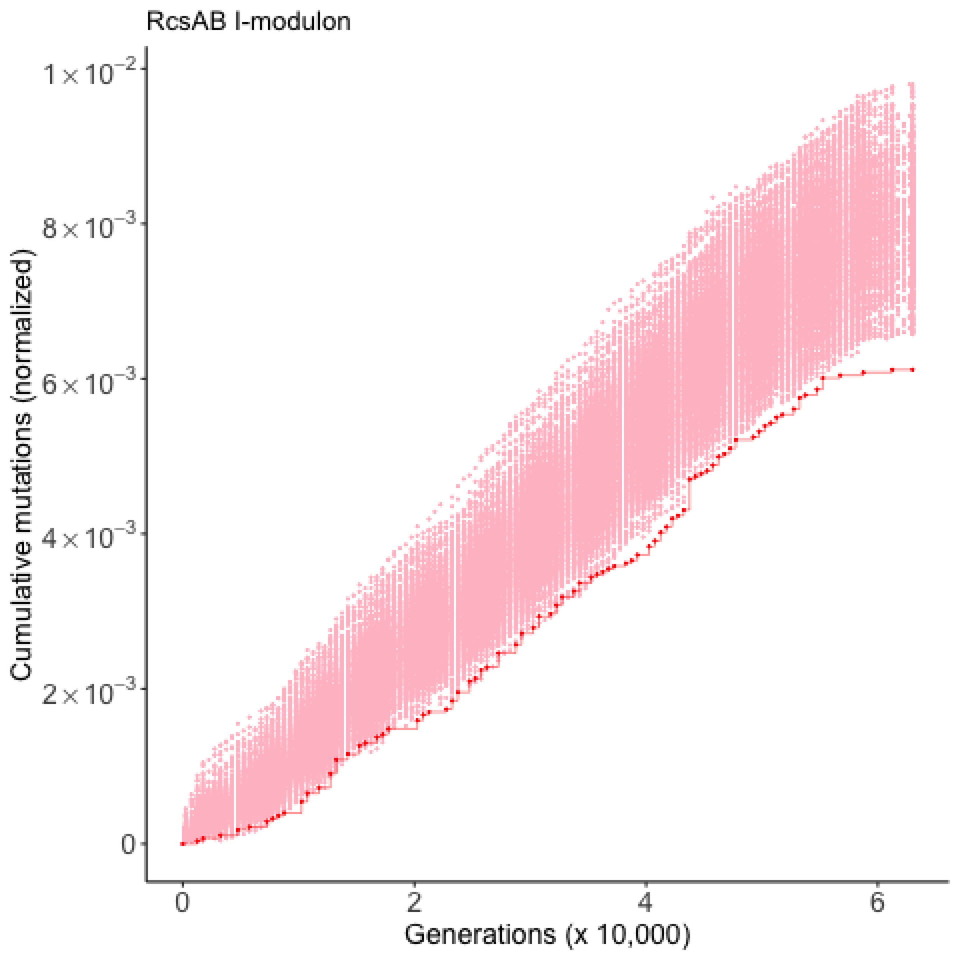

RcsAB I-modulon
Cumulative mutations (normalized)
$1\times10^{-2}$
$8\times10^{-3}$
$6\times10^{-3}$
$4\times10^{-3}$
$2\times10^{-3}$
0
0
2
4
6
Generations (x 10,000)

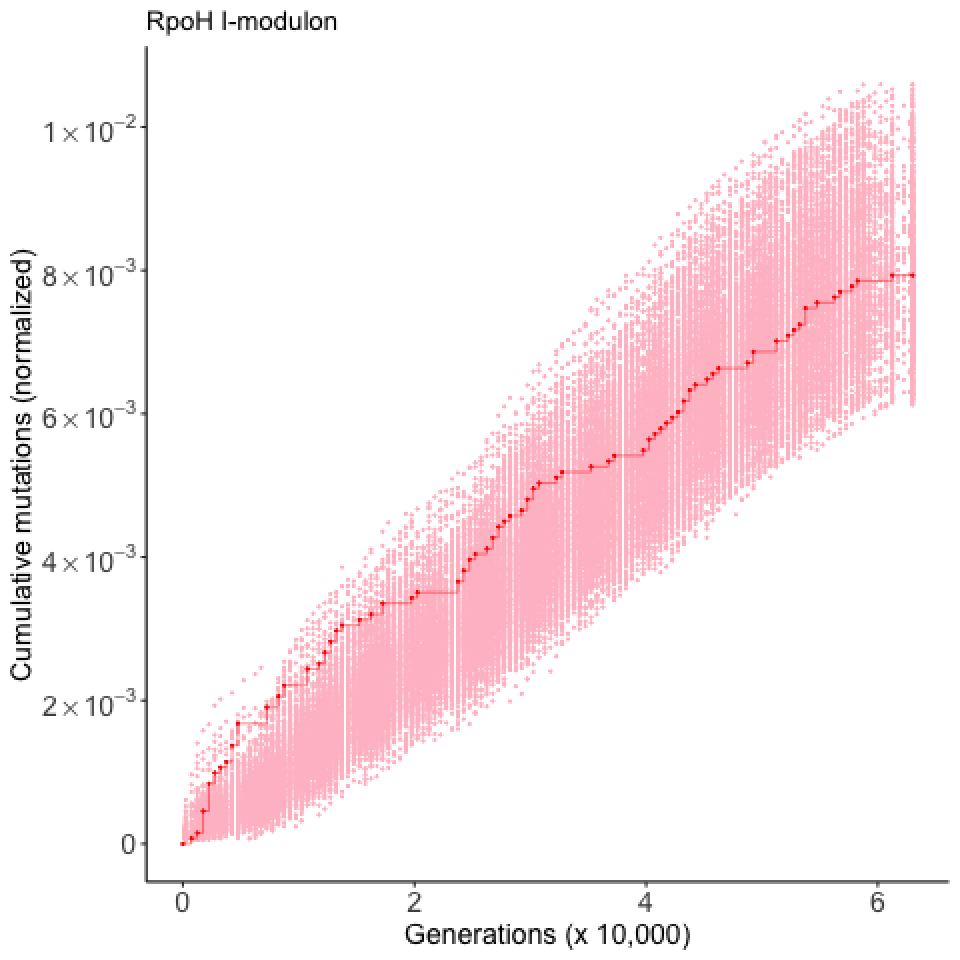

RpoH I-modulon
Cumulative mutations (normalized)
$1\times10^{-2}$
$8\times10^{-3}$
$6\times10^{-3}$
$4\times10^{-3}$
$2\times10^{-3}$
0
0
2
4
6
Generations (x 10,000)

RpoS I-modulon

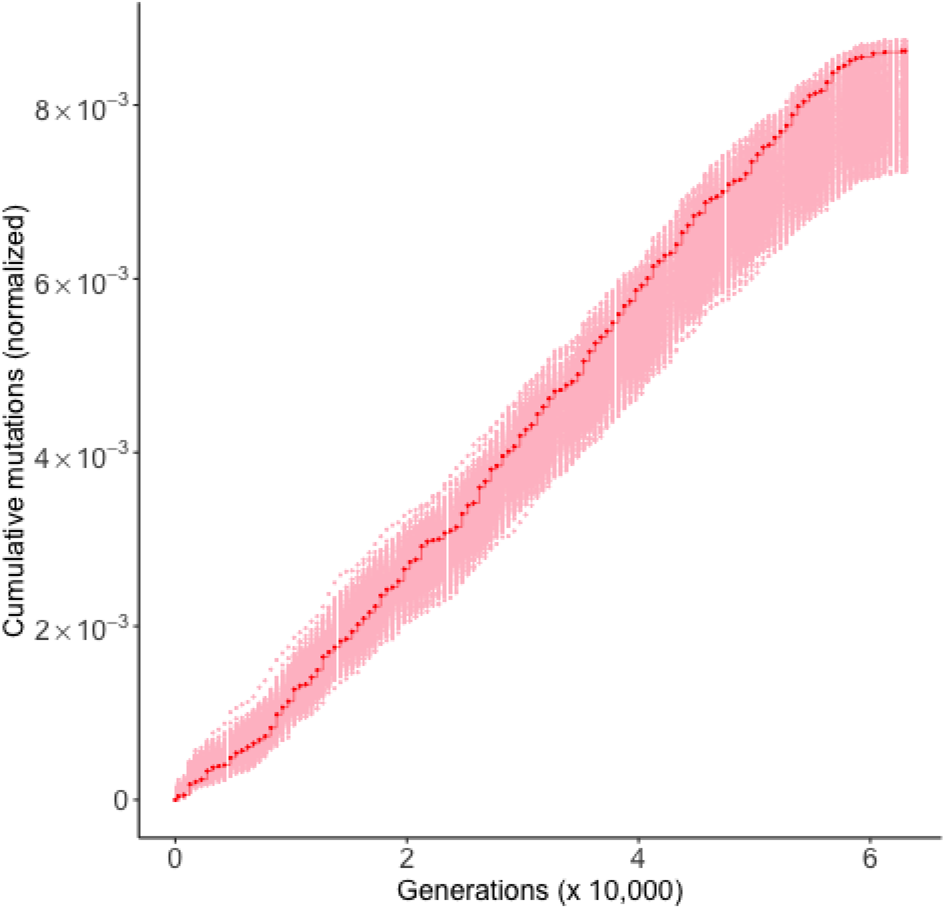

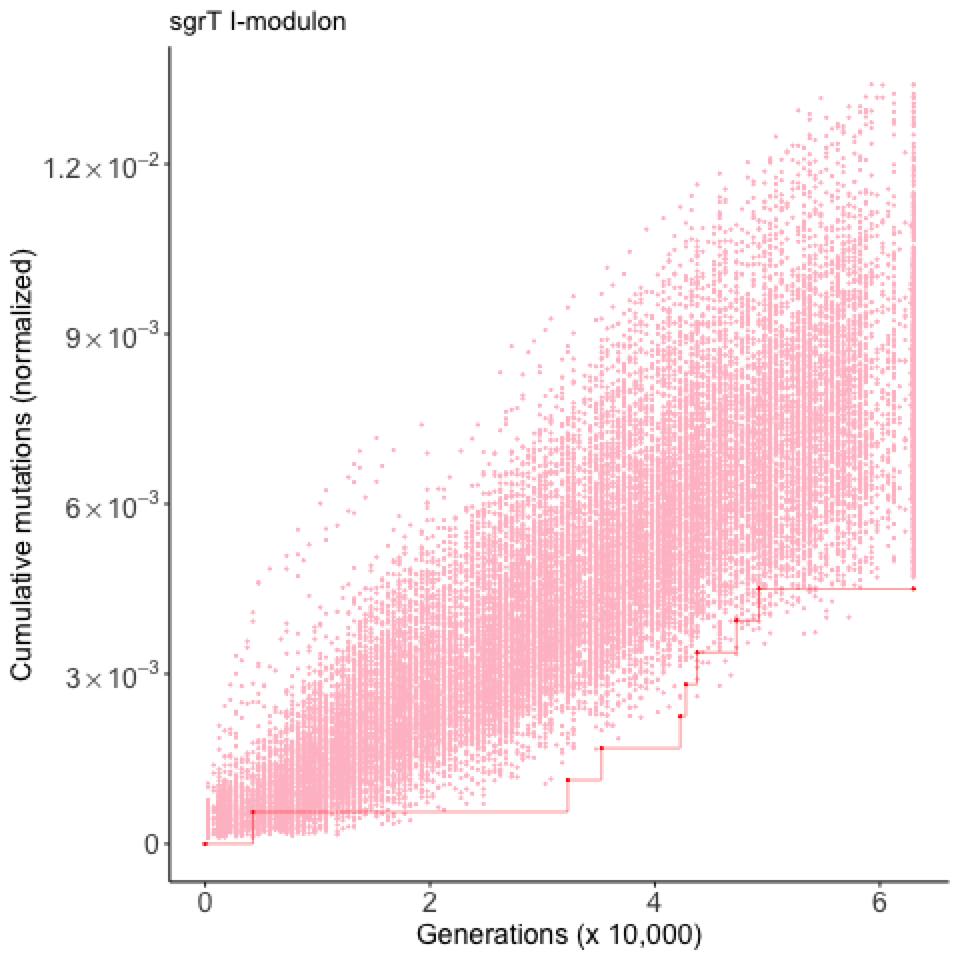
sgrT I-modulon
Cumulative mutations (normalized)
$1.2\times10^{-2}$
$9\times10^{-3}$
$6\times10^{-3}$
$3\times10^{-3}$
0
0
2
4
6
Generations (x 10,000)

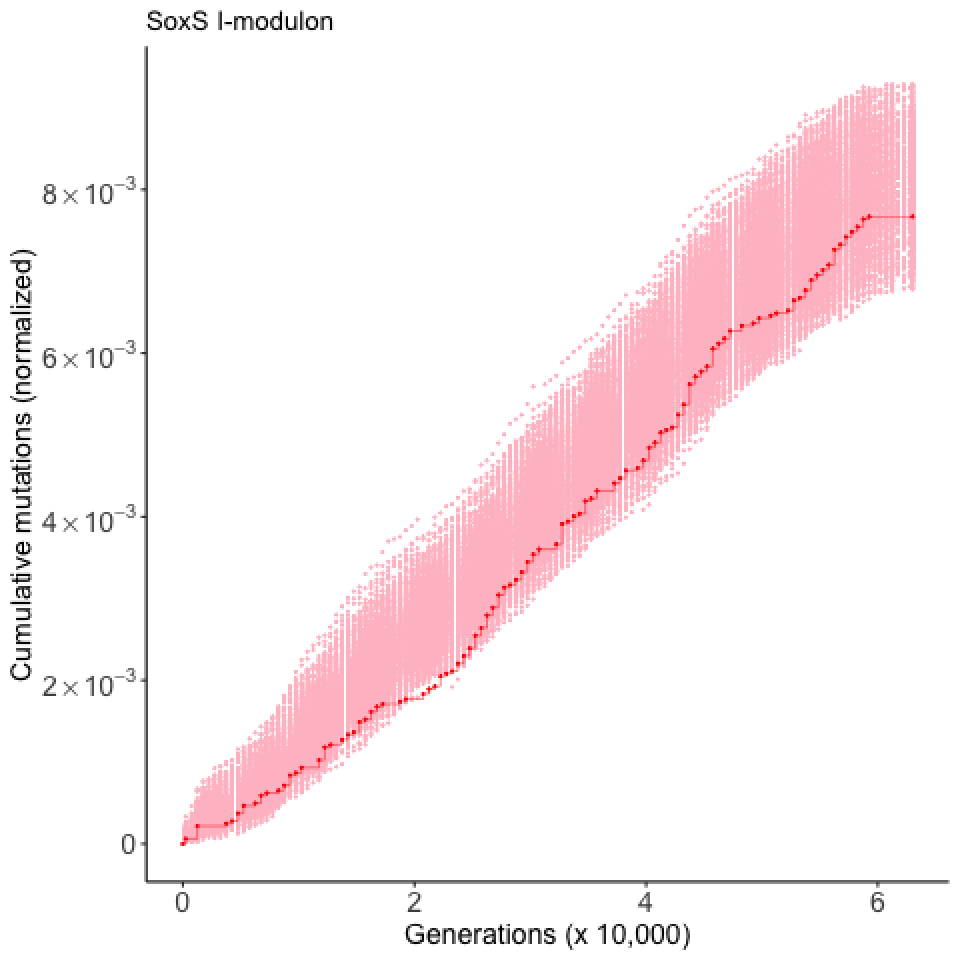

SoxS I-modulon
Cumulative mutations (normalized)
8×10⁻³
6×10⁻³
4×10⁻³
2×10⁻³
0
0
2
4
6
Generations (x 10,000)

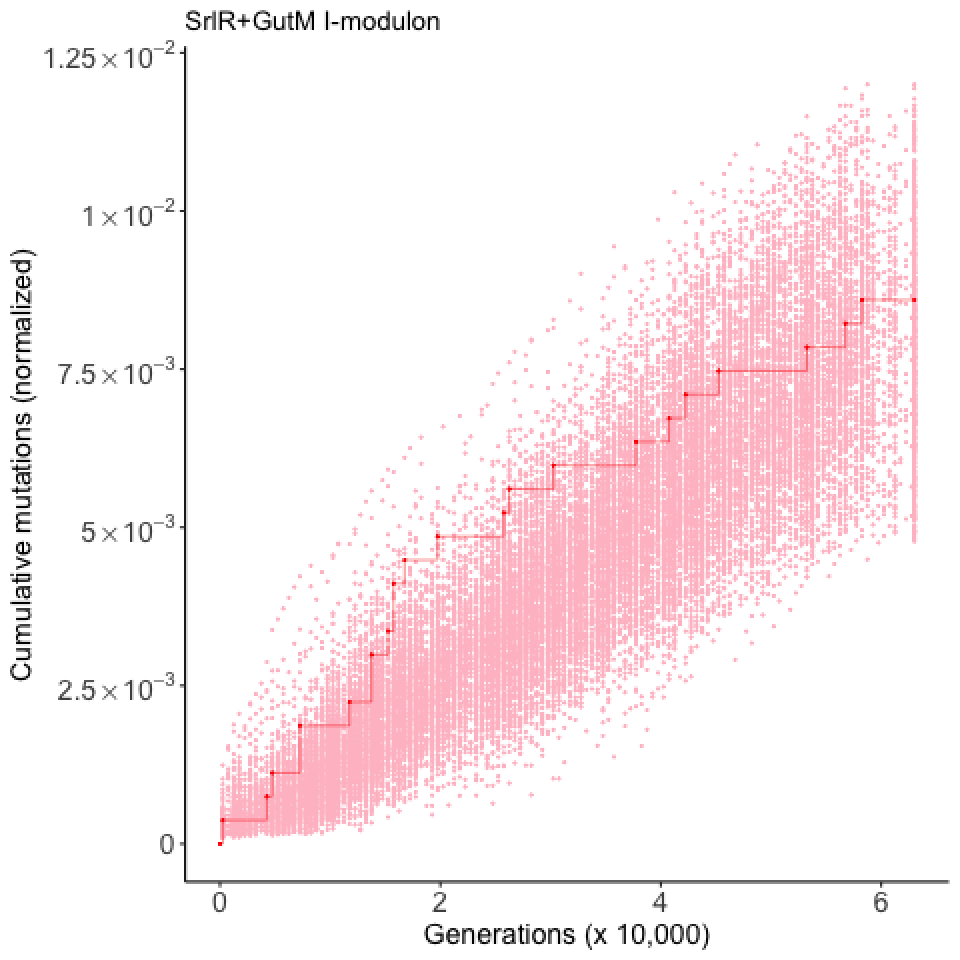

SrlR+GutM I-modulon
Cumulative mutations (normalized)
1.25×10⁻²
1×10⁻²
7.5×10⁻³
5×10⁻³
2.5×10⁻³
0
0
2
4
6
Generations (x 10,000)

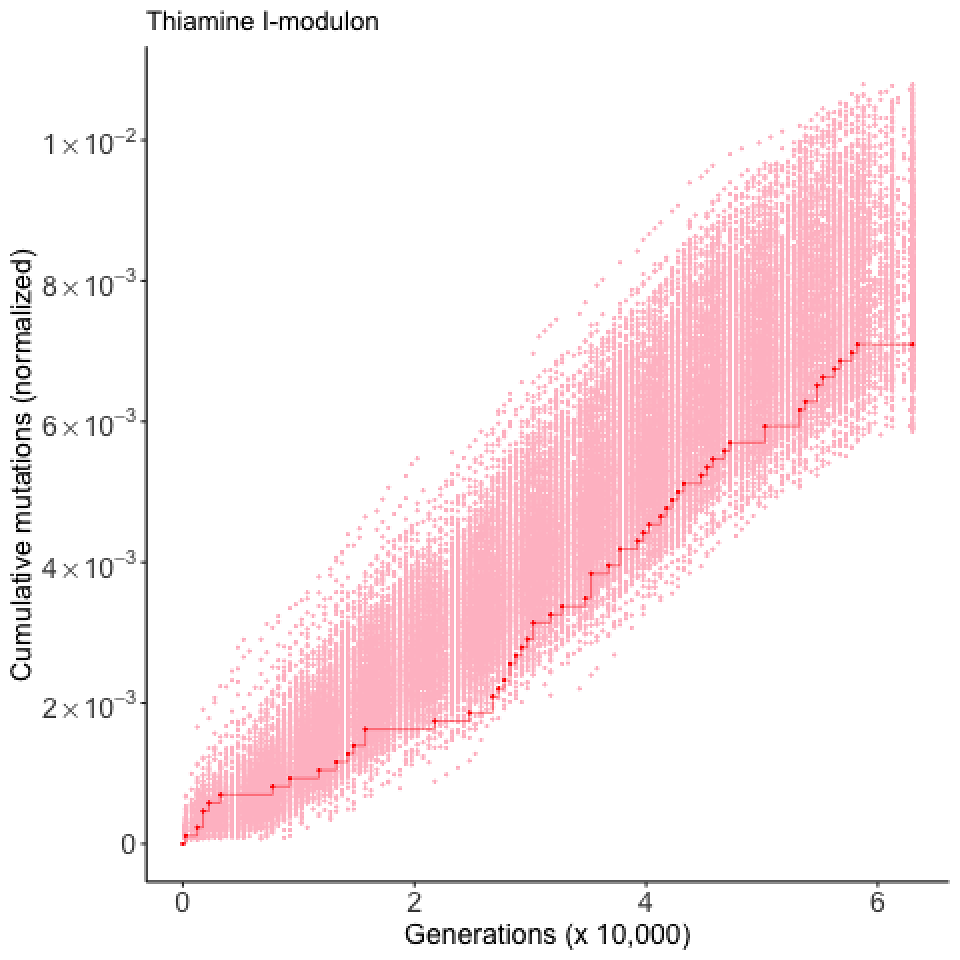

Thiamine I-modulon
Cumulative mutations (normalized)
1×10⁻²
8×10⁻³
6×10⁻³
4×10⁻³
2×10⁻³
0
0
2
4
6
Generations (x 10,000)

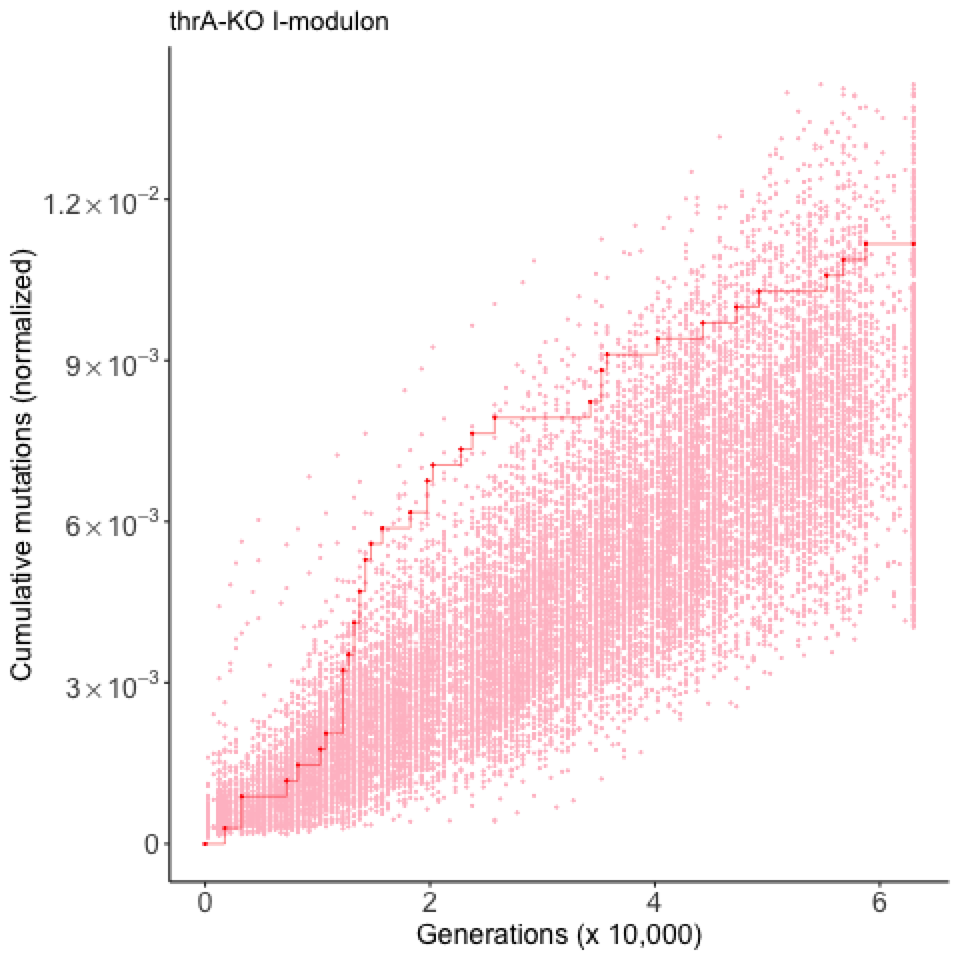

thrA-KO I-modulon
Cumulative mutations (normalized)
$1.2 \times 10^{-2}$
$9 \times 10^{-3}$
$6 \times 10^{-3}$
$3 \times 10^{-3}$
0
0
2
4
6
Generations (x 10,000)

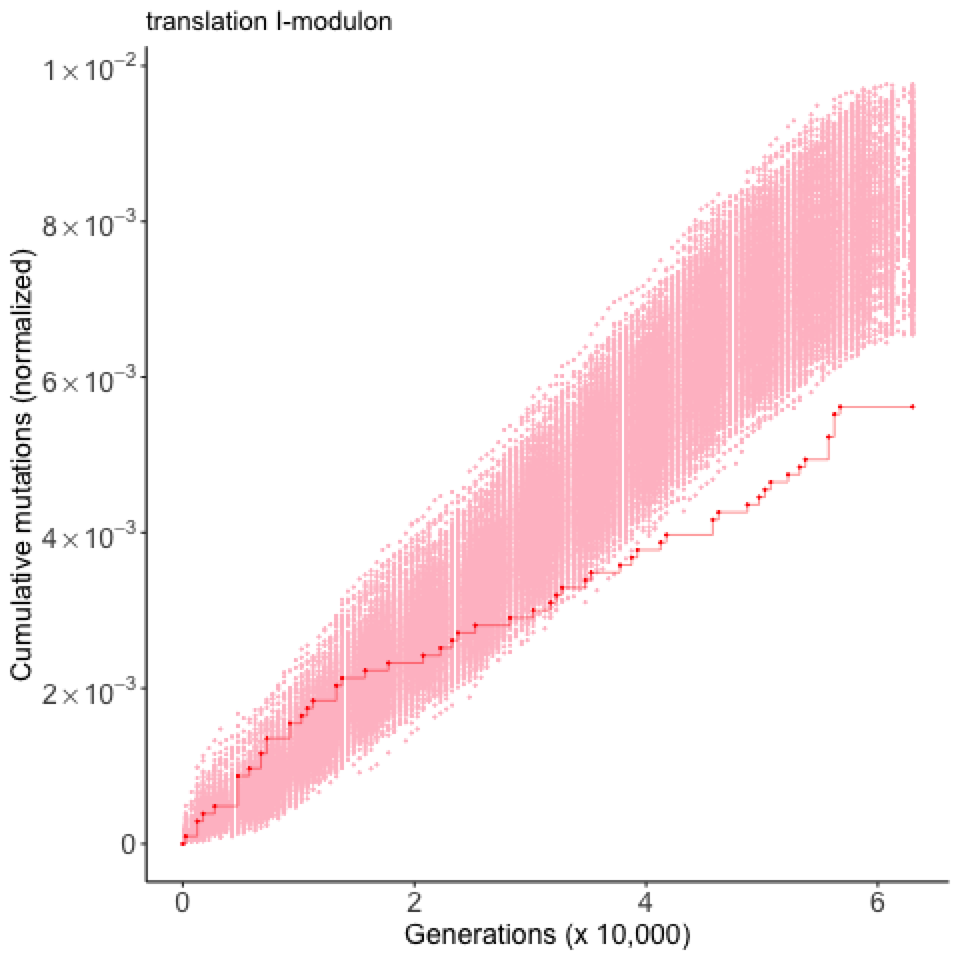

translation I-modulon
Cumulative mutations (normalized)
$1\times10^{-2}$
$8\times10^{-3}$
$6\times10^{-3}$
$4\times10^{-3}$
$2\times10^{-3}$
0
0
2
4
6
Generations (x 10,000)

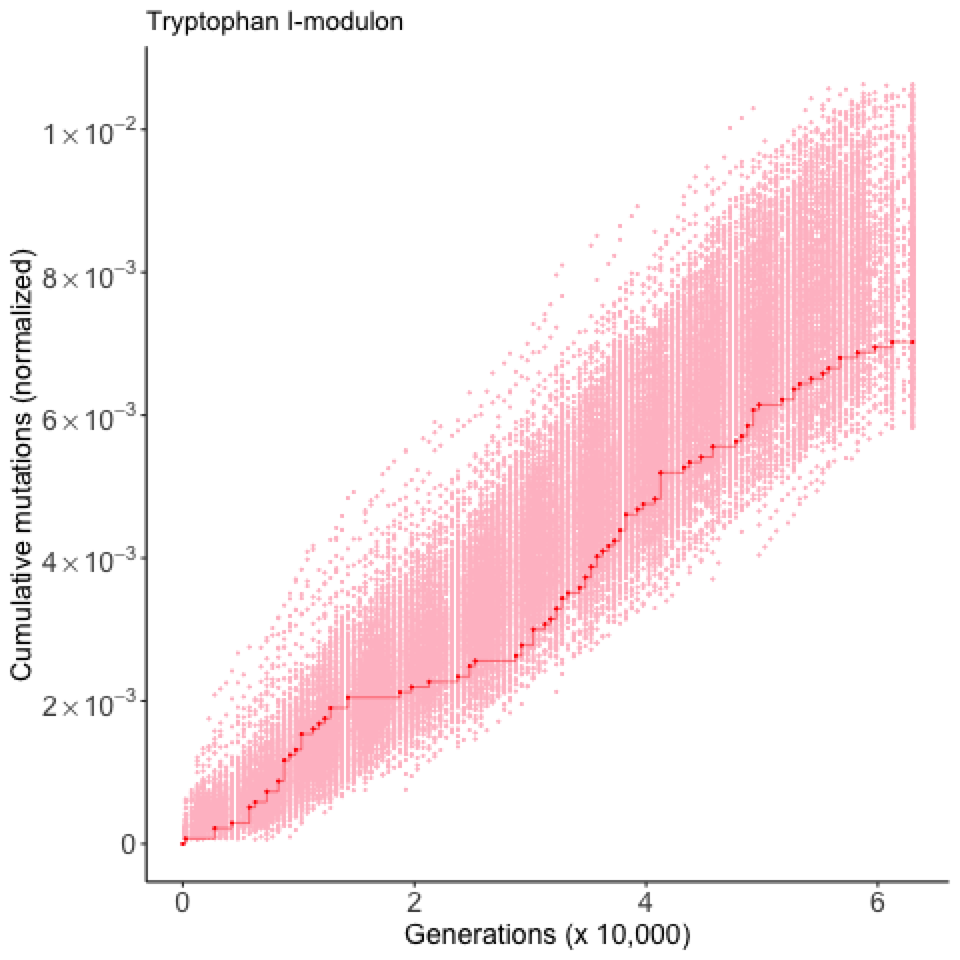

Tryptophan I-modulon
Cumulative mutations (normalized)
1×10⁻²
8×10⁻³
6×10⁻³
4×10⁻³
2×10⁻³
0
0
2
4
6
Generations (x 10,000)

uncharacterized-1 I-modulon

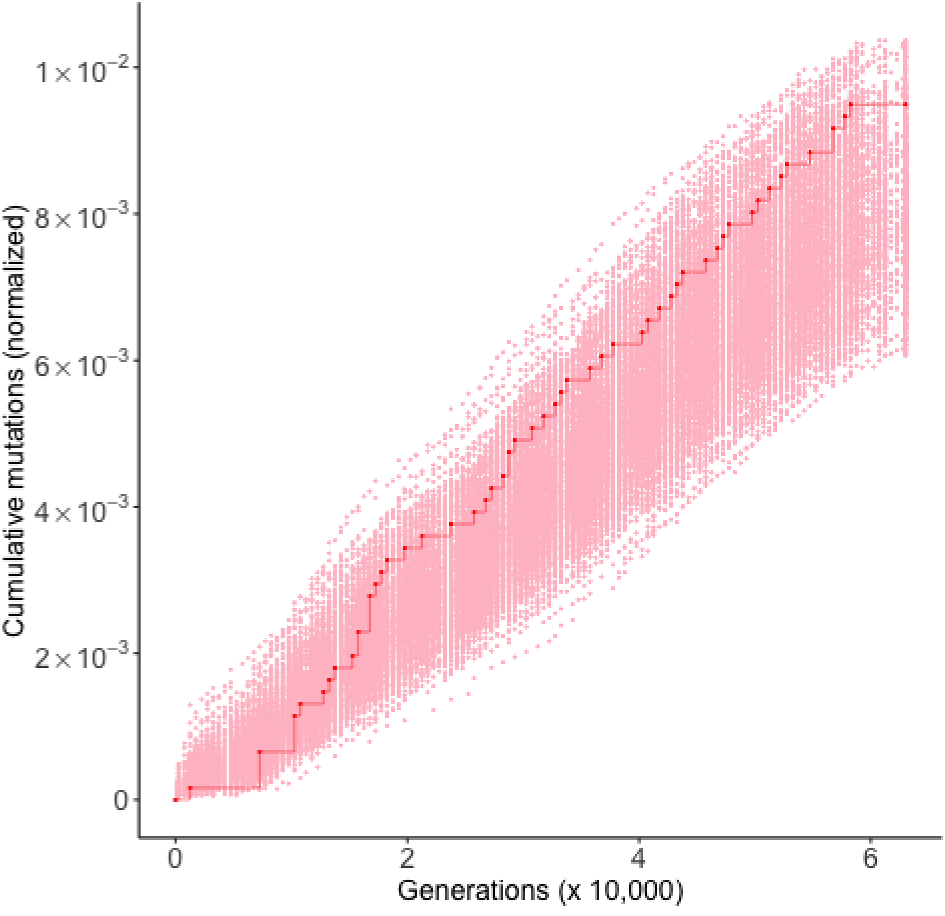

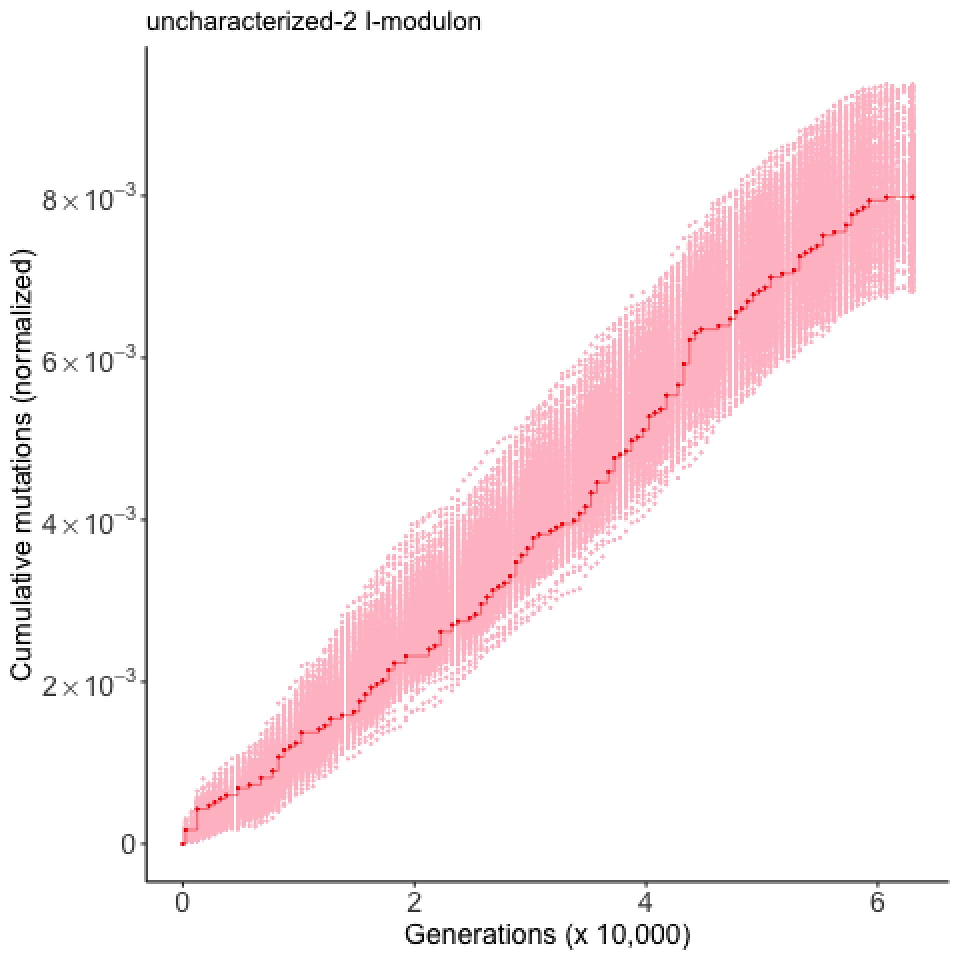

uncharacterized-2 I-modulon
Cumulative mutations (normalized)
8×10⁻³
6×10⁻³
4×10⁻³
2×10⁻³
0
0
2
4
6
Generations (x 10,000)

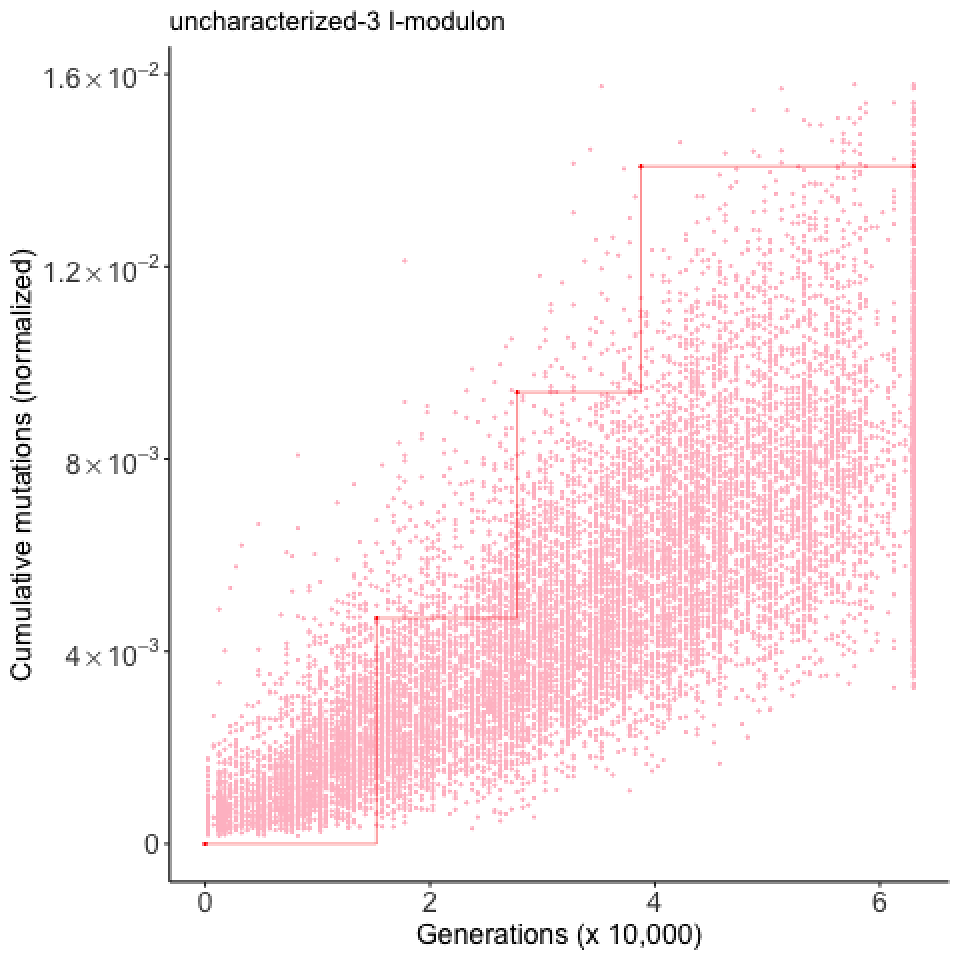

uncharacterized-3 I-modulon
Cumulative mutations (normalized)
1.6×10^-2
1.2×10^-2
8×10^-3
4×10^-3
0
0
2
4
6
Generations (x 10,000)

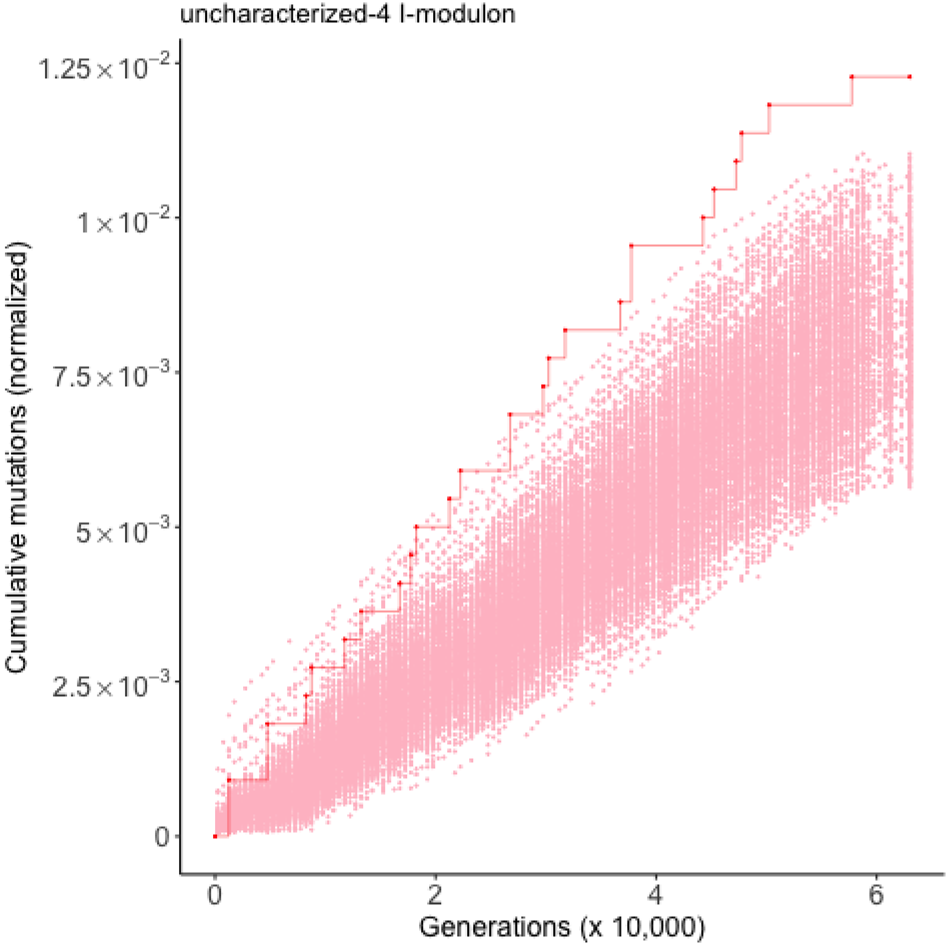

uncharacterized-4 I-modulon
Cumulative mutations (normalized)
$1.25\times10^{-2}$
$1\times10^{-2}$
$7.5\times10^{-3}$
$5\times10^{-3}$
$2.5\times10^{-3}$
0
0
2
4
6
Generations (x 10,000)

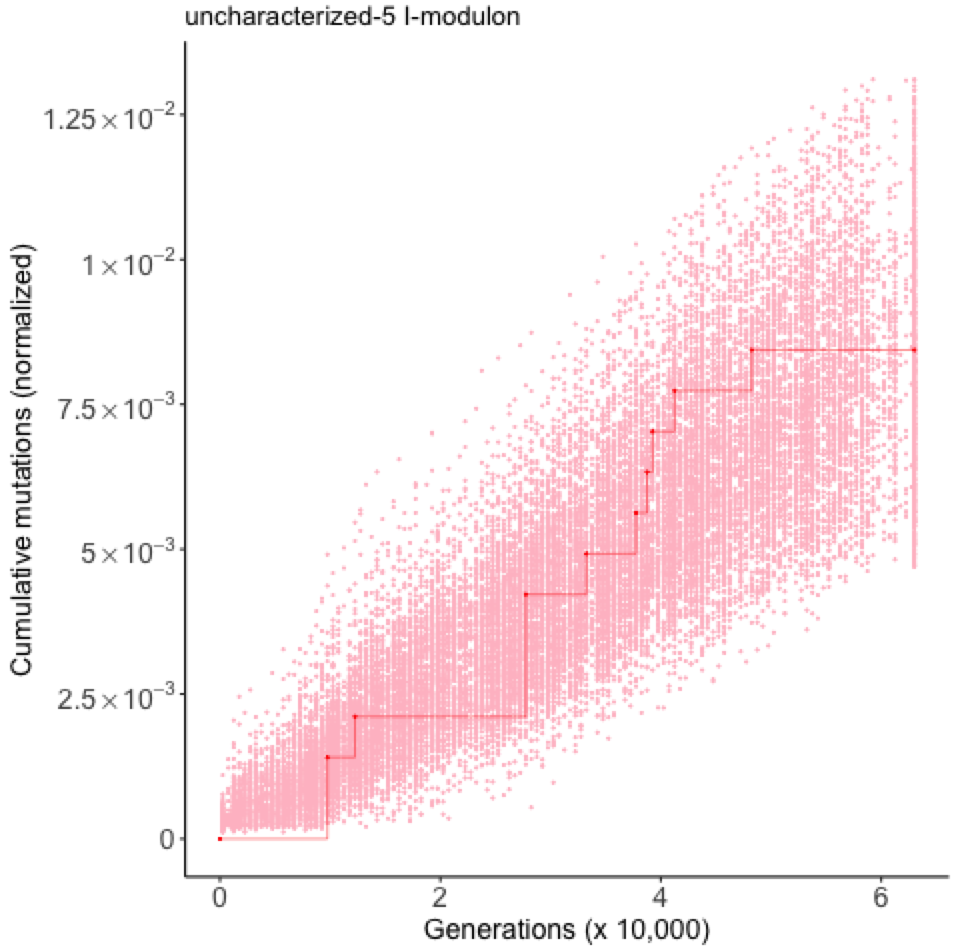

uncharacterized-5 I-modulon
Cumulative mutations (normalized)
$1.25\times10^{-2}$
$1\times10^{-2}$
$7.5\times10^{-3}$
$5\times10^{-3}$
$2.5\times10^{-3}$
0
0
2
4
6
Generations (x 10,000)

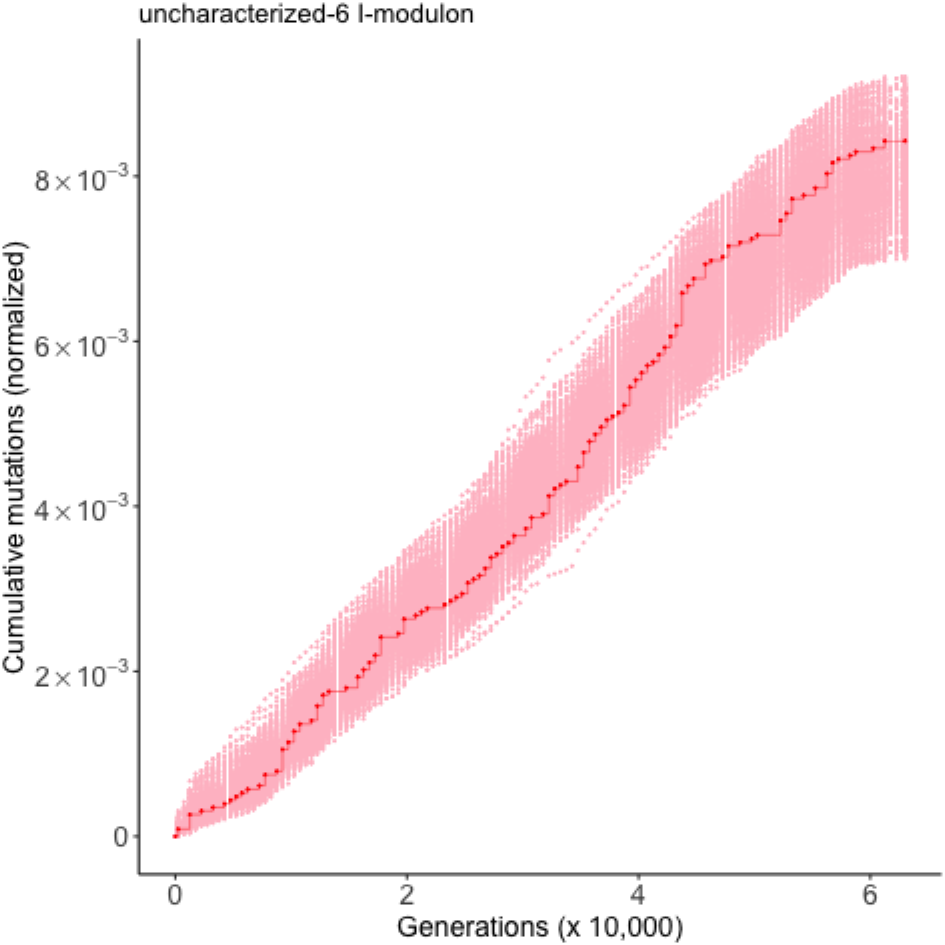

uncharacterized-6 I-modulon
Cumulative mutations (normalized)
0
$2\times10^{-3}$
$4\times10^{-3}$
$6\times10^{-3}$
$8\times10^{-3}$
0
2
4
6
Generations (x 10,000)

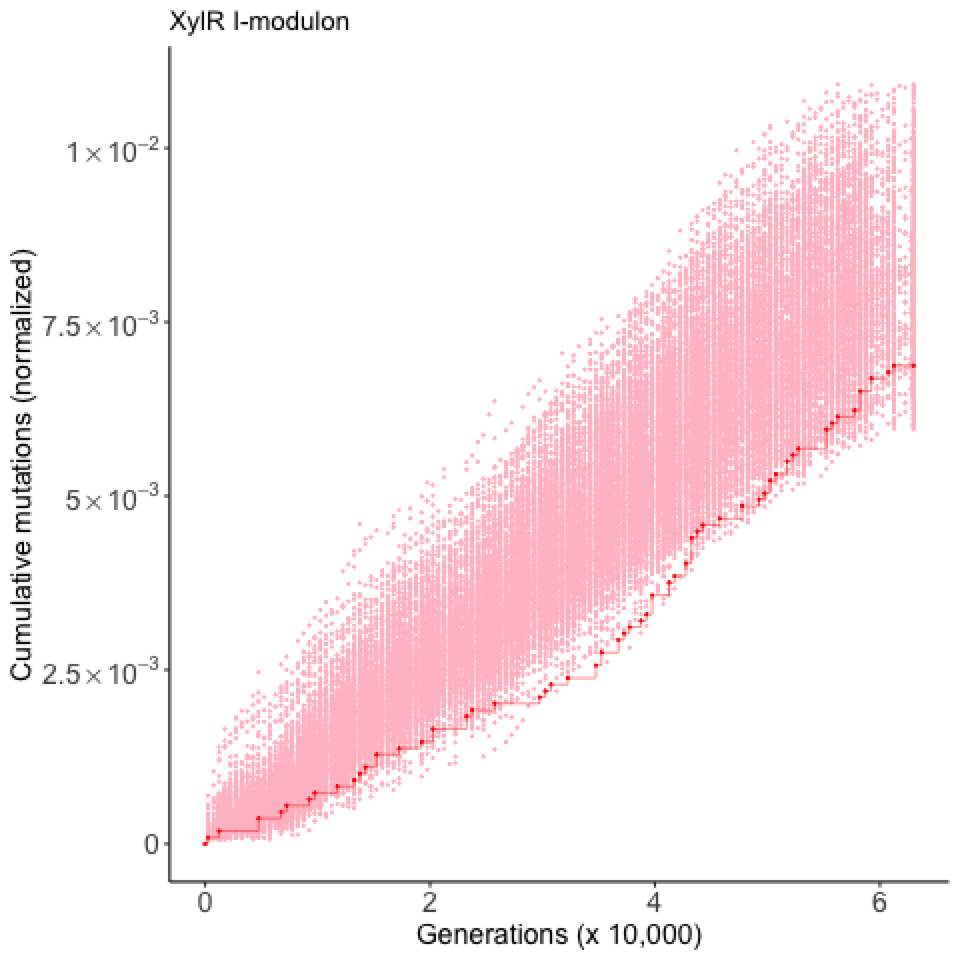
XylR I-modulon
Cumulative mutations (normalized)
1×10^-2
7.5×10^-3
5×10^-3
2.5×10^-3
0
0
2
4
6
Generations (x 10,000)

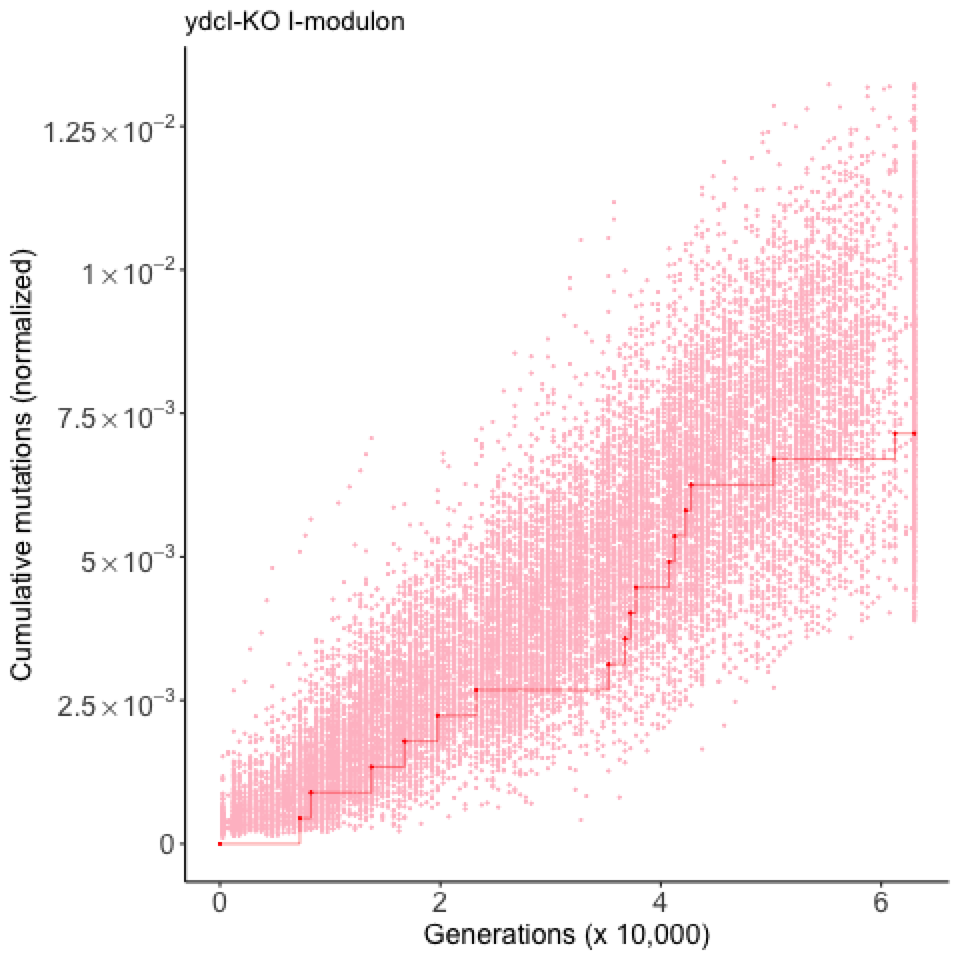

ydcI-KO I-modulon
Cumulative mutations (normalized)
$1.25 \times 10^{-2}$
$1 \times 10^{-2}$
$7.5 \times 10^{-3}$
$5 \times 10^{-3}$
$2.5 \times 10^{-3}$
0
0
2
4
6
Generations (x 10,000)

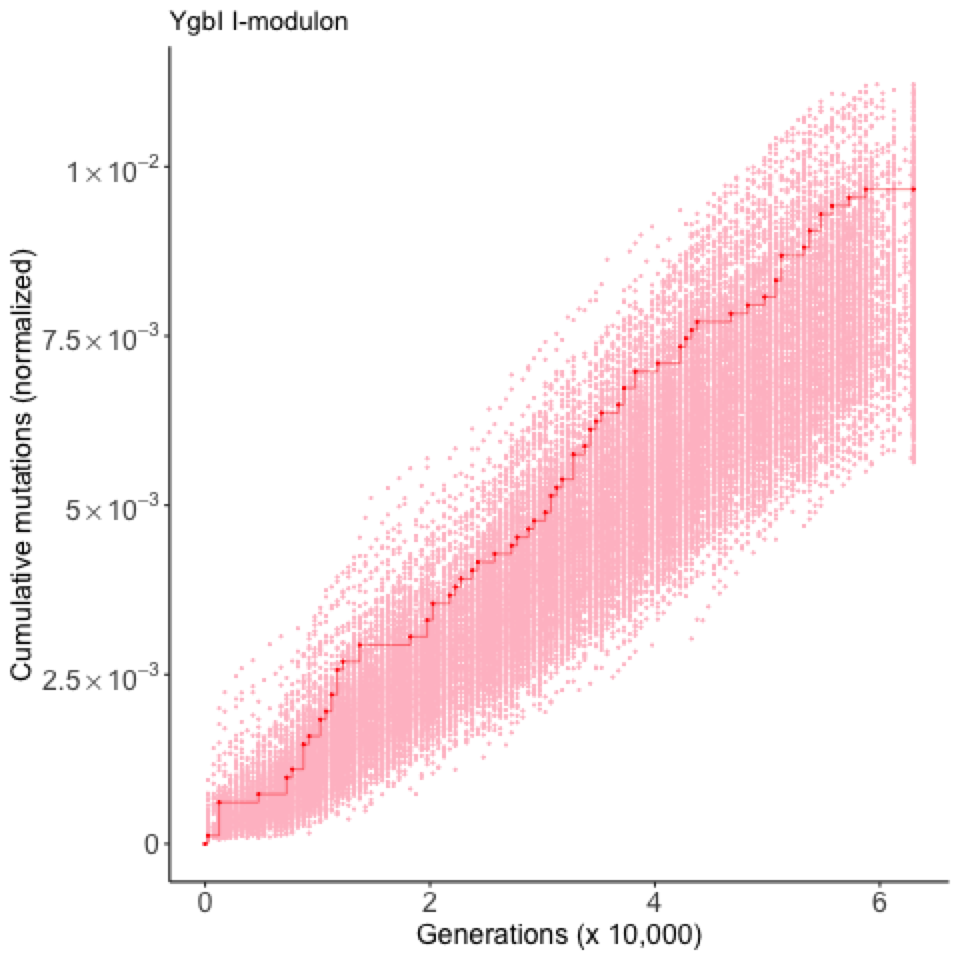

YgbI I-modulon
Cumulative mutations (normalized)
$1\times10^{-2}$
$7.5\times10^{-3}$
$5\times10^{-3}$
$2.5\times10^{-3}$
0
0
2
4
6
Generations (x 10,000)

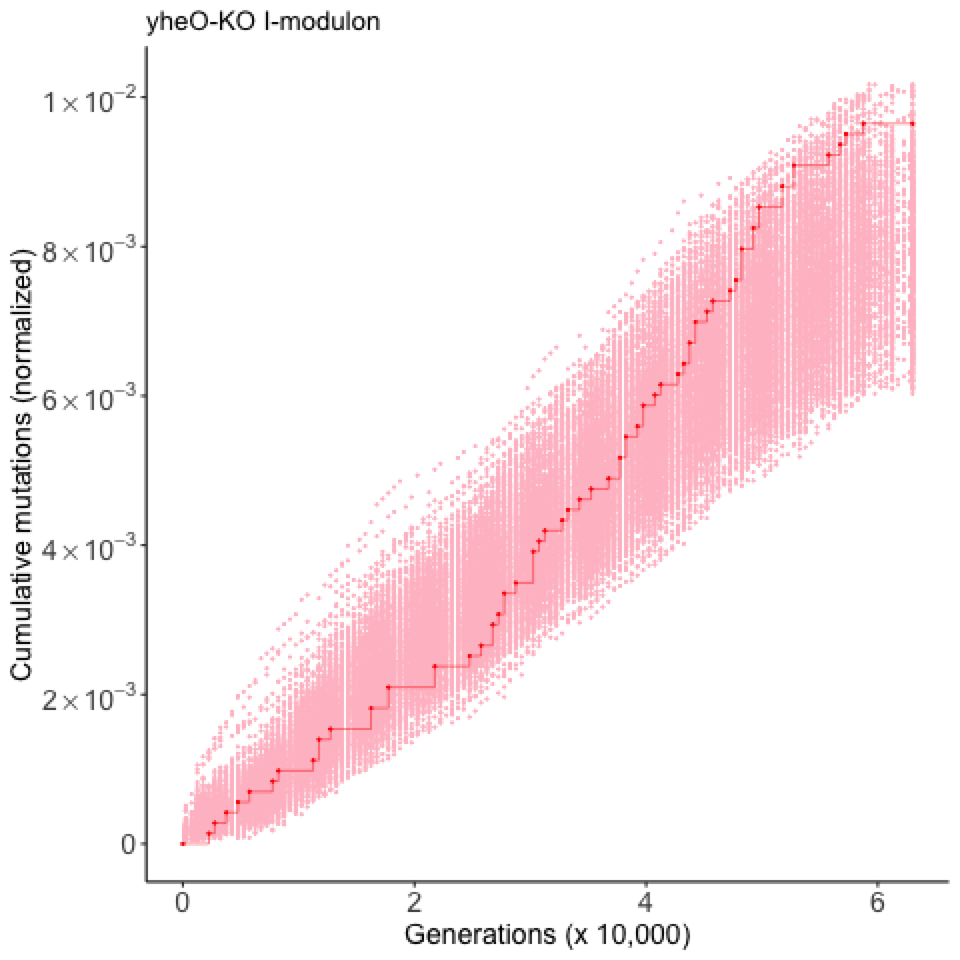

yheO-KO I-modulon
Cumulative mutations (normalized)
1×10^−2
8×10^−3
6×10^−3
4×10^−3
2×10^−3
0
0
2
4
6
Generations (x 10,000)

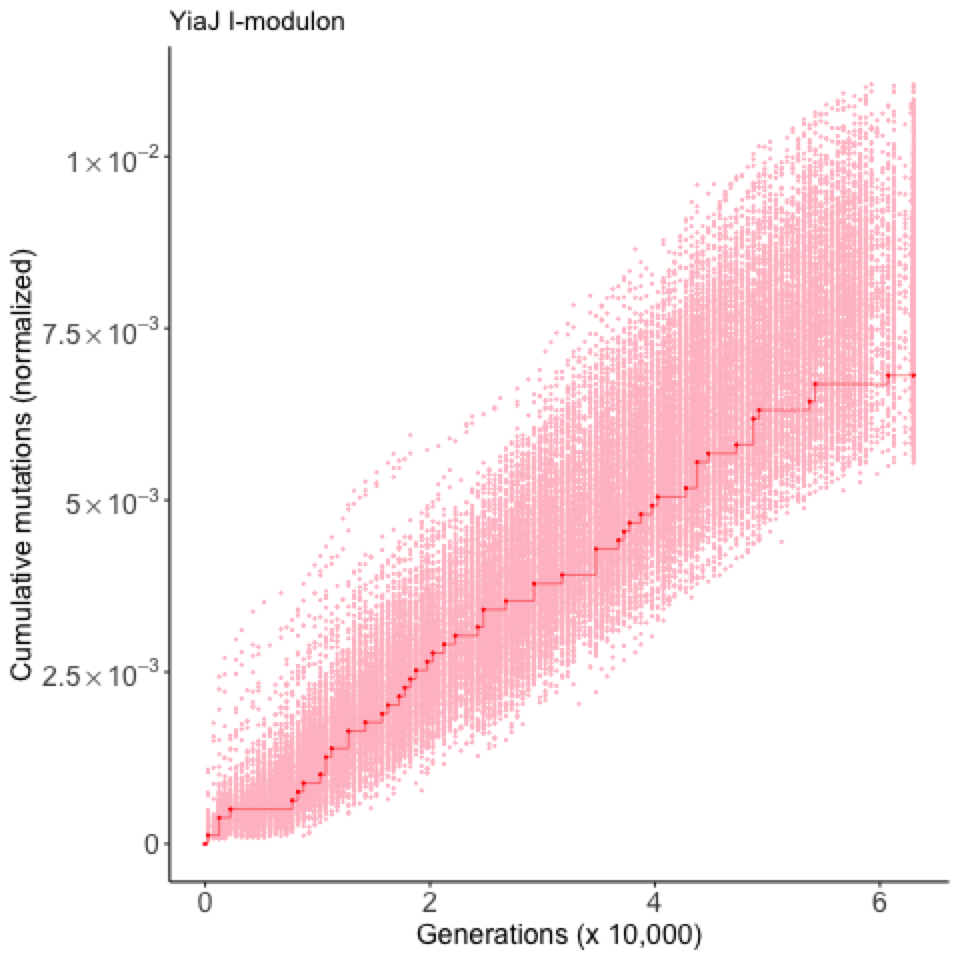

YiaJ I-modulon
Cumulative mutations (normalized)
$1\times10^{-2}$
$7.5\times10^{-3}$
$5\times10^{-3}$
$2.5\times10^{-3}$
0
0
2
4
6
Generations (x 10,000)

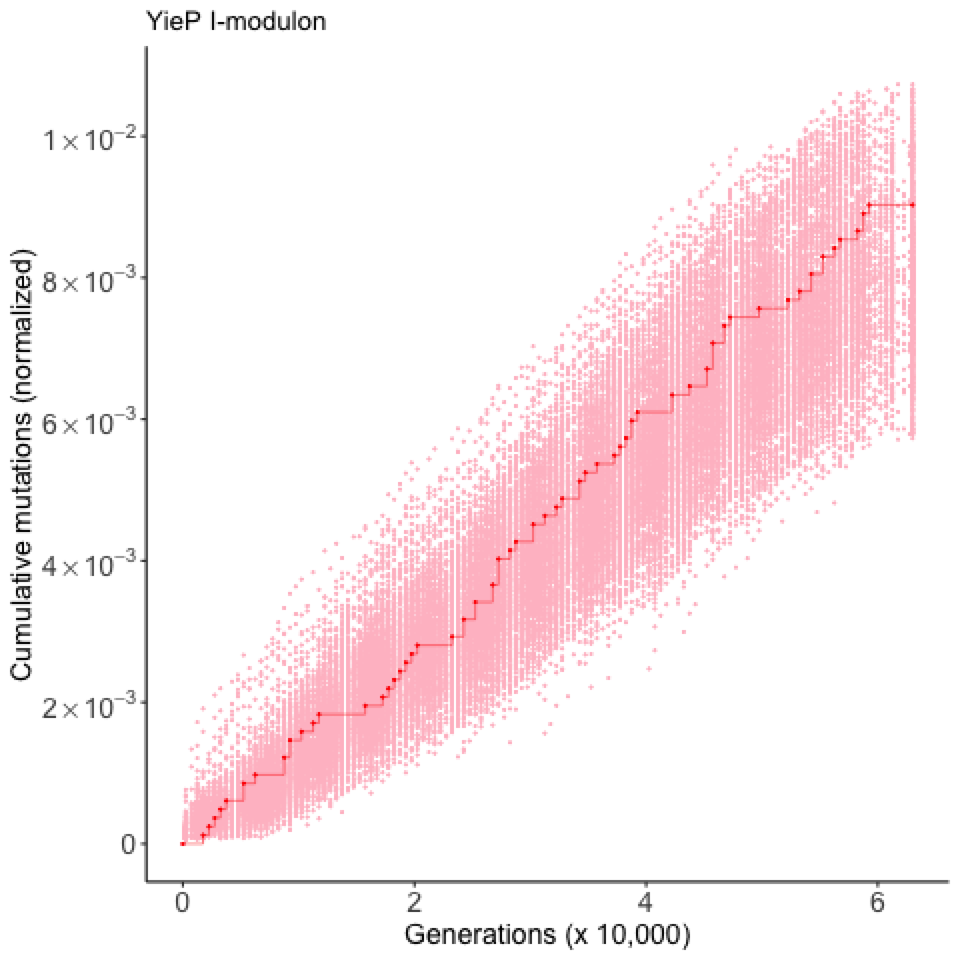

YieP I-modulon
Cumulative mutations (normalized)
1×10^-2
8×10^-3
6×10^-3
4×10^-3
2×10^-3
0
0
2
4
6
Generations (x 10,000)

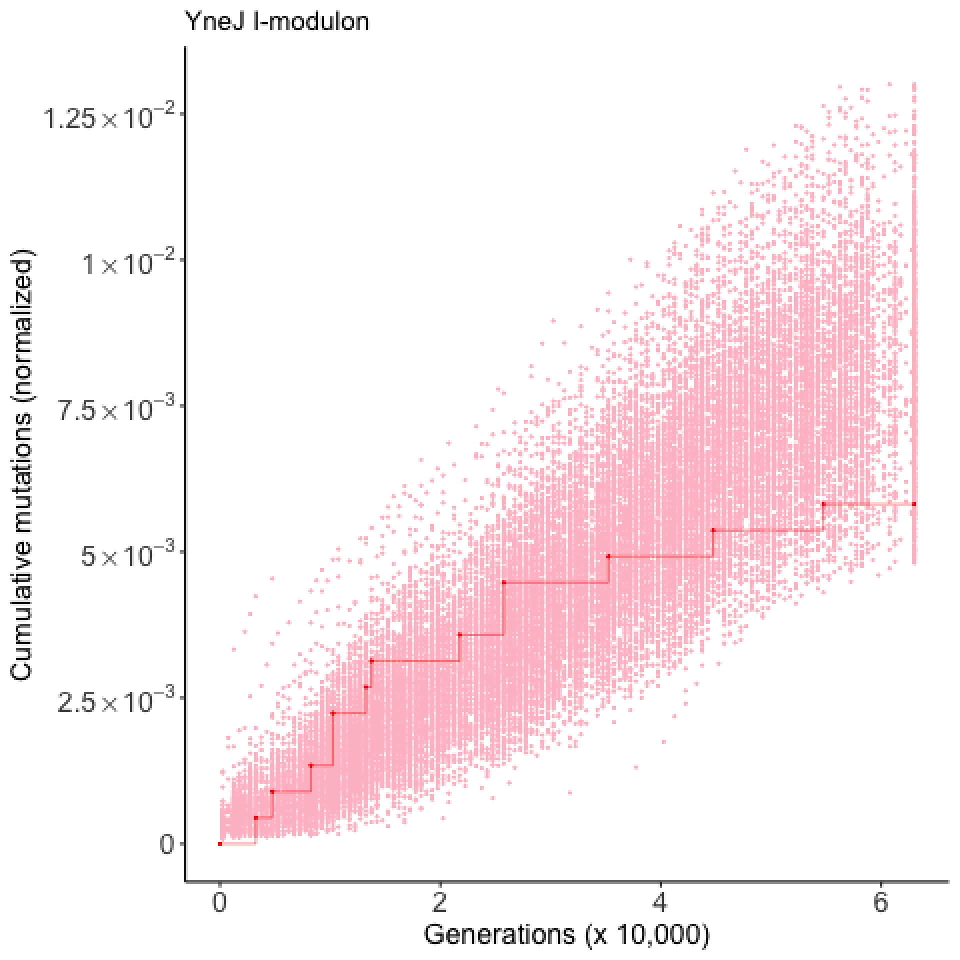

YneJ I-modulon
Cumulative mutations (normalized)
1.25×10^-2
1×10^-2
7.5×10^-3
5×10^-3
2.5×10^-3
0
0
2
4
6
Generations (x 10,000)

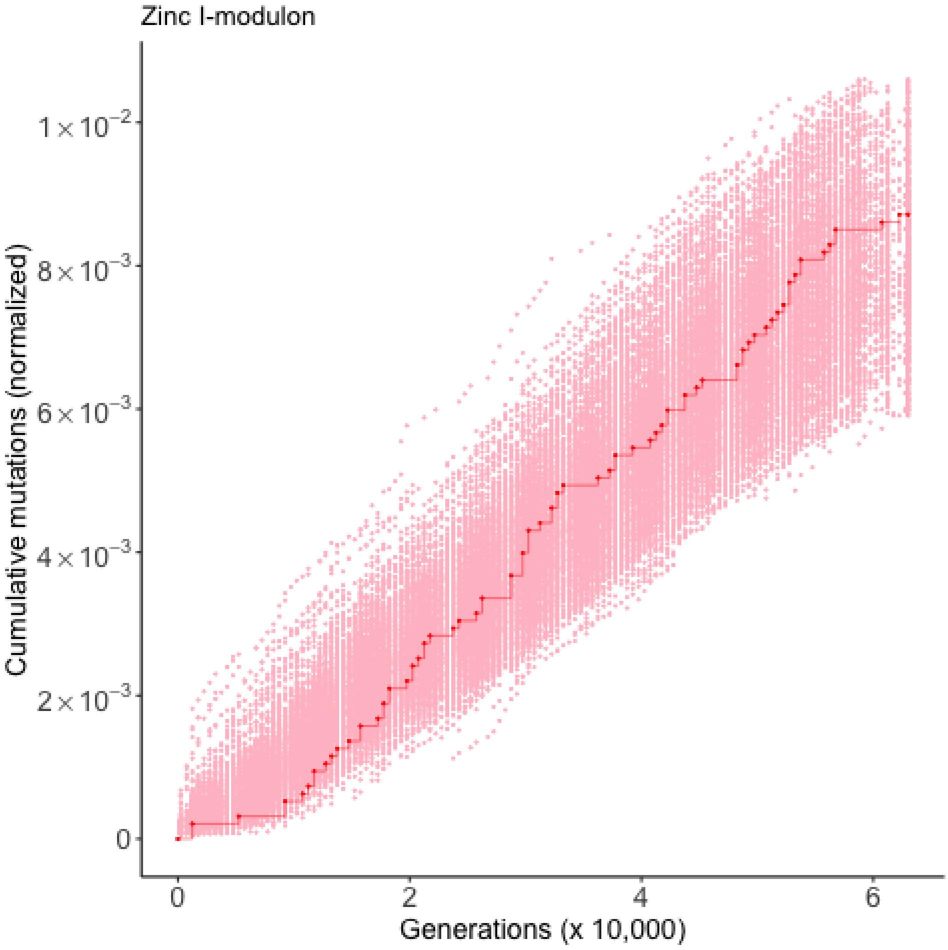
Zinc I-modulon
Cumulative mutations (normalized)
$1\times10^{-2}$
$8\times10^{-3}$
$6\times10^{-3}$
$4\times10^{-3}$
$2\times10^{-3}$
0
0
2
4
6
Generations (x 10,000)
