## Supplementary file S2 for "Discovery of positive and purifying selection in metagenomic time series of hypermutator microbial populations"

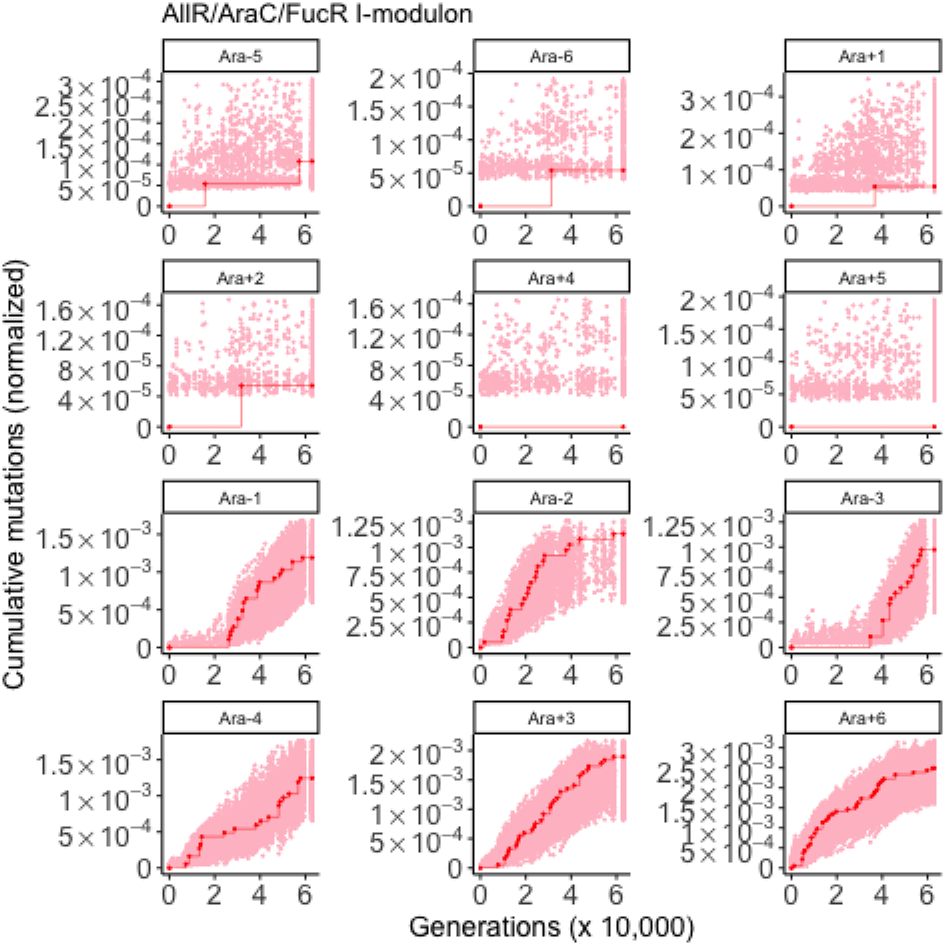
AllR/AraC/FucR I-modulon
Ara-5
Ara-6
Ara+1
Ara+2
Ara+4
Ara+5
Ara-1
Ara-2
Ara-3
Ara-4
Ara+3
Ara+6
Cumulative mutations (normalized)
Generations (x 10,000)

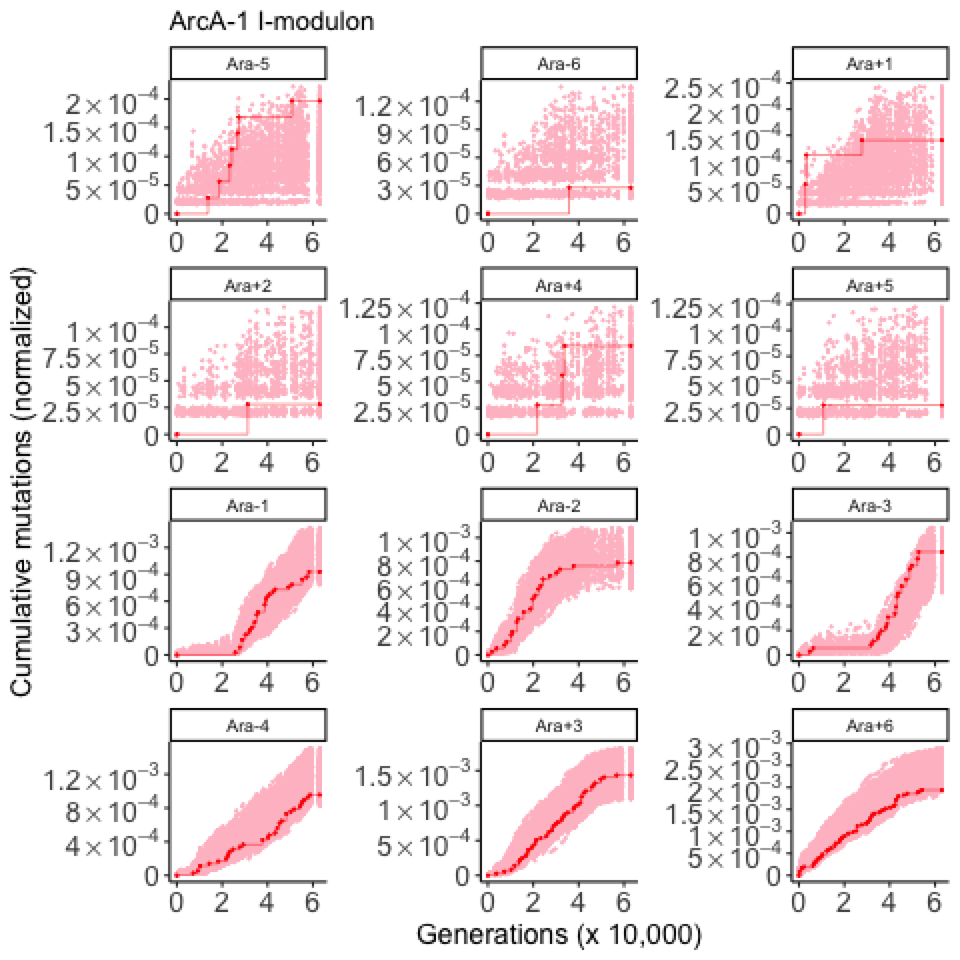
ArcA-1 I-modulon
Ara-5
Ara-6
Ara+1
Ara+2
Ara+4
Ara+5
Ara-1
Ara-2
Ara-3
Ara-4
Ara+3
Ara+6
Cumulative mutations (normalized)
Generations (x 10,000)

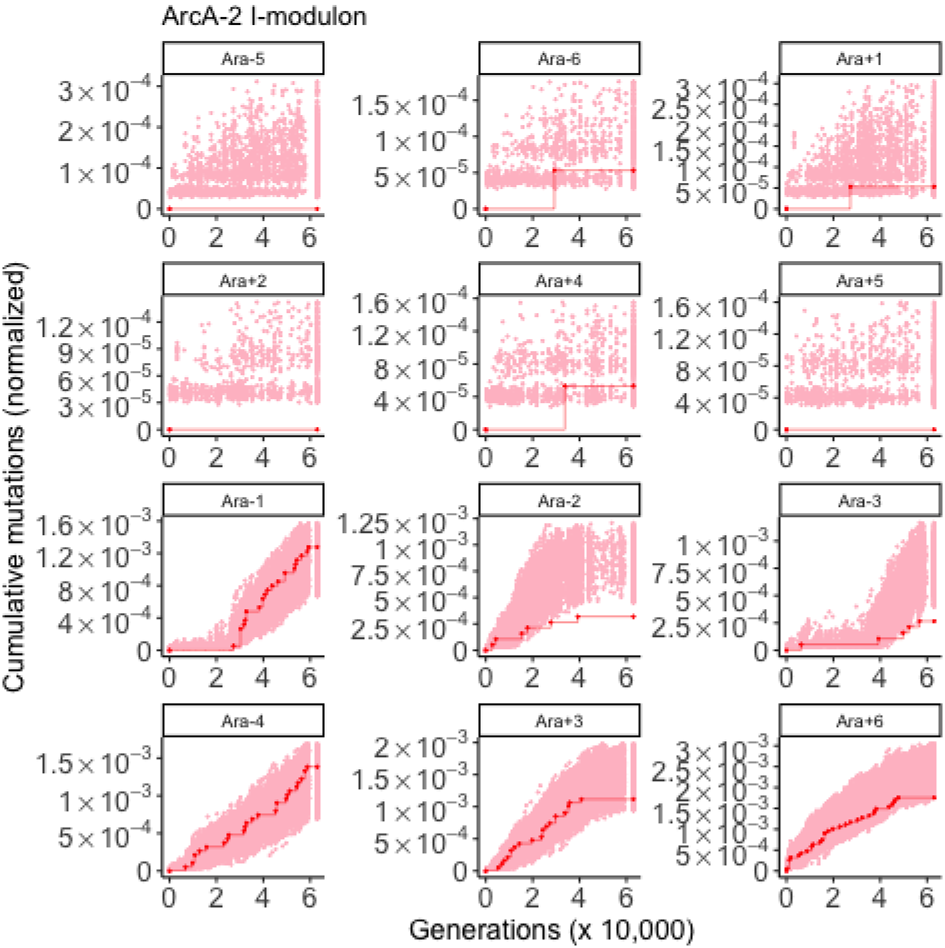
ArcA-2 I-modulon
Ara-5
Ara-6
Ara+1
Ara+2
Ara+4
Ara+5
Ara-1
Ara-2
Ara-3
Ara-4
Ara+3
Ara+6
Cumulative mutations (normalized)
Generations (x 10,000)

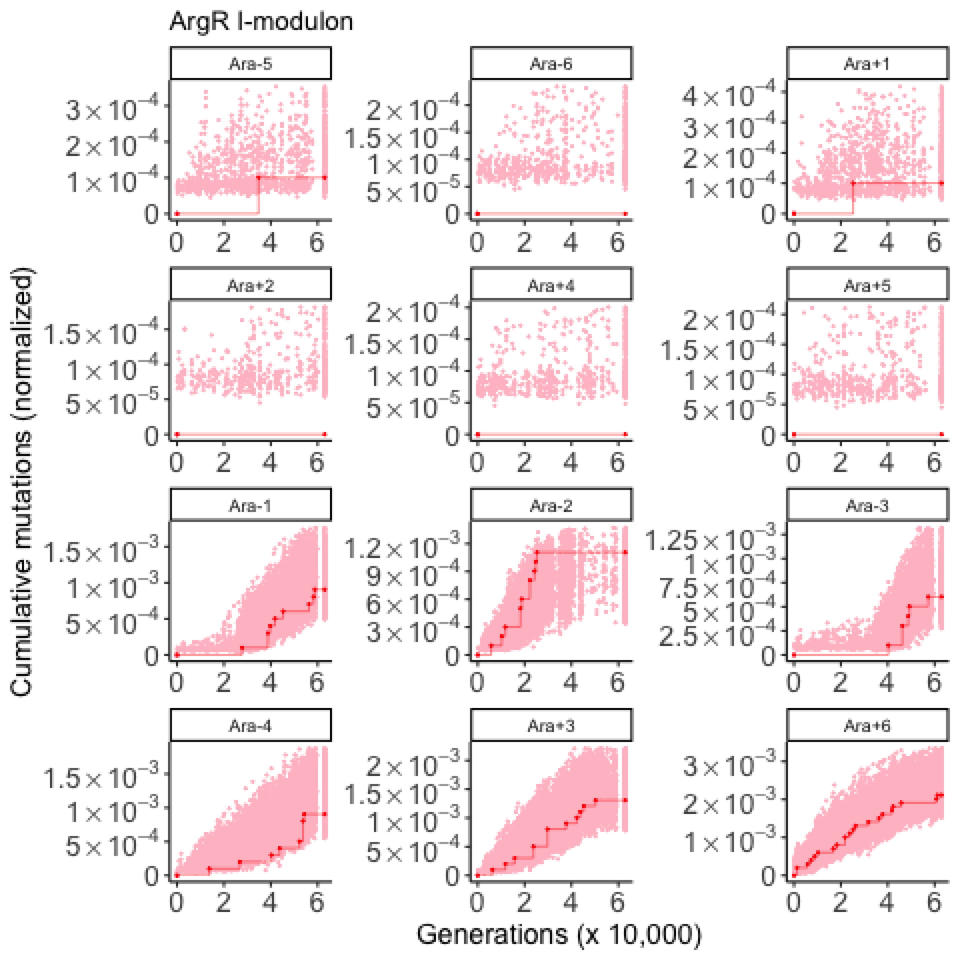

ArgR I-modulon
Ara-5
Ara-6
Ara+1
Ara+2
Ara+4
Ara+5
Ara-1
Ara-2
Ara-3
Ara-4
Ara+3
Ara+6
Cumulative mutations (normalized)
Generations (x 10,000)

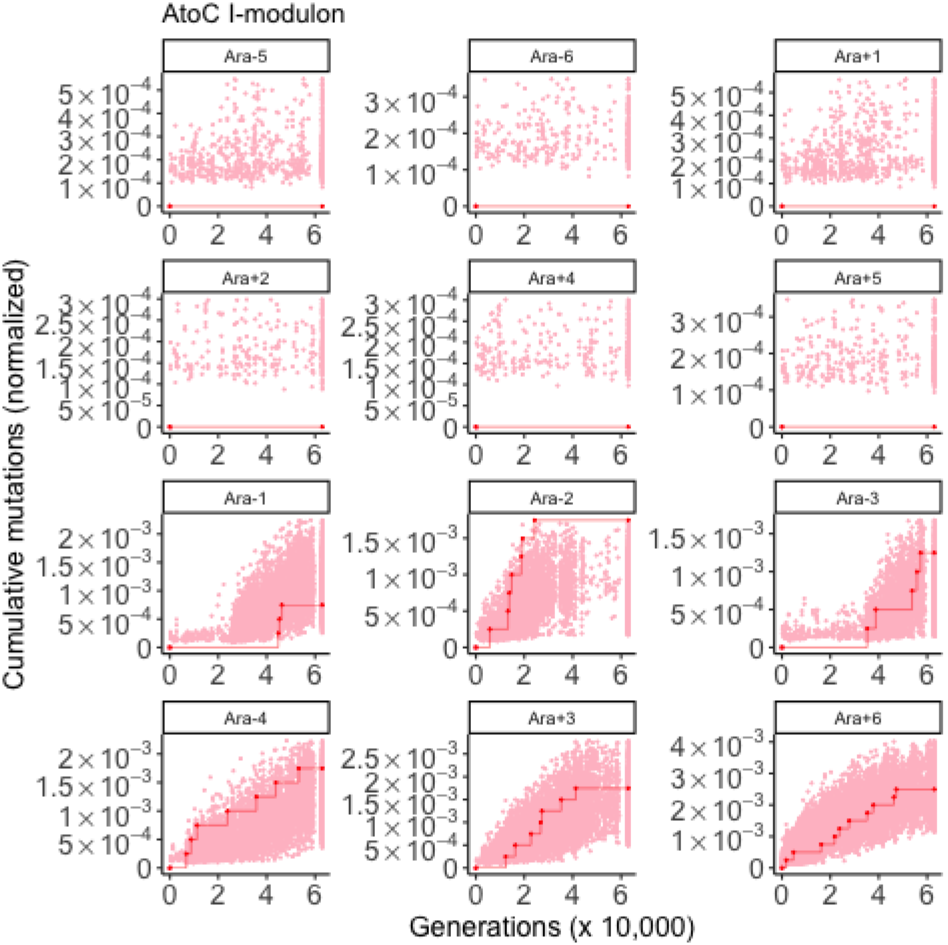
AtoC I-modulon
Ara-5
Ara-6
Ara+1
Ara+2
Ara+4
Ara+5
Ara-1
Ara-2
Ara-3
Ara-4
Ara+3
Ara+6
Cumulative mutations (normalized)
Generations (x 10,000)

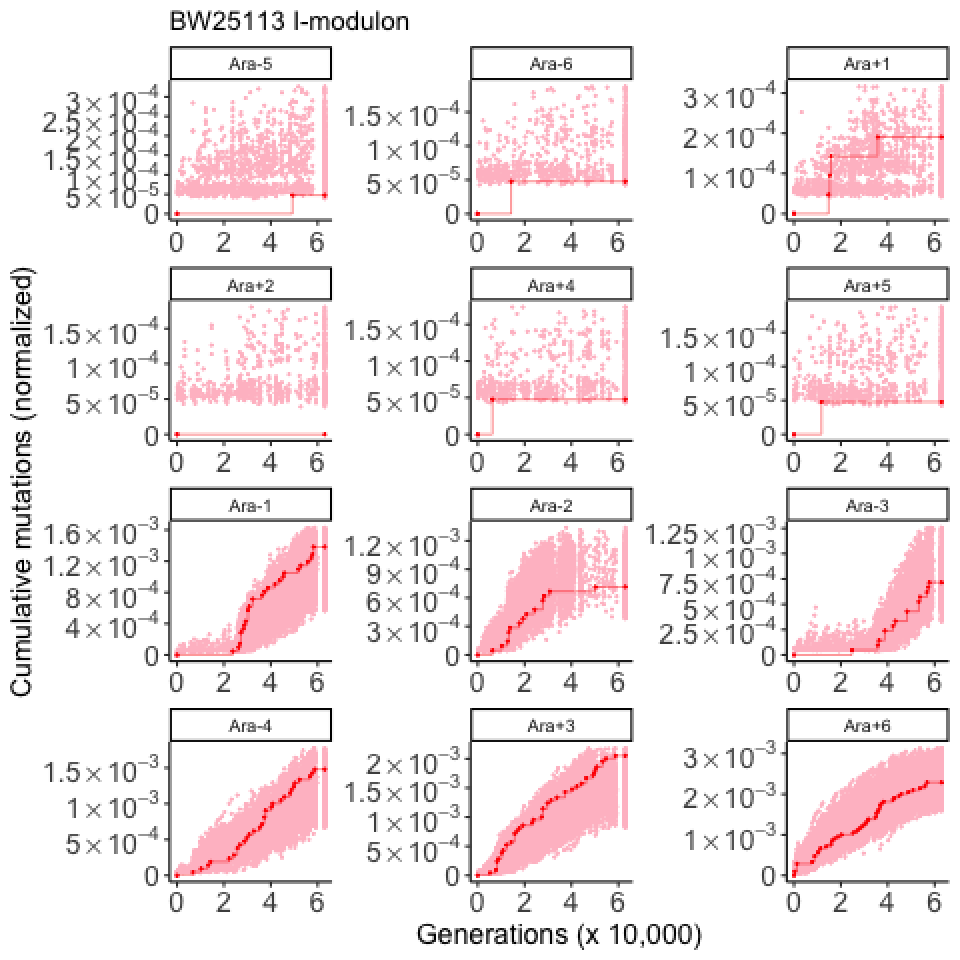
BW25113 I-modulon
Ara-5
Ara-6
Ara+1
Ara+2
Ara+4
Ara+5
Ara-1
Ara-2
Ara-3
Ara-4
Ara+3
Ara+6
Cumulative mutations (normalized)
Generations (x 10,000)

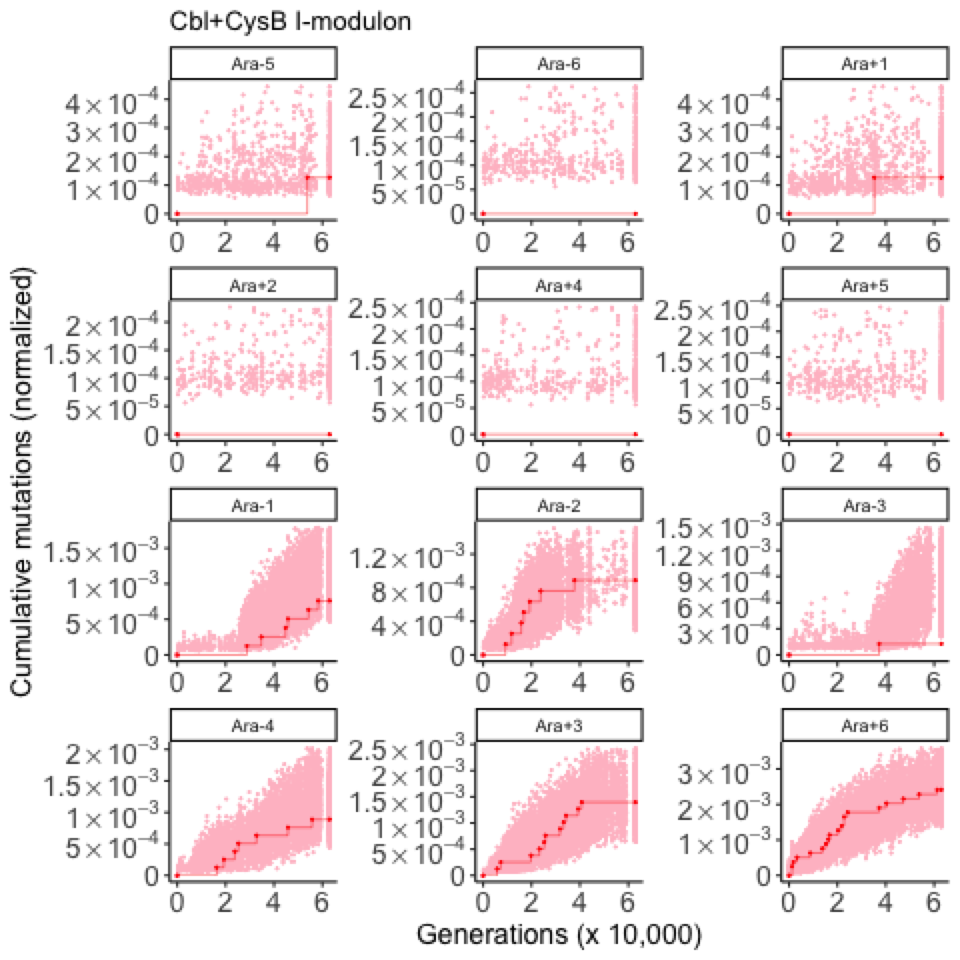

Cbl+CysB I-modulon
Ara-5
Ara-6
Ara+1
Ara+2
Ara+4
Ara+5
Ara-1
Ara-2
Ara-3
Ara-4
Ara+3
Ara+6
Cumulative mutations (normalized)
Generations (x 10,000)

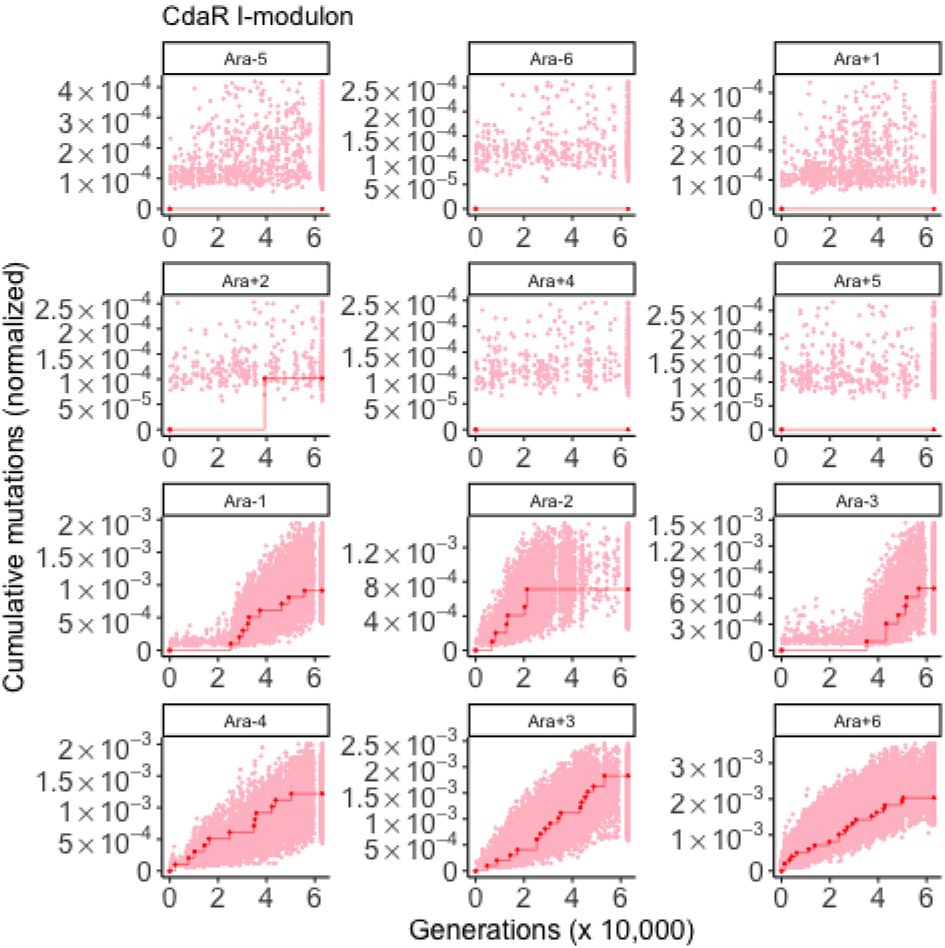
CdaR I-modulon
Ara-5
Ara-6
Ara+1
Ara+2
Ara+4
Ara+5
Ara-1
Ara-2
Ara-3
Ara-4
Ara+3
Ara+6
Cumulative mutations (normalized)
Generations (x 10,000)

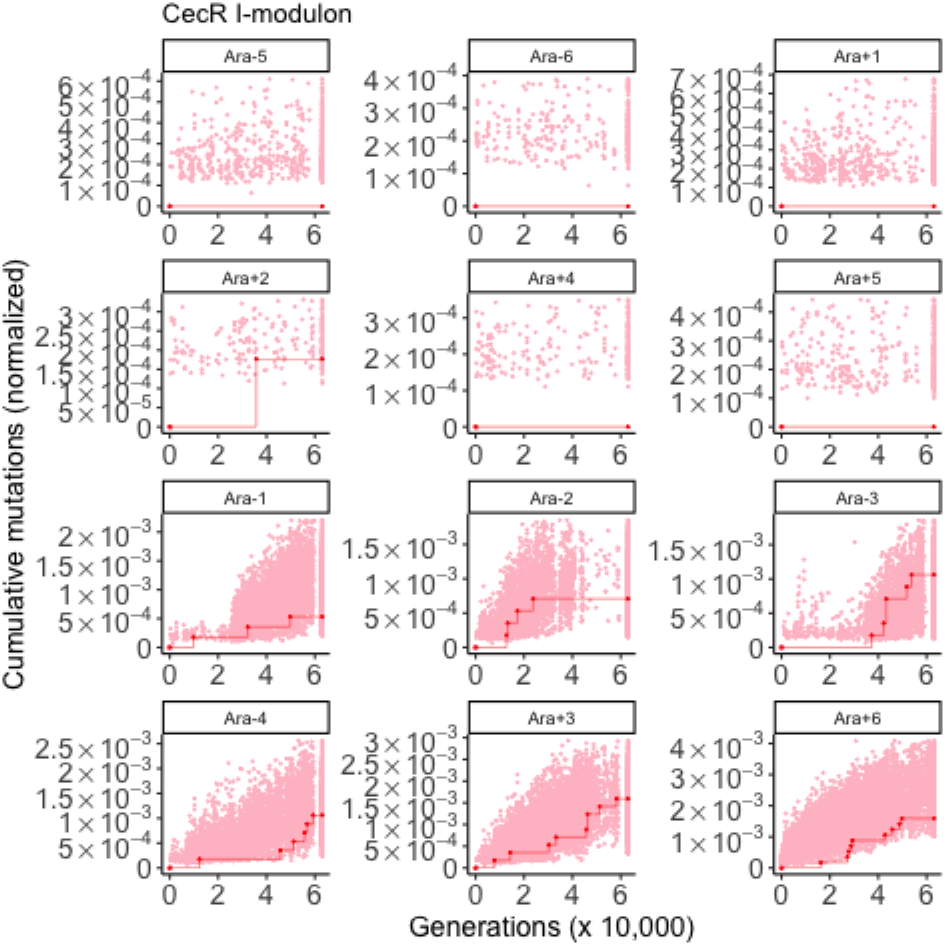
CecR I-modulon
Ara-5
Ara-6
Ara+1
Ara+2
Ara+4
Ara+5
Ara-1
Ara-2
Ara-3
Ara-4
Ara+3
Ara+6
Cumulative mutations (normalized)
Generations (x 10,000)

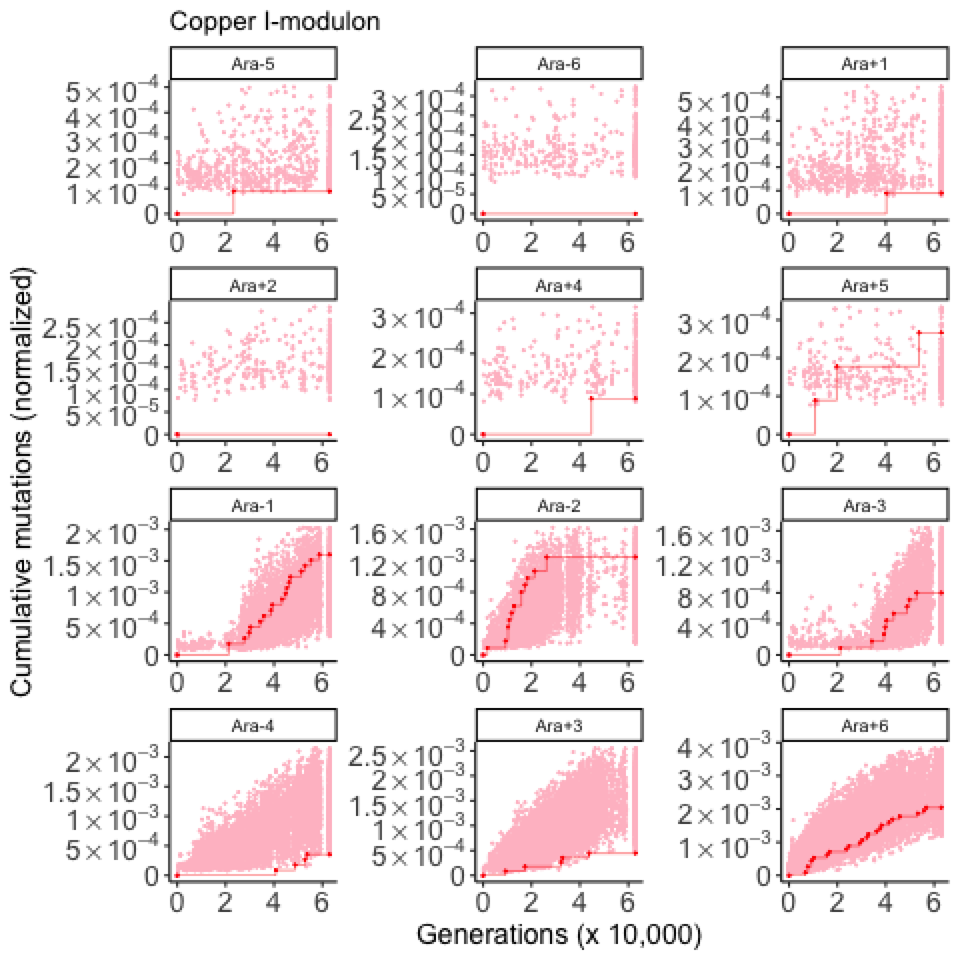
Copper I-modulon
Ara-5
Ara-6
Ara+1
Ara+2
Ara+4
Ara+5
Ara-1
Ara-2
Ara-3
Ara-4
Ara+3
Ara+6
Cumulative mutations (normalized)
Generations (x 10,000)

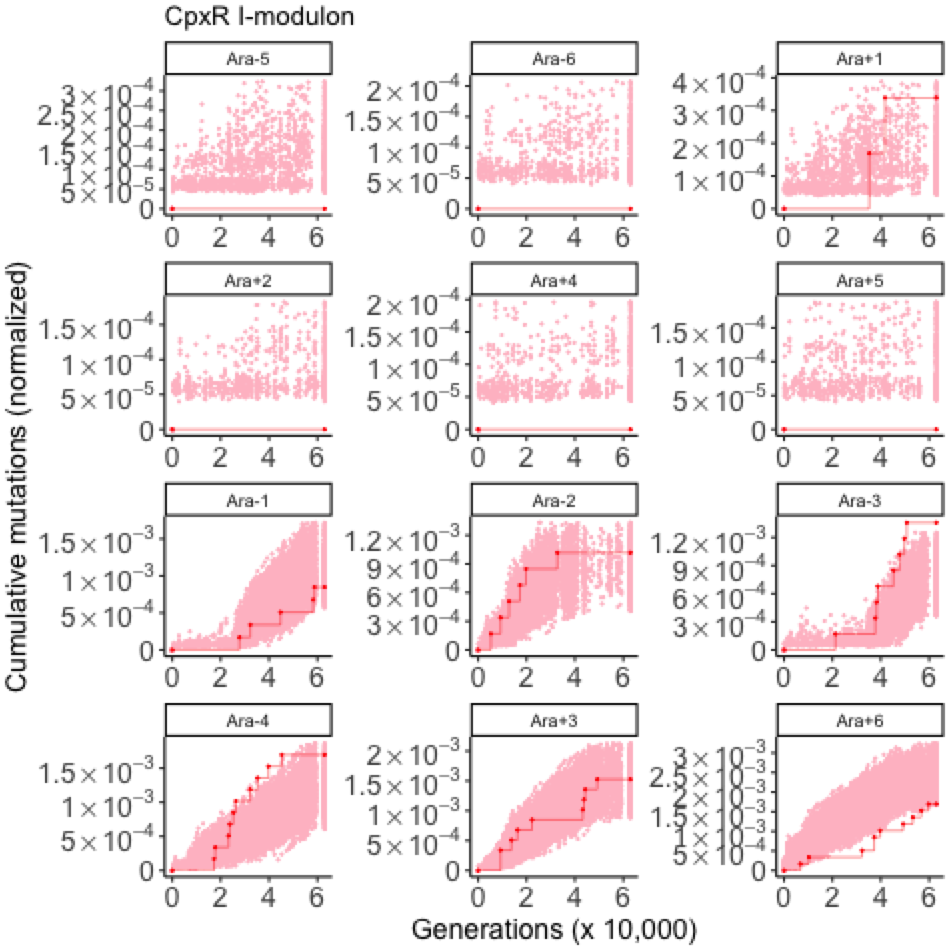
CpxR I-modulon
Ara-5
Ara-6
Ara+1
Ara+2
Ara+4
Ara+5
Ara-1
Ara-2
Ara-3
Ara-4
Ara+3
Ara+6
Cumulative mutations (normalized)
Generations (x 10,000)

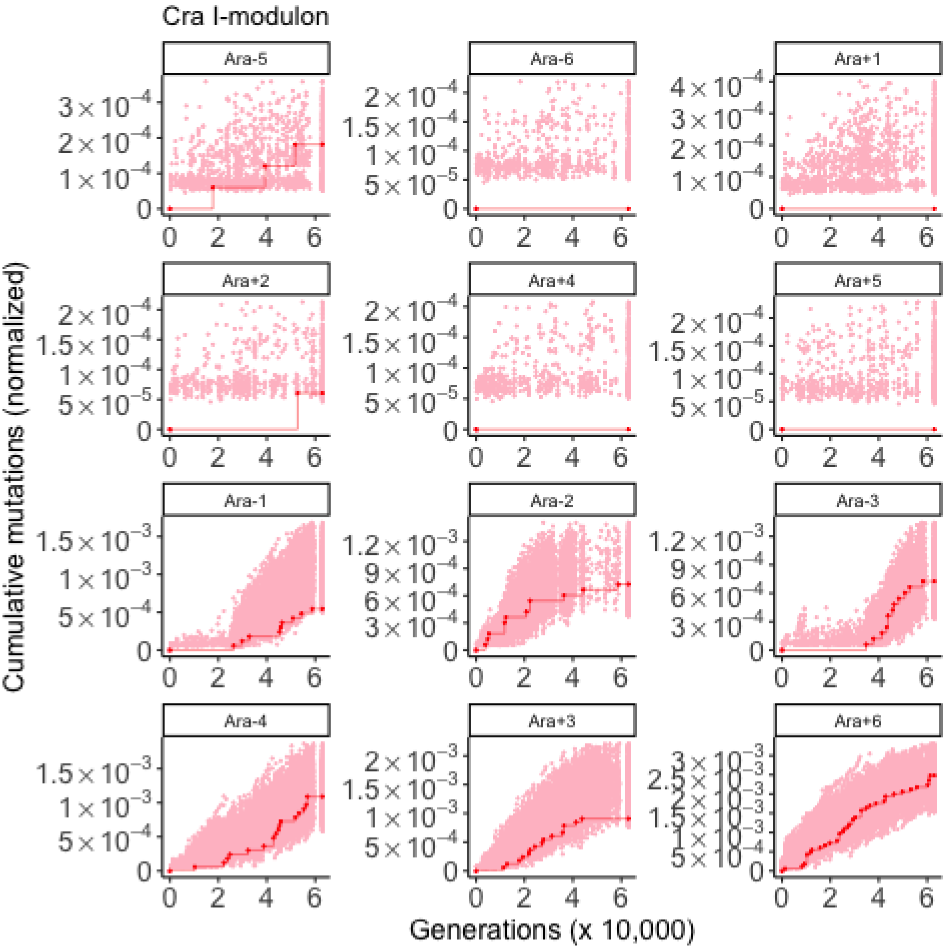
Cra I-modulon
Ara-5
Ara-6
Ara+1
Ara+2
Ara+4
Ara+5
Ara-1
Ara-2
Ara-3
Ara-4
Ara+3
Ara+6
Cumulative mutations (normalized)
Generations (x 10,000)

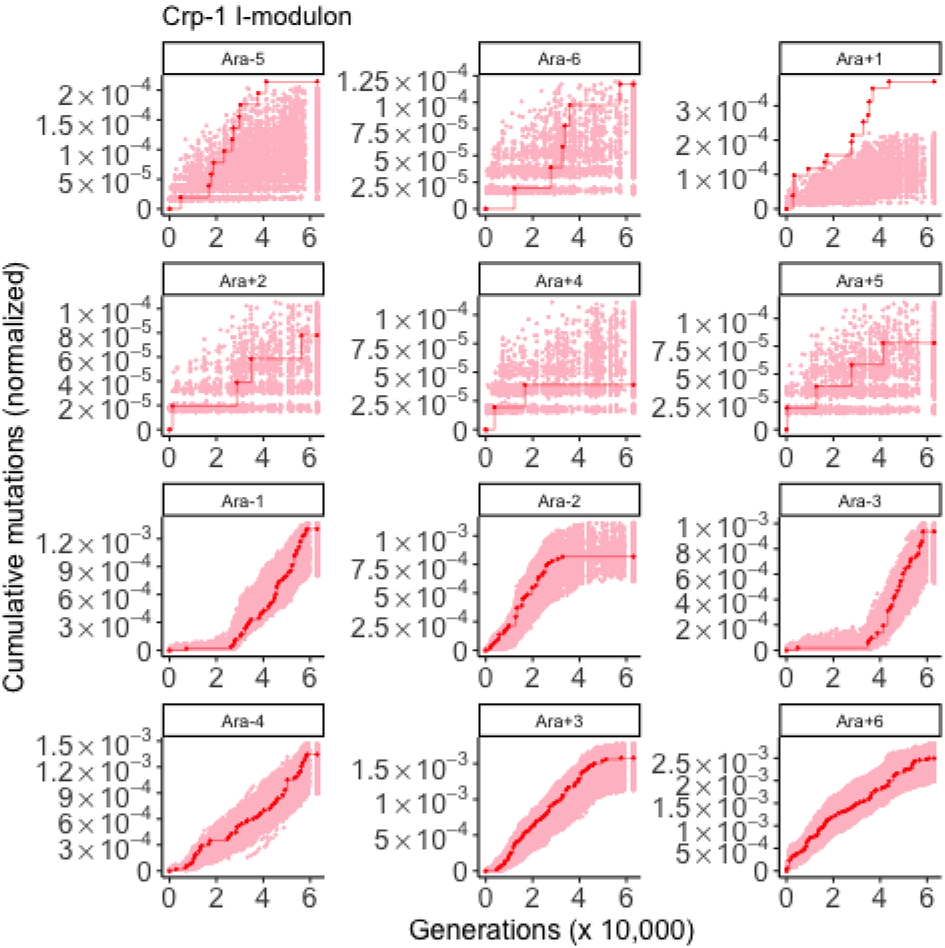

Crp-1 I-modulon
Ara-5
Ara-6
Ara+1
Ara+2
Ara+4
Ara+5
Ara-1
Ara-2
Ara-3
Ara-4
Ara+3
Ara+6
Cumulative mutations (normalized)
Generations (x 10,000)

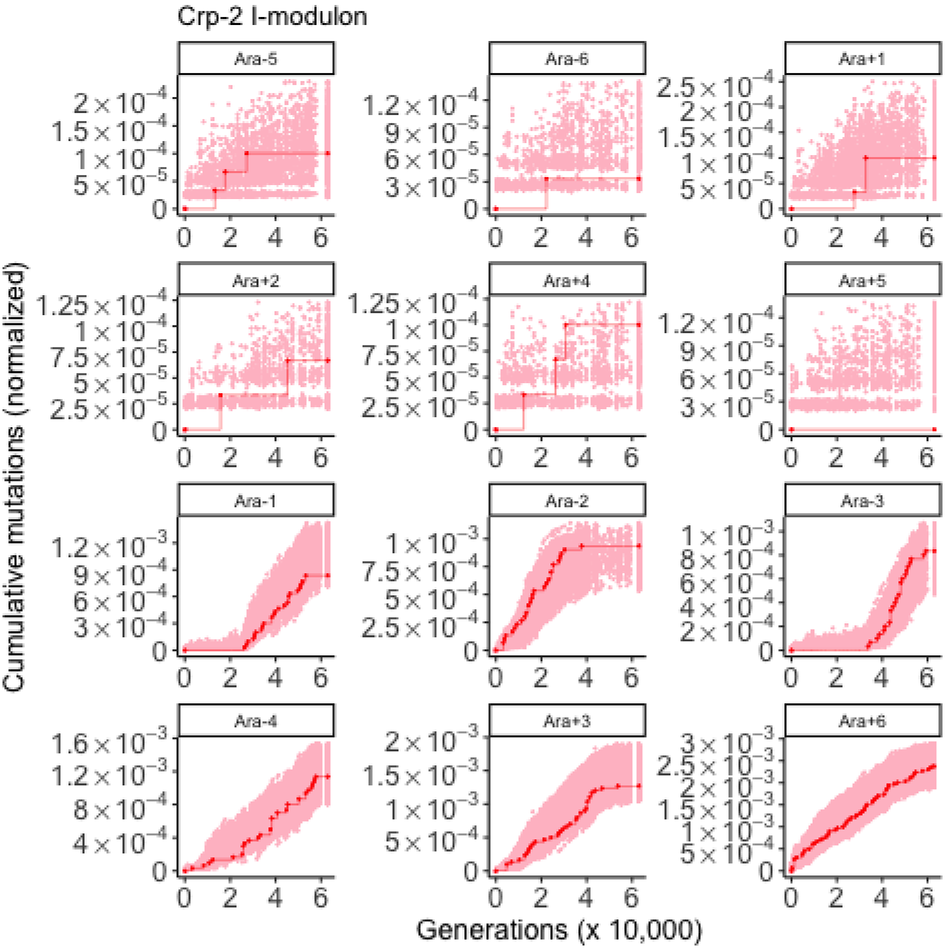

Crp-2 I-modulon
Ara-5
Ara-6
Ara+1
Ara+2
Ara+4
Ara+5
Ara-1
Ara-2
Ara-3
Ara-4
Ara+3
Ara+6
Cumulative mutations (normalized)
Generations (x 10,000)

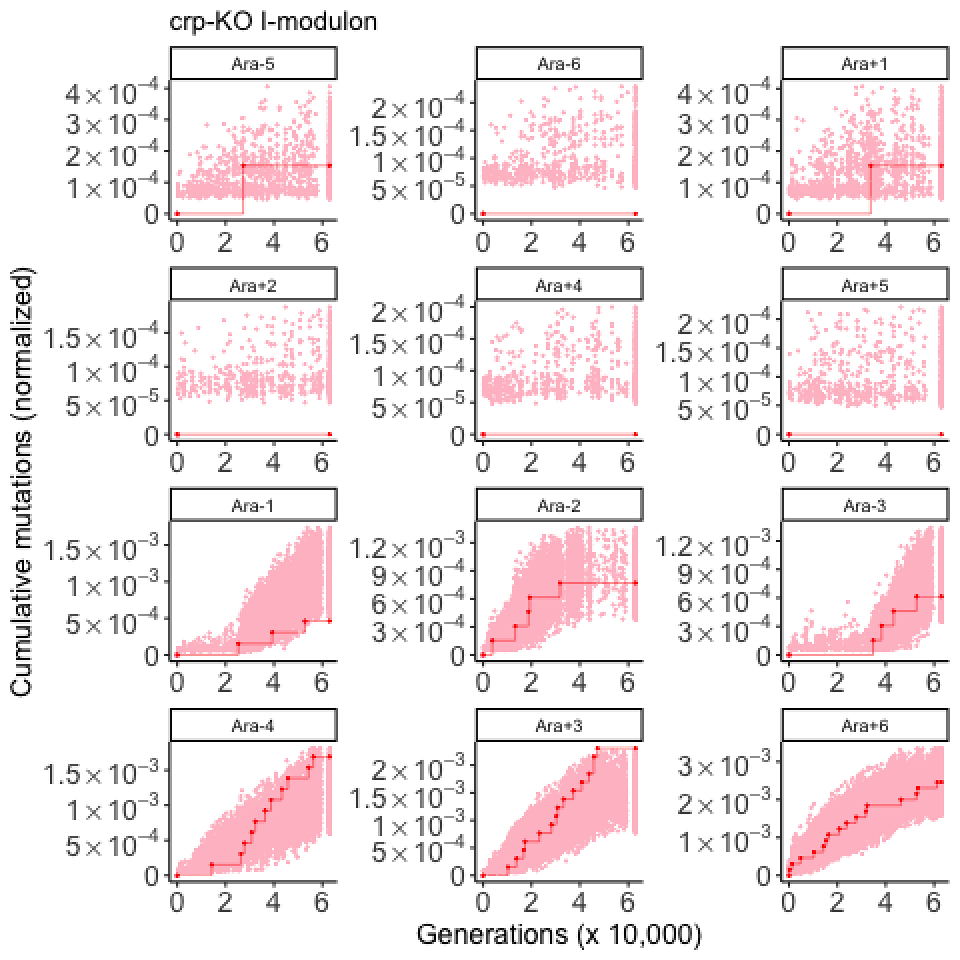

crp-KO I-modulon
Ara-5
Ara-6
Ara+1
Ara+2
Ara+4
Ara+5
Ara-1
Ara-2
Ara-3
Ara-4
Ara+3
Ara+6
Cumulative mutations (normalized)
Generations (x 10,000)

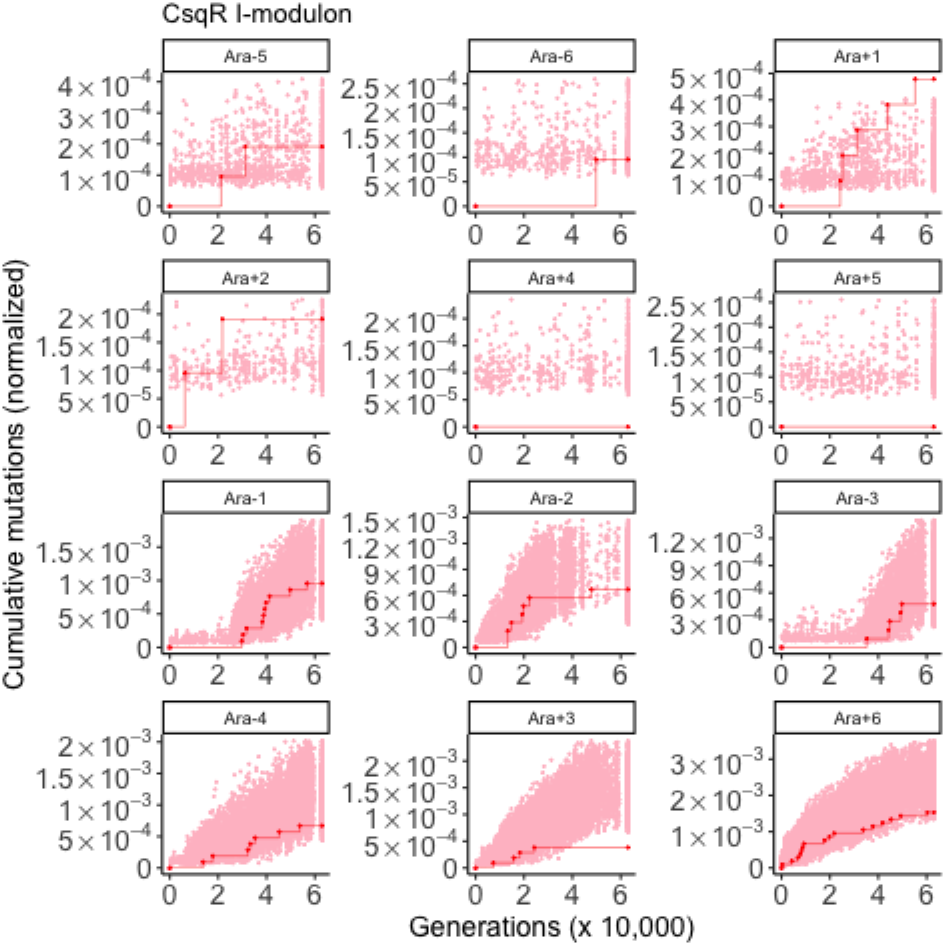

CsqR I-modulon
Ara-5
Ara-6
Ara+1
Ara+2
Ara+4
Ara+5
Ara-1
Ara-2
Ara-3
Ara-4
Ara+3
Ara+6
Cumulative mutations (normalized)
Generations (x 10,000)

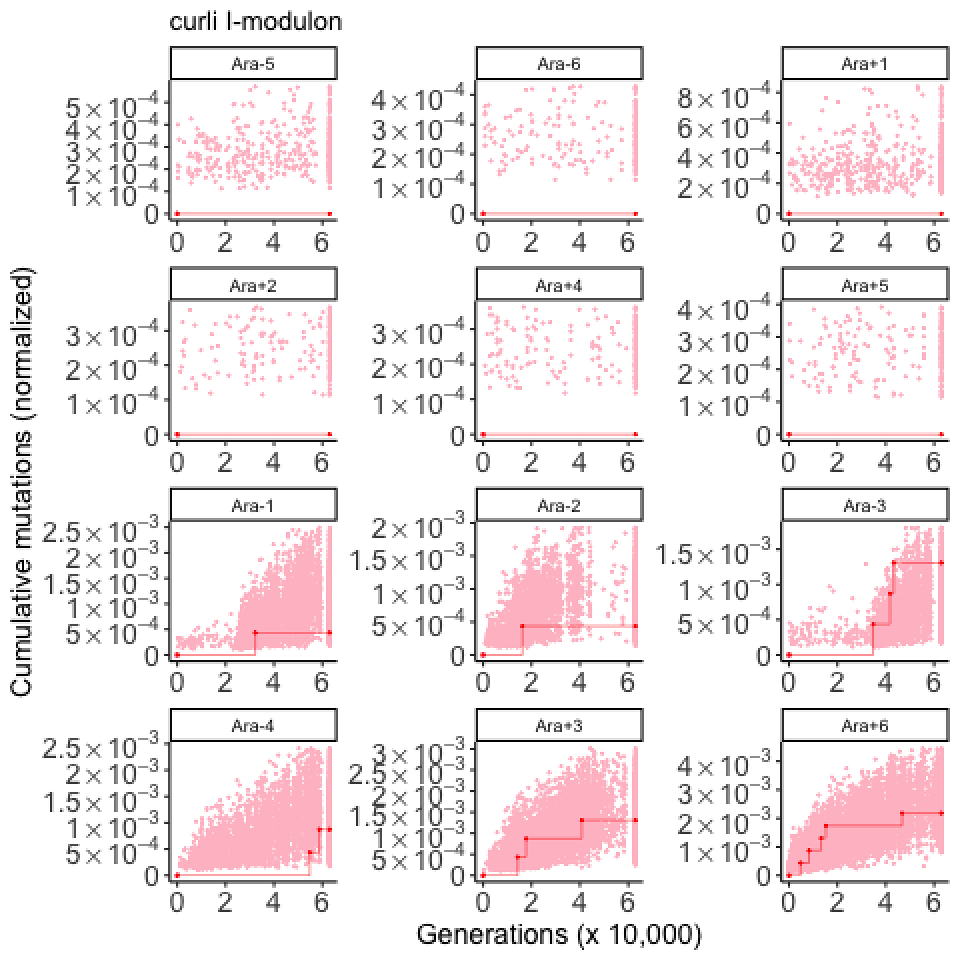
curli I-modulon
Ara-5
Ara-6
Ara+1
Ara+2
Ara+4
Ara+5
Ara-1
Ara-2
Ara-3
Ara-4
Ara+3
Ara+6
Cumulative mutations (normalized)
Generations (x 10,000)

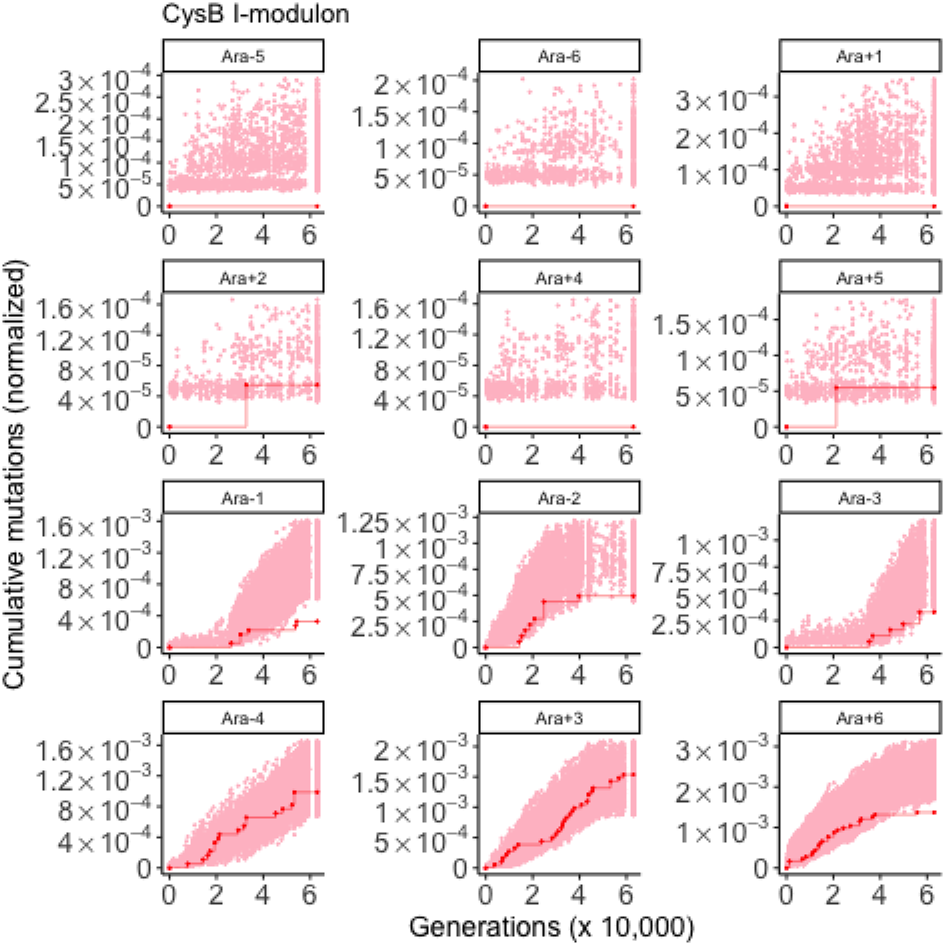
CysB I-modulon
Ara-5
Ara-6
Ara+1
Ara+2
Ara+4
Ara+5
Ara-1
Ara-2
Ara-3
Ara-4
Ara+3
Ara+6
Cumulative mutations (normalized)
Generations (x 10,000)

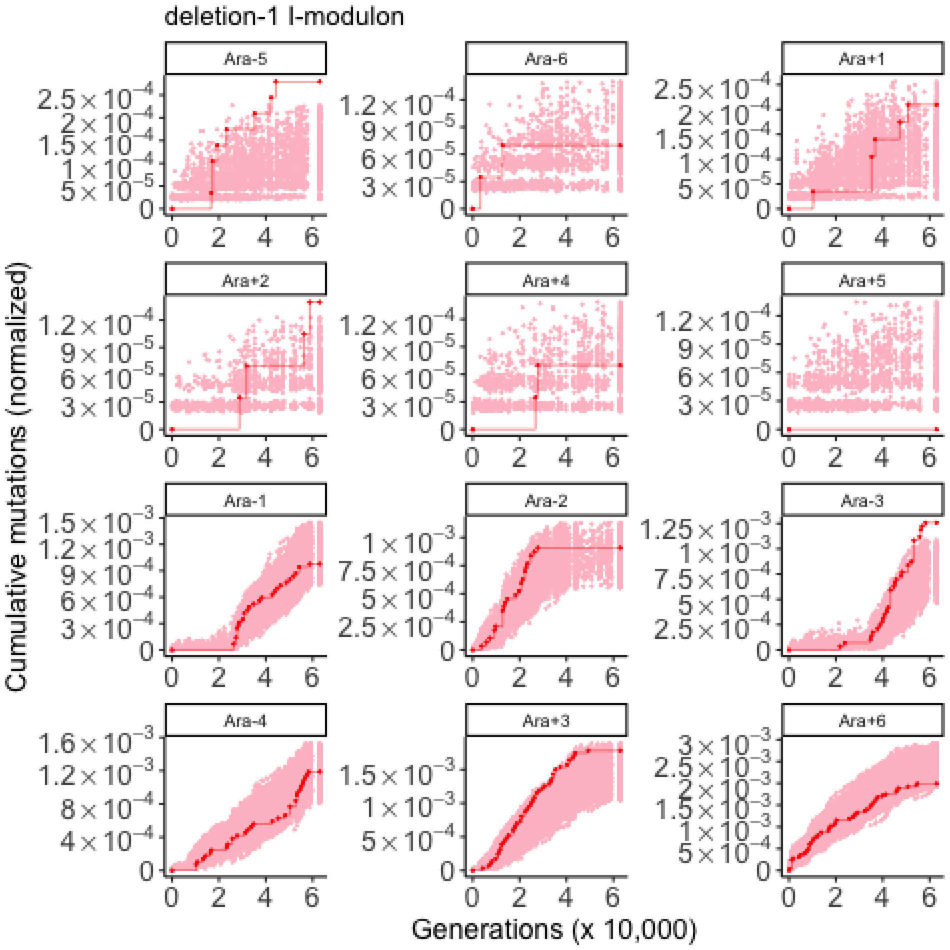
deletion-1 I-modulon
Ara-5
Ara-6
Ara+1
Ara+2
Ara+4
Ara+5
Ara-1
Ara-2
Ara-3
Ara-4
Ara+3
Ara+6
Cumulative mutations (normalized)
Generations (x 10,000)

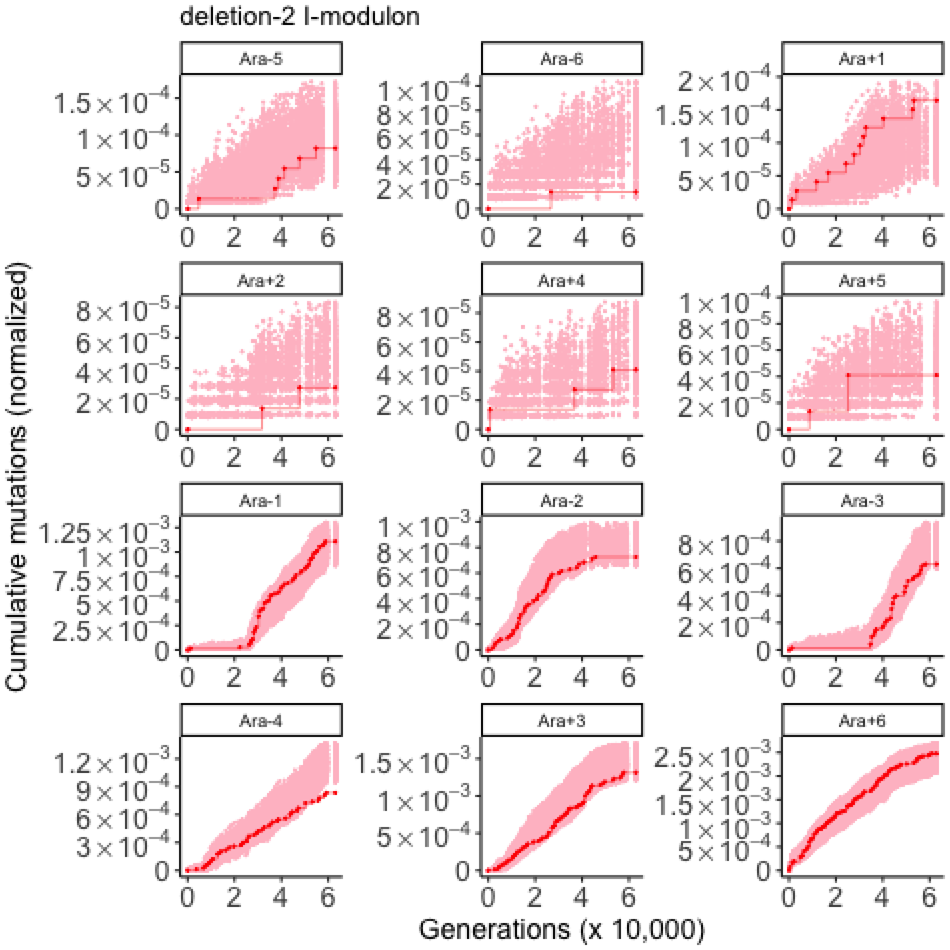

deletion-2 I-modulon
Ara-5
Ara-6
Ara+1
Ara+2
Ara+4
Ara+5
Ara-1
Ara-2
Ara-3
Ara-4
Ara+3
Ara+6
Cumulative mutations (normalized)
Generations (x 10,000)

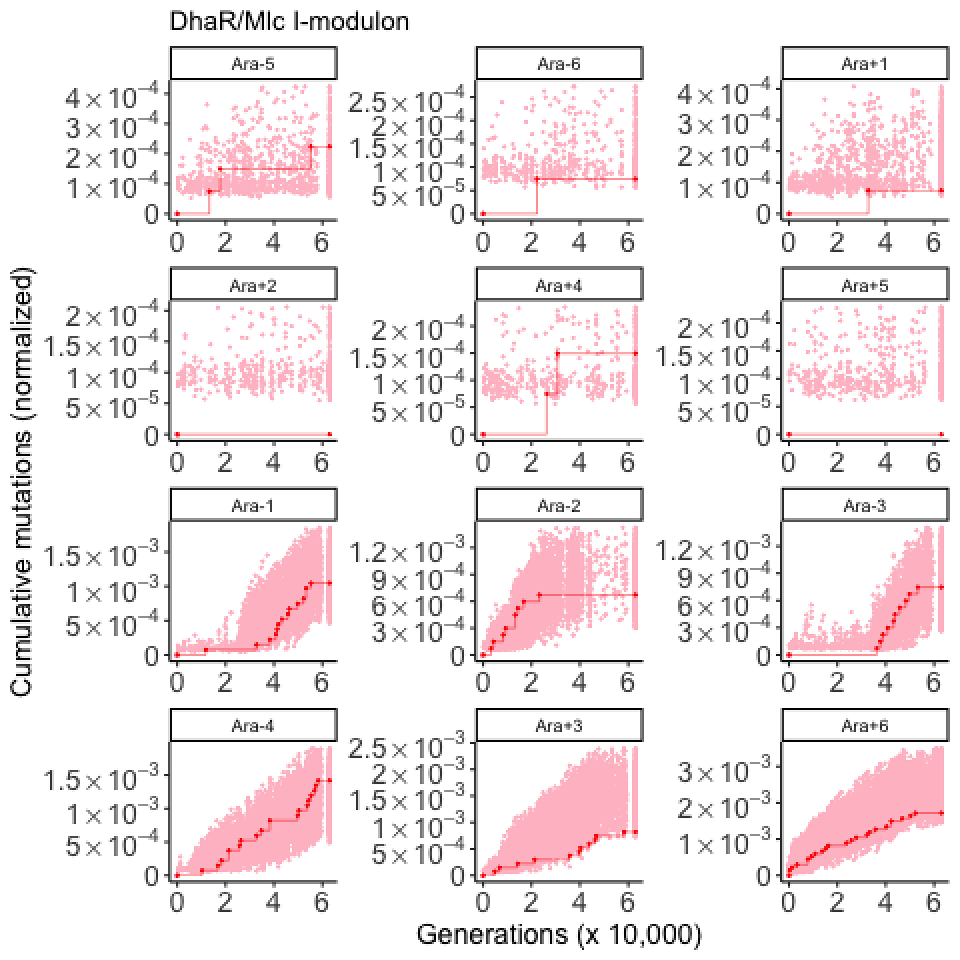
DhaR/Mlc I-modulon
Ara-5
Ara-6
Ara+1
Ara+2
Ara+4
Ara+5
Ara-1
Ara-2
Ara-3
Ara-4
Ara+3
Ara+6
Cumulative mutations (normalized)
Generations (x 10,000)

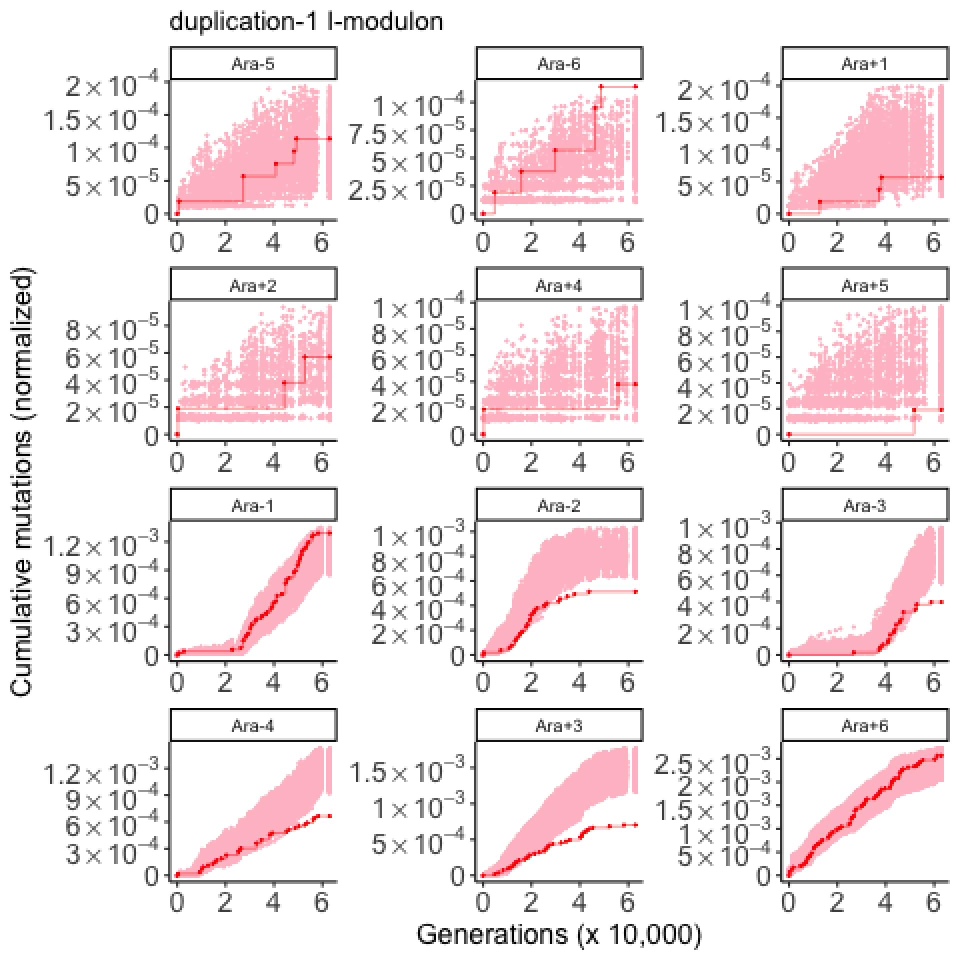

duplication-1 I-modulon
Ara-5
Ara-6
Ara+1
Ara+2
Ara+4
Ara+5
Ara-1
Ara-2
Ara-3
Ara-4
Ara+3
Ara+6
Cumulative mutations (normalized)
Generations (x 10,000)

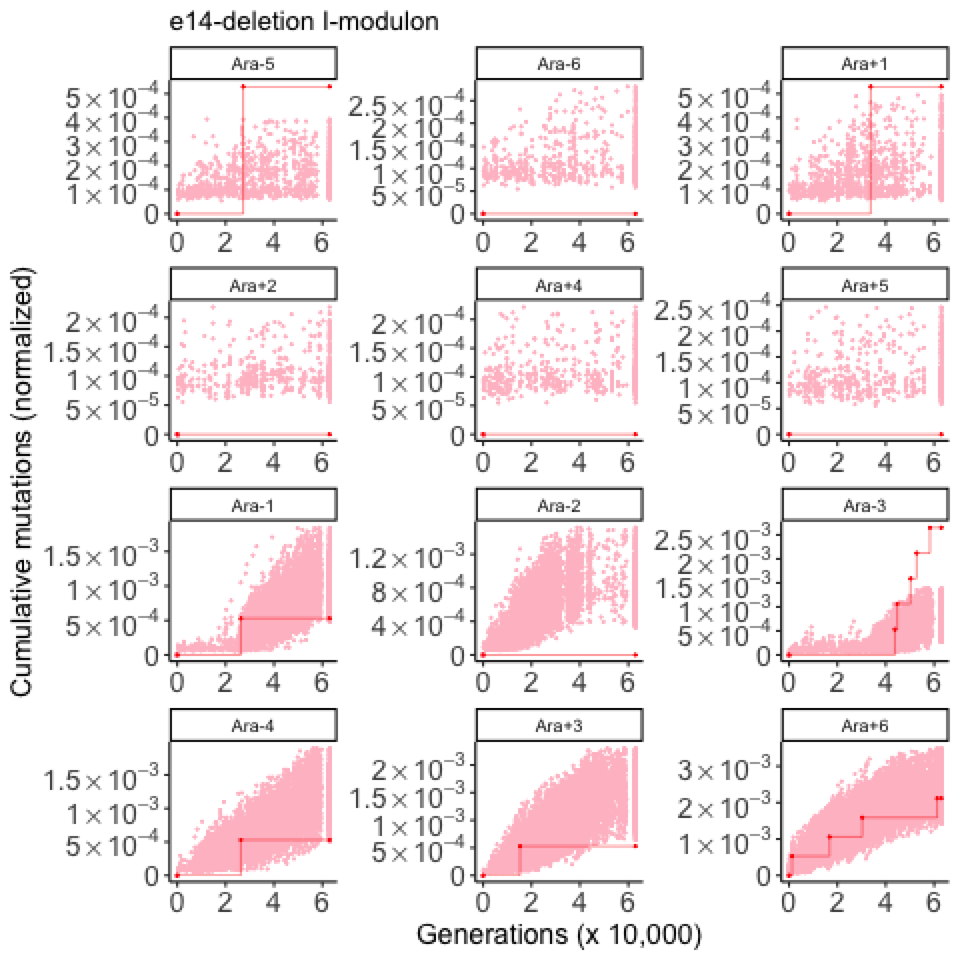

e14-deletion I-modulon
Ara-5
Ara-6
Ara+1
Ara+2
Ara+4
Ara+5
Ara-1
Ara-2
Ara-3
Ara-4
Ara+3
Ara+6
Cumulative mutations (normalized)
Generations (x 10,000)

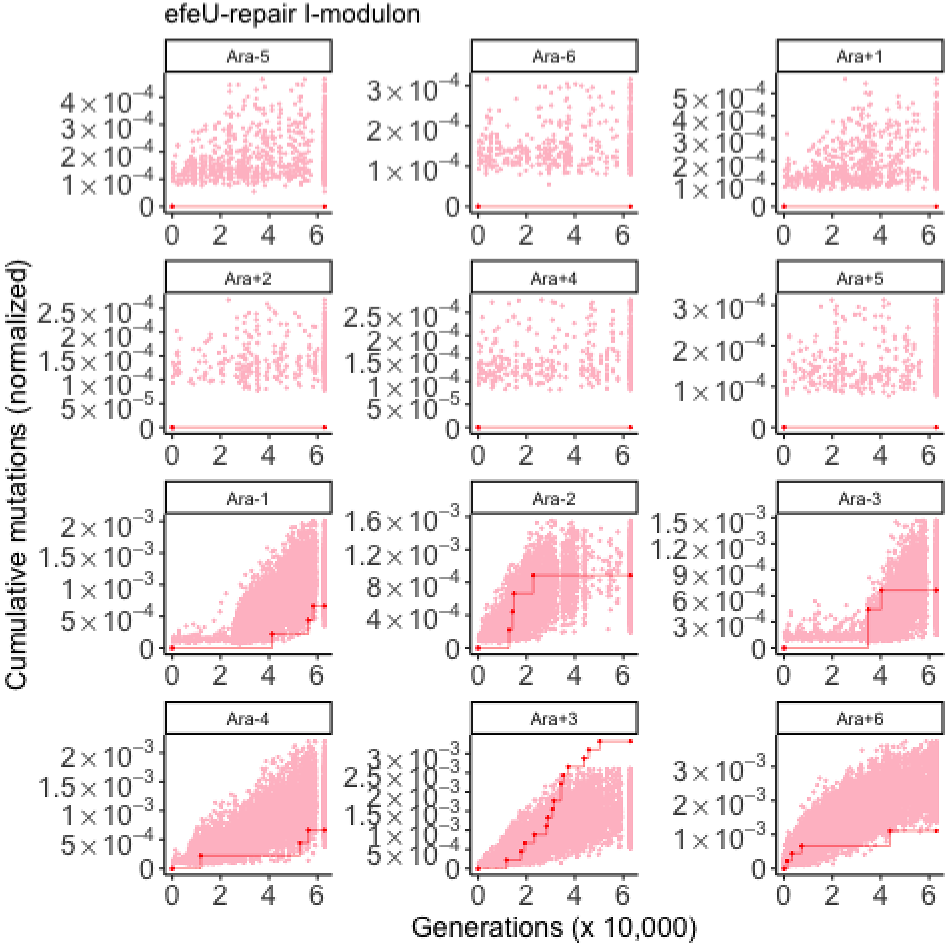
efeU-repair I-modulon
Ara-5
Ara-6
Ara+1
Ara+2
Ara+4
Ara+5
Ara-1
Ara-2
Ara-3
Ara-4
Ara+3
Ara+6
Cumulative mutations (normalized)
Generations (x 10,000)

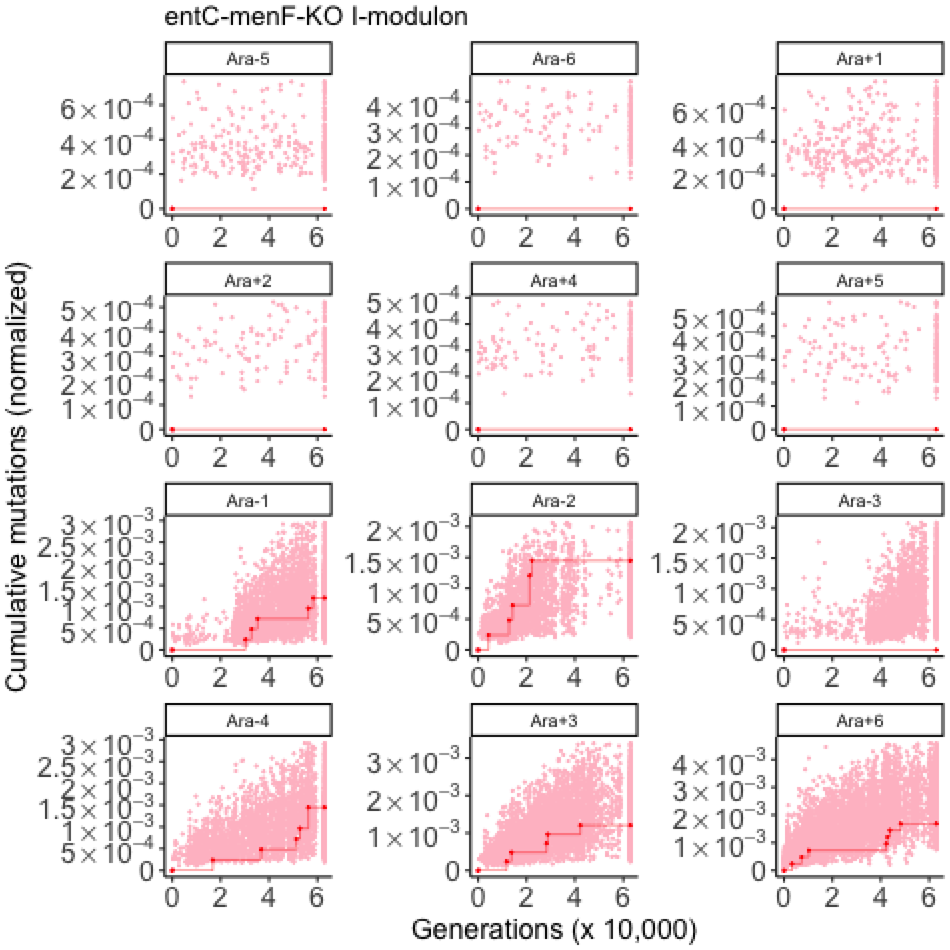
entC-menF-KO I-modulon
Ara-5
Ara-6
Ara+1
Ara+2
Ara+4
Ara+5
Ara-1
Ara-2
Ara-3
Ara-4
Ara+3
Ara+6
Cumulative mutations (normalized)
Generations (x 10,000)

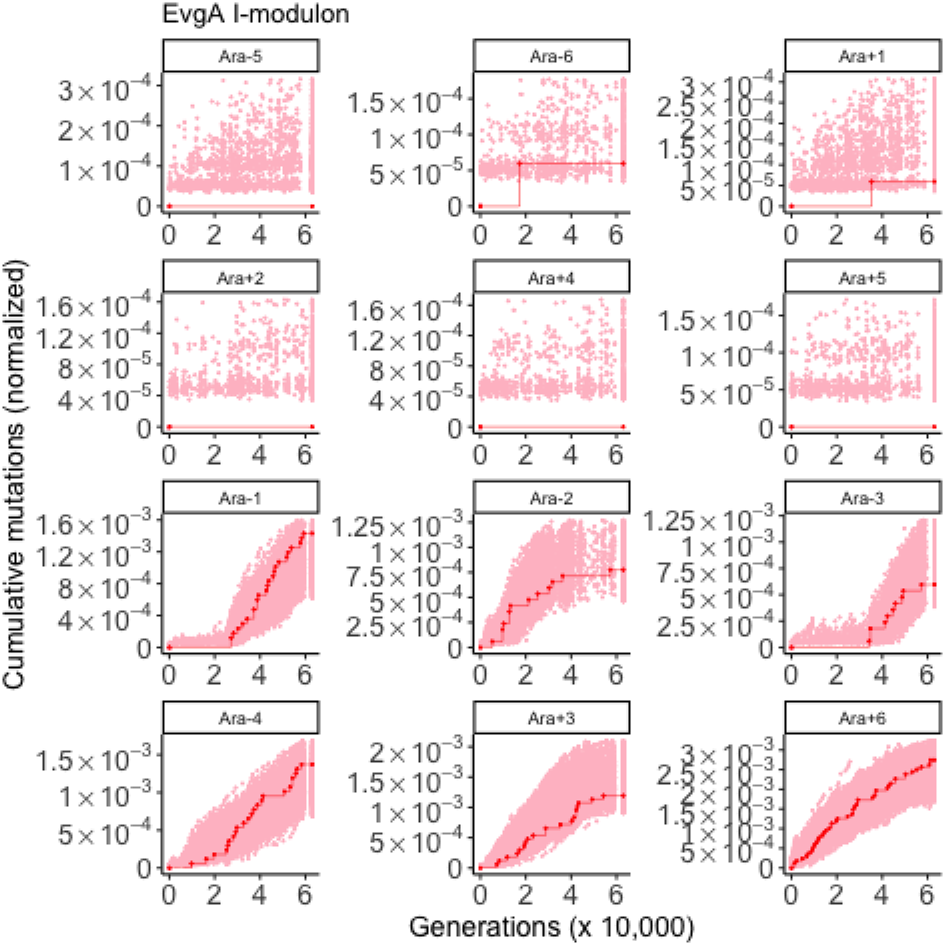
EvgA I-modulon
Ara-5
Ara-6
Ara+1
Ara+2
Ara+4
Ara+5
Ara-1
Ara-2
Ara-3
Ara-4
Ara+3
Ara+6
Cumulative mutations (normalized)
Generations (x 10,000)

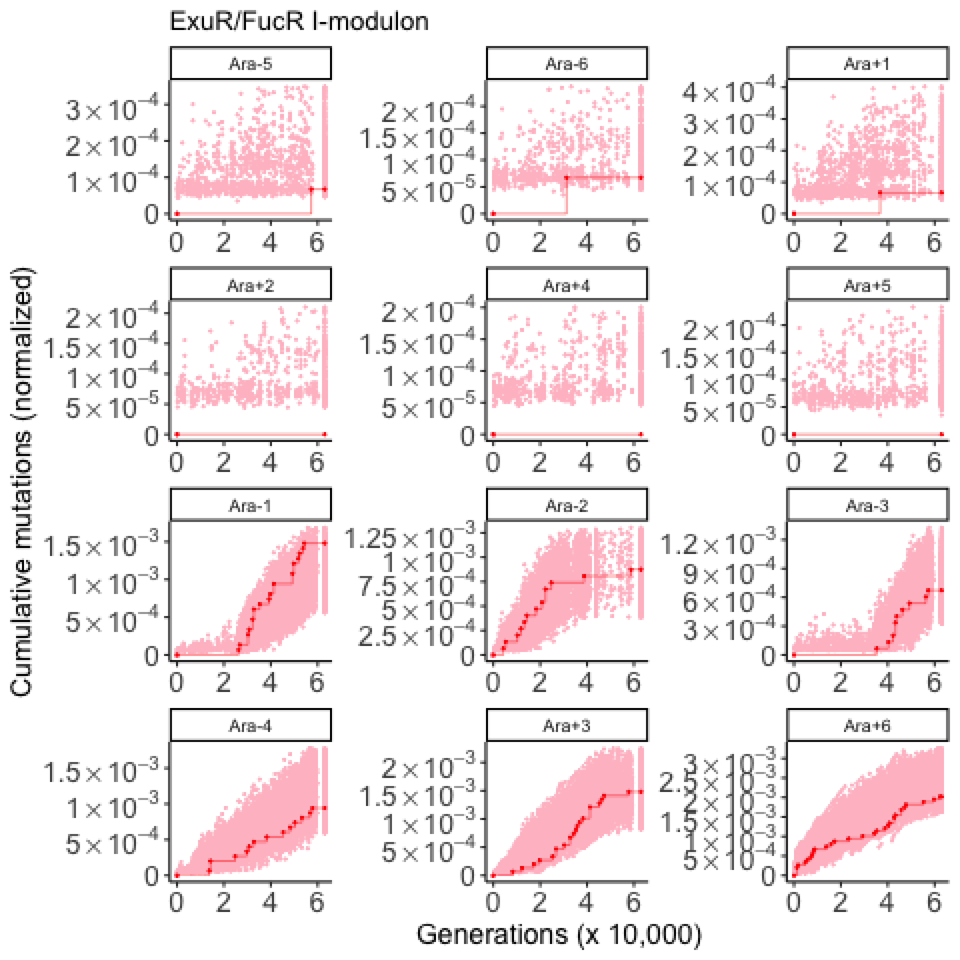
ExuR/FucR I-modulon
Ara-5
Ara-6
Ara+1
Ara+2
Ara+4
Ara+5
Ara-1
Ara-2
Ara-3
Ara-4
Ara+3
Ara+6
Cumulative mutations (normalized)
Generations (x 10,000)

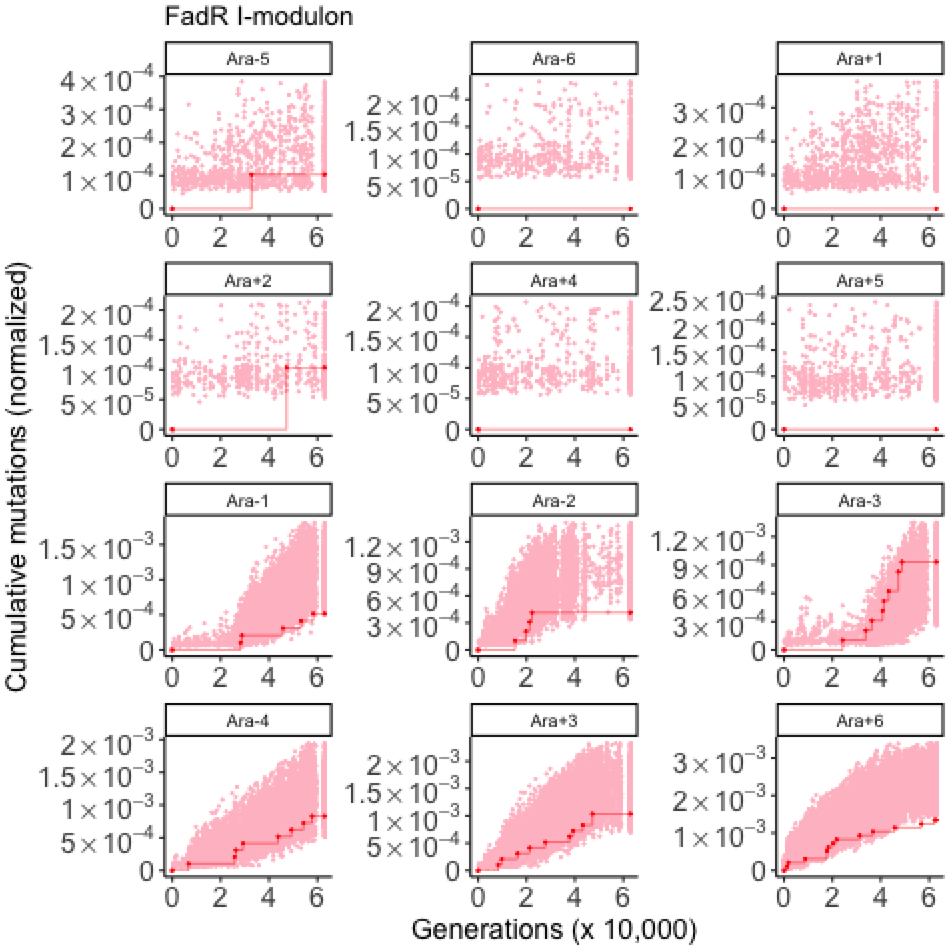

FadR I-modulon
Ara-5
Ara-6
Ara+1
Ara+2
Ara+4
Ara+5
Ara-1
Ara-2
Ara-3
Ara-4
Ara+3
Ara+6
Cumulative mutations (normalized)
Generations (x 10,000)

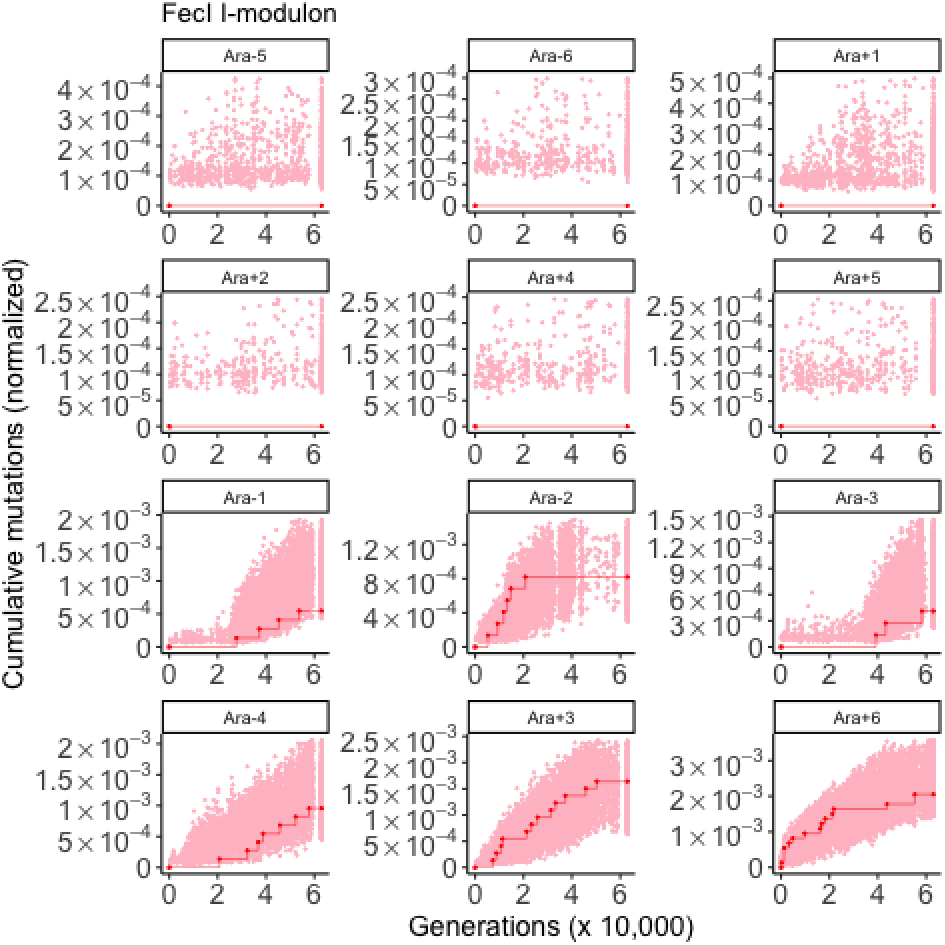

FecI I-modulon
Ara-5
Ara-6
Ara+1
Ara+2
Ara+4
Ara+5
Ara-1
Ara-2
Ara-3
Ara-4
Ara+3
Ara+6
Cumulative mutations (normalized)
Generations (x 10,000)

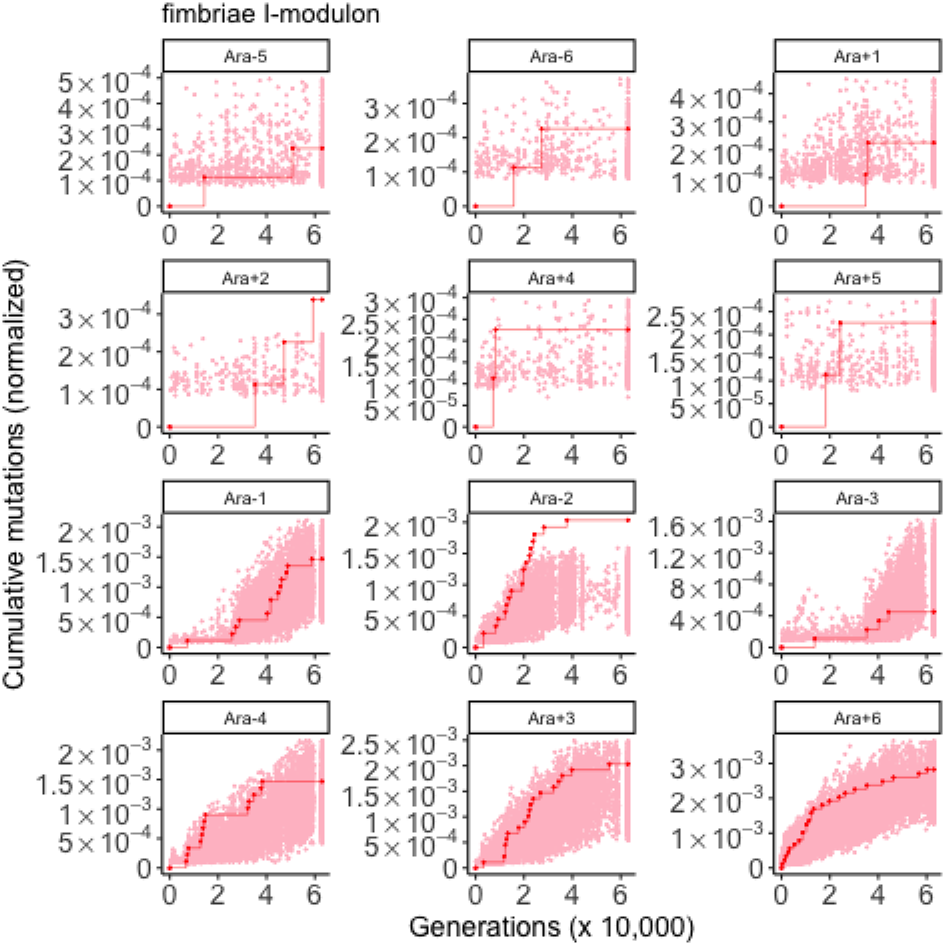

fimbriae I-modulon
Ara-5
Ara-6
Ara+1
Ara+2
Ara+4
Ara+5
Ara-1
Ara-2
Ara-3
Ara-4
Ara+3
Ara+6
Cumulative mutations (normalized)
Generations (x 10,000)

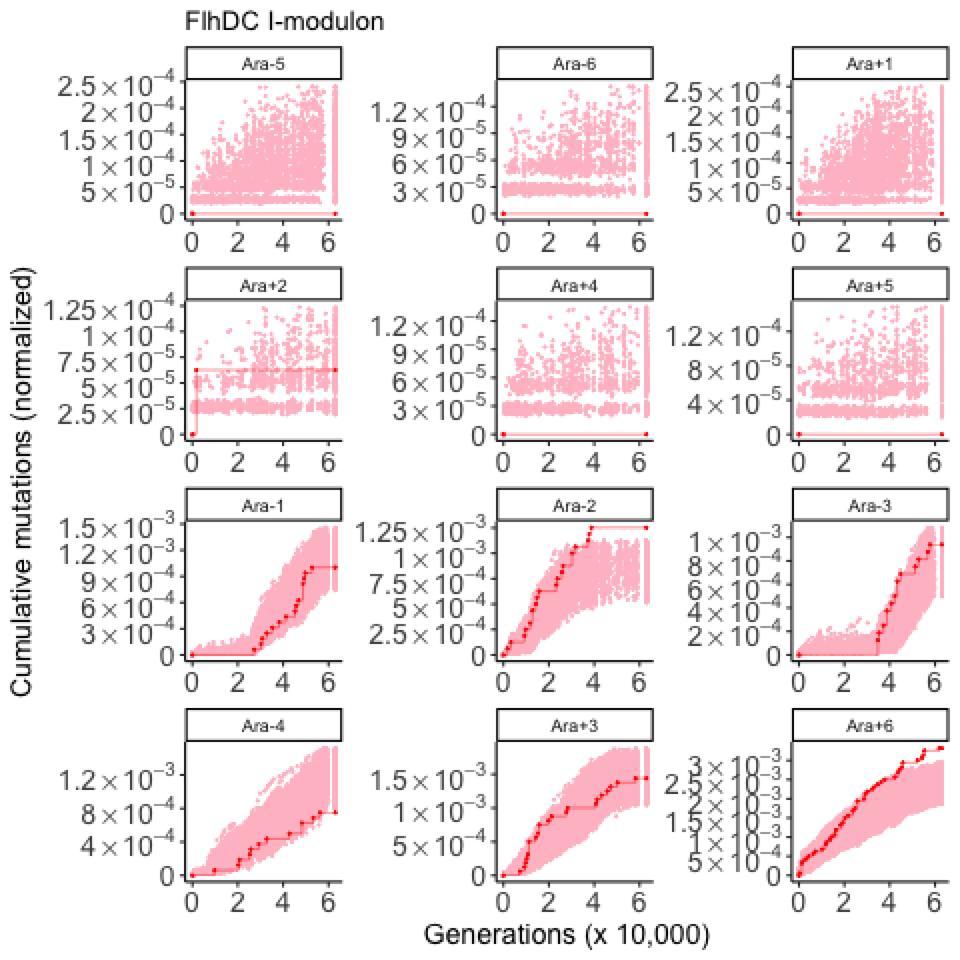
FlhDC I-modulon
Ara-5
Ara-6
Ara+1
Ara+2
Ara+4
Ara+5
Ara-1
Ara-2
Ara-3
Ara-4
Ara+3
Ara+6
Cumulative mutations (normalized)
Generations (x 10,000)

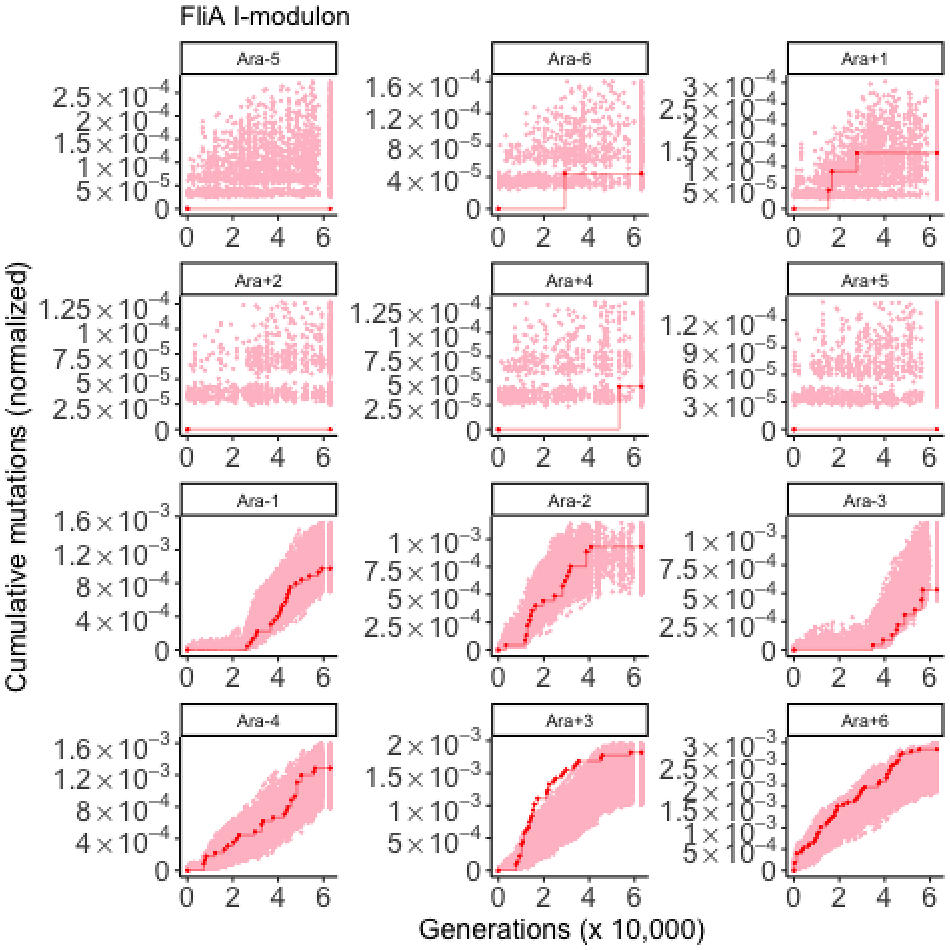
FliA I-modulon
Ara-5
Ara-6
Ara+1
Ara+2
Ara+4
Ara+5
Ara-1
Ara-2
Ara-3
Ara-4
Ara+3
Ara+6
Cumulative mutations (normalized)
Generations (x 10,000)

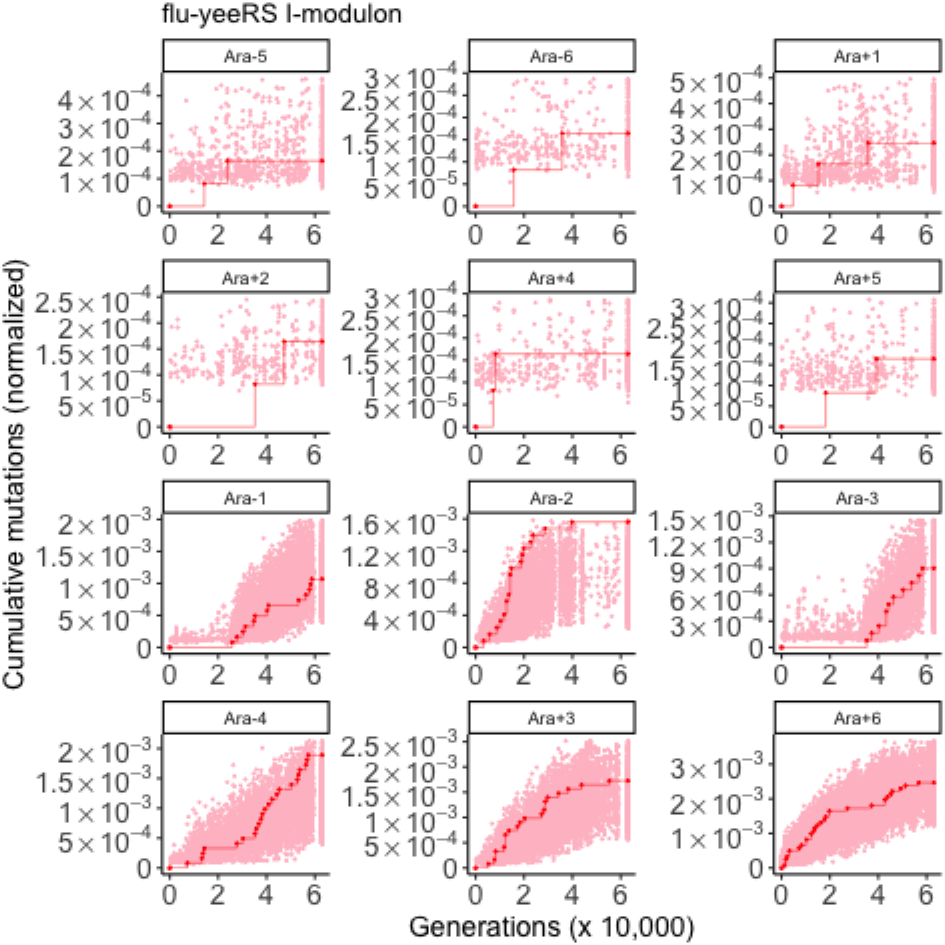
flu-yeeRS I-modulon
Ara-5
Ara-6
Ara+1
Ara+2
Ara+4
Ara+5
Ara-1
Ara-2
Ara-3
Ara-4
Ara+3
Ara+6
Cumulative mutations (normalized)
Generations (x 10,000)

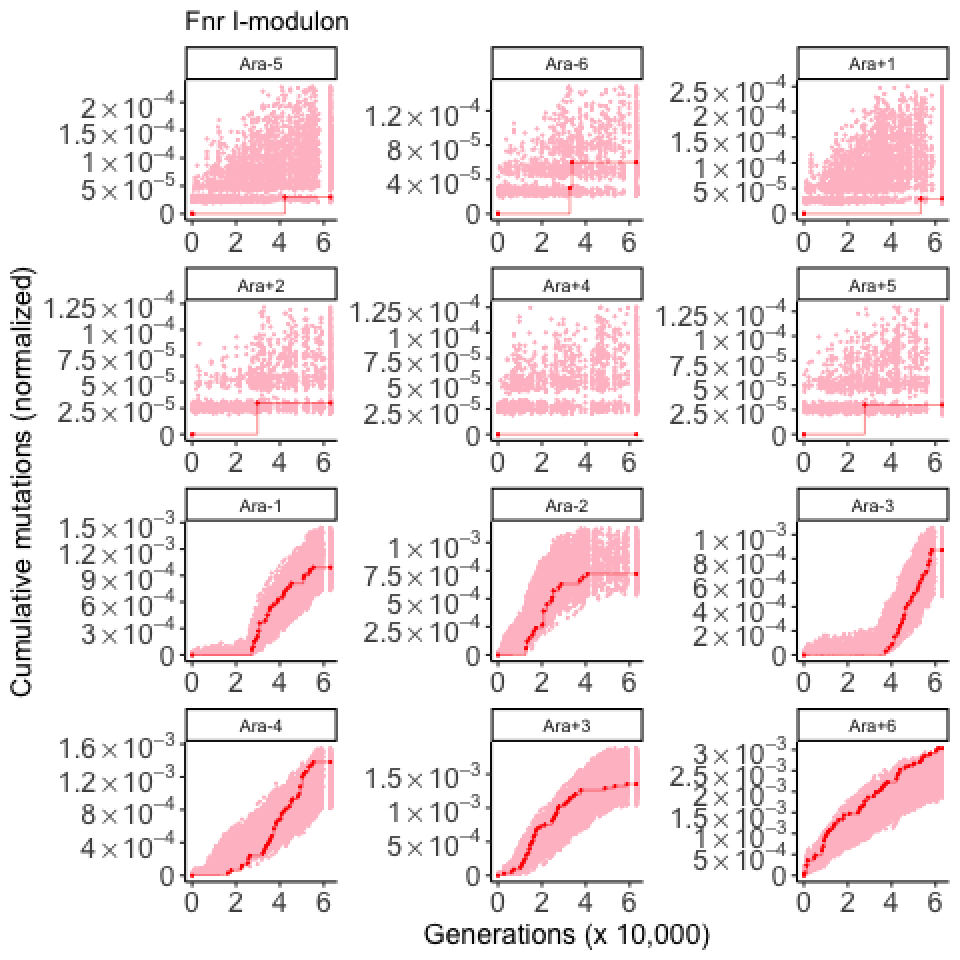

Fnr I-modulon
Ara-5
Ara-6
Ara+1
Ara+2
Ara+4
Ara+5
Ara-1
Ara-2
Ara-3
Ara-4
Ara+3
Ara+6
Cumulative mutations (normalized)
Generations (x 10,000)

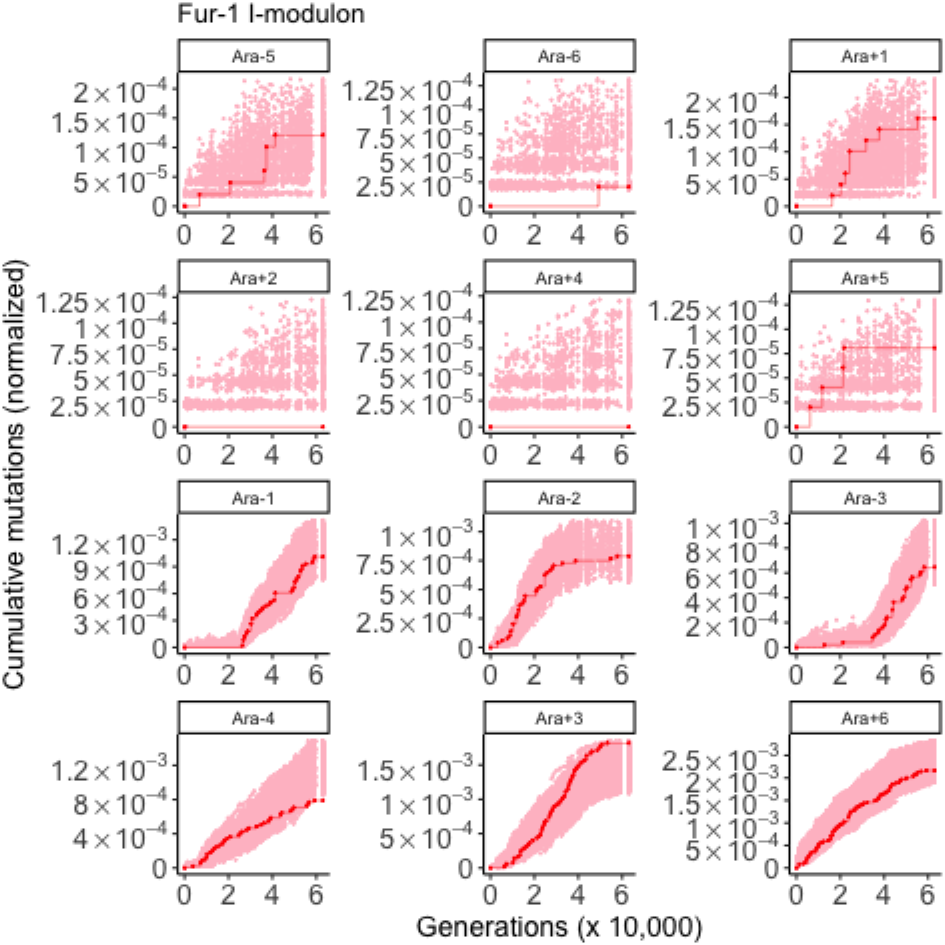
Fur-1 I-modulon
Ara-5
Ara-6
Ara+1
Ara+2
Ara+4
Ara+5
Ara-1
Ara-2
Ara-3
Ara-4
Ara+3
Ara+6
Cumulative mutations (normalized)
Generations (x 10,000)

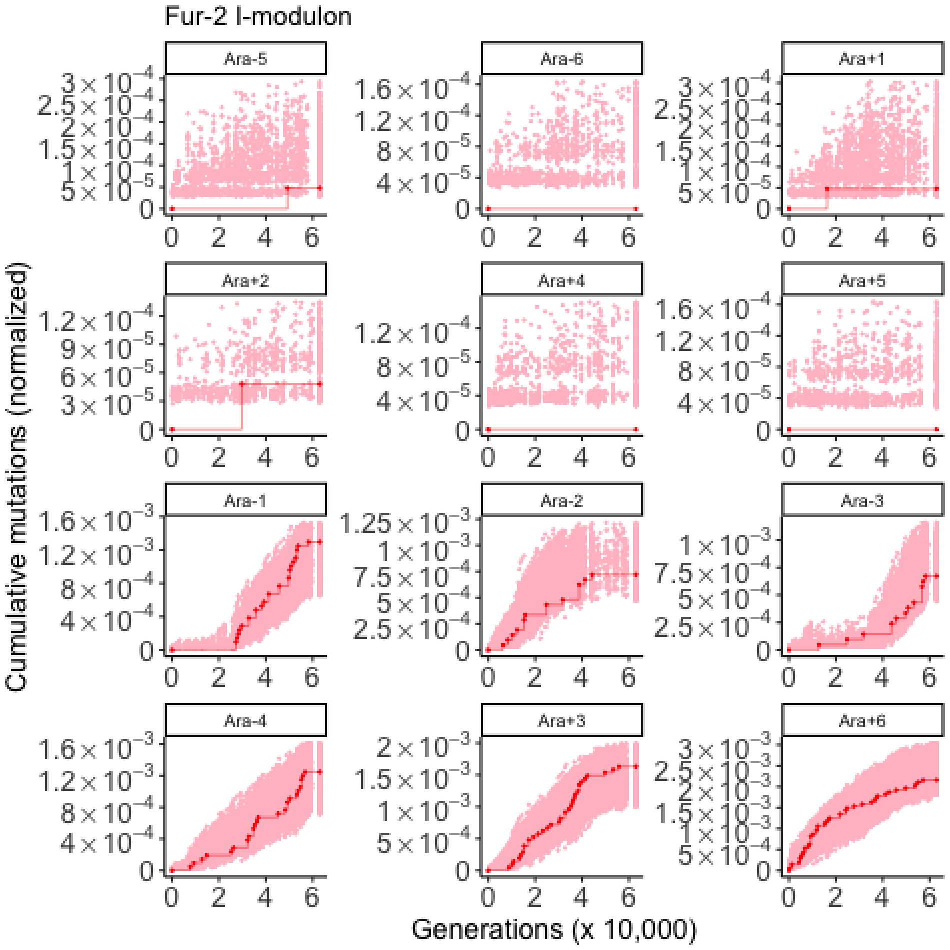

Fur-2 I-modulon
Ara-5
Ara-6
Ara+1
Ara+2
Ara+4
Ara+5
Ara-1
Ara-2
Ara-3
Ara-4
Ara+3
Ara+6
Cumulative mutations (normalized)
Generations (x 10,000)

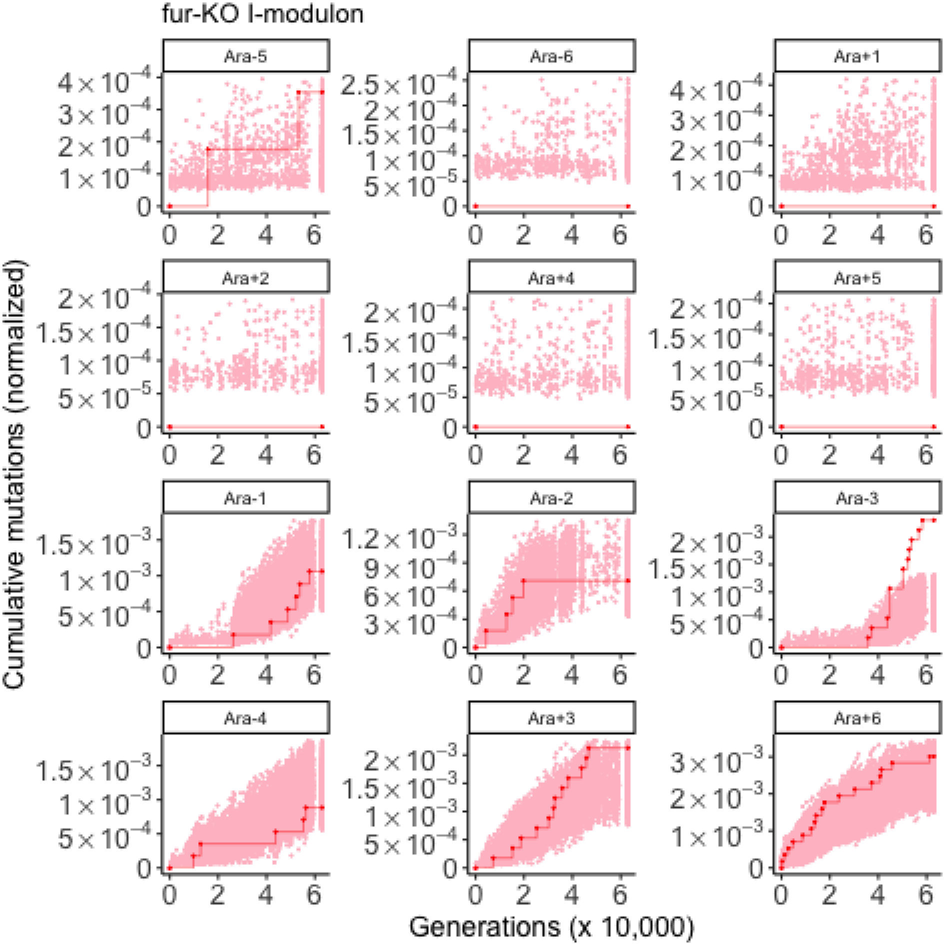
fur-KO I-modulon
Ara-5
Ara-6
Ara+1
Ara+2
Ara+4
Ara+5
Ara-1
Ara-2
Ara-3
Ara-4
Ara+3
Ara+6
Cumulative mutations (normalized)
Generations (x 10,000)

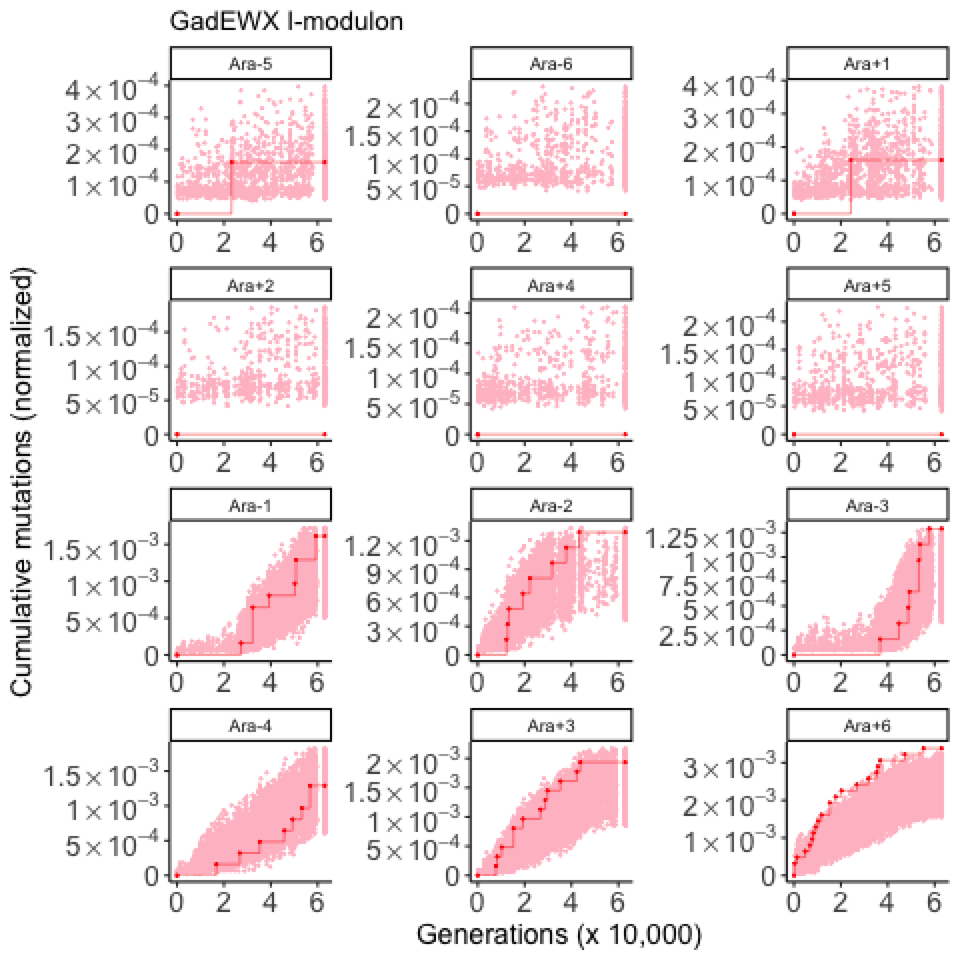
GadEWX I-modulon
Ara-5
Ara-6
Ara+1
Ara+2
Ara+4
Ara+5
Ara-1
Ara-2
Ara-3
Ara-4
Ara+3
Ara+6
Cumulative mutations (normalized)
Generations (x 10,000)

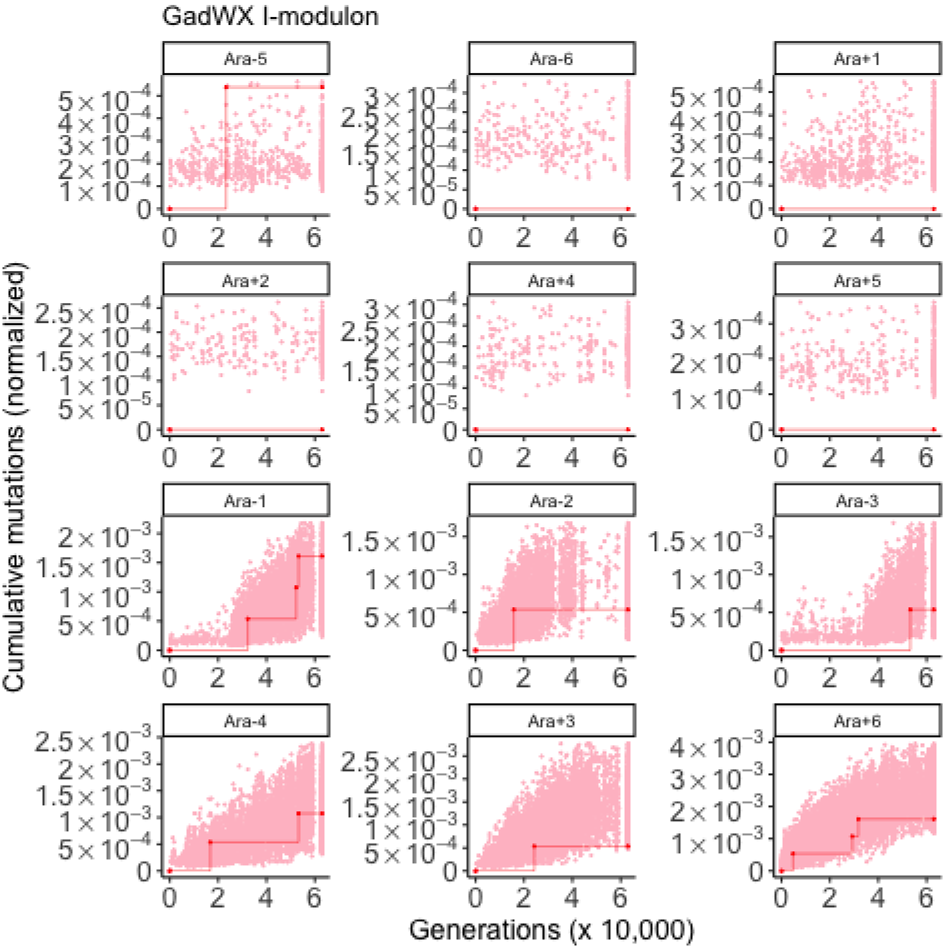
GadWX I-modulon
Ara-5
Ara-6
Ara+1
Ara+2
Ara+4
Ara+5
Ara-1
Ara-2
Ara-3
Ara-4
Ara+3
Ara+6
Cumulative mutations (normalized)
Generations (x 10,000)

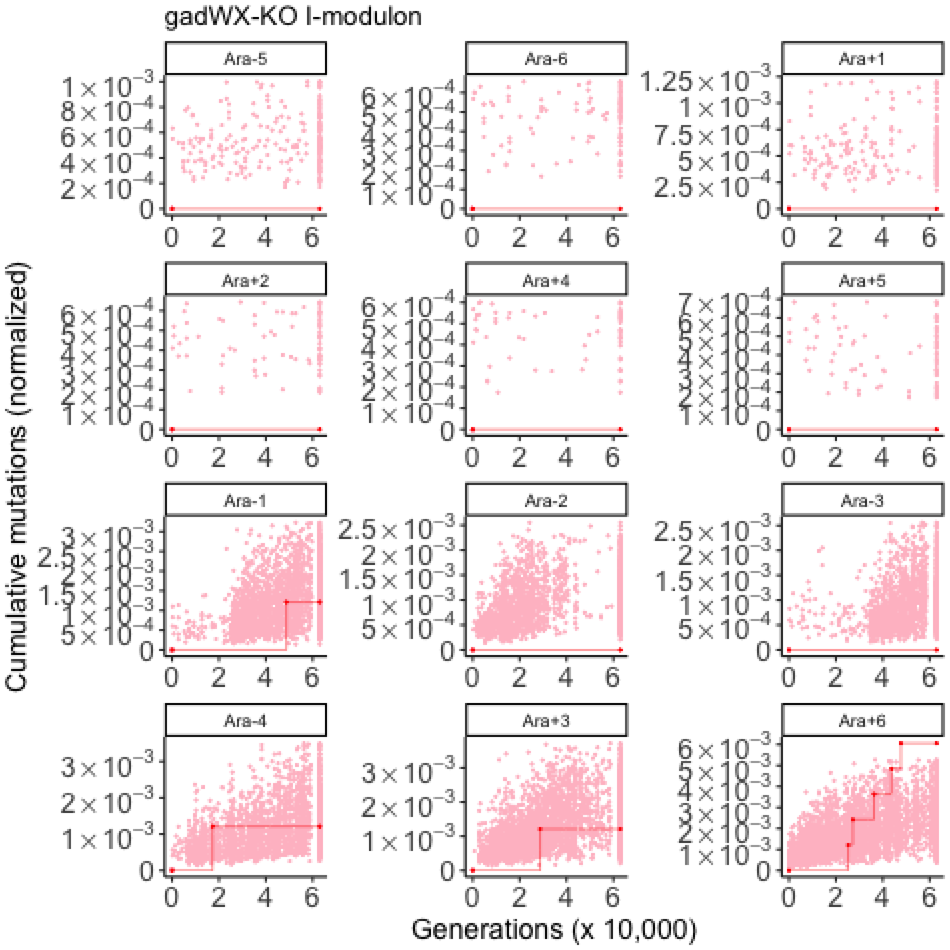

gadWX-KO I-modulon
Ara-5
Ara-6
Ara+1
Ara+2
Ara+4
Ara+5
Ara-1
Ara-2
Ara-3
Ara-4
Ara+3
Ara+6
Cumulative mutations (normalized)
Generations (x 10,000)

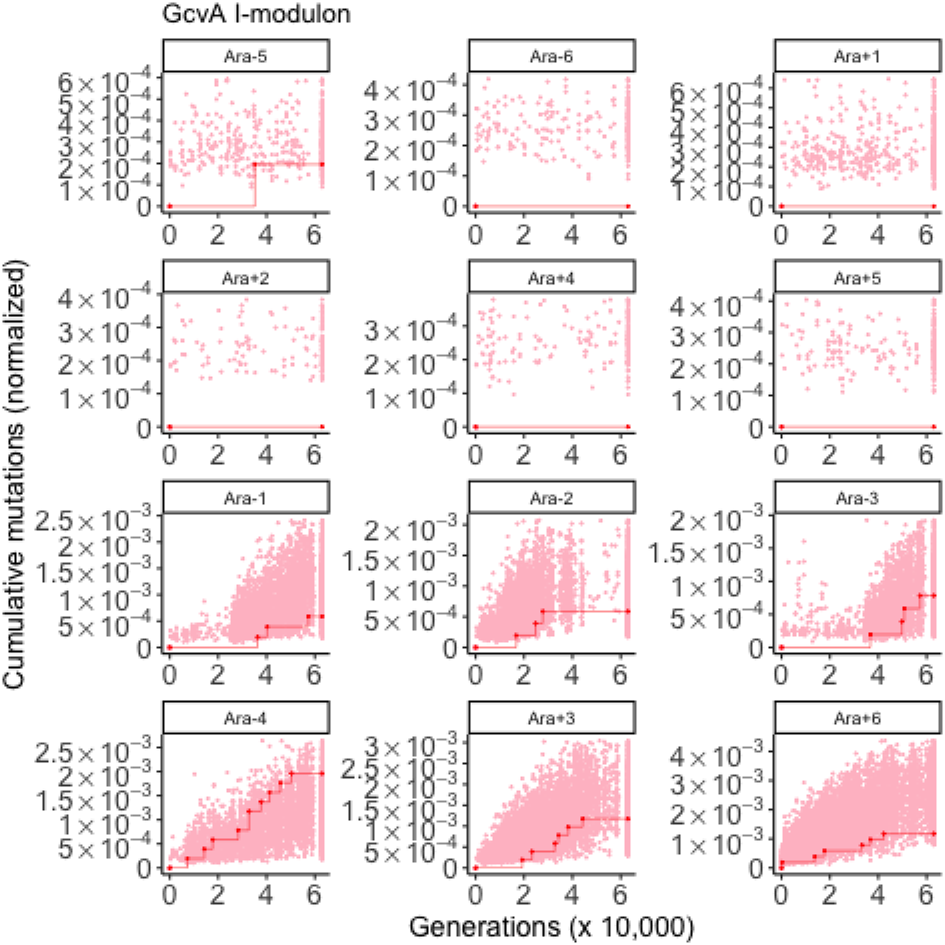

GcvA I-modulon
Ara-5
Ara-6
Ara+1
Ara+2
Ara+4
Ara+5
Ara-1
Ara-2
Ara-3
Ara-4
Ara+3
Ara+6
Cumulative mutations (normalized)
Generations (x 10,000)

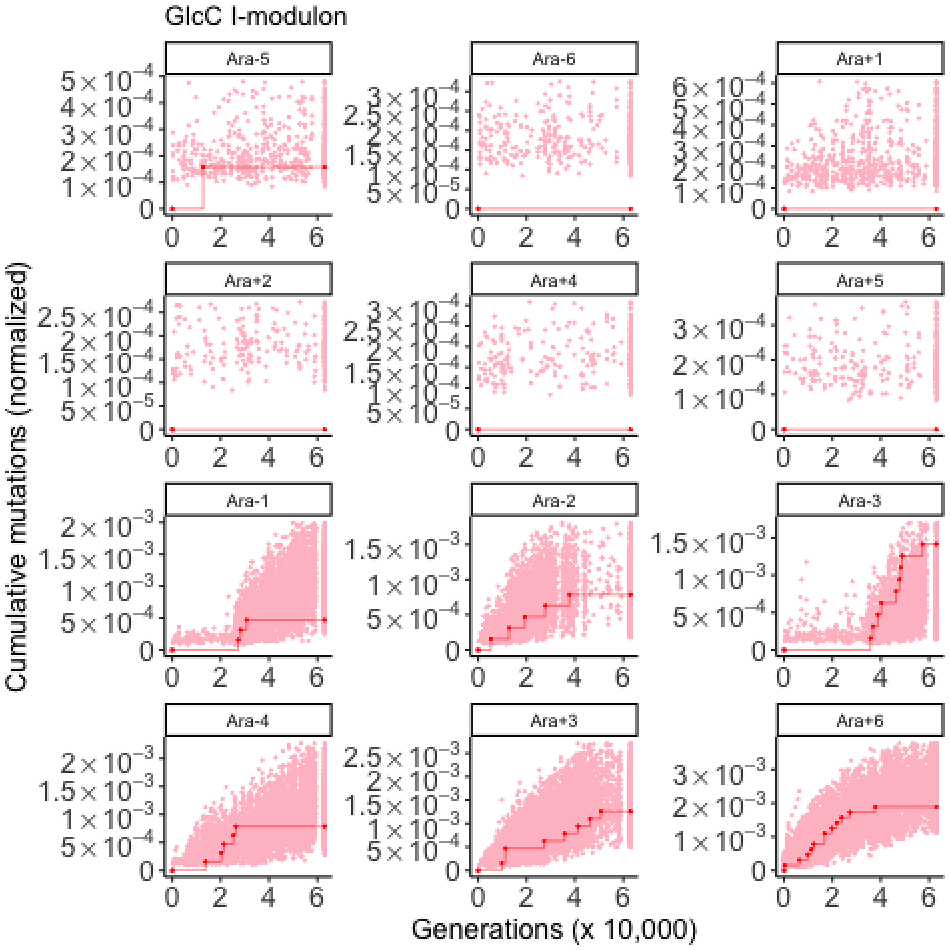
GlcC I-modulon
Ara-5
Ara-6
Ara+1
Ara+2
Ara+4
Ara+5
Ara-1
Ara-2
Ara-3
Ara-4
Ara+3
Ara+6
Cumulative mutations (normalized)
Generations (x 10,000)

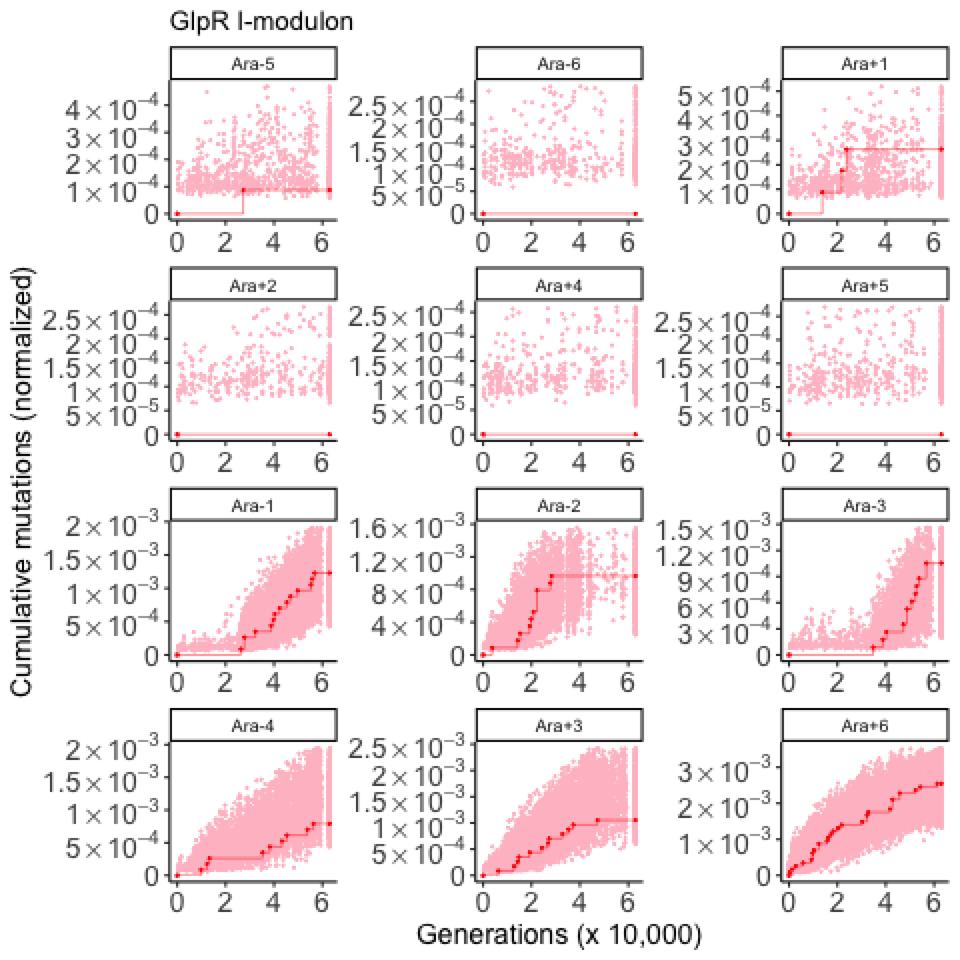

GlpR I-modulon
Ara-5
Ara-6
Ara+1
Ara+2
Ara+4
Ara+5
Ara-1
Ara-2
Ara-3
Ara-4
Ara+3
Ara+6
Cumulative mutations (normalized)
Generations (x 10,000)

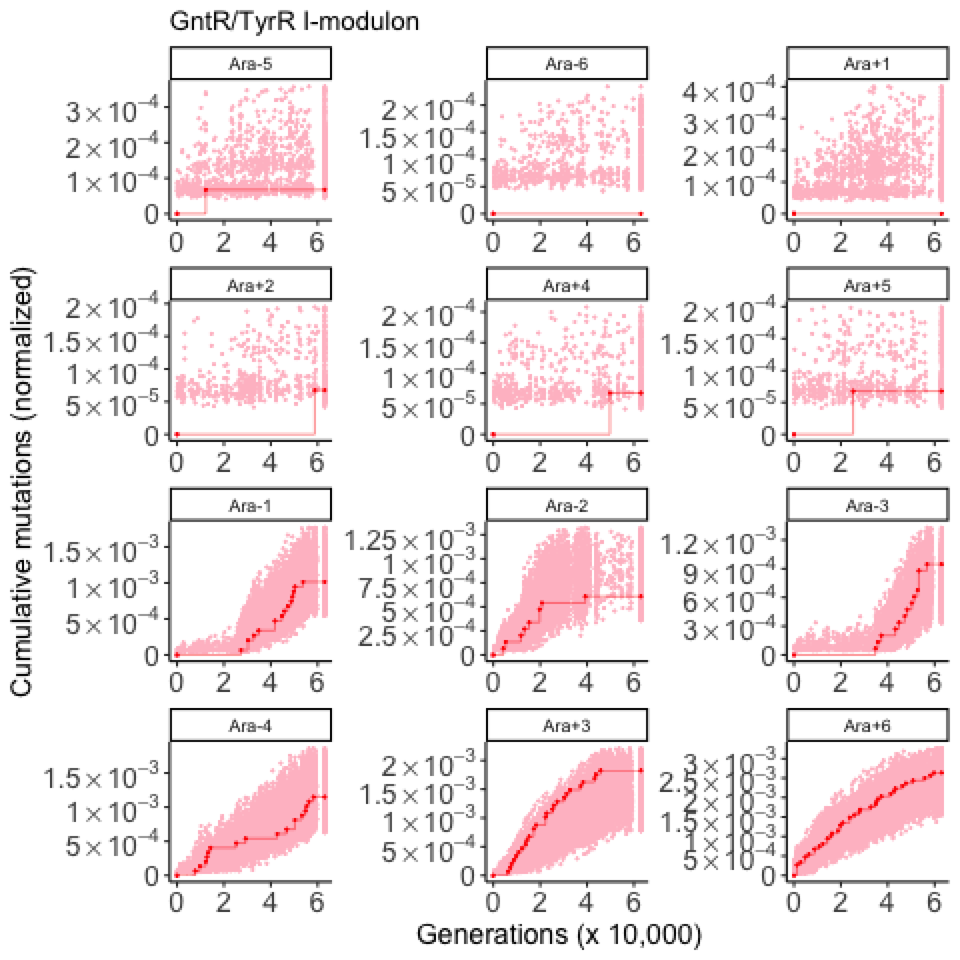

GntR/TyrR I-modulon
Ara-5
Ara-6
Ara+1
Ara+2
Ara+4
Ara+5
Ara-1
Ara-2
Ara-3
Ara-4
Ara+3
Ara+6
Cumulative mutations (normalized)
Generations (x 10,000)

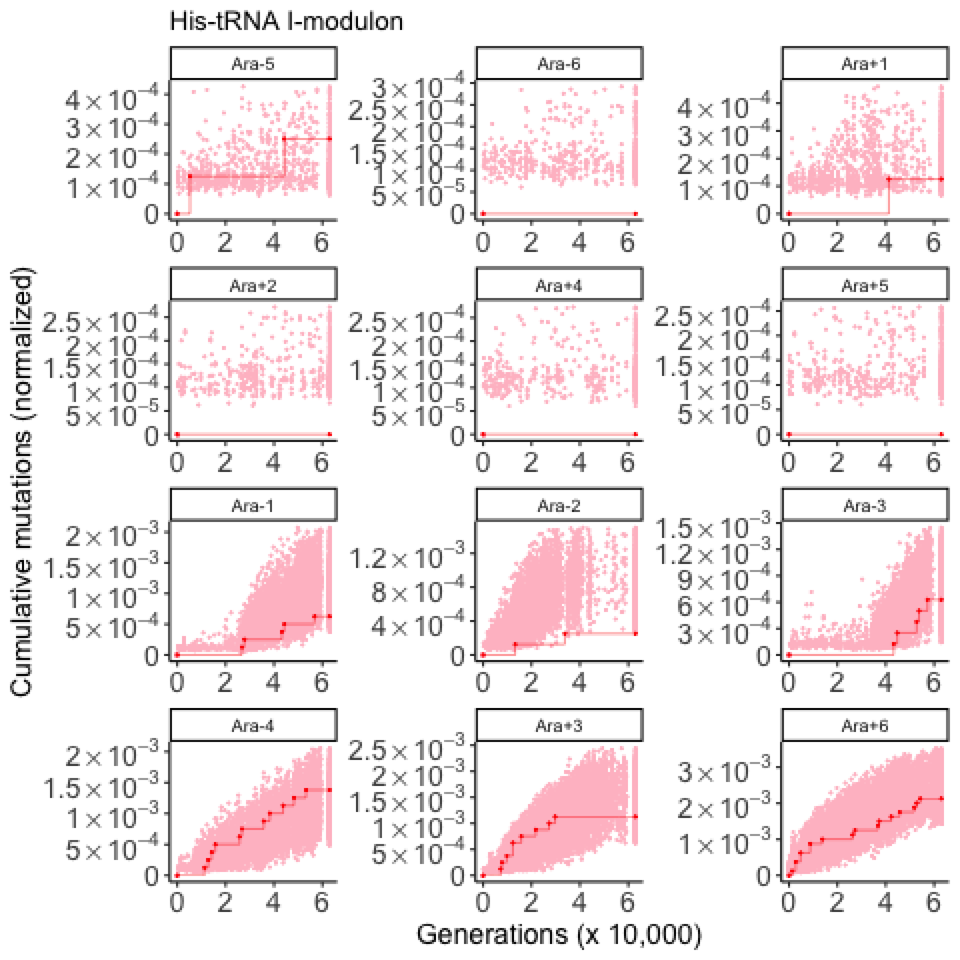
His-tRNA I-modulon
Ara-5
Ara-6
Ara+1
Ara+2
Ara+4
Ara+5
Ara-1
Ara-2
Ara-3
Ara-4
Ara+3
Ara+6
Cumulative mutations (normalized)
Generations (x 10,000)

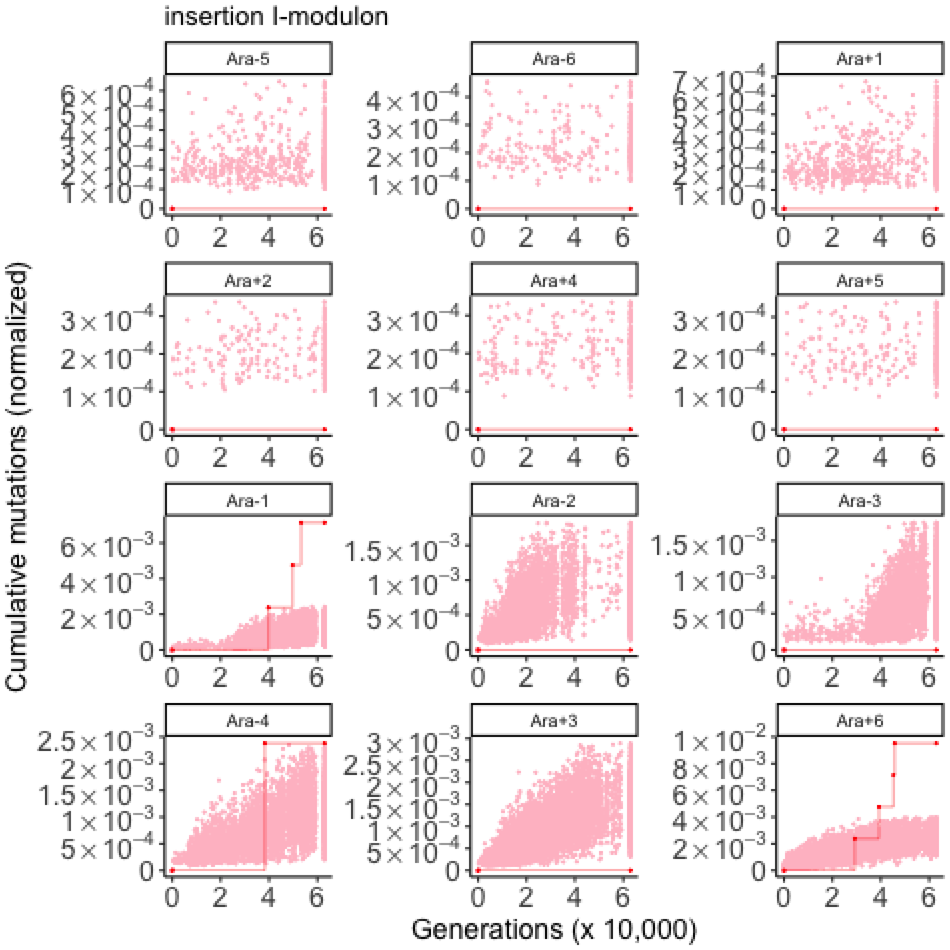
insertion I-modulon
Ara-5
Ara-6
Ara+1
Ara+2
Ara+4
Ara+5
Ara-1
Ara-2
Ara-3
Ara-4
Ara+3
Ara+6
Cumulative mutations (normalized)
Generations (x 10,000)

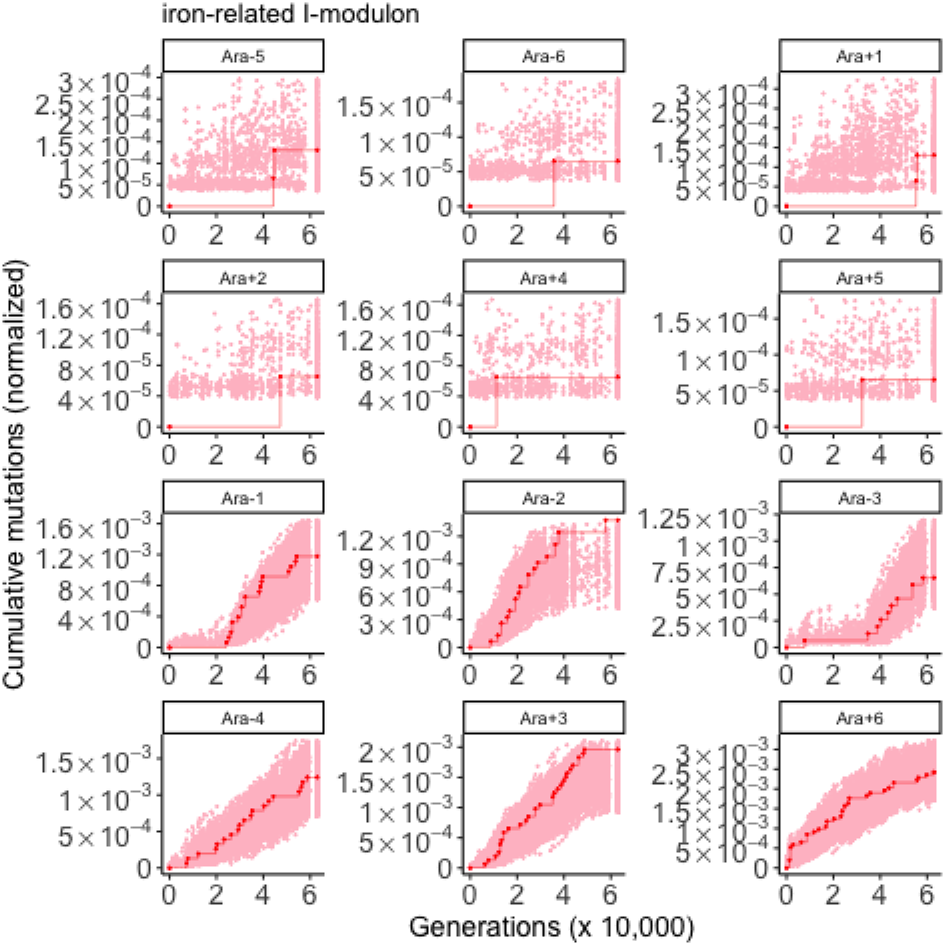

iron-related I-modulon
Ara-5
Ara-6
Ara+1
Ara+2
Ara+4
Ara+5
Ara-1
Ara-2
Ara-3
Ara-4
Ara+3
Ara+6
Cumulative mutations (normalized)
Generations (x 10,000)

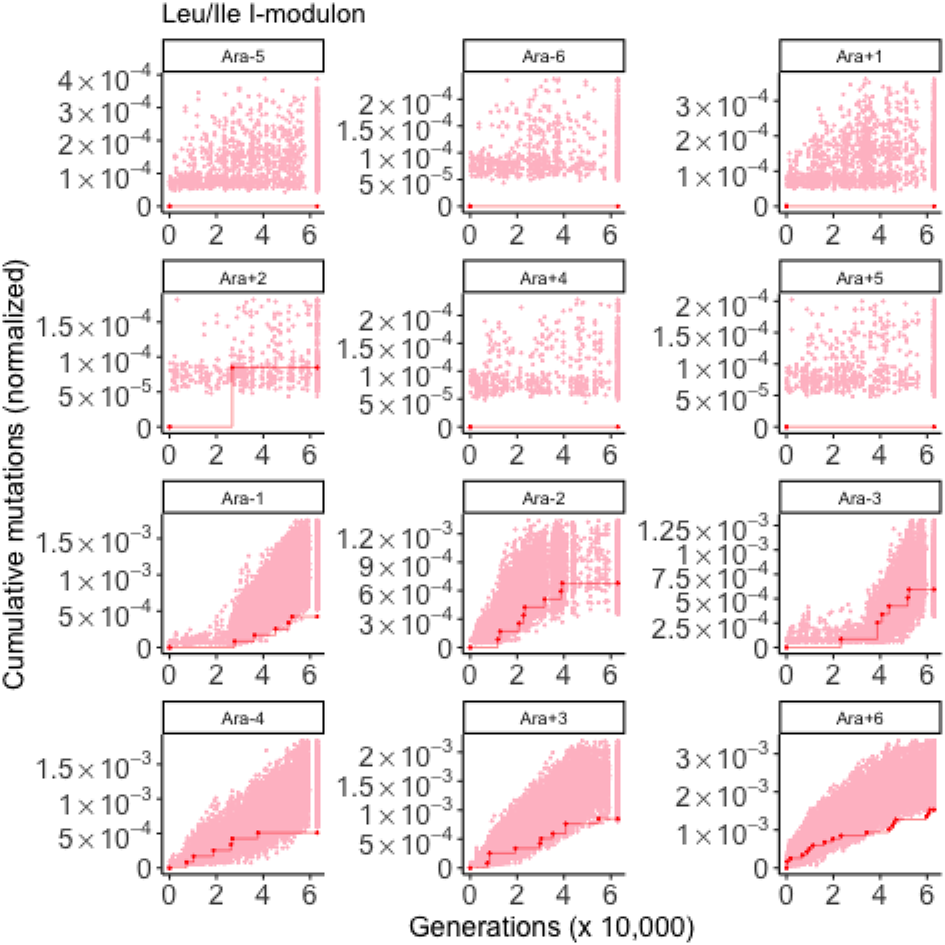

Leu/Ile I-modulon
Ara-5
Ara-6
Ara+1
Ara+2
Ara+4
Ara+5
Ara-1
Ara-2
Ara-3
Ara-4
Ara+3
Ara+6
Cumulative mutations (normalized)
Generations (x 10,000)

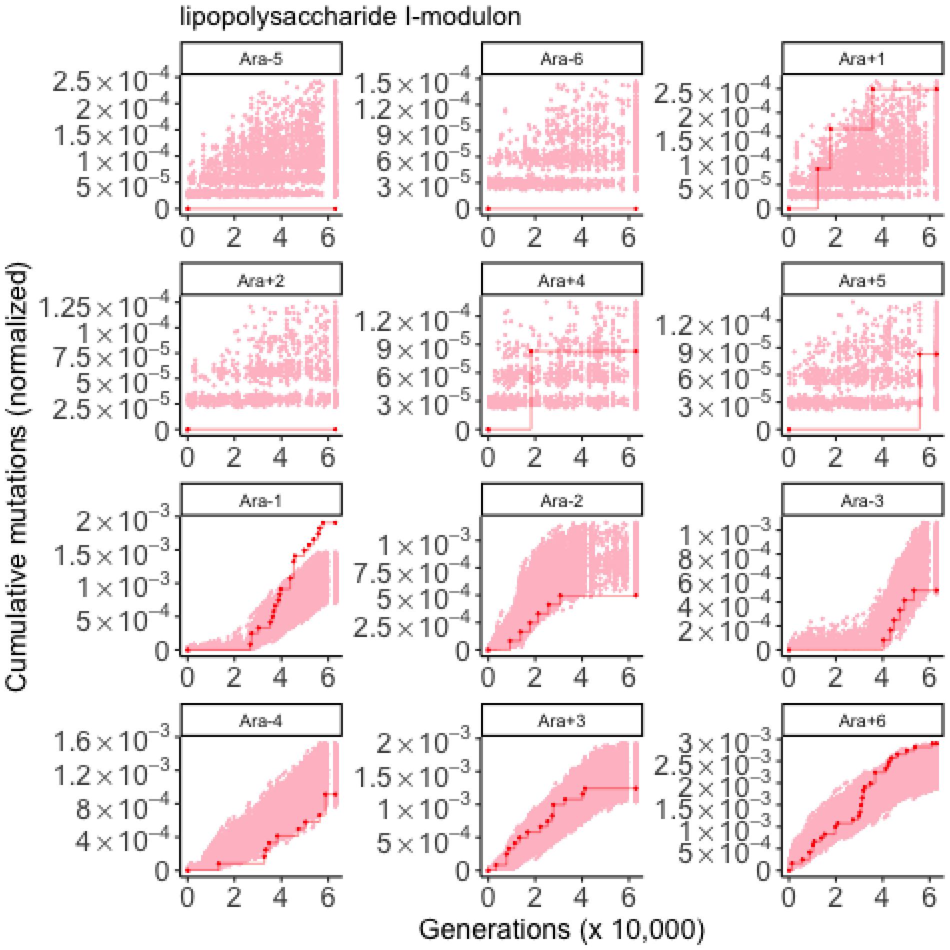
lipopolysaccharide I-modulon
Ara-5
Ara-6
Ara+1
Ara+2
Ara+4
Ara+5
Ara-1
Ara-2
Ara-3
Ara-4
Ara+3
Ara+6
Cumulative mutations (normalized)
Generations (x 10,000)

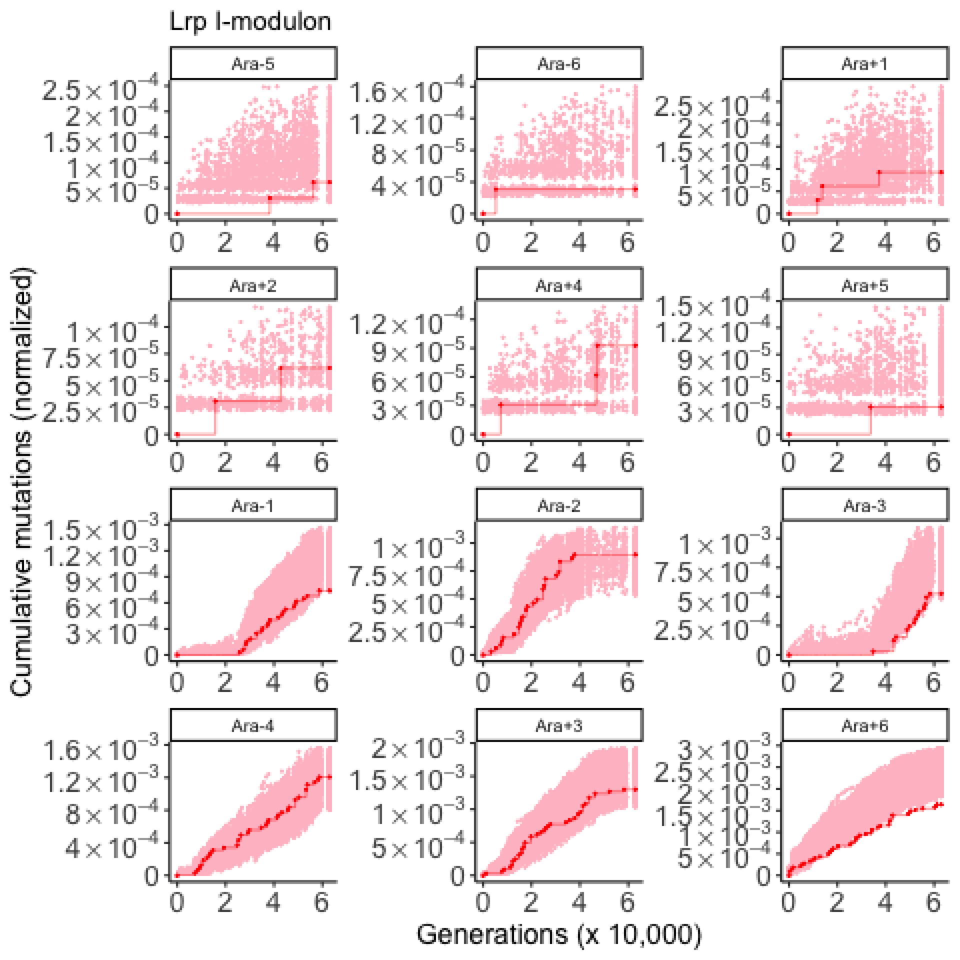

Lrp I-modulon
Ara-5
Ara-6
Ara+1
Ara+2
Ara+4
Ara+5
Ara-1
Ara-2
Ara-3
Ara-4
Ara+3
Ara+6
Cumulative mutations (normalized)
Generations (x 10,000)

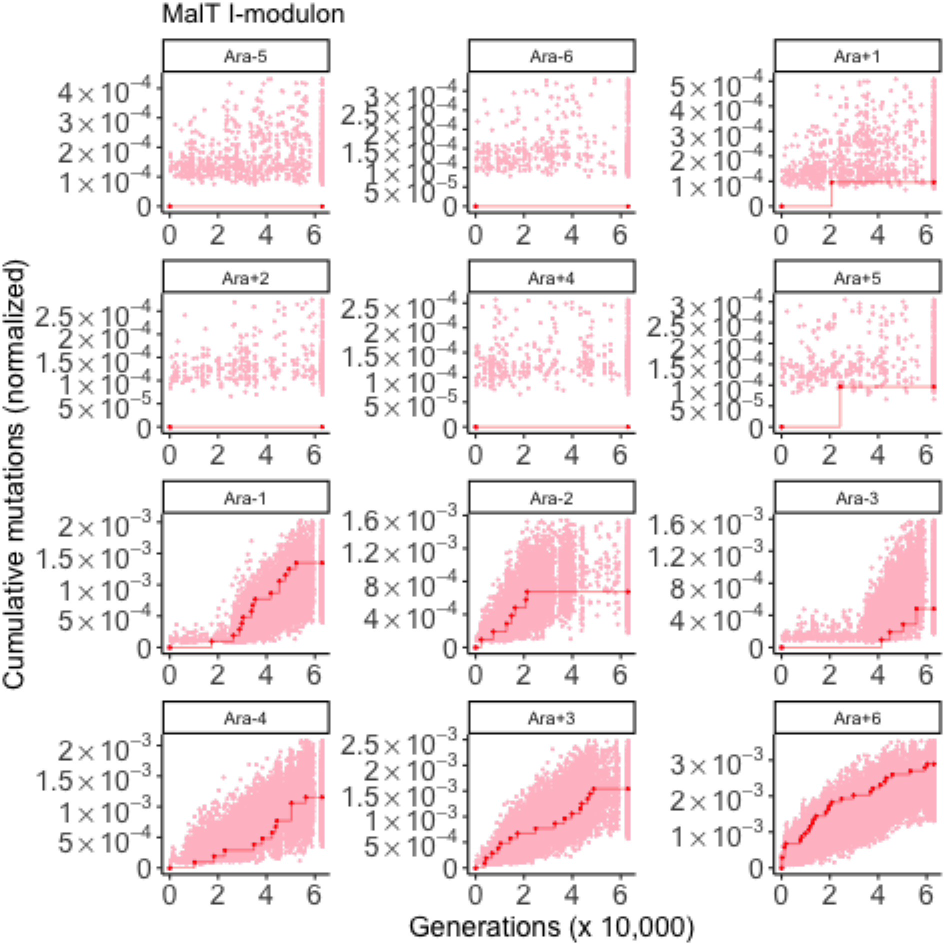

MalT I-modulon
Ara-5
Ara-6
Ara+1
Ara+2
Ara+4
Ara+5
Ara-1
Ara-2
Ara-3
Ara-4
Ara+3
Ara+6
Cumulative mutations (normalized)
Generations (x 10,000)

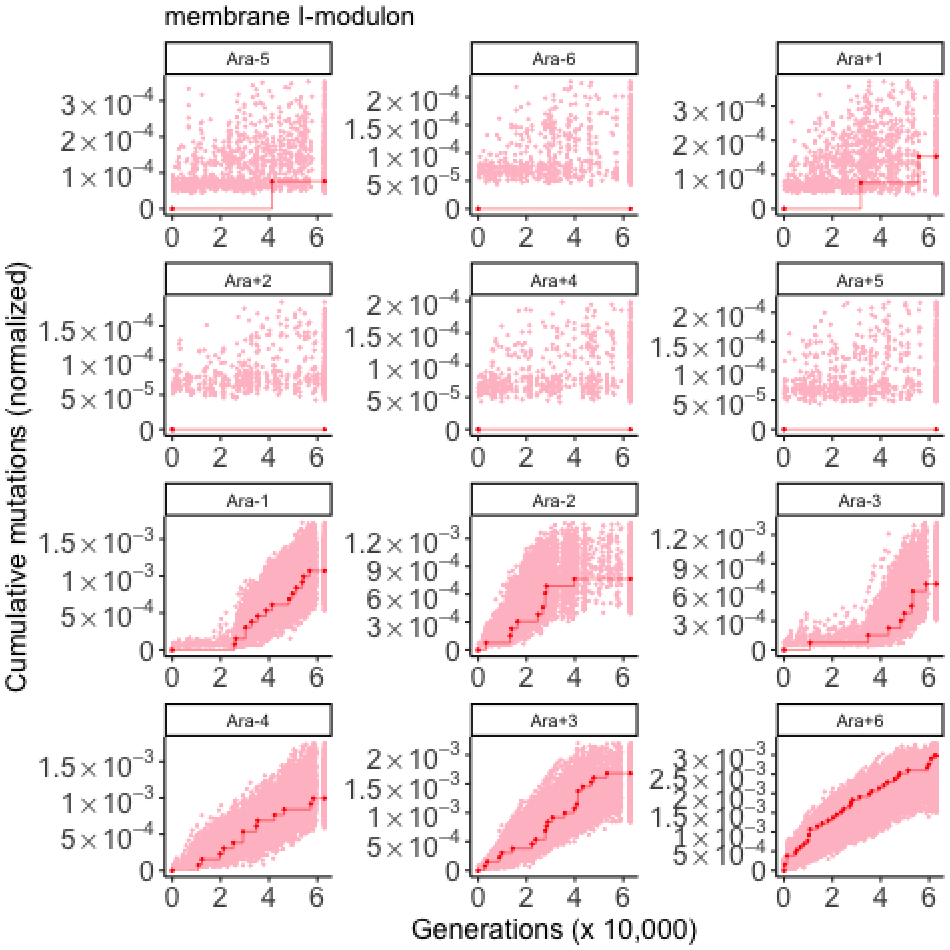

membrane I-modulon
Ara-5
Ara-6
Ara+1
Ara+2
Ara+4
Ara+5
Ara-1
Ara-2
Ara-3
Ara-4
Ara+3
Ara+6
Cumulative mutations (normalized)
Generations (x 10,000)

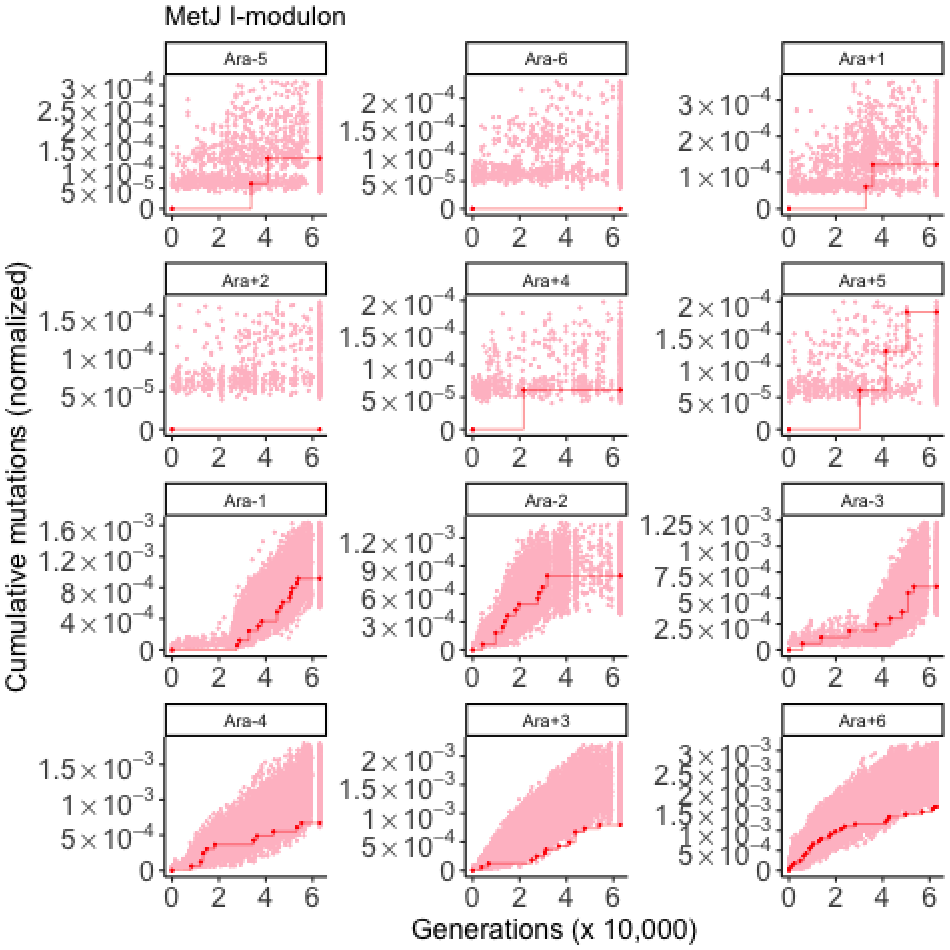
MetJ I-modulon
Ara-5
Ara-6
Ara+1
Ara+2
Ara+4
Ara+5
Ara-1
Ara-2
Ara-3
Ara-4
Ara+3
Ara+6
Cumulative mutations (normalized)
Generations (x 10,000)

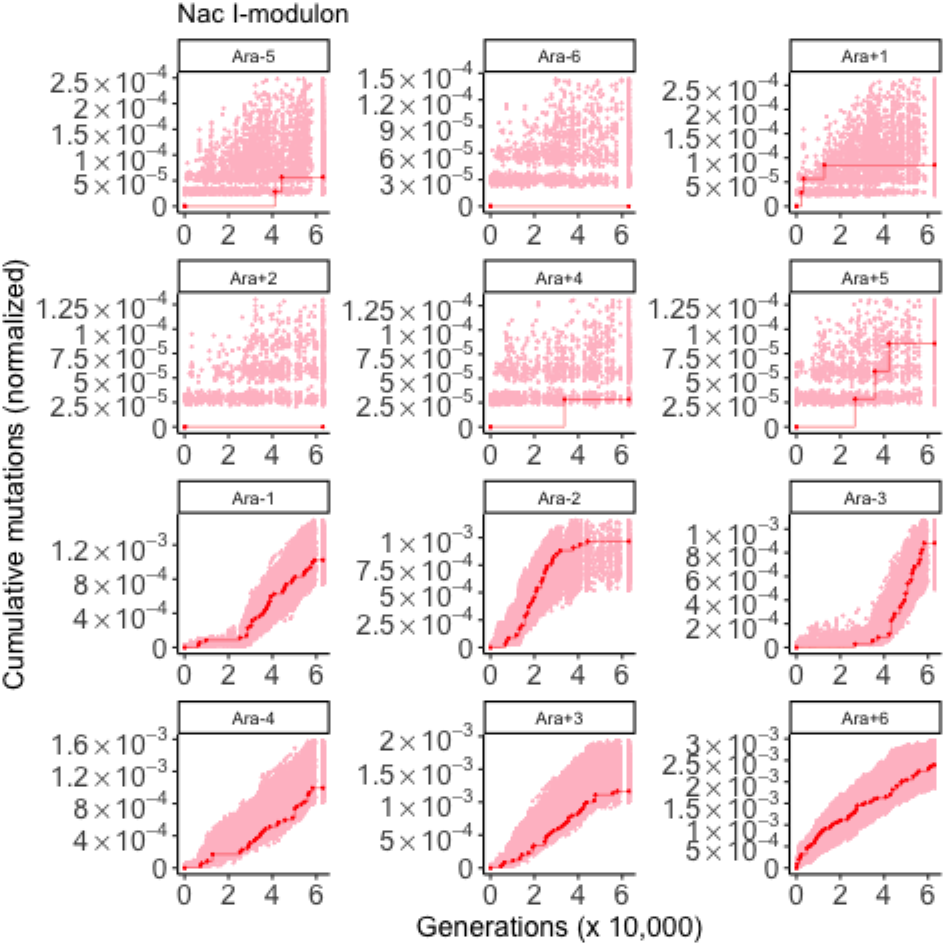

Nac I-modulon
Ara-5
Ara-6
Ara+1
Ara+2
Ara+4
Ara+5
Ara-1
Ara-2
Ara-3
Ara-4
Ara+3
Ara+6
Cumulative mutations (normalized)
Generations (x 10,000)

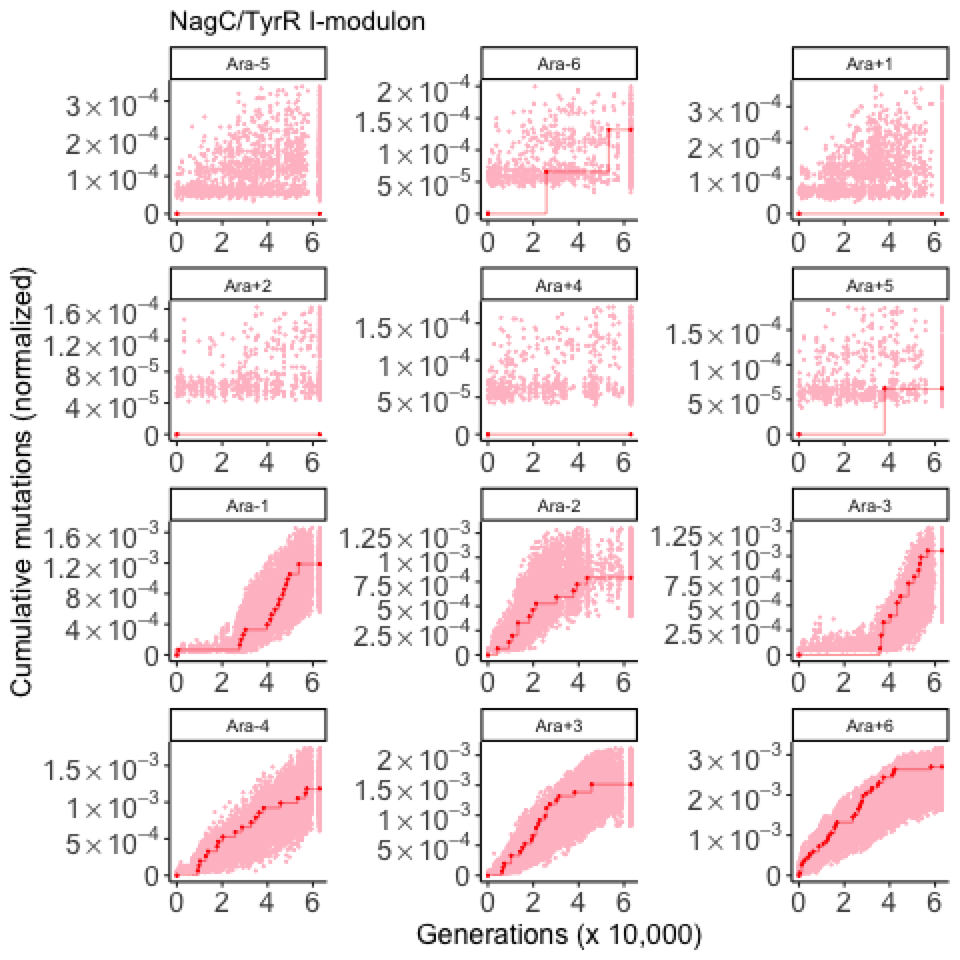
NagC/TyrR I-modulon
Ara-5
Ara-6
Ara+1
Ara+2
Ara+4
Ara+5
Ara-1
Ara-2
Ara-3
Ara-4
Ara+3
Ara+6
Cumulative mutations (normalized)
Generations (x 10,000)

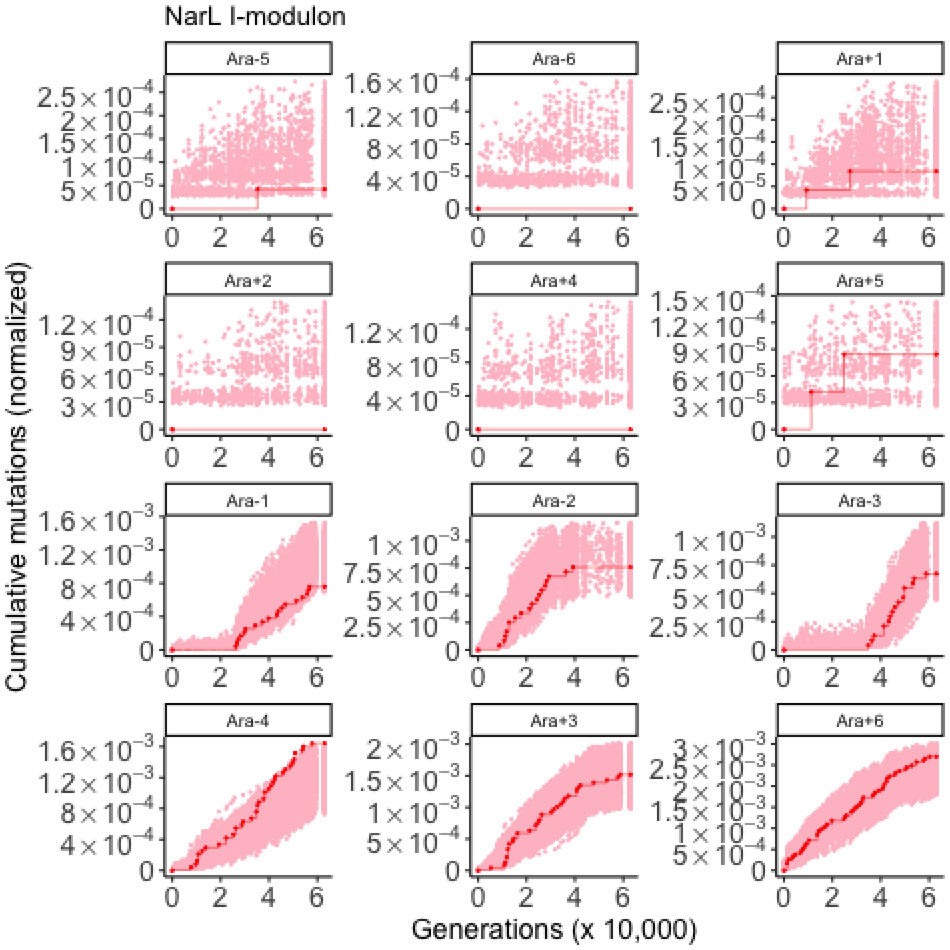

NarL I-modulon
Ara-5
Ara-6
Ara+1
Ara+2
Ara+4
Ara+5
Ara-1
Ara-2
Ara-3
Ara-4
Ara+3
Ara+6
Cumulative mutations (normalized)
Generations (x 10,000)

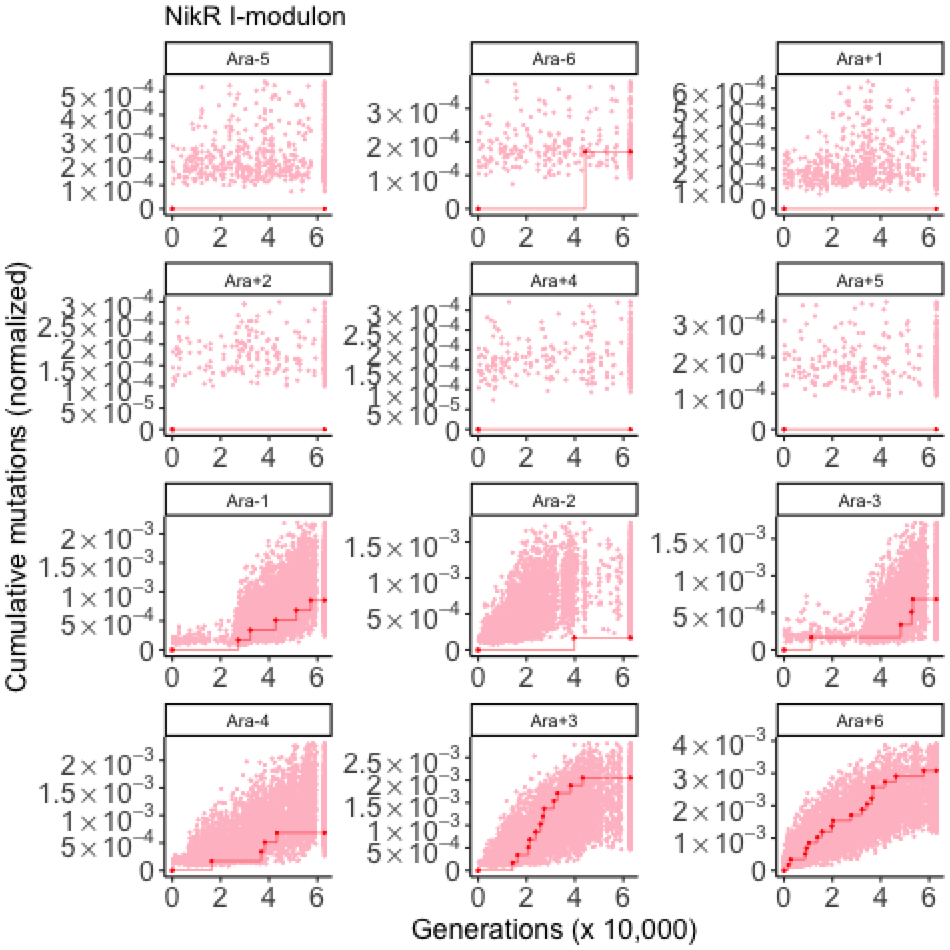
NikR I-modulon
Ara-5
Ara-6
Ara+1
Ara+2
Ara+4
Ara+5
Ara-1
Ara-2
Ara-3
Ara-4
Ara+3
Ara+6
Cumulative mutations (normalized)
Generations (x 10,000)

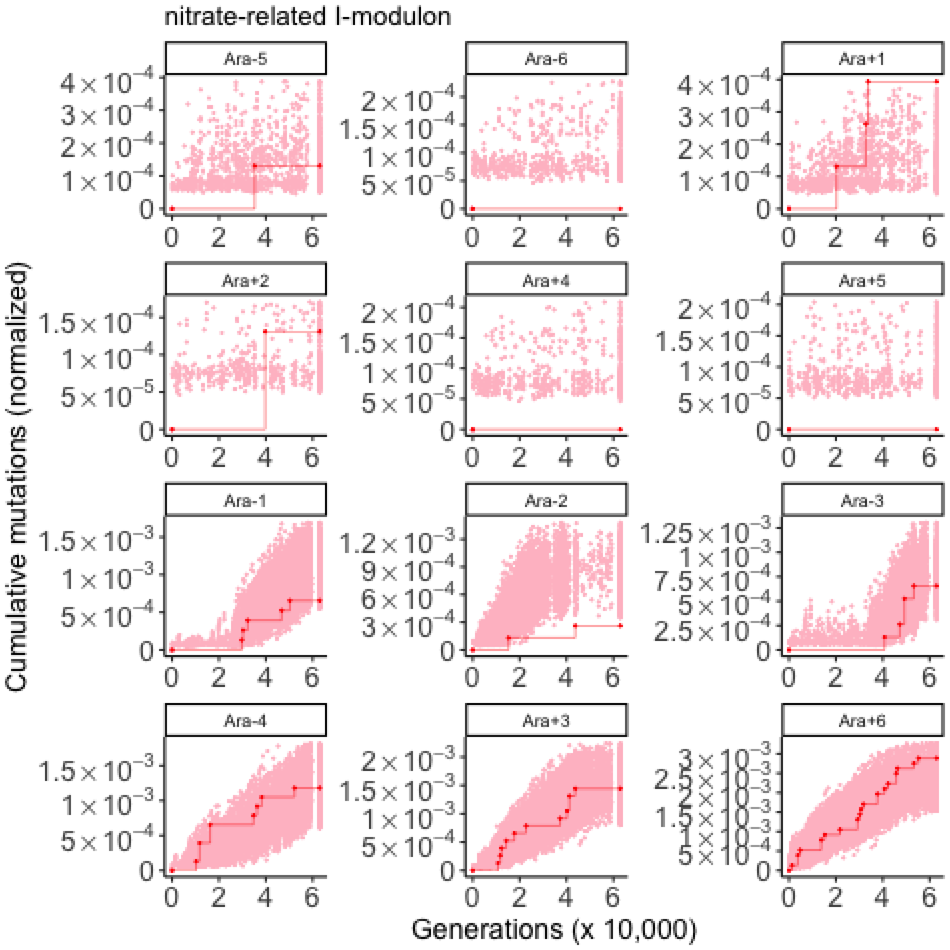
nitrate-related I-modulon
Ara-5
Ara-6
Ara+1
Ara+2
Ara+4
Ara+5
Ara-1
Ara-2
Ara-3
Ara-4
Ara+3
Ara+6
Cumulative mutations (normalized)
Generations (x 10,000)

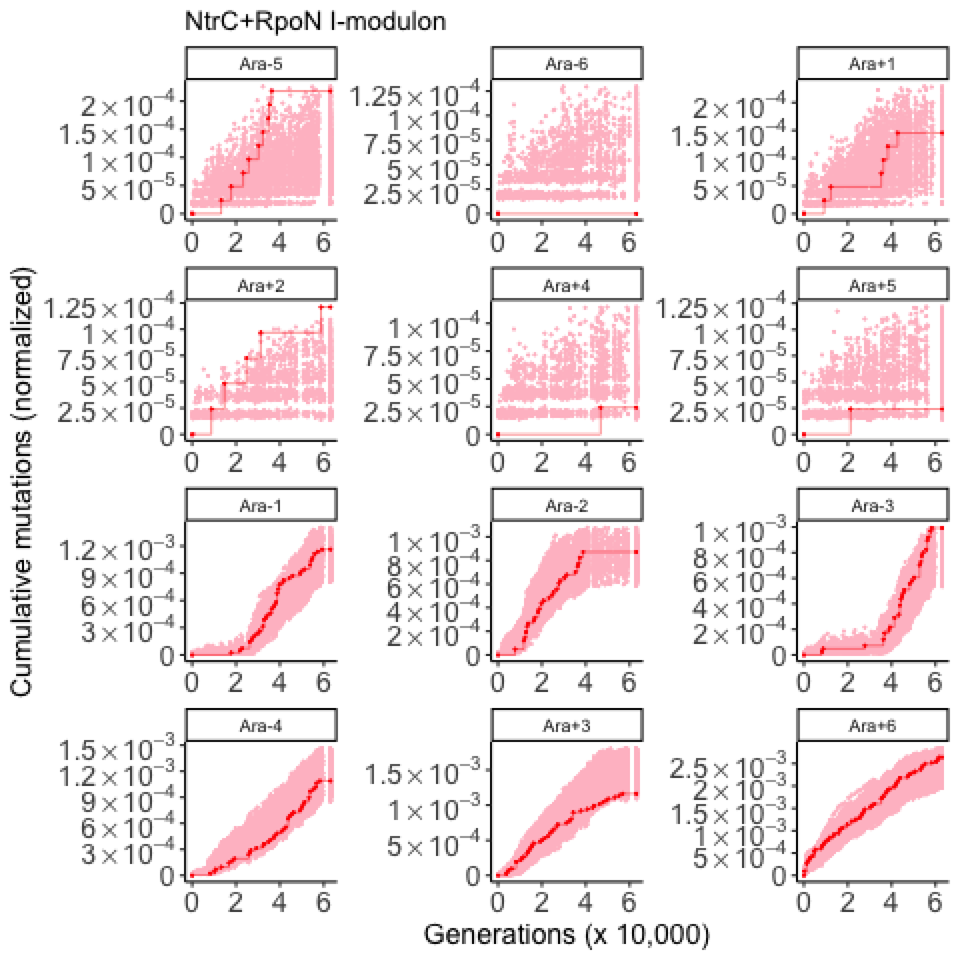
NtrC+RpoN I-modulon
Ara-5
Ara-6
Ara+1
Ara+2
Ara+4
Ara+5
Ara-1
Ara-2
Ara-3
Ara-4
Ara+3
Ara+6
Cumulative mutations (normalized)
Generations (x 10,000)

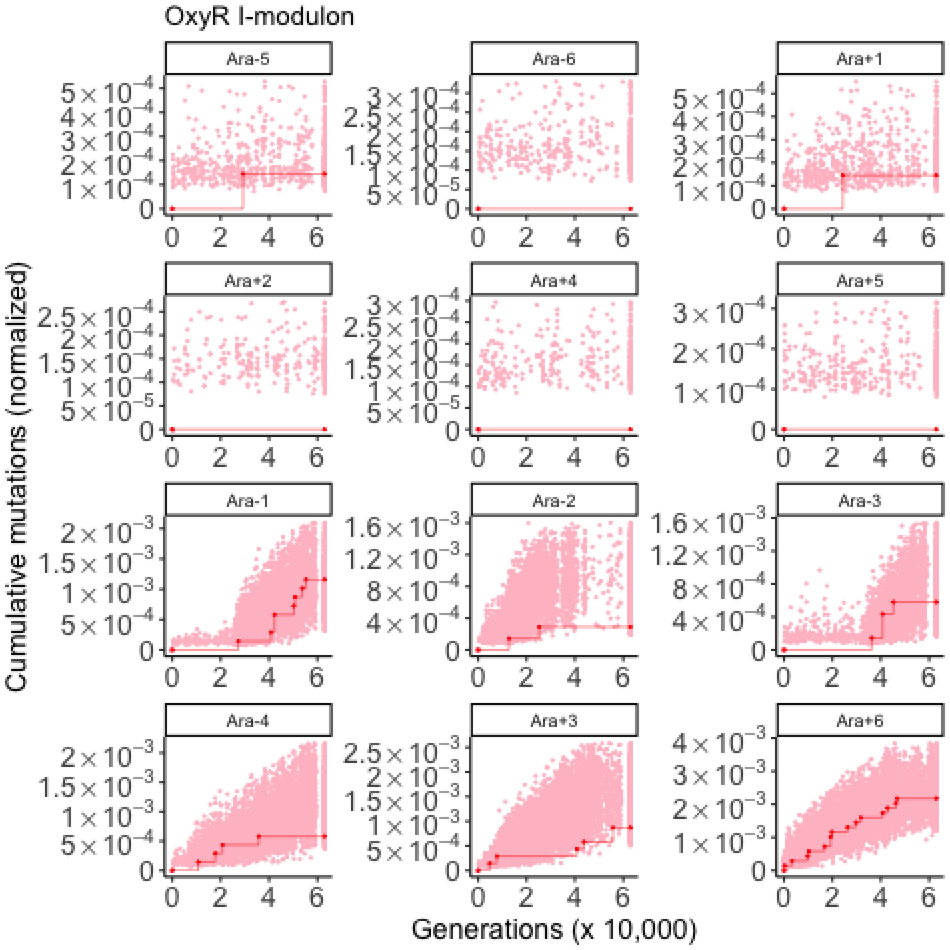
OxyR I-modulon
Ara-5
Ara-6
Ara+1
Ara+2
Ara+4
Ara+5
Ara-1
Ara-2
Ara-3
Ara-4
Ara+3
Ara+6
Cumulative mutations (normalized)
Generations (x 10,000)

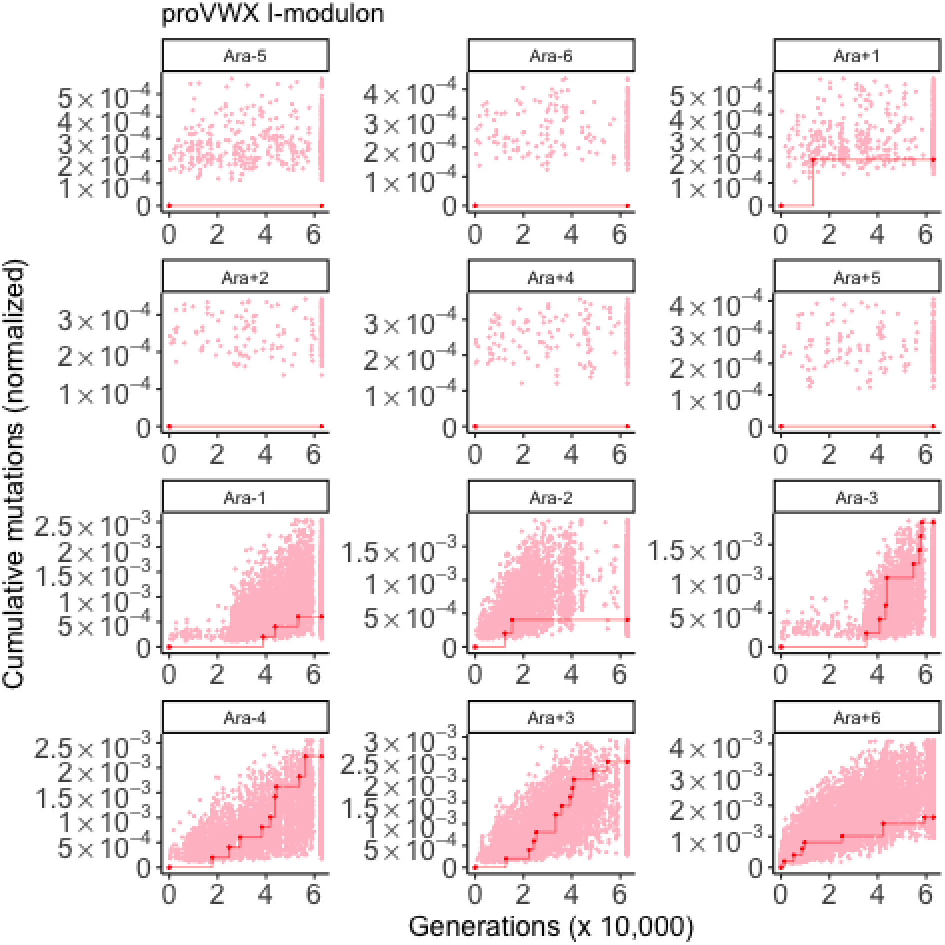
proVWX I-modulon
Ara-5
Ara-6
Ara+1
Ara+2
Ara+4
Ara+5
Ara-1
Ara-2
Ara-3
Ara-4
Ara+3
Ara+6
Cumulative mutations (normalized)
Generations (x 10,000)

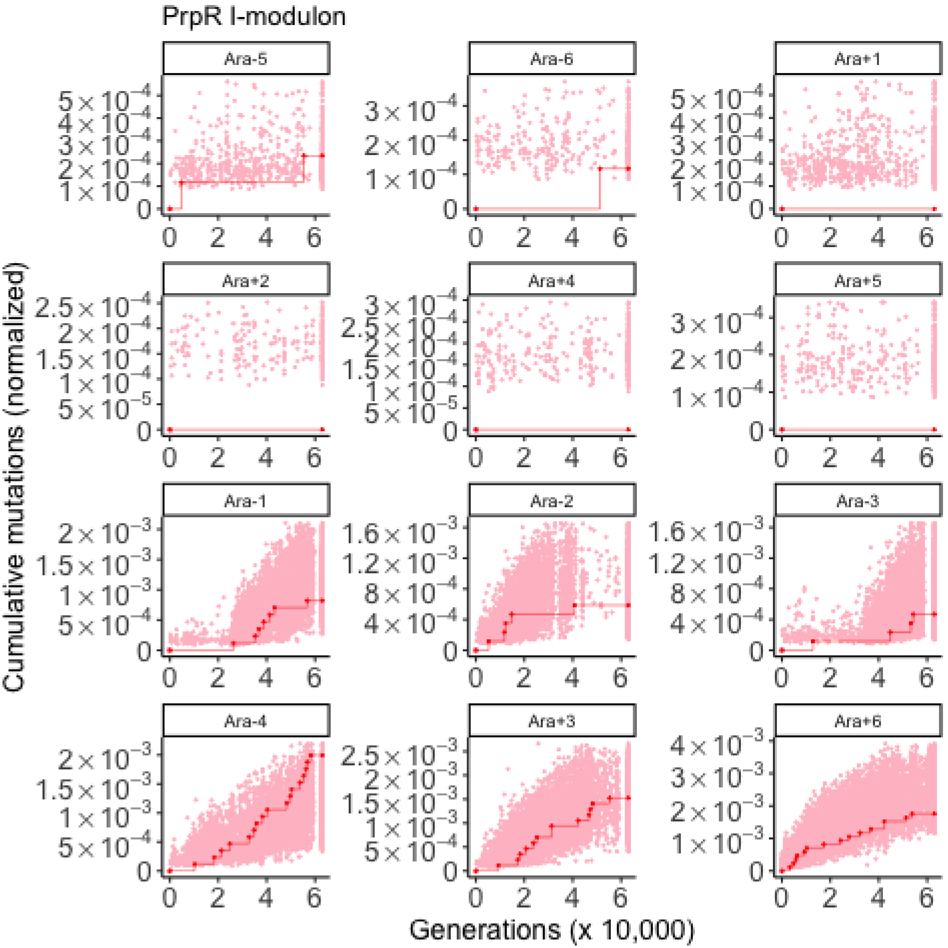

PrpR I-modulon
Ara-5
Ara-6
Ara+1
Ara+2
Ara+4
Ara+5
Ara-1
Ara-2
Ara-3
Ara-4
Ara+3
Ara+6
Cumulative mutations (normalized)
Generations (x 10,000)

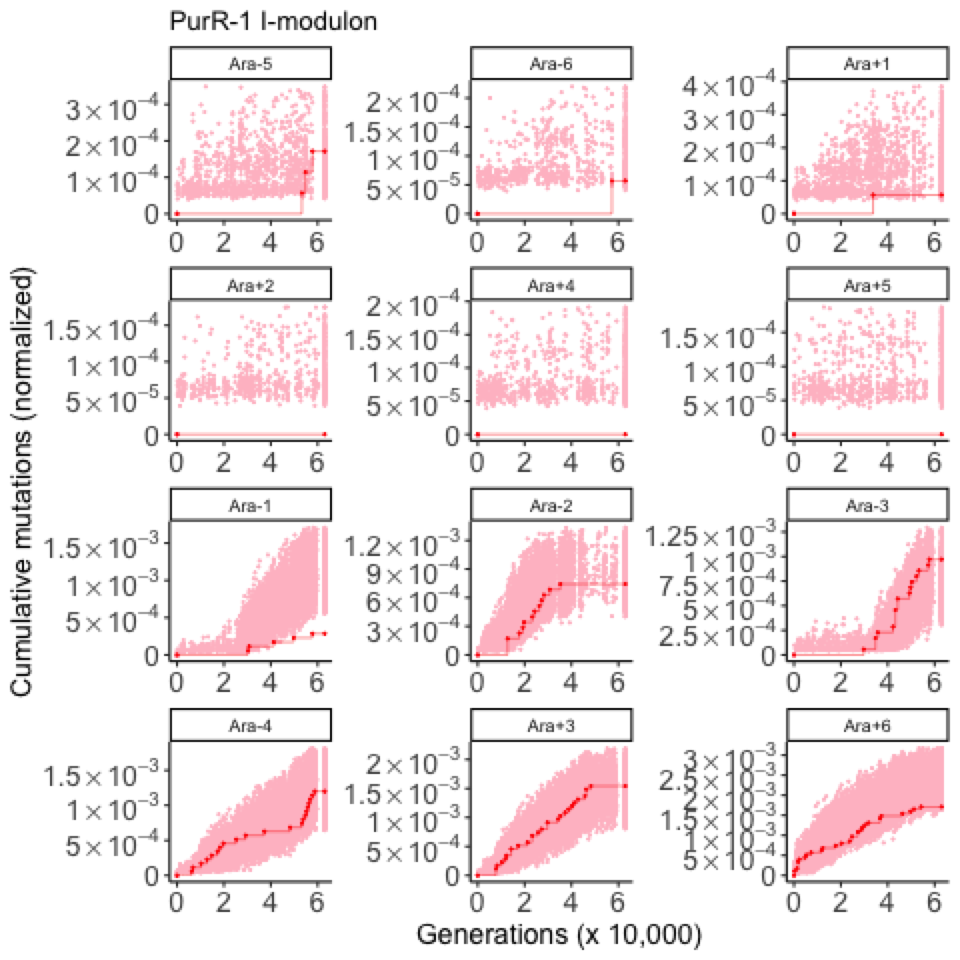
PurR-1 I-modulon
Ara-5
Ara-6
Ara+1
Ara+2
Ara+4
Ara+5
Ara-1
Ara-2
Ara-3
Ara-4
Ara+3
Ara+6
Cumulative mutations (normalized)
Generations (x 10,000)

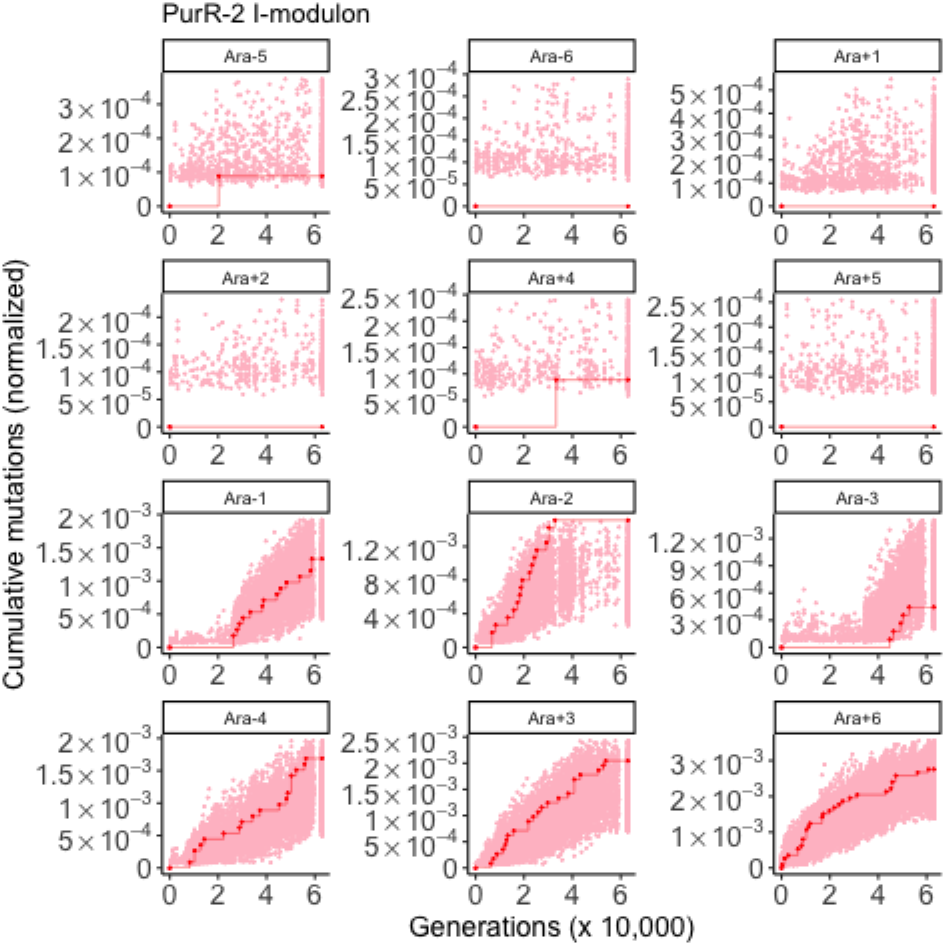

PurR-2 I-modulon
Ara-5
Ara-6
Ara+1
Ara+2
Ara+4
Ara+5
Ara-1
Ara-2
Ara-3
Ara-4
Ara+3
Ara+6
Cumulative mutations (normalized)
Generations (x 10,000)

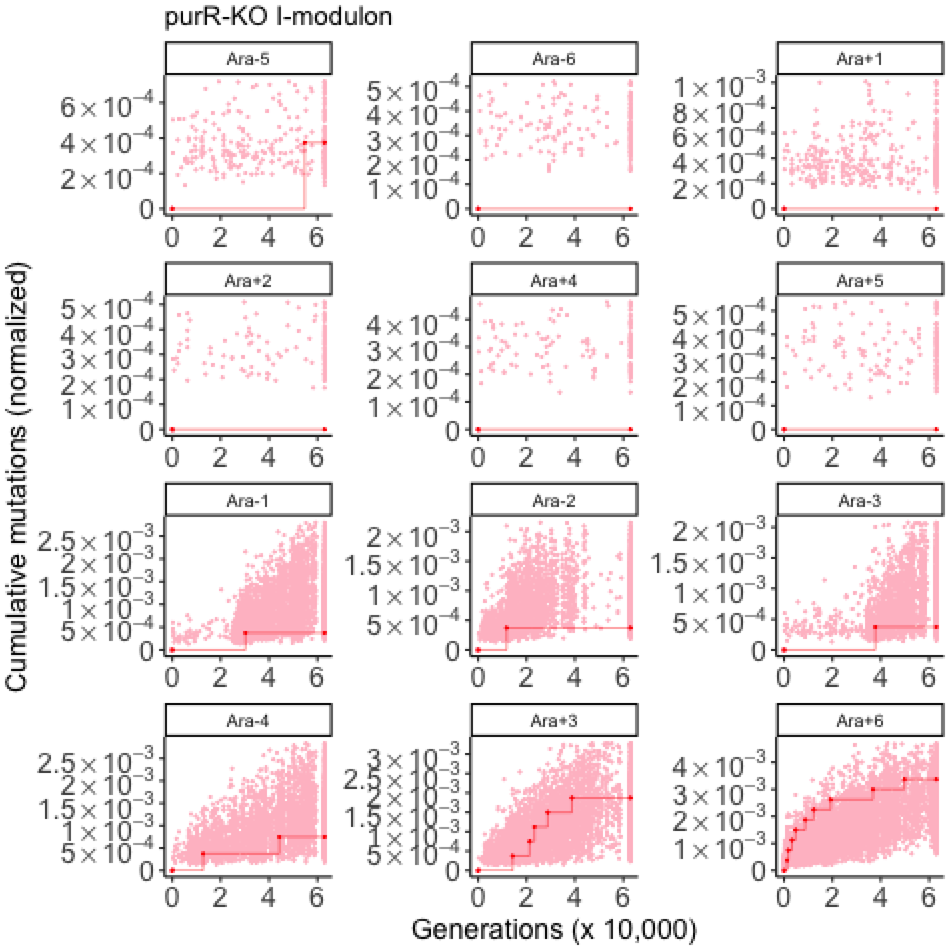

purR-KO I-modulon
Ara-5
Ara-6
Ara+1
Ara+2
Ara+4
Ara+5
Ara-1
Ara-2
Ara-3
Ara-4
Ara+3
Ara+6
Cumulative mutations (normalized)
Generations (x 10,000)

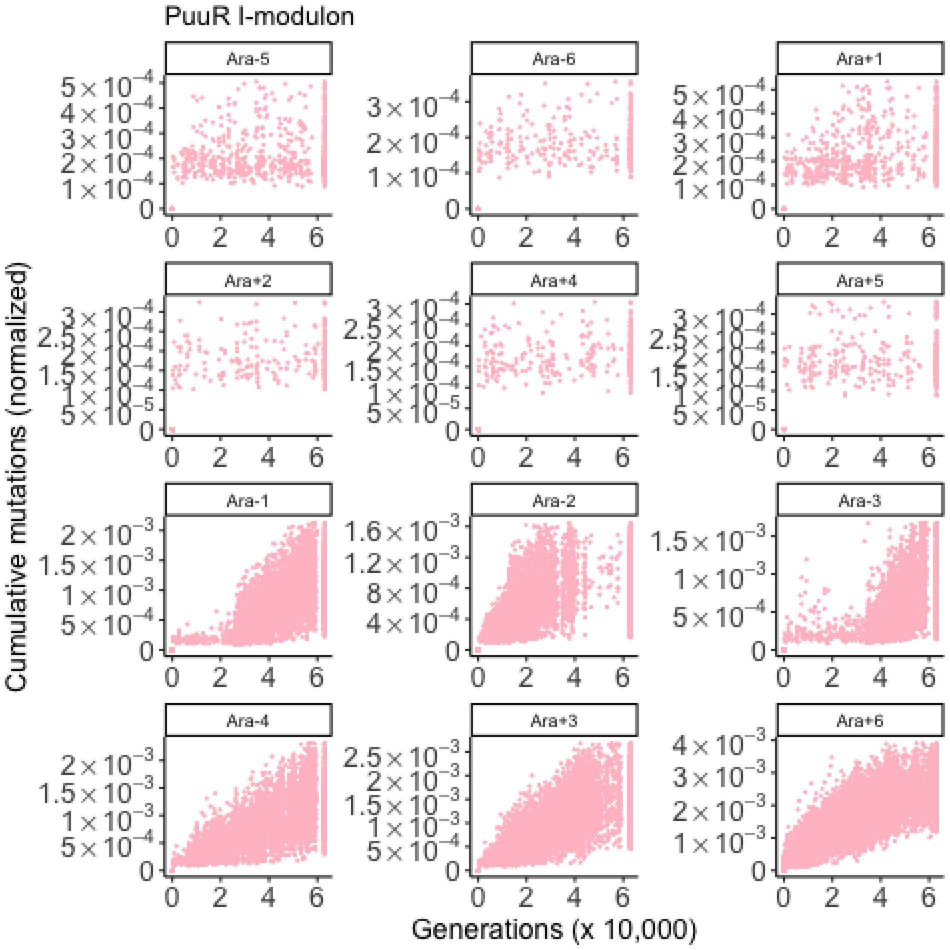
PuuR I-modulon
Ara-5
Ara-6
Ara+1
Ara+2
Ara+4
Ara+5
Ara-1
Ara-2
Ara-3
Ara-4
Ara+3
Ara+6
Cumulative mutations (normalized)
Generations (x 10,000)

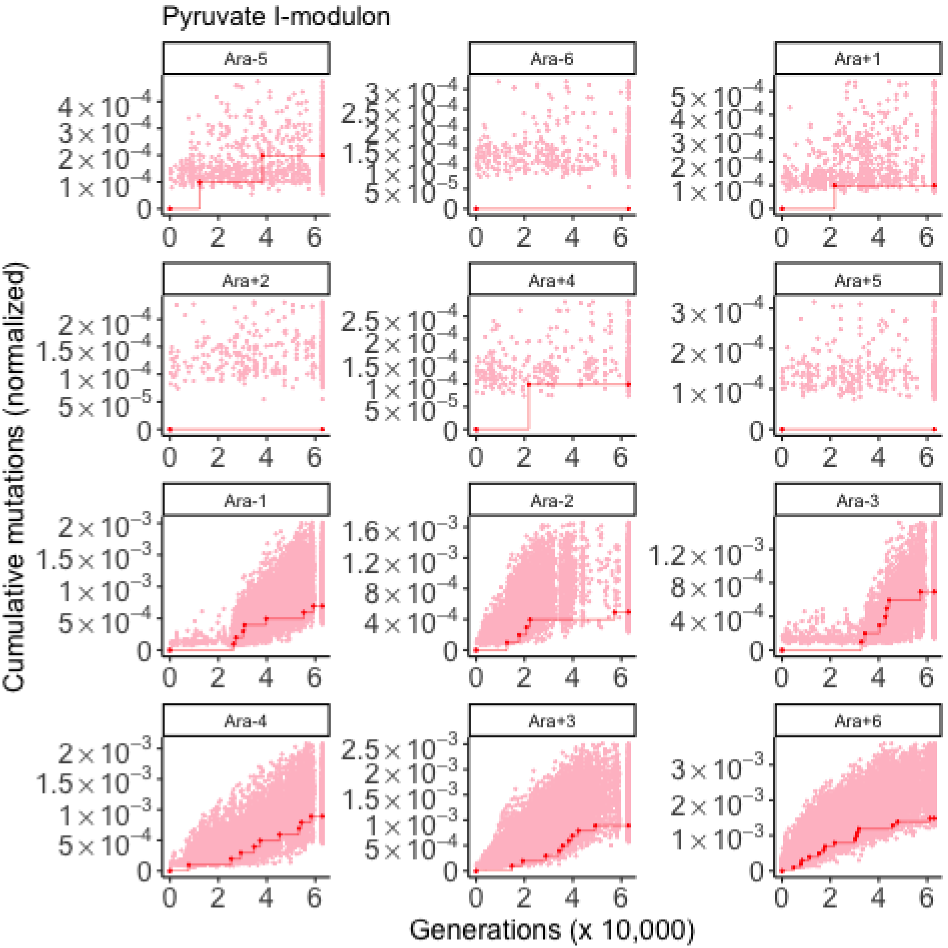

Pyruvate I-modulon
Ara-5
Ara-6
Ara+1
Ara+2
Ara+4
Ara+5
Ara-1
Ara-2
Ara-3
Ara-4
Ara+3
Ara+6
Cumulative mutations (normalized)
Generations (x 10,000)

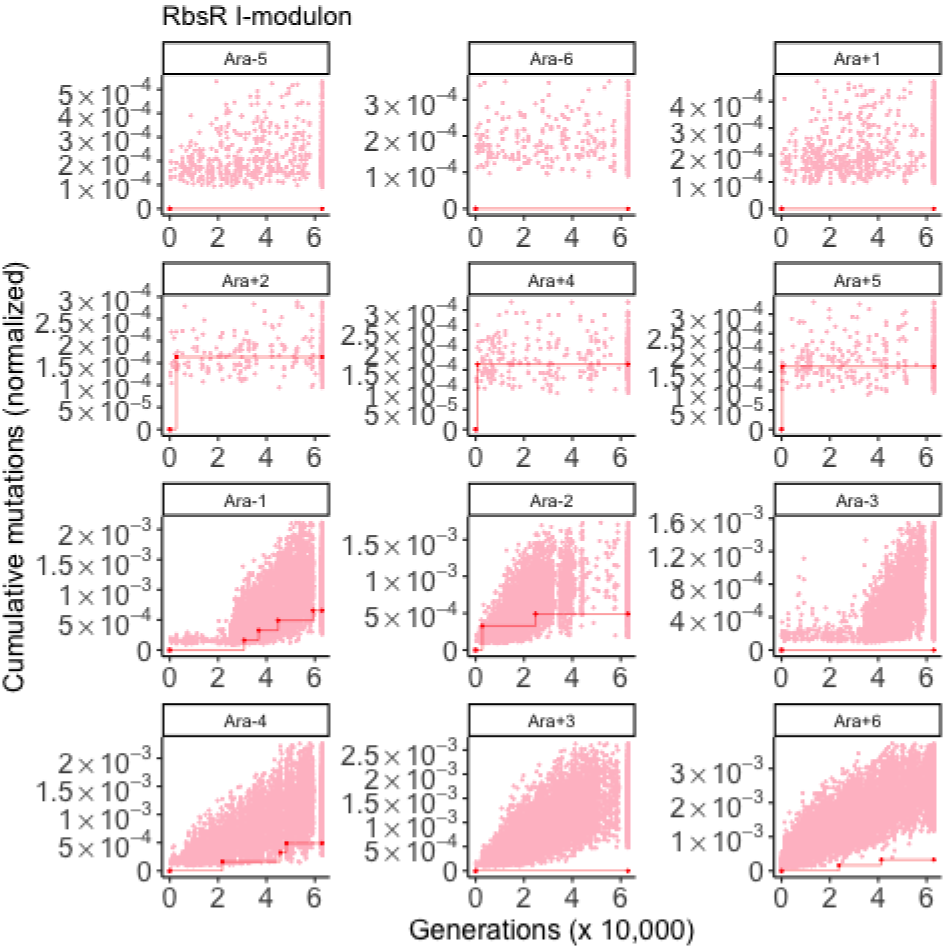

RbsR I-modulon
Ara-5
Ara-6
Ara+1
Ara+2
Ara+4
Ara+5
Ara-1
Ara-2
Ara-3
Ara-4
Ara+3
Ara+6
Cumulative mutations (normalized)
Generations (x 10,000)

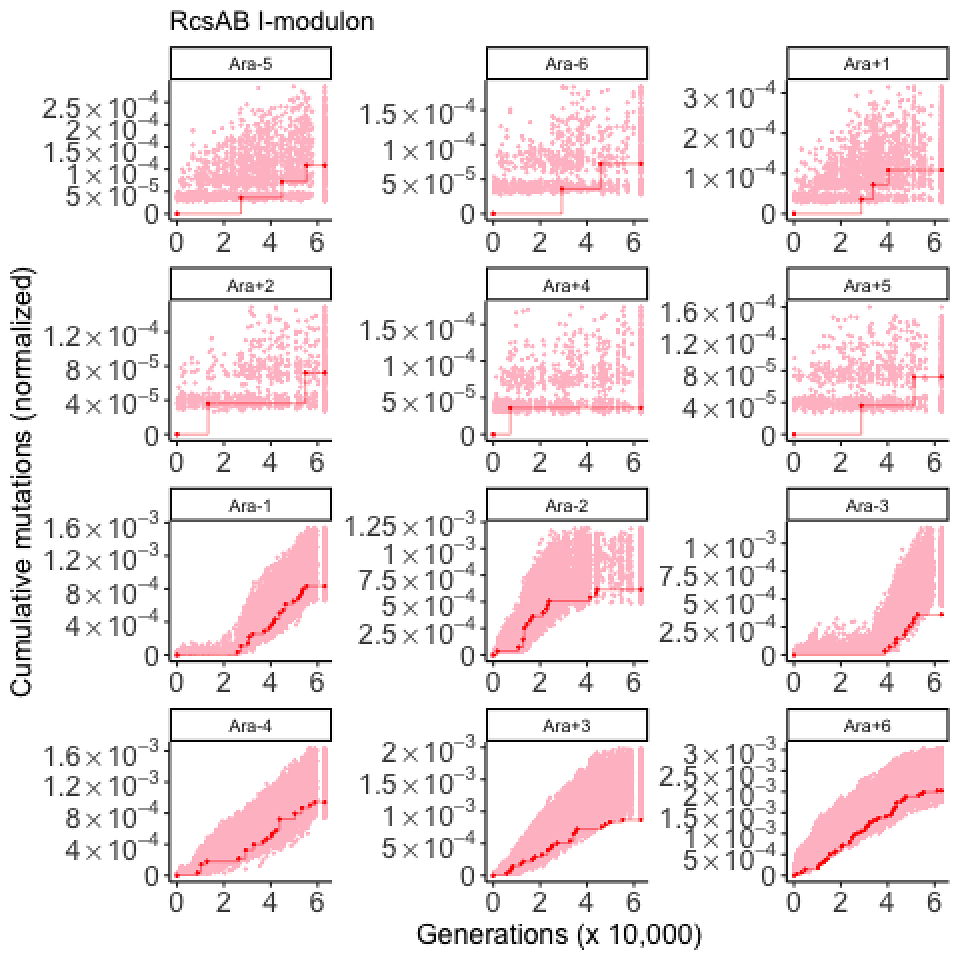

RcsAB I-modulon
Ara-5
Ara-6
Ara+1
Ara+2
Ara+4
Ara+5
Ara-1
Ara-2
Ara-3
Ara-4
Ara+3
Ara+6
Cumulative mutations (normalized)
Generations (x 10,000)

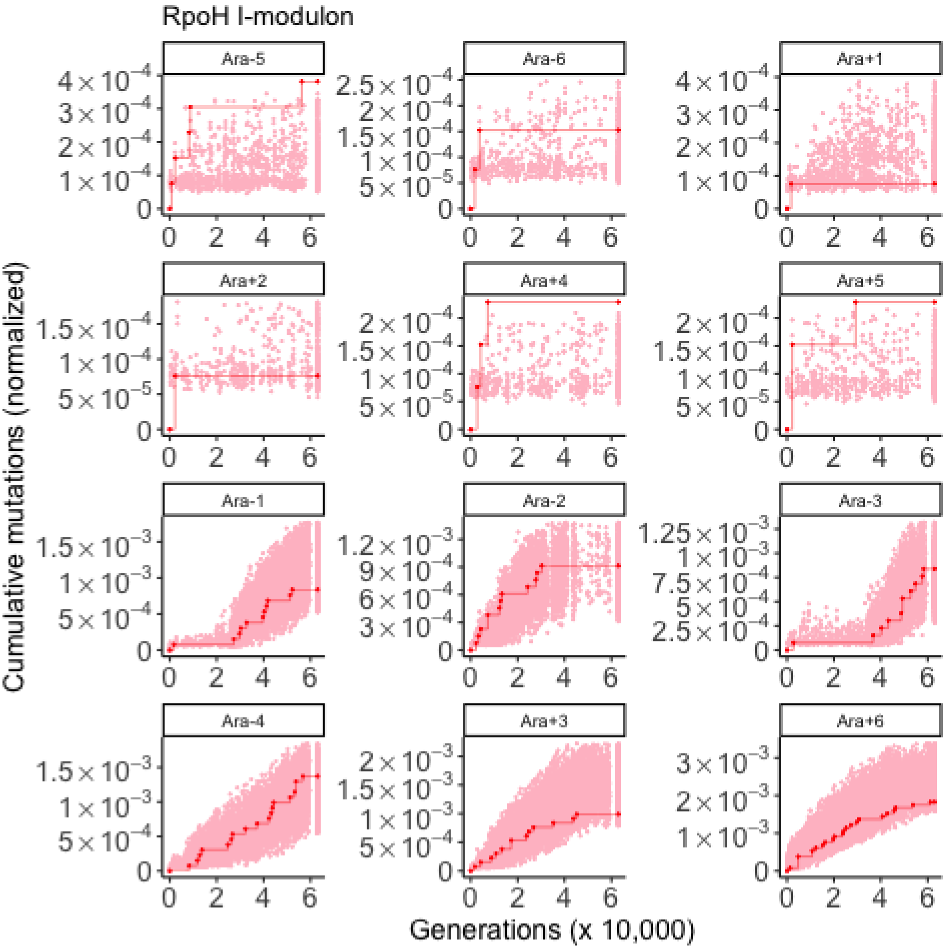

RpoH I-modulon
Ara-5
Ara-6
Ara+1
Ara+2
Ara+4
Ara+5
Ara-1
Ara-2
Ara-3
Ara-4
Ara+3
Ara+6
Cumulative mutations (normalized)
Generations (x 10,000)

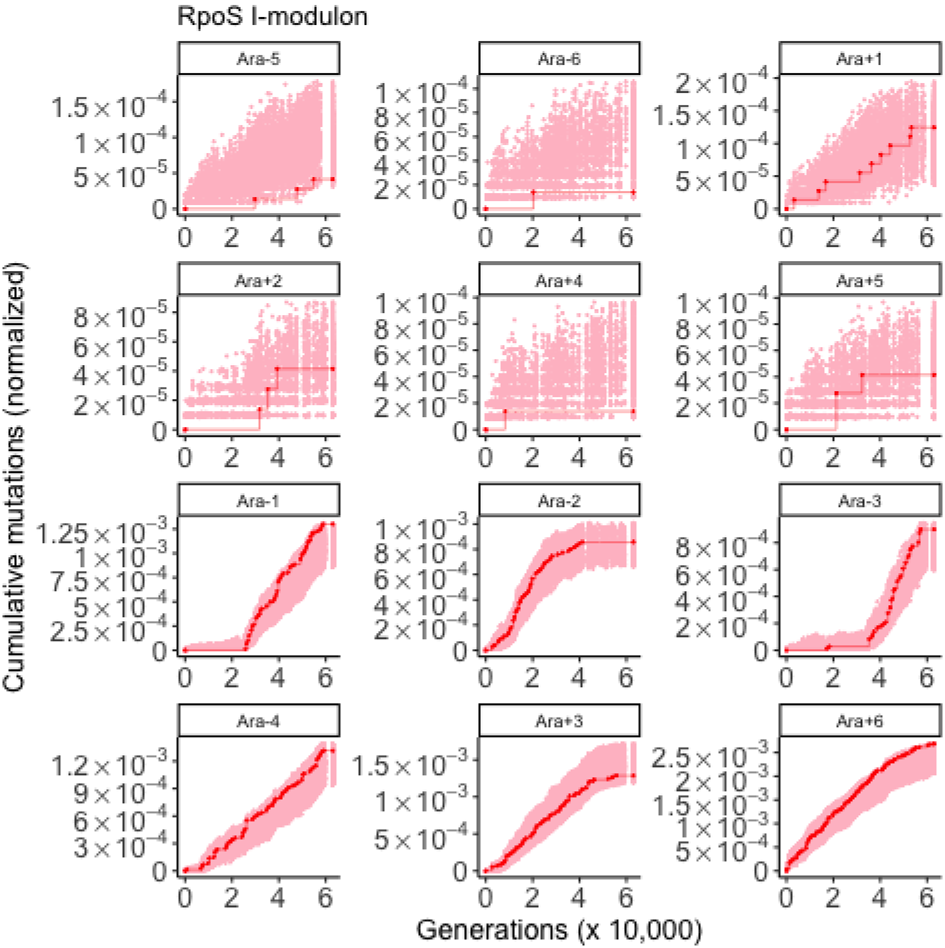

RpoS I-modulon
Ara-5
Ara-6
Ara+1
Ara+2
Ara+4
Ara+5
Ara-1
Ara-2
Ara-3
Ara-4
Ara+3
Ara+6
Cumulative mutations (normalized)
Generations (x 10,000)

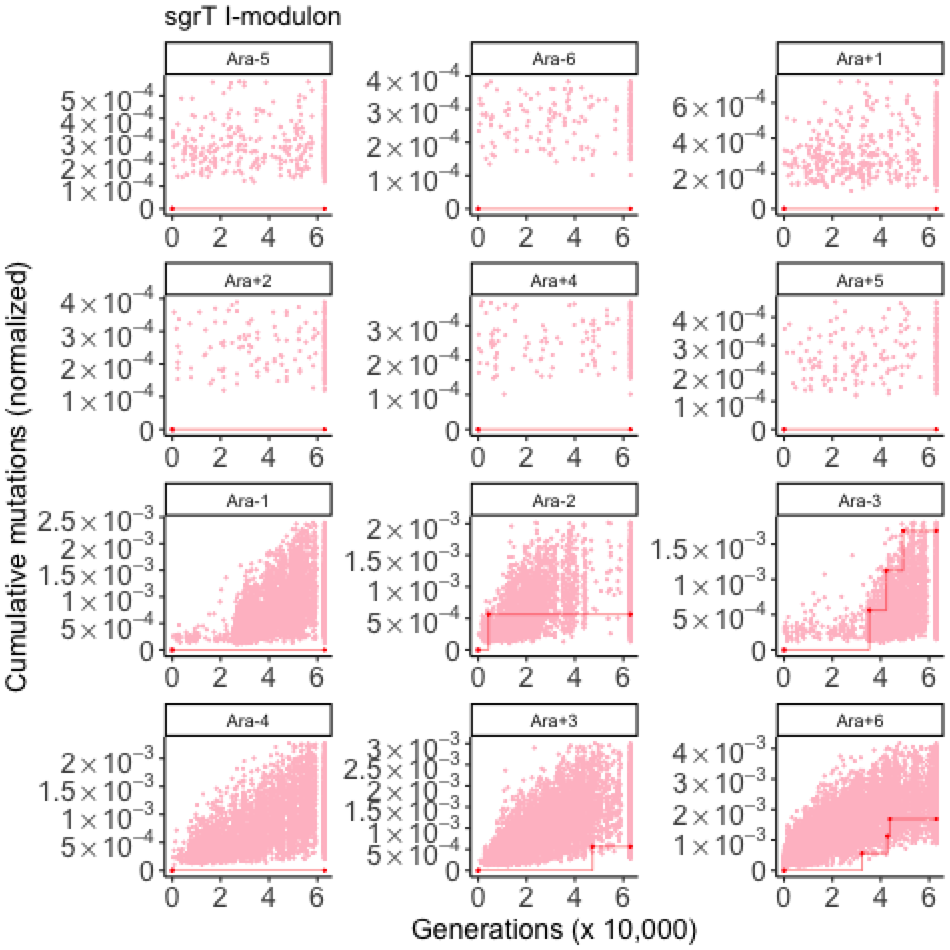
sgrT I-modulon
Ara-5
Ara-6
Ara+1
Ara+2
Ara+4
Ara+5
Ara-1
Ara-2
Ara-3
Ara-4
Ara+3
Ara+6
Cumulative mutations (normalized)
Generations (x 10,000)

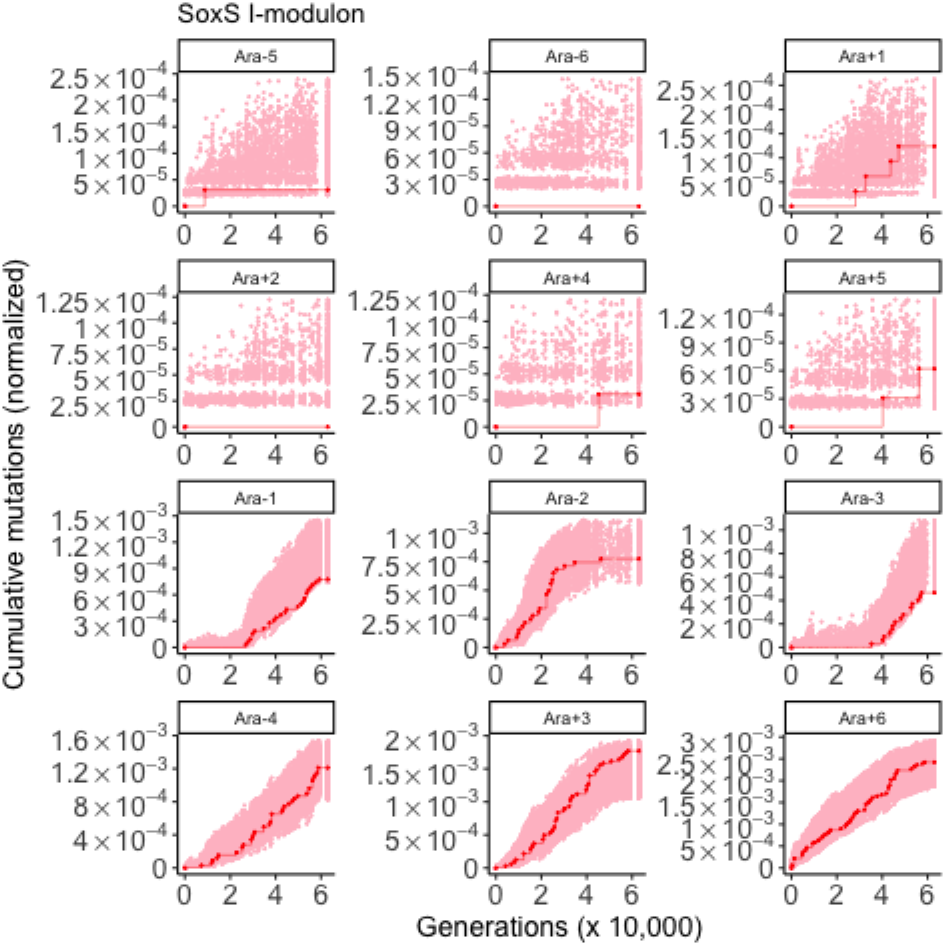

SoxS I-modulon
Ara-5
Ara-6
Ara+1
Ara+2
Ara+4
Ara+5
Ara-1
Ara-2
Ara-3
Ara-4
Ara+3
Ara+6
Cumulative mutations (normalized)
Generations (x 10,000)

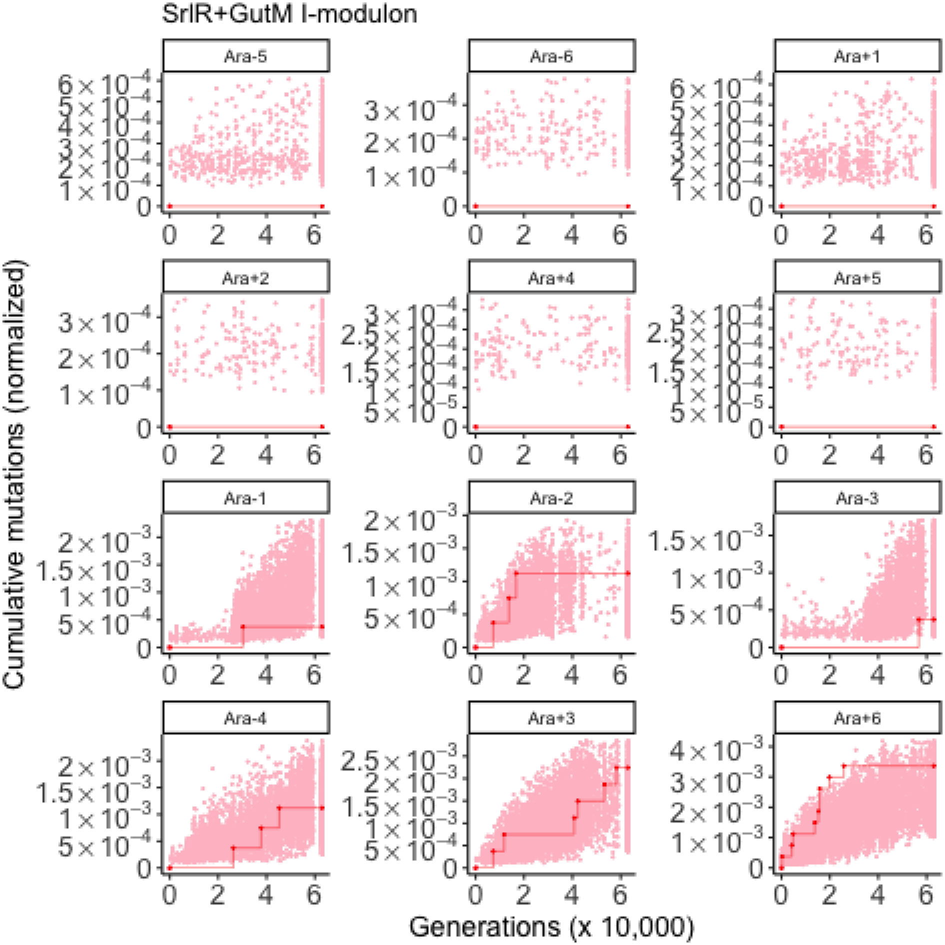

SrlR+GutM I-modulon
Ara-5
Ara-6
Ara+1
Ara+2
Ara+4
Ara+5
Ara-1
Ara-2
Ara-3
Ara-4
Ara+3
Ara+6
Cumulative mutations (normalized)
Generations (x 10,000)

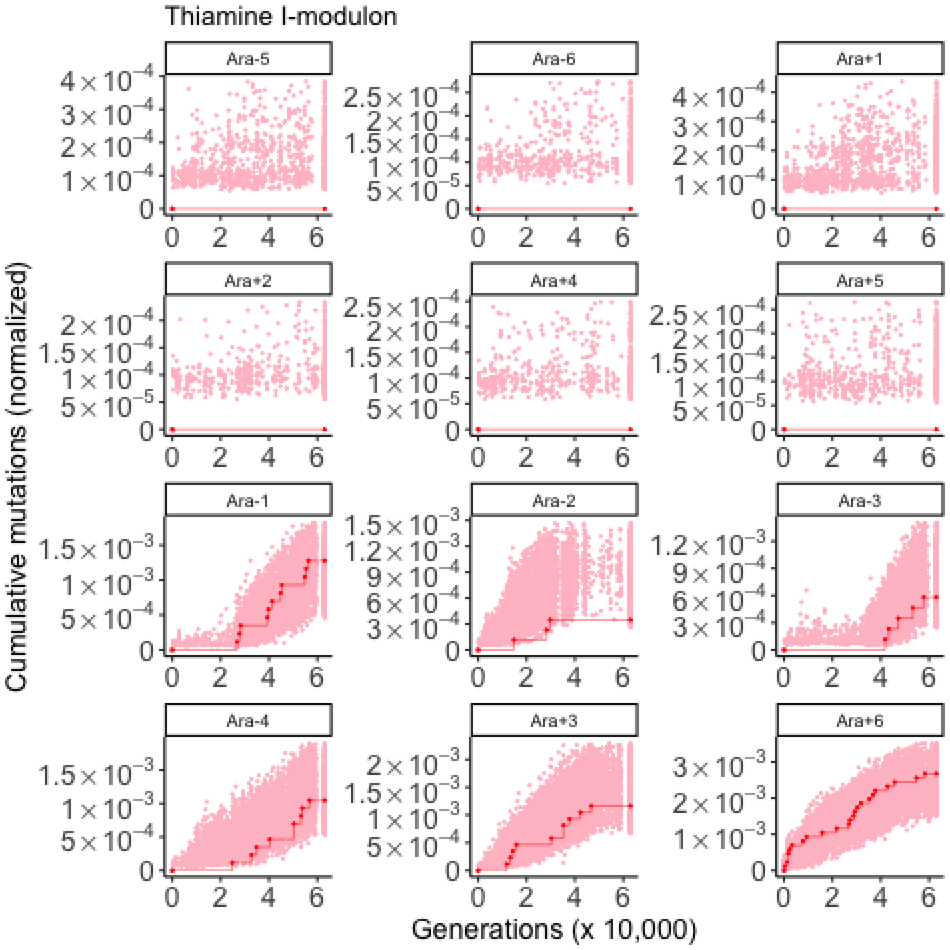

Thiamine I-modulon
Ara-5
Ara-6
Ara+1
Ara+2
Ara+4
Ara+5
Ara-1
Ara-2
Ara-3
Ara-4
Ara+3
Ara+6
Cumulative mutations (normalized)
Generations (x 10,000)

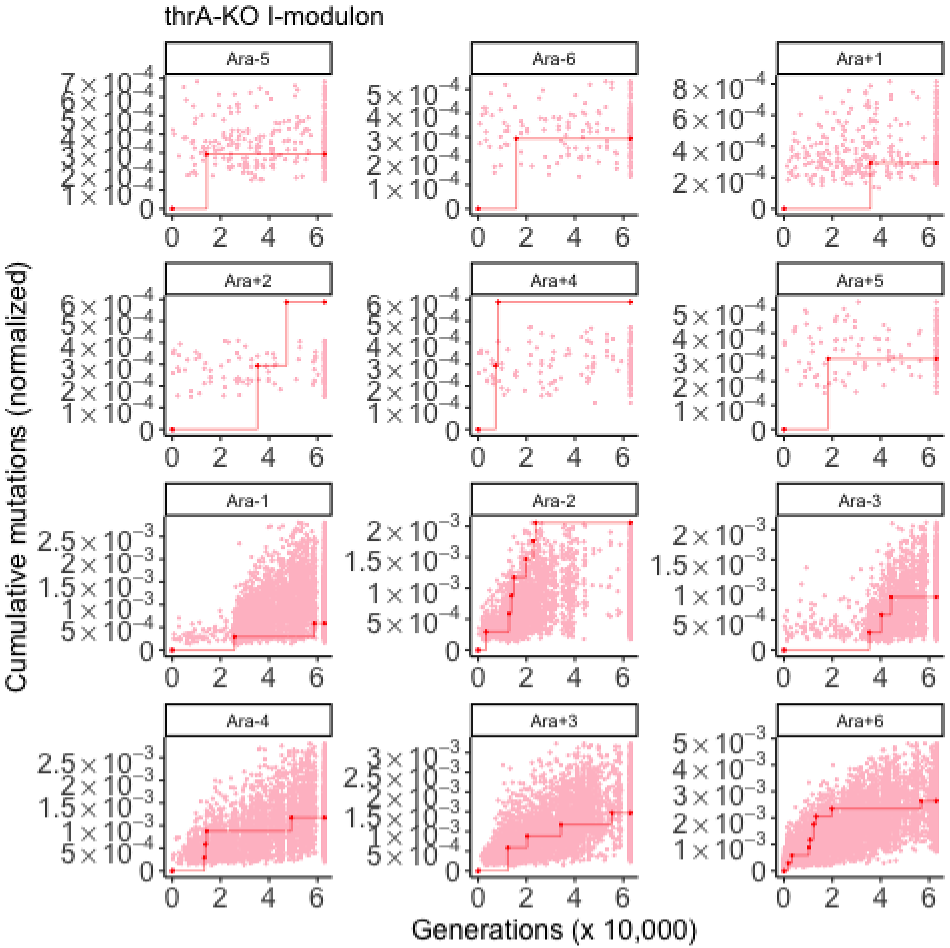
thrA-KO I-modulon
Ara-5
Ara-6
Ara+1
Ara+2
Ara+4
Ara+5
Ara-1
Ara-2
Ara-3
Ara-4
Ara+3
Ara+6
Cumulative mutations (normalized)
Generations (x 10,000)

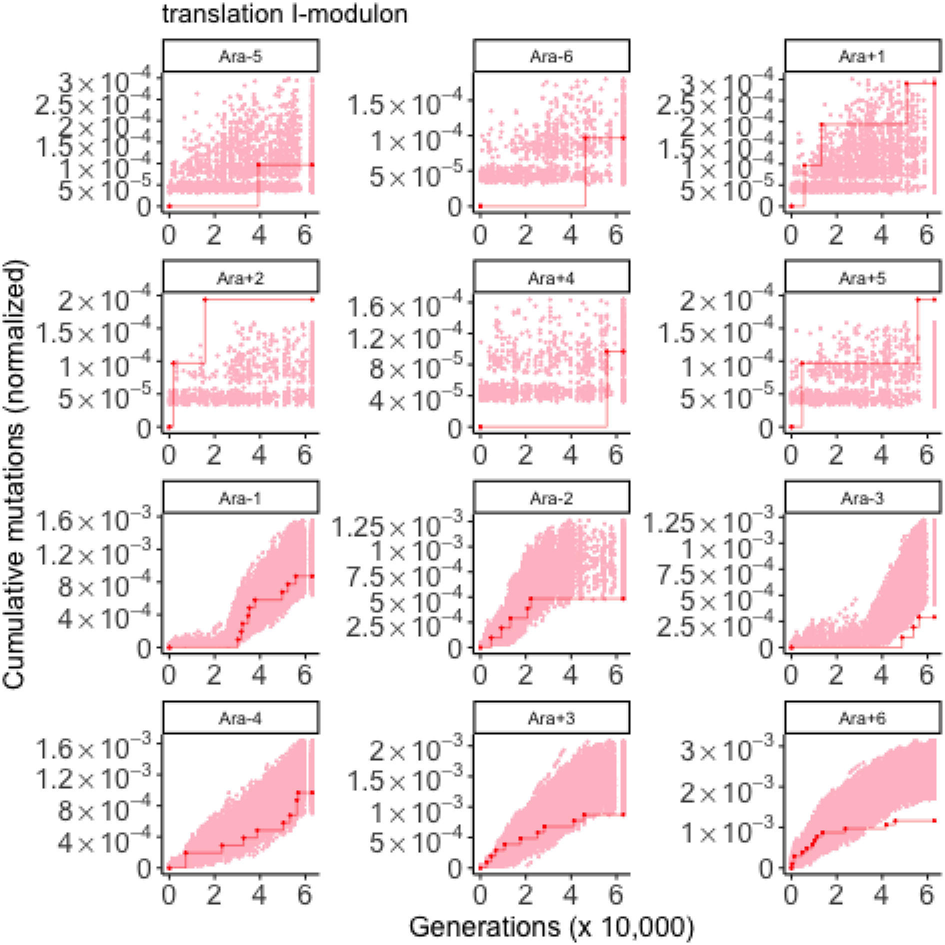

translation I-modulon
Ara-5
Ara-6
Ara+1
Ara+2
Ara+4
Ara+5
Ara-1
Ara-2
Ara-3
Ara-4
Ara+3
Ara+6
Cumulative mutations (normalized)
Generations (x 10,000)

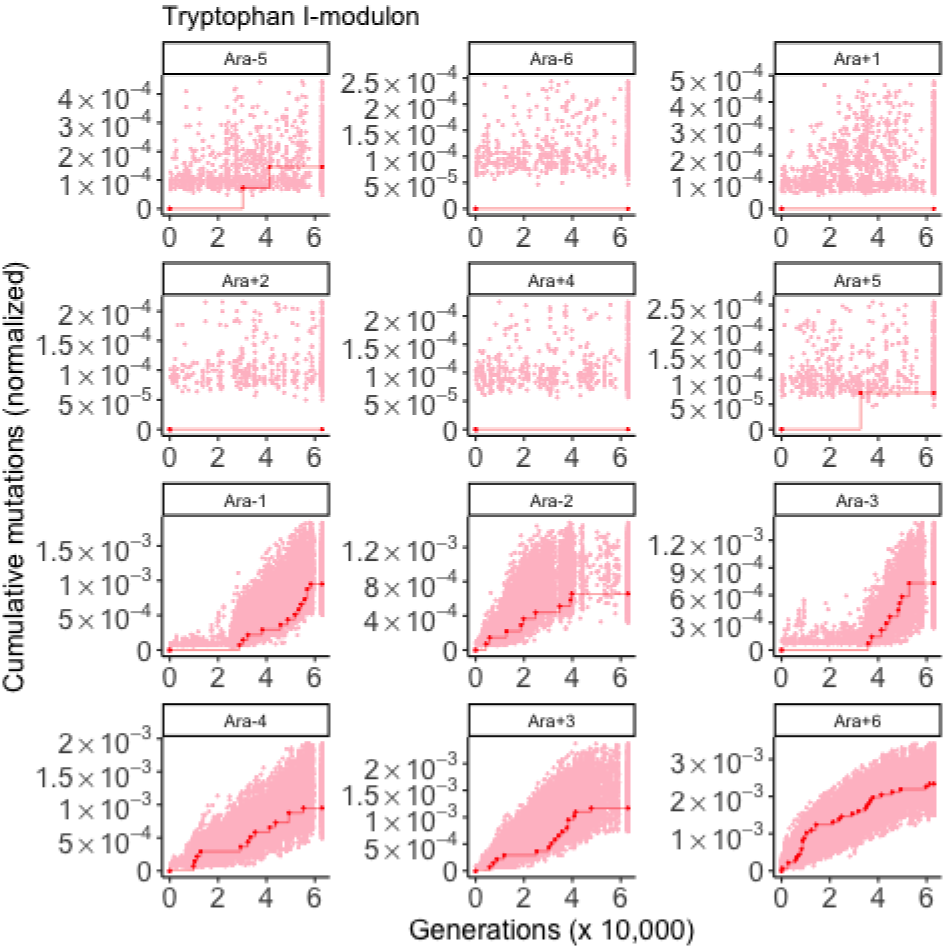
Tryptophan I-modulon
Ara-5
Ara-6
Ara+1
Ara+2
Ara+4
Ara+5
Ara-1
Ara-2
Ara-3
Ara-4
Ara+3
Ara+6
Cumulative mutations (normalized)
Generations (x 10,000)

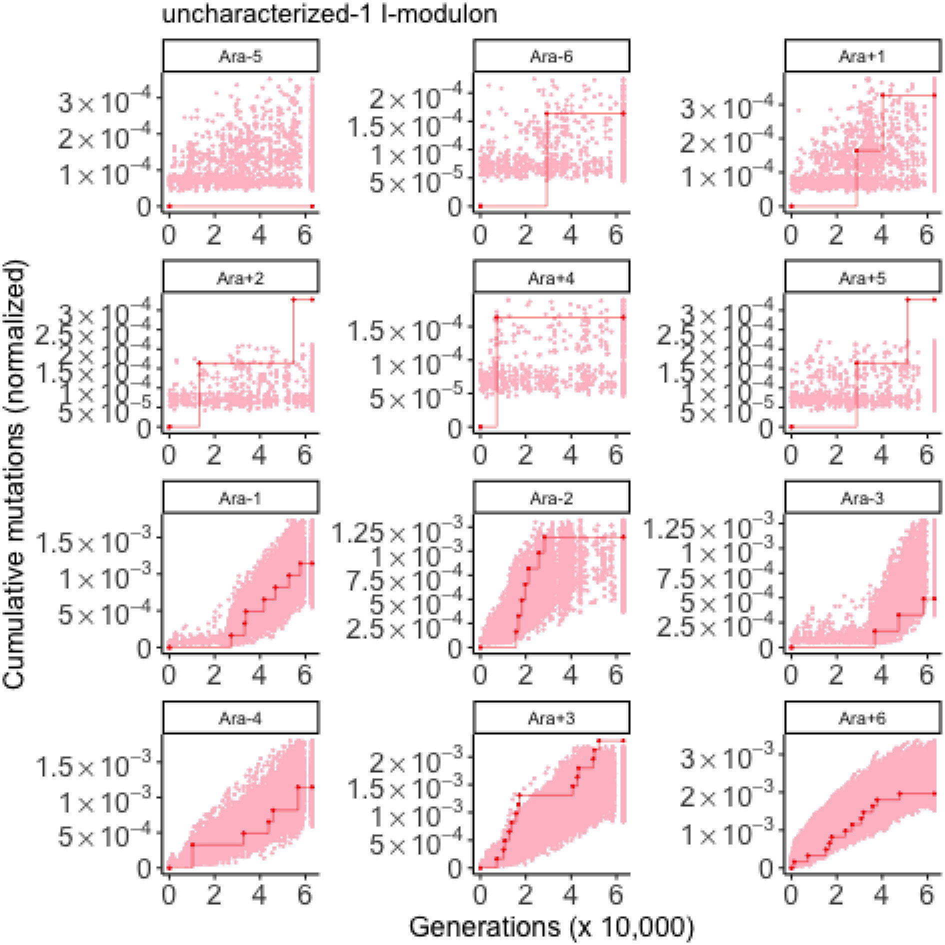
uncharacterized-1 I-modulon
Ara-5
Ara-6
Ara+1
Ara+2
Ara+4
Ara+5
Ara-1
Ara-2
Ara-3
Ara-4
Ara+3
Ara+6
Cumulative mutations (normalized)
Generations (x 10,000)

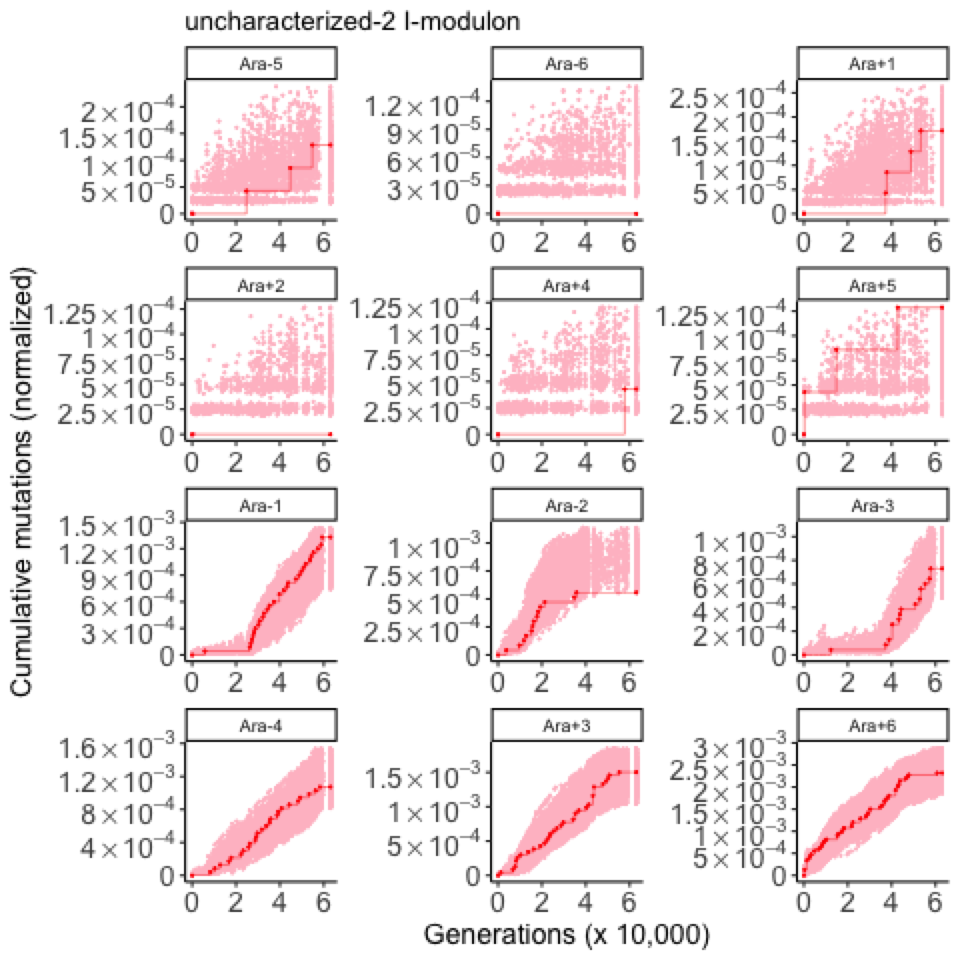

uncharacterized-2 I-modulon
Ara-5
Ara-6
Ara+1
Ara+2
Ara+4
Ara+5
Ara-1
Ara-2
Ara-3
Ara-4
Ara+3
Ara+6
Cumulative mutations (normalized)
Generations (x 10,000)

uncharacterized-3 I-modulon

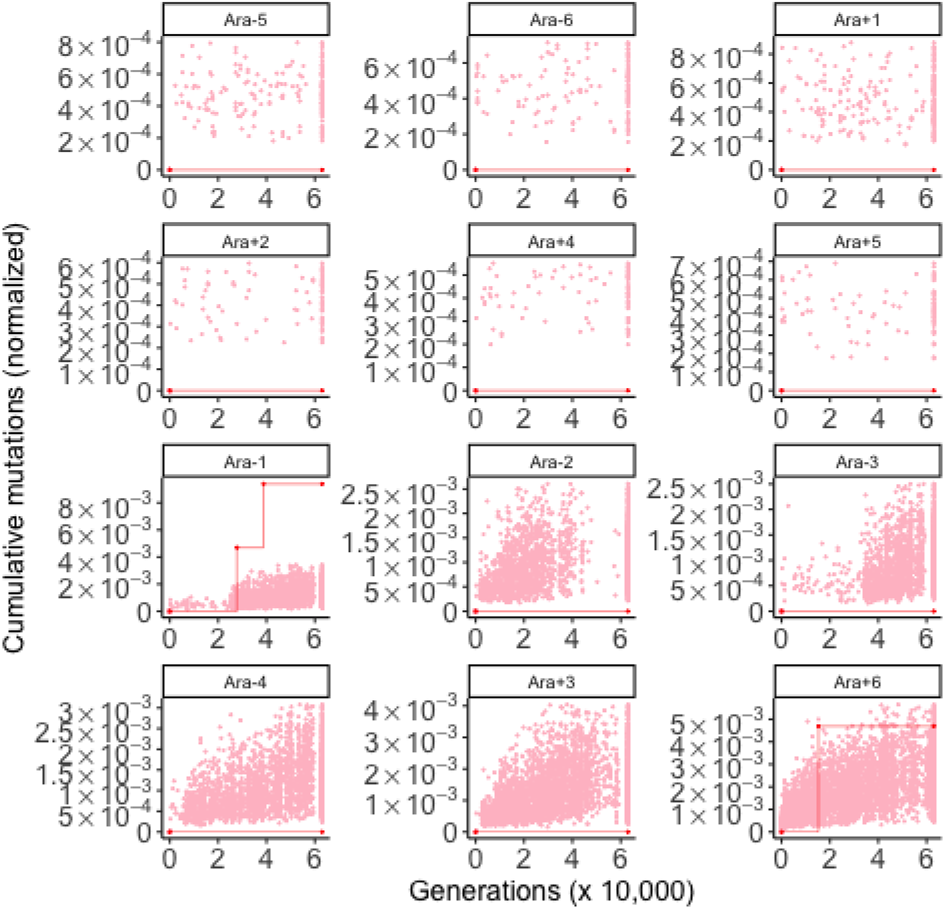

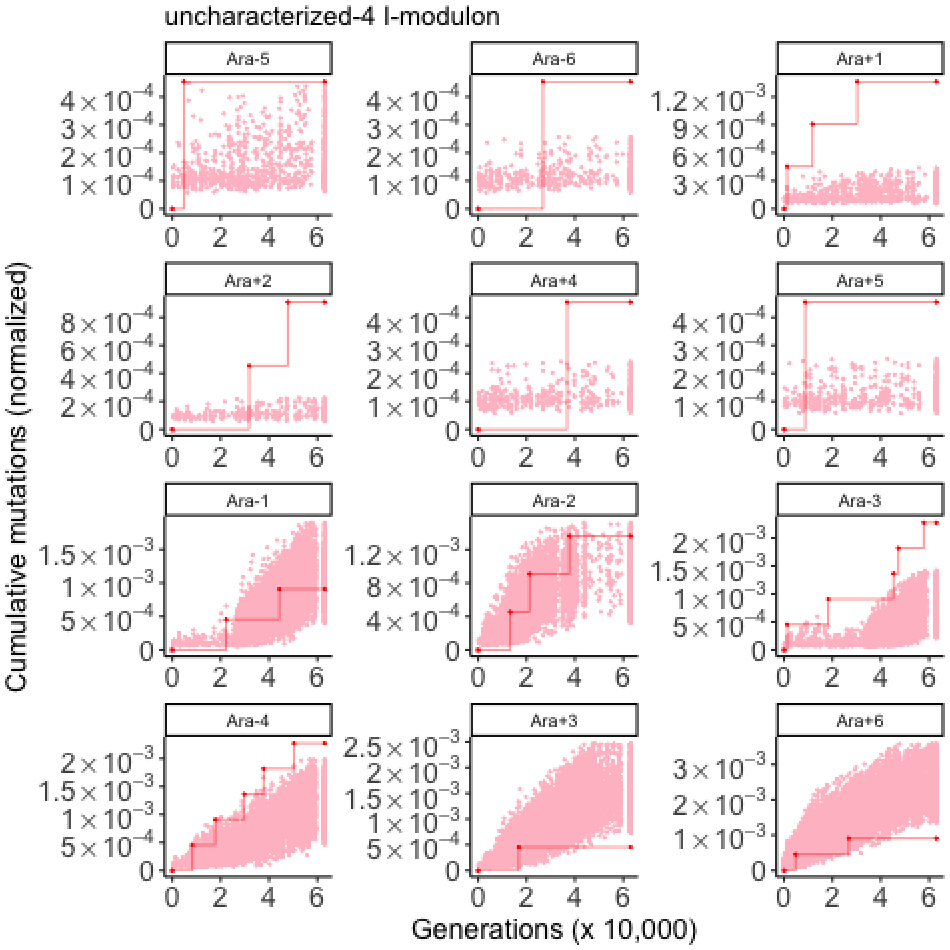
uncharacterized-4 I-modulon
Ara-5
Ara-6
Ara+1
Ara+2
Ara+4
Ara+5
Ara-1
Ara-2
Ara-3
Ara-4
Ara+3
Ara+6
Cumulative mutations (normalized)
Generations (x 10,000)

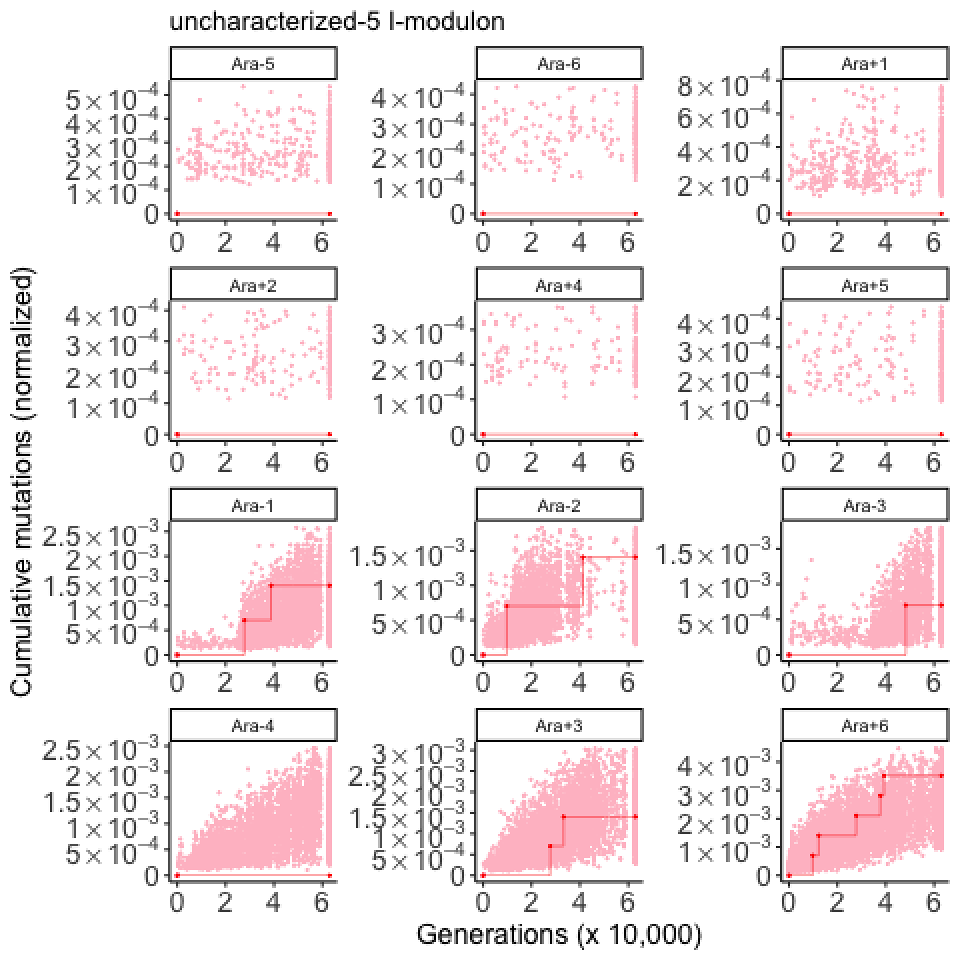

uncharacterized-5 I-modulon
Ara-5
Ara-6
Ara+1
Ara+2
Ara+4
Ara+5
Ara-1
Ara-2
Ara-3
Ara-4
Ara+3
Ara+6
Cumulative mutations (normalized)
Generations (x 10,000)

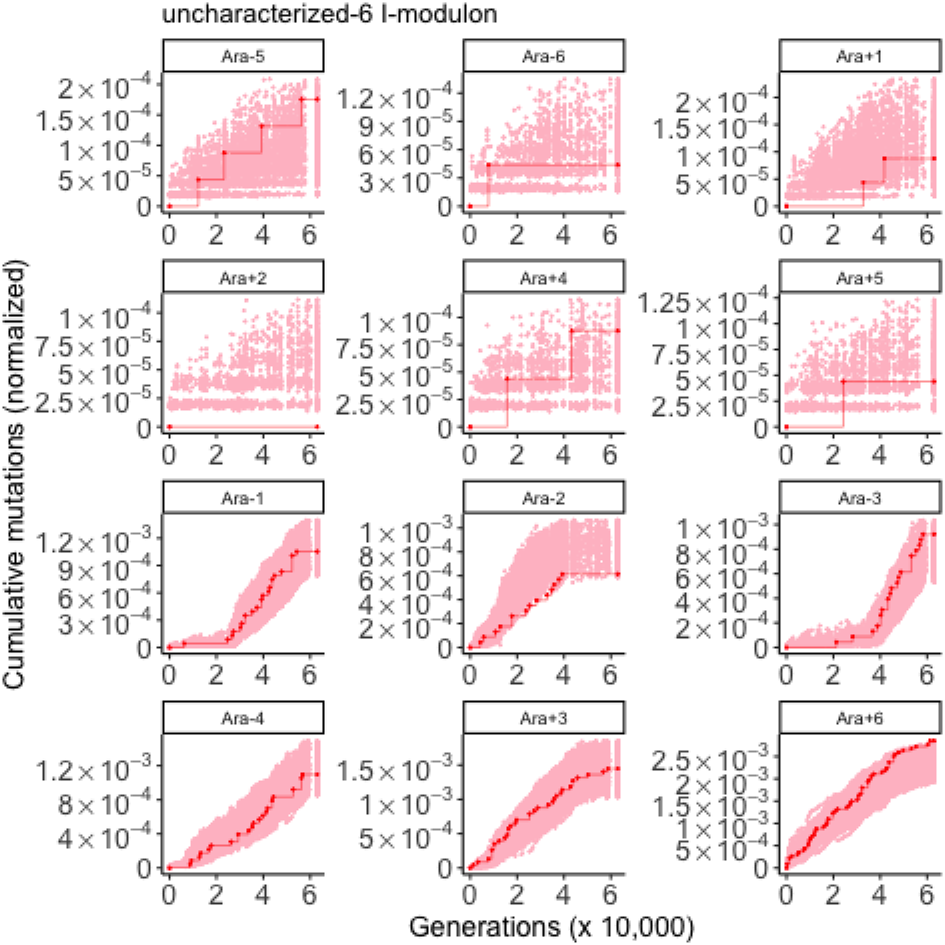

uncharacterized-6 I-modulon
Ara-5
Ara-6
Ara+1
Ara+2
Ara+4
Ara+5
Ara-1
Ara-2
Ara-3
Ara-4
Ara+3
Ara+6
Cumulative mutations (normalized)
Generations (x 10,000)

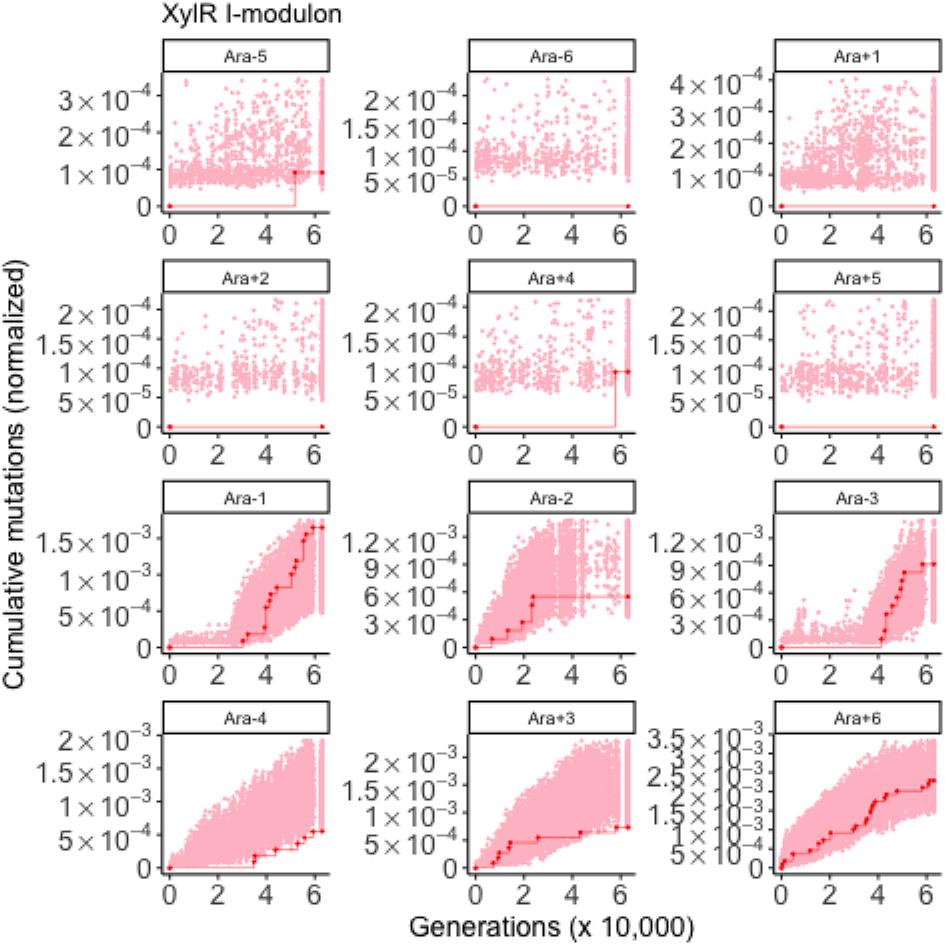

XylR I-modulon
Ara-5
Ara-6
Ara+1
Ara+2
Ara+4
Ara+5
Ara-1
Ara-2
Ara-3
Ara-4
Ara+3
Ara+6
Cumulative mutations (normalized)
Generations (x 10,000)

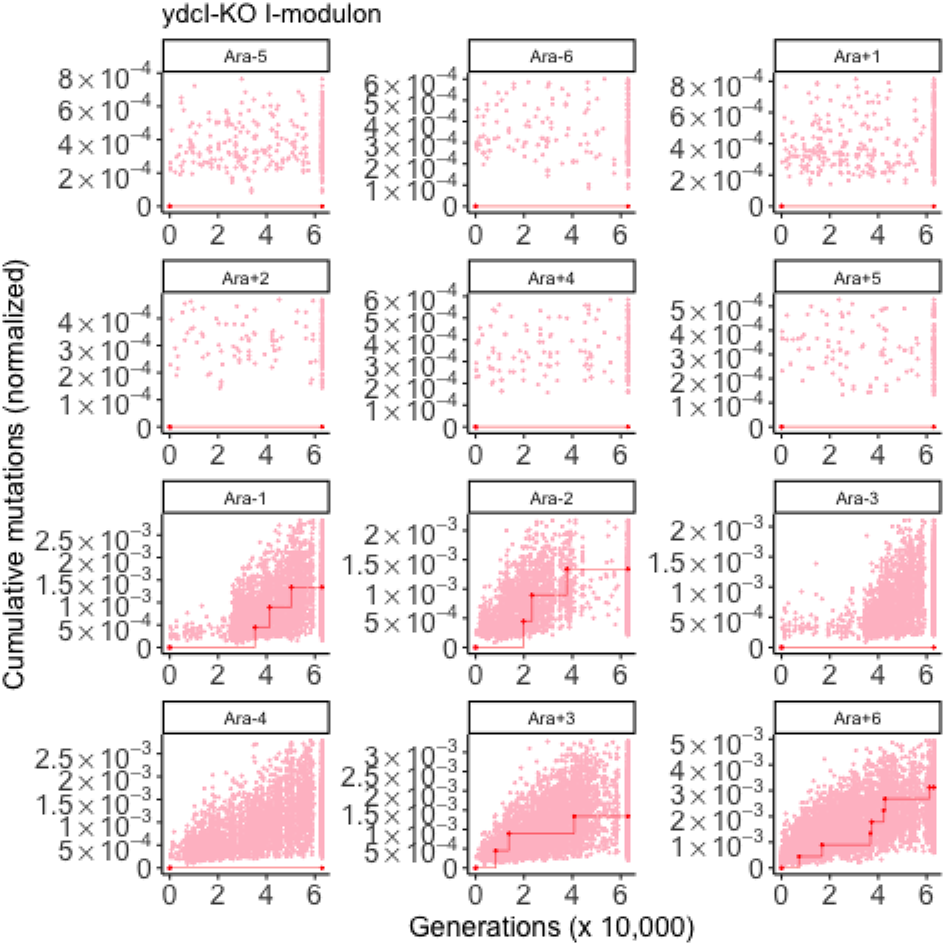

ydcI-KO I-modulon
Ara-5
Ara-6
Ara+1
Ara+2
Ara+4
Ara+5
Ara-1
Ara-2
Ara-3
Ara-4
Ara+3
Ara+6
Cumulative mutations (normalized)
Generations (x 10,000)

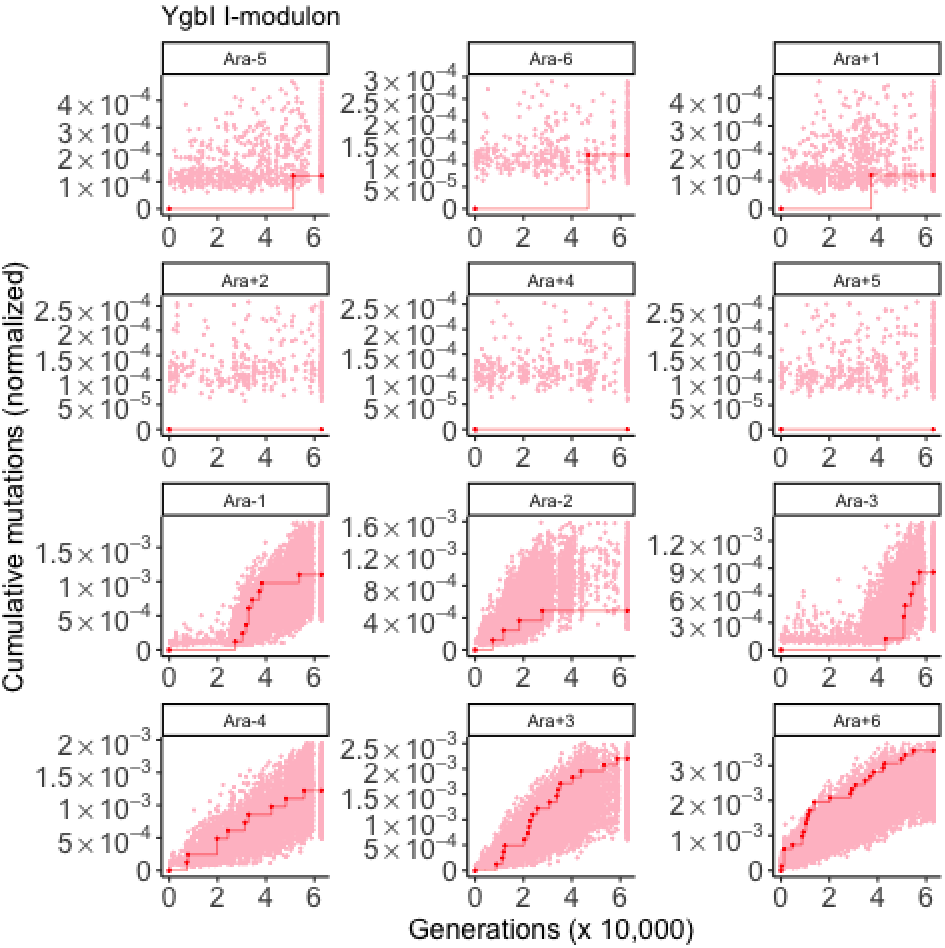
YgbI I-modulon
Ara-5
Ara-6
Ara+1
Ara+2
Ara+4
Ara+5
Ara-1
Ara-2
Ara-3
Ara-4
Ara+3
Ara+6
Cumulative mutations (normalized)
Generations (x 10,000)

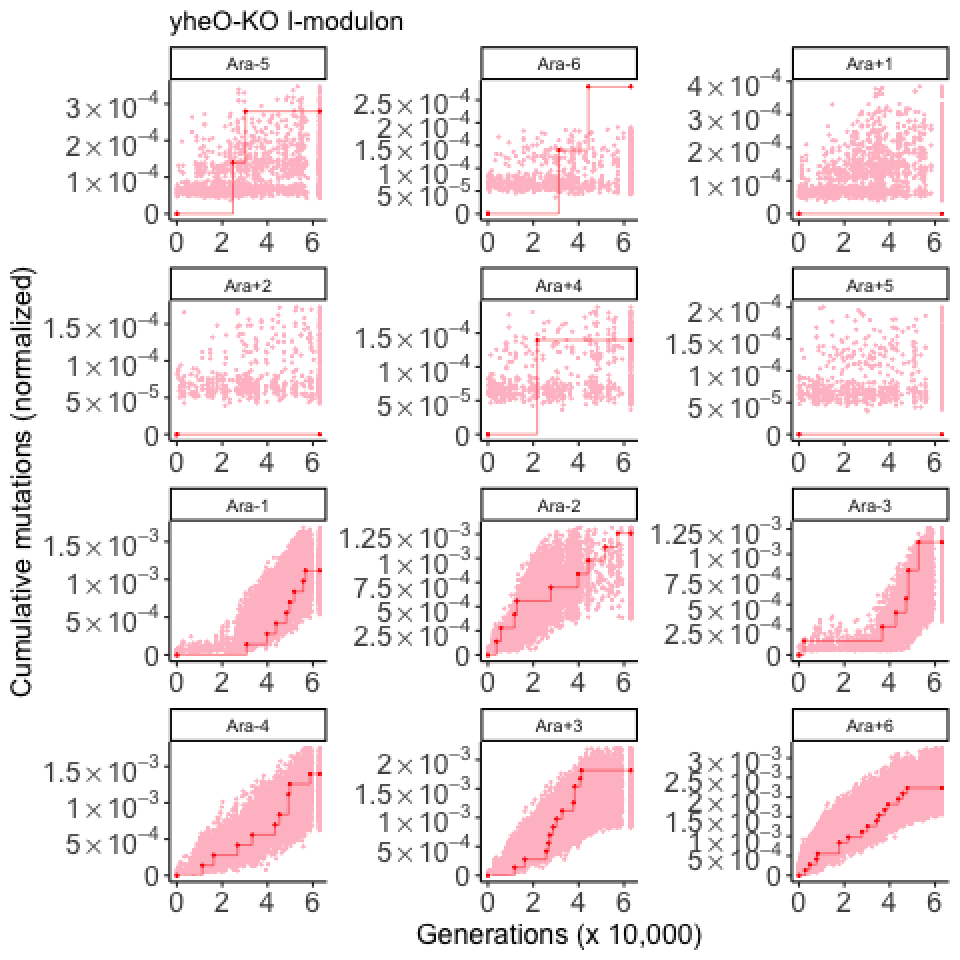
yheO-KO I-modulon
Ara-5
Ara-6
Ara+1
Ara+2
Ara+4
Ara+5
Ara-1
Ara-2
Ara-3
Ara-4
Ara+3
Ara+6
Cumulative mutations (normalized)
Generations (x 10,000)

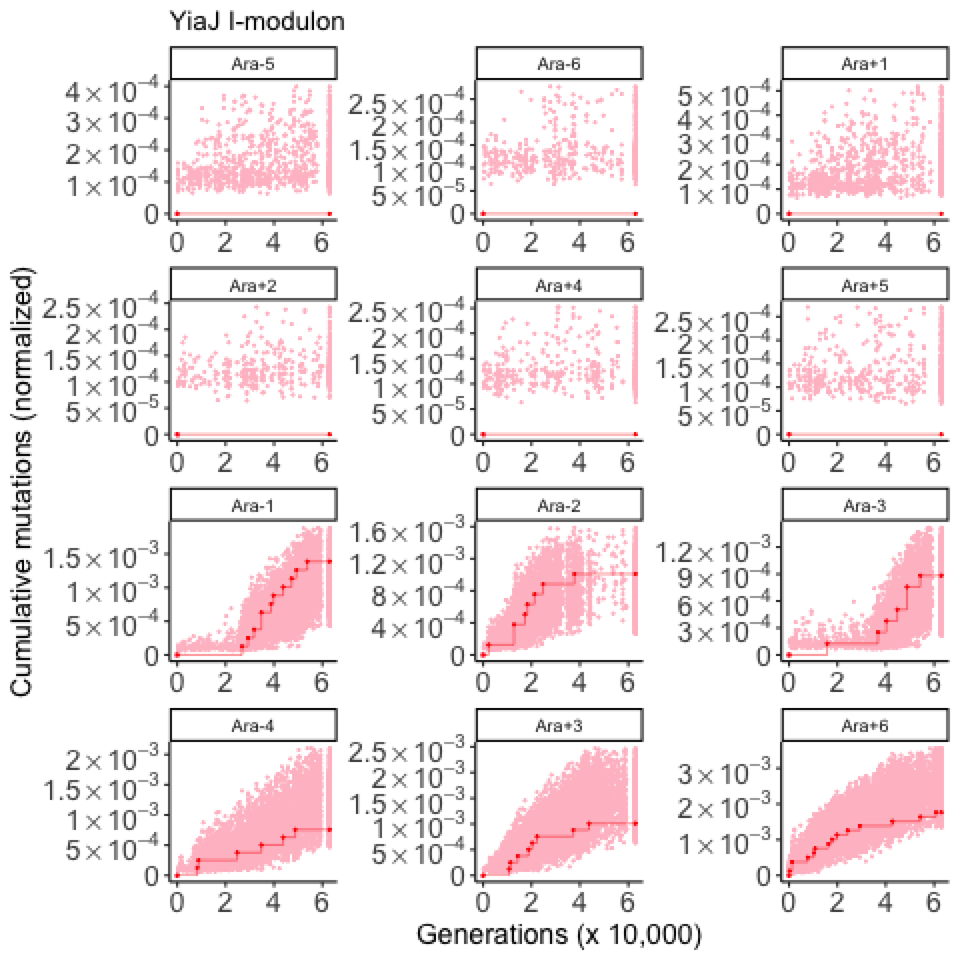

YiaJ I-modulon
Ara-5
Ara-6
Ara+1
Ara+2
Ara+4
Ara+5
Ara-1
Ara-2
Ara-3
Ara-4
Ara+3
Ara+6
Cumulative mutations (normalized)
Generations (x 10,000)

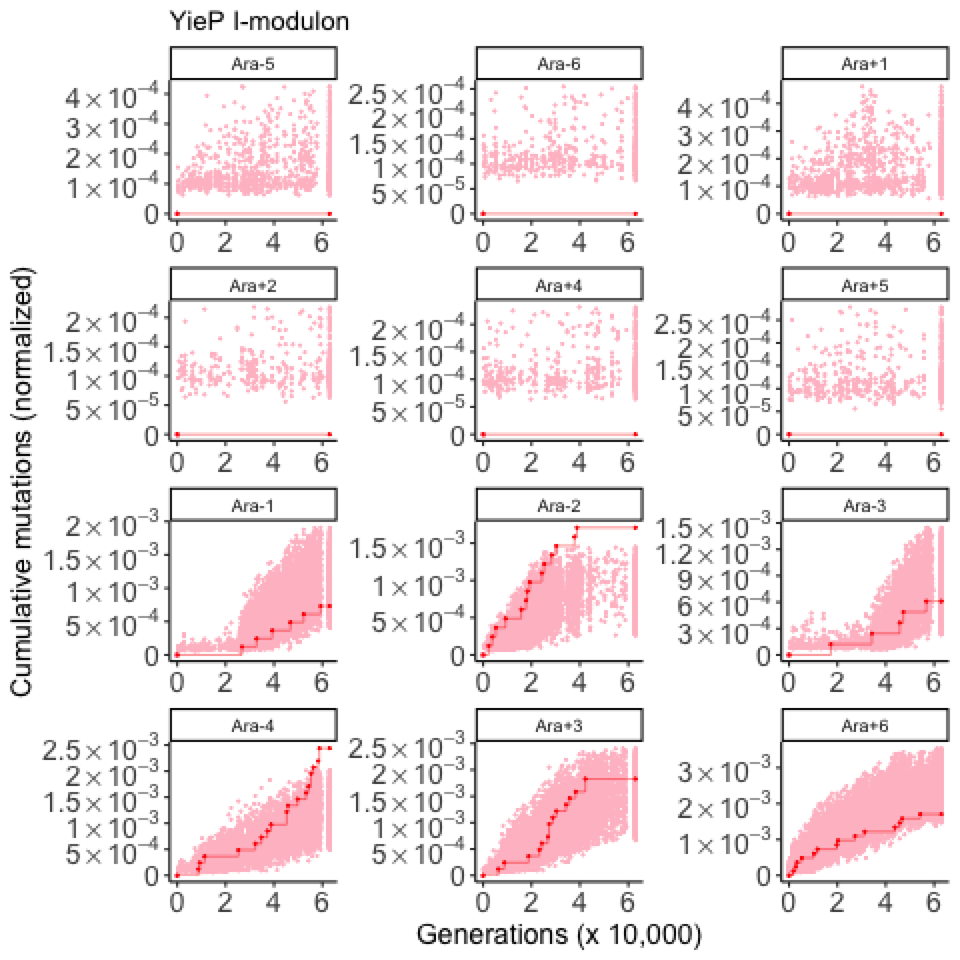

YieP I-modulon
Ara-5
Ara-6
Ara+1
Ara+2
Ara+4
Ara+5
Ara-1
Ara-2
Ara-3
Ara-4
Ara+3
Ara+6
Cumulative mutations (normalized)
Generations (x 10,000)

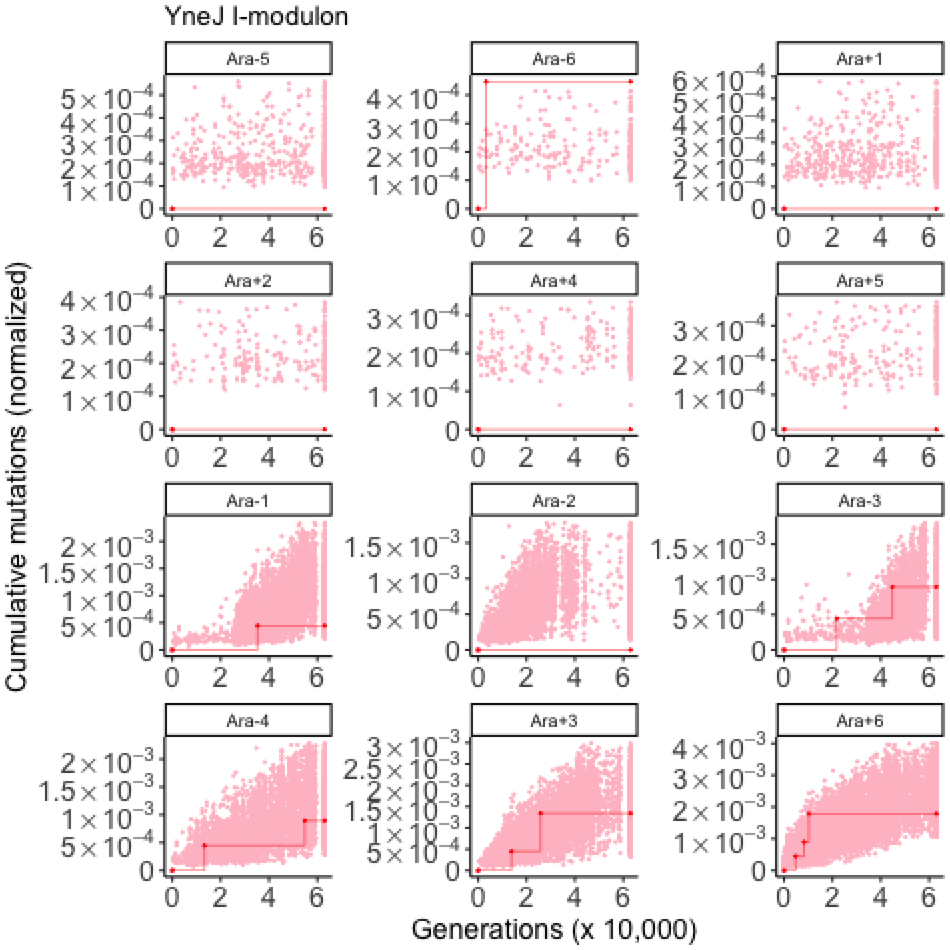

YneJ I-modulon
Ara-5
Ara-6
Ara+1
Ara+2
Ara+4
Ara+5
Ara-1
Ara-2
Ara-3
Ara-4
Ara+3
Ara+6
Cumulative mutations (normalized)
Generations (x 10,000)

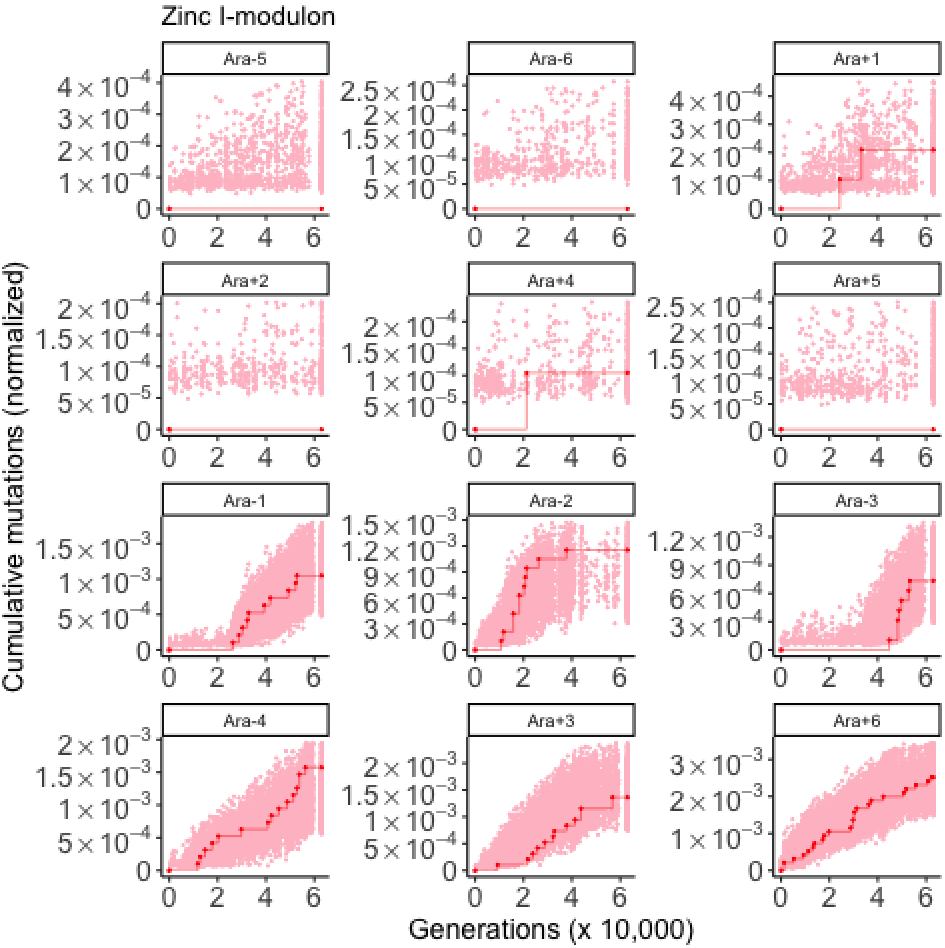
Zinc I-modulon
Ara-5
Ara-6
Ara+1
Ara+2
Ara+4
Ara+5
Ara-1
Ara-2
Ara-3
Ara-4
Ara+3
Ara+6
Cumulative mutations (normalized)
Generations (x 10,000)
